# Circle-Mediated HGT shapes the multichromosomal mitochondrial genome of the endoparasite *Mitrastemon yamamotoi*

**DOI:** 10.1101/2025.06.25.661533

**Authors:** Maria Emilia Roulet, Laura Evangelina Garcia, Runxian Yu, Chenyixin Wang, Renchao Zhou, M. Virginia Sanchez-Puerta

## Abstract

Horizontal gene transfer (HGT), a well-established driver of genome evolution in prokaryotes, was historically considered rare in plants. However, accumulating genomic evidence supports its occurrence in angiosperms, impacting both nuclear and mitochondrial genomes, particularly in parasitic species that establish vascular connections with their hosts. Despite the increasing recognition of HGT in a few clades of parasitic plants (e.g., Balanophoraceae, Rafflesiaceae, and Cynomoriaceae), the underlying mechanisms and evolutionary consequences of these transfers are still not fully understood. *Mitrastemon yamamotoi*, a holoparasitic endoparasite in the order Ericales, invades the roots of host trees in the Fagaceae family, creating favorable conditions for HGT. In this study, we assembled for the first time the mtDNA of *Mitrastemon*, revealing a multipartite structure consisting of 51 circular-mapping chromosomes. Phylogenetic and comparative genomic analyses uncovered extensive HGT from Fagaceae hosts, affecting both coding and non-coding regions. Notably, more than 60% of the *Mitrastemon* mtDNA is of foreign origin, and seven chromosomes are entirely foreign, with structural signatures in the donor mtDNA consistent with the recently proposed circle-mediated HGT model. Additionally, we detected six protein-coding genes of foreign origin and one chimeric gene. Remarkably, a foreign *atp1* gene has replaced the missing native copy and represents a rare event of functional HGT in plant mitochondria. These results position *Mitrastemon* as a valuable model for studying mtDNA evolution and deepening our understanding of the HGT process. Our findings expand the range of lineages in which circle-mediated HGT has been documented, suggesting it is a more widespread and fundamental mode of mitochondrial HGT in plants.

**Significant Statement:** Parasitic plants form intimate connections with their hosts, but how these interactions influence genome evolution remains poorly understood. Our study shows that *Mitrastemon yamamotoi* has acquired over 60% of its mitochondrial DNA from its host through horizontal gene transfer, including entire foreign chromosomes. These findings provide strong support for a recently proposed mechanism, circle-mediated HGT, and suggest that this process may be a more widespread driver of mitochondrial genome evolution in flowering plants.

## Introduction

Horizontal gene transfer (HGT), the movement of genetic material between different species by means other than sexual reproduction, was historically considered rare in plants (Richardson & Palmer, 2007). Over the past two decades, most documented cases of HGT in angiosperms have involved plant-to-plant transfers impacting the nuclear and mitochondrial genomes, especially in parasitic plants that form intimate connections with their hosts (Cusimano *et al*., 2008; Xi *et al*., 2013; Davis & Xi, 2015; Sanchez-Puerta *et al*., 2017; Roulet *et al*., 2020; Cai *et al*., 2021). This phenomenon has been frequently reported in parasitic plants with prolonged endophytic stages, such as species in the families Balanophoraceae, Rafflesiaceae, and Cynomoriaceae (Bellot *et al*., 2016; Cusimano & Wicke, 2016; Roulet *et al*., 2020; Sanchez-Puerta *et al*., 2023; Roulet *et al*., 2024).

Two complementary mechanisms have been proposed to explain plant mitochondrial HGT. The first is the fusion-compatibility model, which posits that HGT occurs via the acquisition of entire foreign mitochondria from species in the green lineage, subsequently capable of fusing with the native mitochondria (Rice *et al*., 2013). Several lines of evidence support this model, including the horizontal transfer of large tracts (or even entire) mitochondrial genomes (mtDNAs), but not nuclear or plastid DNA, into both free-living and parasitic plant mitochondria (Rice *et al*., 2013; Sanchez-Puerta *et al*., 2019). The second proposed mechanism is the circle-mediated HGT, in which foreign mitochondrial DNA enters the mitochondria of a recipient plant but remains unintegrated, and forms subgenomic, circular DNA molecules (Roulet *et al*., 2024). These circular molecules are primarily formed through microhomology-mediated recombination across direct repeats within the donor mtDNA. Once formed, they can replicate autonomously and persist until they are either lost or eventually recombine with the native mtDNA. The circle-mediated HGT model has been described in two species of parasitic plants that suffered extraordinary and continuous horizontal transfers of mitochondrial DNA from their host plants (Roulet *et al*., 2024; Gatica-Soria *et al*., 2025). It is unclear how widespread this model is for angiosperm mitochondrial HGT and which conditions are relevant to set the stage for the circularization process. Examining diverse plants with high levels of HGT is instrumental to evaluate this model.

The holoparasitic endoparasite *Mitrastemon yamamotoi* (Ericales) offers a unique and compelling system to investigate mitochondrial HGT, and represents the first study describing the mitochondrial genome of this parasitic lineage. It completely lacks roots, stems, and leaves, remaining concealed within the root tissues of its host, typically *Castanopsis* species (Fagaceae), from which it extracts water, nutrients, and metabolites via direct vascular connections (Teixeira-Costa et al., 2023). Its vegetative body consists of a parenchymatous endophyte that extends into both the phloem and xylem of the host plant, and only emerges externally during flowering, which may occur years after the initial infection (Watanabe, 1933). This prolonged endophytic phase facilitates continuous acquisition of host DNA, which upon flowering can be passed onto the offspring, a key step in the establishment and maintenance of HGT events across generations.

In this study, we aimed to characterize the mtDNA of *Mitrastemon* to evaluate its organization, gene content, and the extent of HGT. Specifically, we investigated the presence and distribution of foreign mitochondrial DNA and whether there is evidence of circle-mediated HGT. For this, we assembled the mtDNA of *Mitrastemon* and of 16 species of *Castanopsis*, which provided the necessary comparative genomics data to uncover a multichromosomal mtDNA in *Mitrastemon*, with large tracts of host-derived mitochondrial DNA. Finally, we examined the transcription and RNA editing of foreign genes and identified in the transcriptome of *Mitrastemon* the expression of genes involved in the processes of replication, repair, and recombination of the mitochondrial DNA.

## Results

### Overview of the mitochondrial genome of *Mitrastemon yamamotoi*

The assembly of the *Mitrastemon yamamotoi* mitochondrial genome (mtDNA), from an individual parasitizing *Castanopsis* sp. roots, revealed a highly fragmented, multipartite structure comprising 51 circular-mapping chromosomes ranging in size from 9 to 37 kb, with a total length of 929,557 bp and a GC content of 45% (Figure 1 and Table 1). Direct and inverted repeats constitute 17% of the mtDNA (159,368 bp) (Table S1 and Table 1). The mtDNA exhibits even read coverage, without gaps or low-coverage regions and with an average read depth of 700x, varying between 600x and 900x across the 51 mitochondrial chromosomes (Figure S1).

**Figure 1.**
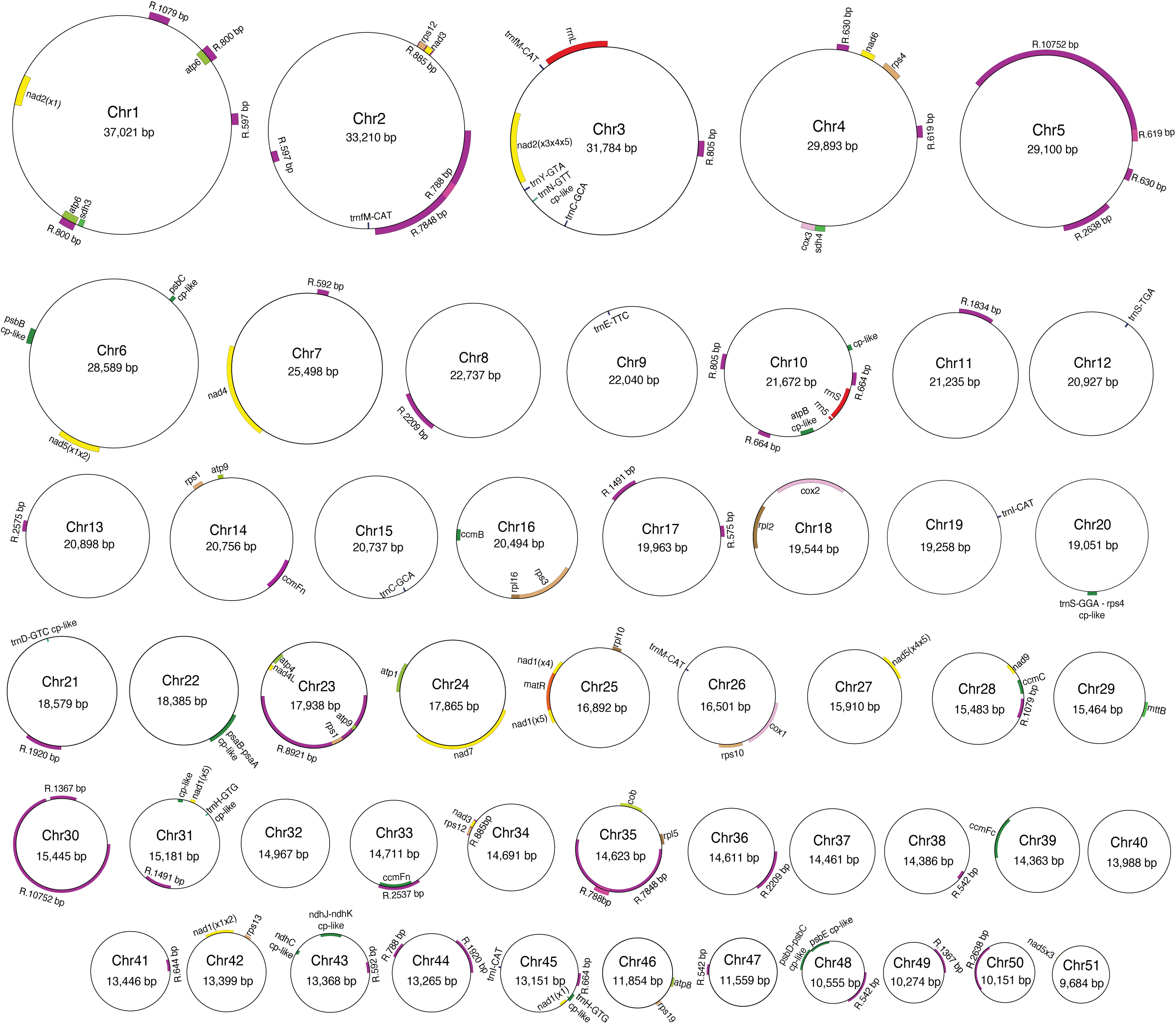
Map of the mitochondrial genome (mtDNA) of *Mitrastemon yamamotoi*. The mtDNA is 929,557 bp long and is subdivided in 51 circular chromosomes (Chr) of diverse lengths. Genes drawn inside and outside of each circle are transcribed clockwise and counterclockwise, respectively. Shown are full-length genes, repeats >500 bp (labeled “R” followed by the length), and sequences derived from the chloroplast (cp-like) longer than 200 bp.

**Table 1.** General features of the mitochondrial genome of *Mitrastemon yamamotoi*.

|  |  |
| --- | --- |
| Genome length (bp) | 929,557 bp |
| GC content | 45.13% |
| Protein-coding genes (including repeats) | 37 (43) |
| rRNA-coding genes | 3 |
| tRNA-coding genes | 12 (14) |
| Pseudogenes | 11 |
| Group II introns |  |
| cis-splicing | 19 |
| trans-splicing | 5 |
| Group I introns | 1 |
| Repeats* (bp; % of total repeats) | 159,368 bp; 17% |
| Chloroplast derived sequences (# MTPTs) | 11 |
| Chloroplast derived sequences (bp, % of genome) | 8,340 bp; 0.9% |
| Mitochondrial-like sequences (including genes) | 817,087 bp; 88.76% |

Among the 51 chromosomes, only 31 contain mitochondrial genes, whereas the remaining 20 lack any full-length gene (Figure 1). The coding portion of the mtDNA constitutes 7.5%, encompassing three ribosomal RNAs (rRNAs), 14 transfer RNAs (tRNAs), and 37 protein-coding genes. Some of these genes occur in multiple copies, bringing the total protein gene count to 43 (Table 1, Table S2). The mtDNA also contains 19 *cis*-spliced and five *trans*-spliced introns, alongside a single group I intron in the *cox1* gene (Table 1). Five of the tRNAs (*trnN*, *trnD*, *trnM*, and two copies of *trnH*) are of plastid origin (Table S2). We identified 11 mitochondrial pseudogenes that were truncated or frameshifted (Table S2) and 11 chloroplast-derived sequences representing 1% of the genome (Table S3).

### Foreign origin of the *Mitrastemon yamamotoi* mitochondrial protein coding genes

To assess the phylogenetic origin of the protein-coding genes in *Mitrastemon* we performed Maximum Likelihood phylogenetic analyses and evaluated their chimeric origin using GeneConv. Phylogenetic trees of each protein gene revealed that, out of the 43 protein-coding genes, six are of foreign origin (*atp1, atp6-1, atp6-2, ccmFn-1, nad3-1, rps12-1*), one is chimeric (*nad4*), and the remaining 36 are classified as native (Table 2, Figure S2). Genes associated with Ericales, as well as those with an unsupported position, were categorized as native. However, there were several cases (e.g. *atp4*, *cox2*, *nad4L*, *nad9*) in which the gene of *Mitrastemon* clusters with Fagales but with low to moderate bootstrap and were conservatively considered native.

**Table 2.** Features of protein-coding genes in the *Mitrastemon yamamotoi* mtDNA.

| Gene name † | Mitochondrial chromosome | Gene Start | Gene End | CDS length (gene length) in bp | Phylogenetic origin | AU test |
| --- | --- | --- | --- | --- | --- | --- |
| *atp1* | Chr24 | 7350 | 8882 | 1533 | Foreign | $p < 0.05$ |
| *atp4* | Chr23 | 7025 | 6384 | 642 | Native | na |
| *atp6_1* | Chr01 | 4465 | 3668 | 798 | Foreign | ns |
| *atp6_2* | Chr01 | 25183 | 24332 | 852 | Foreign | ns |
| *atp8* | Chr46 | 11163 | 11663 | 501 | Native | na |
| *atp9_1* | Chr14 | 5628 | 5852 | 225 | Native | na |
| *atp9_2* | Chr23 | 15915 | 16136 | 225 | Native | na |
| *ccmB* | Chr16 | 10386 | 9766 | 621 | Native | na |
| *ccmC* | Chr28 | 1057 | 314 | 744 | Native | na |
| *ccmFC (1)* | Chr39 | 7618 | 5292 | 1338 (2327) | Native | na |
| *ccmFN_1* | Chr14 | 17829 | 19559 | 1731 | Foreign | $p < 0.05$ |
| *ccmFN_2* | Chr33 | 12038 | 10311 | 1728 | Native | na |
| *cob* | Chr35 | 2382 | 3570 | 1189 | Native | na |
| *cox1* | Chr26 | 13571 | 16126 | 1584 | Native | na |
| *cox2 (2)* | Chr18 | 6856 | 2955 | 768 (3902) | Native | na |
| *cox3* | Chr04 | 20987 | 21784 | 798 | Native | na |
| *matR* | Chr25 | 7134 | 9095 | 1962 | Native | na |
| *mttB* | Chr29 | 14463 | 15251 | 789 | Native | na |
| *nad1x1 (1)* | Chr45 | 11477 | 11091 | 387 | Native | na |
| *nad1x2x3 (1)* | Chr42 | 2997 | 4461 | 273 (1465) | Native | na |
| *nad1x4x5 (1)* | Chr25 | 6425 | 9798 | 318 (3374) | Native | na |
| *nad2x1x2 (1)* | Chr01 | 17412 | 15721 | 545 (1692) | Native | na |
| *nad2x3x4x5 (2)* | Chr03 | 18261 | 14322 | 922 (3940) | Native | na |
| *nad3_1* | Chr02 | 4644 | 5000 | 357 | Foreign | ns |
| *nad3_2* | Chr34 | 5875 | 6231 | 357 | Native | na |
| *nad4 (2)* | Chr07 | 11568 | 16580 | 1476 (5013) | Chimeric | na |
| *nad4L* | Chr23 | 7489 | 7187 | 303 | Native | na |
| *nad5x1x2 (1)* | Chr06 | 18441 | 20683 | 1446 (2243) | Native | na |
| *nad5x3* | Chr51 | 3903 | 3924 | 22 | Native | na |
| *nad5x4x5 (1)* | Chr27 | 973 | 2357 | 543 (1490) | Native | na |
| *nad6* | Chr04 | 5004 | 5620 | 617 | Native | na |
| *nad7 (4)* | Chr24 | 11413 | 16965 | 1179 (5553) | Native | na |
| *nad9* | Chr28 | 2078 | 1506 | 573 | Native | na |
| *rpl2 (1)* | Chr18 | 10420 | 8057 | 1091 (2419) | Native | na |
| *rpl5* | Chr35 | 244 | 798 | 555 | Native | na |
| *rpl10* | Chr25 | 3065 | 3547 | 483 | Native | na |
| *rpl16* | Chr16 | 15575 | 15060 | 516 | Native | na |
| *rps1_1* | Chr14 | 6656 | 7339 | 684 | Native | na |
| *rps1_2* | Chr23 | 15108 | 14425 | 684 | Native | na |
| *rps3 (1)* | Chr16 | 18740 | 15466 | 1668 (3275) | Native | na |
| *rps4* | Chr04 | 3320 | 4390 | 1071 | Native | na |
| *rps10 (1)* | Chr26 | 12032 | 13197 | 420 (1223) | Native | na |
| *rps12_1* | Chr02 | 5049 | 5426 | 378 | Foreign | $p < 0.05$ |
| *rps12_2* | Chr34 | 6282 | 6659 | 378 | Native | na |
| *rps13* | Chr42 | 2021 | 2395 | 348 | Native | na |
| *rps19* | Chr46 | 9974 | 10255 | 282 | Native | na |
| *sdh3* | Chr01 | 25603 | 25316 | 288 | Native | na |
| *sdh4* | Chr04 | 21712 | 22177 | 466 | Native | na |

† The number of cis-spliced introns are indicated in parenthesis next to the gene name. The genes nad1, nad2 and nad5 also have 2, 1 and 2 trans-spliced introns, respectively.

In contrast, genes categorized as foreign consistently clustered phylogenetically as either a sister group to or within sequences of the host family Fagaceae, with bootstrap support exceeding 70%. Besides, BLAST analysis of complete chromosomes revealed that all foreign genes are embedded within larger foreign regions (Figure S3 and Figure S4). Several factors ruled out contamination from the host plant, providing strong evidence of HGT (see below). Furthermore, vertical inheritance of three of the six foreign gene copies (*atp1*, *ccmFN*, and *rps12*) was rejected by the AU test (Table 2). Unconstrained trees placing the foreign gene copy within the Fagaceae family exhibited high *p*-values in the AU test (Figure S5). Figure 2A presents examples of trees showing a foreign gene (*atp1*) located in the tree with high bootstrap support within the Fagales. Interestingly, the alignment of RNAseq data available from another individual of *Mitrastemon* (SRR28027597) shows that the *atp1* is expressed and edited at both predicted sites (Figure 2A).

**Figure 2.**
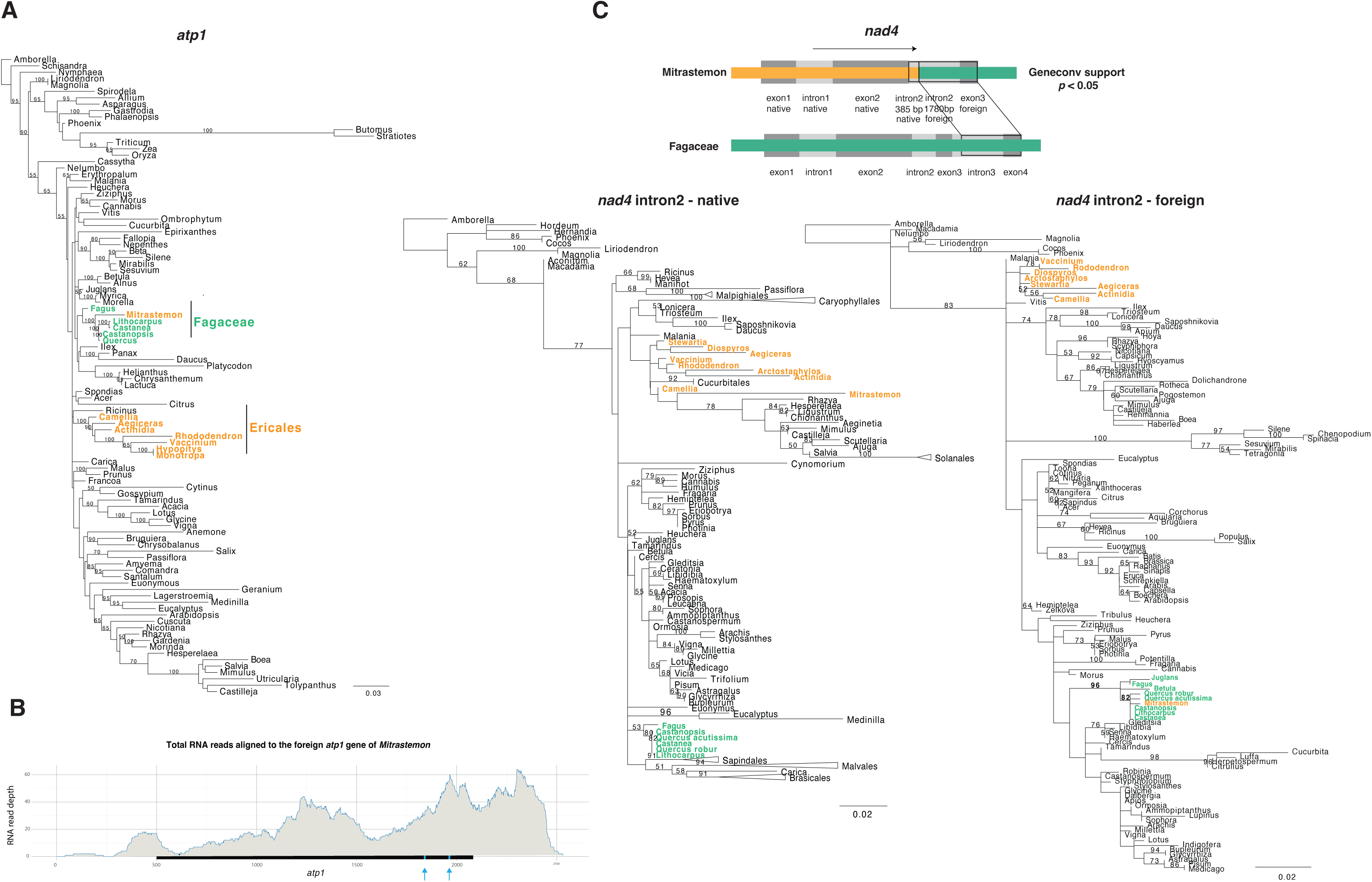
Analyses of a foreign (*atp1*) and a chimeric (*nad4*) mitochondrial gene of *Mitrastemon yamamotoi*. **A.** Maximum likelihood (ML) phylogenetic analysis of the *atp1* gene. **B.** RNA-seq data from another individual of *Mitrastemon* (SRR28027597) mapped to the foreign gene *atp1*, showing both transcription and RNA editing at the two predicted sites (blue vertical arrows). **C.** A schematic graph of the *nad4* gene with its exons in dark gray and introns in light gray. The native and foreign regions are indicated in orange and in green, respectively. The inferred recombination site is located within intron 2 between positions 329 and 385. Phylogenetic trees of the native (1-329) and the foreign (385-2165) regions of intron 2 are shown. Members of the family Fagaceae and Ericales are shown in green and orange, respectively. ML bootstrap support values > 50% are shown; some branches were collapsed to facilitate visualization. Bars correspond to substitutions per site.

One gene (*nad4*) in the mtDNA of *Mitrastemon* is classified as chimeric based on GeneConv results and phylogenetic analyses of gene subregions (Figure S2). The *nad4* gene shows a foreign origin of intron 2 and exon 3. While members of Fagales have three introns in the *nad4* gene, *Mitrastemon*, like other Ericales, has only two introns. However, recombination occurs in *Mitrastemon* intron 2, which corresponds to intron 3 in Fagales (Figure 2B).

### Chloroplast-derived sequences in *Mitrastemon yamamotoi* are foreign

We identified 11 chloroplast-derived sequences in the mtDNA of *Mitrastemon* (MTPTs), which together account for 0.9% of the genome (Table 1 and Figure 1). Phylogenetic analyses reveal that one of these sequences (*atpB*) has a long branch and it is not associated with any angiosperm lineage (Figure S6 and Table S3). The gene *atpB* is not present in the chloroplast genome of *Mitrastemon* (NC_080971); thus, the origin of this MTPT may be traced to an ancestral Intracellular Gene Transfer (IGT) prior to the loss of *atpB* from the cpDNA.

The remaining ten MTPTs are grouped with species of the Fagales, with strong support in the phylogenetic trees, indicating their origin through HGT (Figure S6). The majority of these MTPTs were located within larger segments of foreign mitochondrial sequences from the same lineage as the MTPT (Table S3, Figure S3 and Figure S4). These findings imply that chloroplast sequences were transferred through mitochondrion-to-mitochondrion HGT from a donor species of Fagales that had earlier incorporated plastid sequences into its mtDNA via IGT.

### Large-scale HGT from Fagaceae hosts to the mtDNA of *Mitrastemon yamamotoi*

To explore the impact of HGT along the noncoding regions of the mtDNA of *Mitrastemon* and tRNA and rRNA genes, similarity searches were performed. To increase the chances of identifying foreign mtDNA, we assembled mitochondrial genomic data from 16 *Castanopsis* spp. using the sequencing data available at SRA (NCBI). Similarity searches using local BLASTn against databases containing all published angiosperm mitochondrial genomes up to December 2024, including species of the host lineage (Fagales), revealed that 89% of the mitochondrial genome of *Mitrastemon* contains hits longer than 200 bp with identity percentages exceeding 70%, while the rest had no mitochondrial hits meeting this threshold and are considered of unknown origin (Figure 3 and Table S4). The phylogenetic origin of the noncoding mitochondrial regions was inferred with an R script that selects the best BLAST hits for each chromosome based on sequence identity and alignment length minimizing redundancy and maximizing coverage. This approach allowed us to retain the most relevant matches while ensuring minimal overlap between consecutive hits. About 61% (559 kb) of the mtDNA of *Mitrastemon* show Fagales species as the best hit (i.e. inferred as foreign DNA), with a mean identity of 96% (Figure 3 and Figure S4). In contrast, those native regions with Ericales as best hit exhibit a mean sequence identity of 90%.

**Figure 3.**
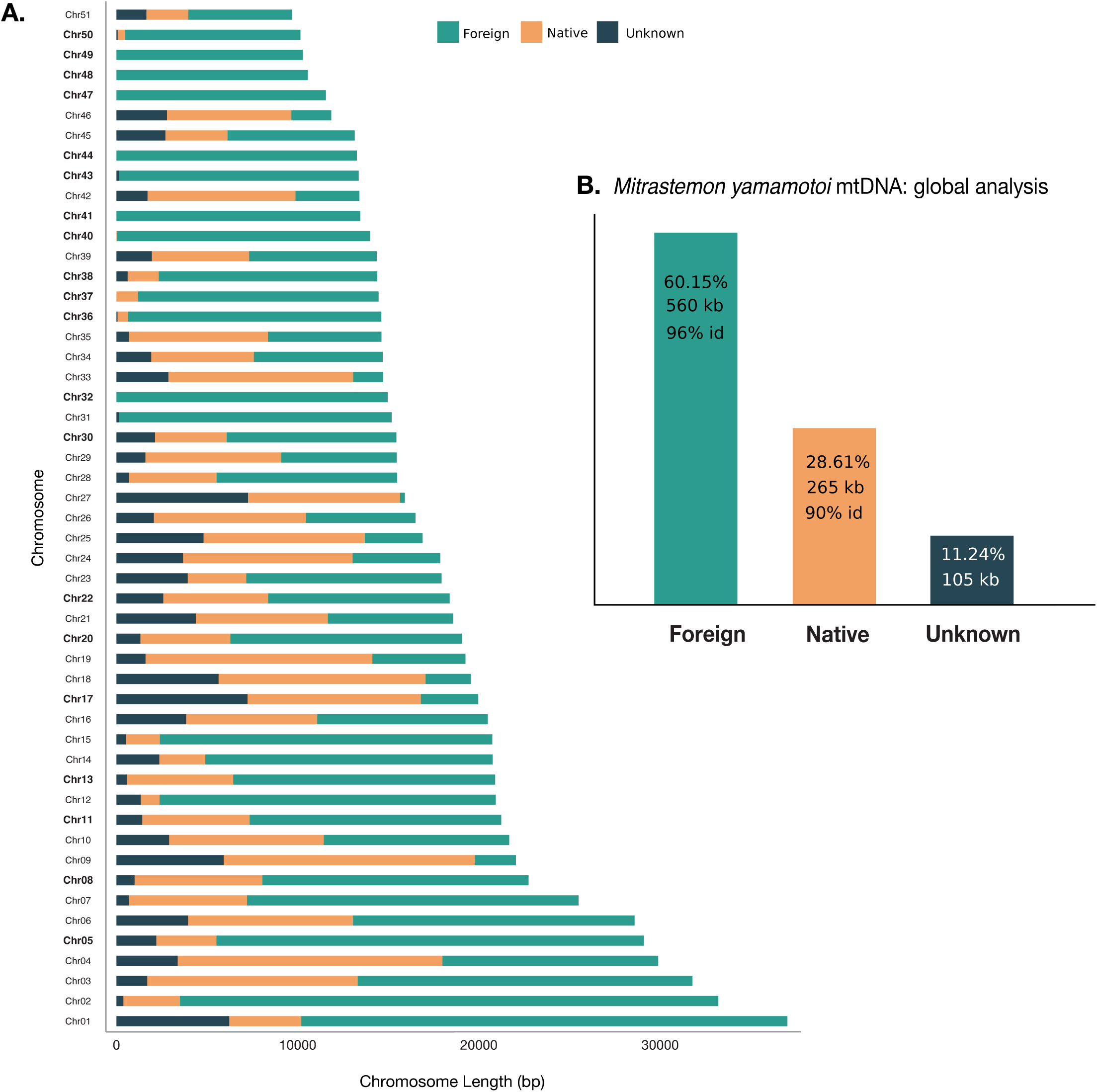
Evolutionary origin of the mtDNA of *Mitrastemon yamamotoi*. Shown is the amount of sequences with similarity to the Order Fagales (foreign; in green), Ericales or other angiosperm mitochondrial genomes (putatively native; in orange), and those regions with no significant BLASTn hits against the angiosperm mtDNAs (unknown origin; in blue), considering each mitochondrial chromosome individually (A) or the whole genome (B). The total amount, percent of the genome, and average sequence identity (id) of native and foreign regions are indicated for the global analysis. Chromosome names in boldface are noncoding (Table S4).

The foreign regions are spread across all 51 chromosomes of *Mitrastemon*, encompassing both coding and noncoding sequences and covering approximately 2–100% of each chromosome (Figure 3 and Table S4). BLASTn searches identified seven completely foreign chromosomes (32, 40, 41, 44, 47, 48, 49) and five chromosomes (31, 36, 37, 43, 50) with over 90% of its length of foreign origin (Table S4).

### Evidence of circle-mediated HGT in *Mitrastemon*

To evaluate the evidence supporting the circle-mediated HGT model (Roulet *et al*., 2024), we analyzed the seven completely foreign chromosomes in the *Mitrastemon* mtDNA. This involved searching for a continuous mitochondrial tract in the host mtDNA (*Castanopsis*), flanked by short direct repeats, which became a circular molecule with a single repeat in *Mitrastemon*.

The continuous mitochondrial regions in *Castanopsis* spp. (that were transferred to *Mitrastemon*) were identified by examining alternative conformations of the *Castanopsis* mitochondrial assemblies (Tu et al, 2024). This approach allowed us to demonstrate the existence of these continuous tracts in the mitochondria of the donor and confirm that they are flanked by direct repeats, which are 8-19 bp in length (Figure 4).

**Figure 4.**
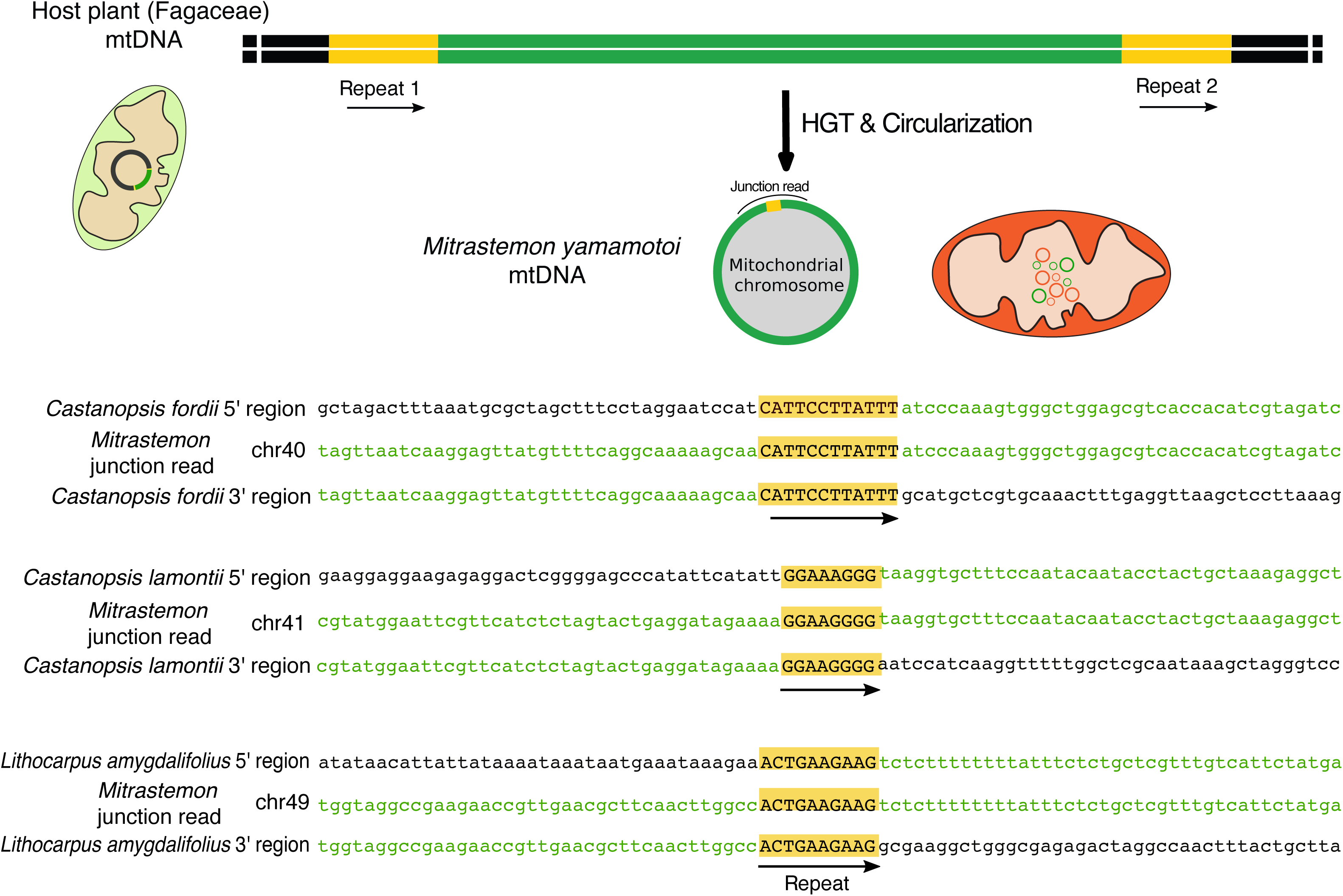
Formation of circular chromosomes according to the “circle-mediated HGT” model. In *Mitrastemon*, at least three foreign mitochondrial chromosomes originated from a region of the host mtDNA flanked by short direct repeats that mediate the circularization process.

### Mitochondrial-targeted DNA-RRR proteins in *Mitrastemon*

We examined the transcriptome of *Mitrastemon* to assess the expression of genes involved in DNA replication, repair, and recombination (DNA-RRR) that might be responsible for the rare events of circularization through microhomology-mediated pathways. A total of 24 nuclear-encoded DNA-RRR genes whose products are targeted to the mitochondrion in Arabidopsis were used as queries in BLASTN and TBLASTX searches against the transcriptome assembly and the identified open reading frames (ORFs) of *Mitrastemon*. For selected genes with multiple homologs, we ran phylogenetic analyses to identify which one is expressed in *Mitrastemon* (Figure S7). Overall, we identified all mitochondrial DNA-RRR genes in the transcriptome of *Mitrastemon*, except for OSB1 (Table S6). The completeness of the assembled transcriptome was assessed with Benchmarking Universal Single-Copy Ortholog (BUSCO). This assessment showed that the transcriptome assembly of *Mistrastemon* may be considered fairly complete for an endoparasitic plant (Table S6), as it shows similar results as those in other holoparasites (Garcia & Sanchez-Puerta, 2021; Yu *et al*., 2022; Chen *et al*., 2023).

## Discussion

### Multichromosomal mitochondrial genome and non-coding chromosomes in *Mitrastemon yamamotoi*

The parasitic family Mitrastemonaceae is monotypic and the genus *Mitrastemon* includes two species, *M. matudae* (often in the Americas) and *M. yamamotoi* (Southeast Asia), that are root endoparasites of Fagaceae species, mainly *Quercus* spp. and *Castanopsis* spp. (Teixeira-Costa & Suetsugu, 2023). Here, we assembled the mtDNA of an individual of *M. yamamotoi* that was parasitizing the roots of *Castanopsis* sp.

The mitochondrial genome of *Mitrastemon yamamotoi* is organized into 51 circular-mapping chromosomes, ranging from 9,684 to 37,021 bp. This multipartite configuration represents only one of several possible structures, given the presence of large repeats, e.g. a 10,752 bp repeat shared between Chr5 and Chr30. However, 21 chromosomes do not contain any intermediate or large repeats (>500 bp), and mate-pair information provides no evidence of recombination events involving these chromosomes. The lack of intermediate or large repeats in these chromosomes prevents the assembly of the mtDNA as a master circle, indicating that *Mitrastemon* has a multichromosomal mitochondrial genome. This complex architecture resembles that reported in other angiosperms, both in heterotrophic taxa such as those in the Balanophoraceae, in *Sapria himalayana* (Rafflesiaceae), or in the Orchidaceae (Sanchez-Puerta *et al*., 2017; Roulet *et al*., 2020; Yu *et al*., 2022; Guo *et al*., 2023; Zhou *et al*., 2023; Shen *et al*., 2024) (Yuan *et al*., 2018; Yang *et al*., 2023), and autotrophic, taxa such as *Silene* spp., *Pulsatilla patens*, or *Lilium tsingtauense* (Sloan *et al*., 2012; Szandar *et al*., 2022; Qu *et al*., 2024). This recurring theme of multipartite mitochondrial genomes in many heterotrophic and free-living plants suggests that while parasitism may exacerbate this complexity, there might be underlying genomic instability or other molecular mechanisms that predispose certain plant lineages to such highly fragmented configurations.

An especially intriguing aspect of *Mitrastemon* mitochondrial genome is the presence of 20 non-coding chromosomes. The persistence of such a large number of non-coding chromosomes suggests either a very relaxed selective pressure on mitochondrial genome size and content in *Mitrastemon*, or alternatively, a highly efficient, yet uncharacterized, mechanism for their replication and segregation. Similar patterns have been observed in other angiosperms with multichromosomal mtDNAs, such as *Lophophytum mirable* and *Silene noctiflora* (Sloan *et al*., 2012; Sanchez-Puerta *et al*., 2017). Analyses of multiple individuals of these species revealed a high degree of variability in the presence/absence of the non-coding chromosomes, both within and between populations, suggesting they are evolving by genetic drift (Wu *et al*., 2015; Gatica-Soria *et al*., 2025). While there is still considerable uncertainty regarding the origin of the non-coding chromosomes in *Silene* (Wu & Sloan, 2019), those of *L. mirabile* are clearly the result of HGT (Gatica-Soria *et al*., 2025). This precedent raises compelling questions about the origin and variability of non-coding chromosomes in *Mitrastemon*. Similar to *L. mirabile*, this study supports a foreign mitochondrial origin for most of the mtDNA of *Mitrastemon*. Examining multiple individuals of this holoparasite will be necessary to assess the variability of these chromosomes at the population level, to define the *Mitrastemon* mitochondrial pangenome, to infer the rates of ongoing host-to-parasite transfers, and to understand the evolutionary trade-offs and molecular mechanisms involved in maintaining so many non-coding molecules.

### High prevalence of HGT in the mtDNA of *Mitrastemon* from Fagaceae hosts

A defining characteristic of parasitic plants is the vascular and intimate connections formed by haustoria between the parasite and its host (Mower *et al*., 2010; Xi *et al*., 2013; Davis & Xi, 2015; Góralski *et al*., 2021). *Mitrastemon* establishes both xylem and phloem connections with their hosts (Teixeira-Costa & Suetsugu, 2023), creating a close interaction that facilitates the transfer of genetic material. Through sequence similarity searches across the entire mtDNA and phylogenetic analyses of coding regions, we identified extensive foreign mitochondrial regions derived from Fagales hosts in the *Mitrastemon* mtDNA, comprising more than 60% of the genome. This extraordinary scale of genetic exchange underscores the hypothesis that intimate, prolonged host-parasite interactions, characteristic of endoparasites like *Mitrastemon*, are potent facilitators of extensive HGT.

The host-derived DNA is not evenly distributed across the mitochondrial chromosomes of *Mitrastemon*, which carry 2 to 100% of foreign DNA. The non-coding regions take up 99 % of the foreign DNA. This disproportionate transfer of non-coding DNA, despite the massive overall HGT, suggests that while the mechanism of HGT (e.g., cell-to-cell connection) is highly efficient in facilitating DNA uptake, the selection pressure for integrating and maintaining functional foreign genes is considerably lower.

Given that *Castanopsis* is the reported host for *M. yamamotoi*, we generated mitochondrial assemblies of 16 *Castanopsis* species, and incorporated data from other Fagaceae (*Quercus* spp., *Lithocarpus* sp., *Castanea* spp., and *Fagus* sp.) to further evaluate the HGT process. In the BLASTn searches, *Castanopsis* was consistently identified as the best hit based on hit length and sequence identity. These results demonstrate that *Castanopsis* mitochondrial data provides a large number of foreign DNA tracts including completely foreign chromosomes, which are consistent with the circle-mediated HGT model (see below). The limited number of complete mtDNA sequences currently available for Fagaceae (only 36 out of >1000 described species) likely leads to an underestimation of the actual number and length of the foreign tracts. The 11% of the *Mitrastemon* mtDNA with unknown origin that had no mitochondrial hits longer than 200 bp are particularly intriguing. These regions could represent additional host-derived DNA, which are present in a yet unsampled host plant, or nuclear-derived DNA obtained by IGT.

### Circle-mediated HGT model

Foreign regions in the mtDNA of *Mitrastemon* include both coding and non-coding sequences, and in seven chromosomes, they span the entire length of the chromosome. The discovery of fully foreign chromosomes that consist of continuous DNA tracts found in the host mtDNA flanked by short direct repeats provides compelling evidence for the recently proposed circle-mediated HGT model (Roulet *et al*., 2024). According to this model, mitochondrial DNA segments from the donor plant can enter the recipient mitochondria and form autonomous circular molecules without immediate integration with the resident mitochondrial genome. The circularization process seems to be mediated by microhomologies involving rare DNA repair pathways across short direct repeats flanking the DNA segment that becomes circular. Over time, these circular chromosomes may persist independently, replicate, or undergo recombination with other mitochondrial chromosomes through homologous recombination, ultimately giving rise to chimeric chromosomes. Our identification of these completely foreign circular chromosomes, their continuity as tracts in *Castanopsis* mitochondria in three of them, and the recovery of flanking short direct repeats (8-19 bp) represent direct evidence for this mechanism.

The support of the circle-mediated HGT mechanism in *Mitrastemon* represents the first evidence of this process outside of the Santalales family, expanding the occurrence of this mechanism in the evolution of mitochondrial genomes in parasitic plants. This mechanism, where foreign DNA forms autonomous circular molecules, has also been implicated in interkingdom HGT events. For instance, a circularization process mediated the horizontal transfer of a fungal mitochondrial region, including tRNA genes, into orchid mitochondria (Sinn & Barrett, 2020). Other studies also point to circularization as a key step in HGT across diverse lineages, highlighting its potential as a general mechanism for foreign DNA transfer and maintenance (Kempken & Kück, 1998; Rosewich & Kistler, 2000; Zhang *et al*., 2024)

### Missing DNA-RRR gene expression may impact the mtDNA stability and repair in *Mitrastemon*

The circumstances under which foreign DNA can form subgenomic circular molecules and undergo autonomous replication within the mitochondria of the recipient plant are unclear. Depletion of DNA-RRR factors or genomic loss of relevant DNA-RRR genes have been suggested as potential conditions for circularization (Roulet *et al*., 2024) based on the involvement of rare microhomology-mediated repair pathways, which are normally inhibited in plant mitochondria (Gualberto & Newton, 2017). This is an interesting mechanistic hypothesis, suggesting a link between the intrinsic genomic maintenance machinery of the recipient plant and its susceptibility to specific HGT mechanisms. It implies that the recipient’s recombination and repair pathways might be a key factor in determining the fate of HGT.

Of the 24 DNA-RRR genes described in Arabidopsis whose products are targeted to the mitochondrion, all but one gene (*osb1*) were identified in the transcriptome of *Mitrastemon*. The missing gene *osb1* is plant-specific and codes for a single stranded binding protein that promotes homologous recombination and suppresses ectopic rearrangements (Gualberto & Newton, 2017). Several Arabidopsis mutants centered on mitochondrion-targeted DNA-RRR factors have demonstrated that recombination/repair deficiencies in plant mitochondria lead to genomic instability and to the generation of episomes that often show increased stoichiometry (Cappadocia *et al*., 2010; Maréchal & Brisson, 2010; Wallet *et al*., 2015; Chevigny *et al*., 2020; Schatz *et al*., 2025). In particular, *osb1* mutants show increased ectopic homologous recombination in the mitochondrial genome (Zaegel *et al*., 2006; Qian *et al*., 2022). The holoparasitic *Lophophytum* spp., which harbors dozens of foreign mitochondrial chromosomes acquired from mimosoid hosts through circle-mediated HGT, lack expression of the *why2* gene (Roulet *et al*., 2024). It has been shown that *why2* mutants exhibit increased illegitimate recombination involving sequence microhomologies when treated with a genotoxic agent that produces double strand breaks in the organellar genomes (Cappadocia *et al*., 2010). Overall, altered expression or missing mitochondrial DNA-RRR genes may allow the rare circularization of foreign DNA in angiosperm mitochondria.

### Foreign and chimeric genes in the mtDNA of *Mitrastemon*

Initially, plant mitochondrial HGT results in gene duplication (duplicative HGT), where both native and foreign gene copies are present as full-length genes. In *Mitrastemon* mtDNA we found three cases of duplicative HGT. In addition, foreign copies frequently become pseudogenes and are eventually lost (Bergthorsson *et al*., 2003; Davis & Wurdack, 2004; Davis *et al*., 2005; Rice *et al*., 2013; Xi *et al*., 2013; Park *et al*., 2015; Cusimano & Renner, 2019; Forgione *et al*., 2019; Garcia *et al*., 2021). We found evidence of seven foreign pseudogenes in *Mitrastemon* mtDNA.

Previous studies in *Amborella trichopoda* (Rice *et al*., 2013) and *Lophophytum* spp. (Roulet *et al*., 2024) (García et al., 2021; Roulet et al., 2024) have shown that, when full-length foreign and native copies coexist, the native copies are the only functional alleles, based on efficient transcription and RNA editing. This is likely the case for the duplicative HGT cases in *Mitrastemon* as well, but the limited depth of the transcriptomic data available precluded this analysis. The presence of so many duplicated foreign genes or pseudogenes in *Mitrastemon* suggests that while the mechanism of DNA uptake is highly efficient, the selective pressure for maintaining functional foreign genes is considerably lower. However, even if non-functional, the persistence of these foreign genes and pseudogenes highlights the remarkable permissiveness of the recipient mitochondrial genome to foreign material, possibly due to relaxed selection, and opens the possibility of future recombination events giving rise to genetic diversity in the functional mitochondrial genes. In several plant mtDNAs (Bergthorsson *et al*., 2003; Barkman *et al*., 2007; Hao *et al*., 2010; Mower *et al*., 2010; Hepburn *et al*., 2012; Park *et al*., 2015; Sanchez-Puerta *et al*., 2017; Roulet *et al*., 2020; Yu *et al*., 2023) recombination between the native and foreign gene copies led to the formation of chimeric genes (chimeric or partial replacement HGT). A single chimeric gene (*nad4*) was identified in *Mitrastemon*.

In rare instances, native genes are completely replaced by foreign homologs (full replacement HGT). This has been documented in a few plant species to date (Bergthorsson *et al*., 2003; Xi *et al*., 2013; Bellot *et al*., 2016; Sanchez-Puerta *et al*., 2017; Garcia *et al*., 2021; Roulet *et al*., 2024). In the mtDNA of *Mitrastemon*, we found two cases of full replacement HGT in the *atp1* and *atp6* genes. Notably, the foreign *atp1* gene shows accurate and efficient RNA editing, consistent with a functional integration. This essential gene represents the sole mitochondrial copy, and no evidence of a nuclear-encoded version was found in transcriptomic data. The functional horizontal transfer of *atp1* is quite remarkable as the vast majority of the mitochondrial HGT events in plants involve non-functional duplicated copies of mitochondrial genes or pseudogenes. To date, only a few studies have provided strong evidence of functional mitochondrial HGT from other plants (Bergthorsson *et al*., 2003; Hao *et al*., 2010; Garcia *et al*., 2021) and tRNAgenes from fungi (Warren *et al*., 2025).

## Conclusions

The study reinforces the hypothesis that species undergoing a prolonged endophytic stage are more prone to HGT, as highlighted by the large-scale mitochondrial HGT in the endoparasite *Mitrastemon yamamotoi*. Our findings support the fusion compatibility model of mitochondrial HGT because long tracts of mtDNA from the hosts, and no plastid or nuclear DNA, were horizontally transferred indicating that mitochondrial fusion likely mediated the HGT process. Furthermore, it provides compelling evidence for the circle mediated HGT model, suggesting that under certain circumstances (e.g. depletion of DNA-RRR factors or genomic loss of relevant DNA-RRR genes), foreign DNA can become circular and undergo autonomous replication in the mtDNA increasing the likelihood of being maintained across generations consolidating the HGT event. By demonstrating this mechanism in *Mitrastemon yamamotoi* (Ericales), the study provides critical independent validation, suggesting that this is not an isolated evolutionary event identified in Balanophoraceae (Santalales), but potentially a more general mechanism for foreign DNA acquisition and maintenance in plant mitochondria.

## Materials and Methods

### Sample collection and DNA extraction and sequencing

An individual of *Mitrastemon yamamotoi* was collected in Menghai, Yunnan, China (22.0122° N, 100.3775° E) on Oct 1, 2020, while parasitizing the roots of a *Castanopsis* sp. Total DNA from flower tissue (to avoid contamination from the host) was isolated using the CTAB method and a DNA library with an insert size of 350 bp was constructed and sequenced on Illumina platform. The clean data have been deposited in NCBI (SRR34031518).

### Assembly of mitochondrial genome

A *de novo* assembly of the DNAseq reads from *Mitrastemon yamamotoi* was performed using GetOrganelle v.1.7.1 (Jin *et al*., 2020) with default parameters for plant mitogenome assembly, except k set to 127. We identified putative mitochondrial contigs if they have a BLASTn hit to a custom dataset of angiosperm mitochondrial genes and high read coverage (>500x). The final mitochondrial assembly comprised circular chromosomes, with circularity validated by paired-end reads showing one mate mapping to the contig’s start and the other to its end, as visualized in CONSED v.29.0 (Gordon & Green, 2013). The DNA read depth for each chromosome was calculated using BOWTIE2 v.2.2.2 (Langmead & Salzberg, 2012) with the presets: –*end-to-end* –*very-fast*, SAMTOOLS v.1.10 (Li *et al*., 2009) and BEDTOOLS v.2.26.0 (Quinlan & Hall, 2010), while the plots were generated using R (Figure S1).

### Mitochondrial genome annotation

The mitochondrial genome annotation was performed using BLASTN searches (Camacho *et al*., 2009) with angiosperm mitochondrial genes as queries. The tRNA genes were identified with tRNAscan-SE algorithm with default and infernal search modes without HMM filter and using the bacterial, organellar, and general sequence sources (Lowe & Eddy, 1997). Plastid-derived mitochondrial sequences (MTPTs) > 200 bp were detected by BLASTn searches against a custom chloroplast database including Ericales and Fagaceae chloroplast databases. Pseudogenes greater than 100 bp were also annotated. The repetitive content of the genome was analyzed using the *get_repeats.sh* script developed by Gandini et al. (Gandini *et al*., 2019) (Table S1). The annotated mitochondrial chromosomes were deposited in Genbank (PV791430-PV791480).

### Mitochondrial genome assemblies of *Castanopsis* spp

To obtain additional data to identify HGT events in the mitochondrial genome of *Mitrastemon*, we assembled the mitochondrial genomes of 16 *Castanopsis* species using SPAdes v.3.15.2 (Bankevich *et al*., 2012). Assemblies were performed with the expected coverage and coverage cutoff set to automatic, and k-mer lengths of 71, 87, and 91 (Table S5A). Alternative arrangements were evaluated based on paired-end read information and the presence of repeats.

### Comparative genomics analyses

To further evaluate the phylogenetic origin of each mitochondrial chromosome of *Mitrastemon*, we performed BLASTn searches of each chromosome against the custom angiosperm mitochondrial database (Table S5A-B), including the mitochondrial assemblies of 16 *Castanopsis* spp. (typical hosts of *Mitrastemon*). Thus, a database was compiled containing all sequenced mitochondrial genomes of angiosperms up to December 2024 (Table S5A-B). BLASTn searches were performed with the 51 mitochondrial chromosomes of *Mitrastemon* as queries, and the database divided into five groups: *Castanopsis spp.*, Fagaceae, Fagales, Ericales, and the remaining angiosperms (excluding Fagales and Ericales) for better visualization. BLASTn hits with an *e*-value < 2 x10^-10^ were plotted using the SUSHI R package v.1.20.0 (Phanstiel *et al*., 2014).

To identify the most relevant BLAST hits while minimizing redundancy, we applied a filtering and selection strategy to the BLAST results dataset. First, we retained only hits with an alignment length greater than 500 bp and a sequence identity greater than 70%. For each chromosome, we iteratively selected the best BLAST hit for each genomic segment. The process followed these steps:

1. The algorithm started at the beginning of the chromosome and iteratively selected the best hit that covered the current genomic segment.
2. Hits were first ranked by sequence identity (pident), and in case of ties, by alignment length. The highest-ranking hit was selected.
3. The process advances to the end of the selected hit, ensuring minimal overlap between consecutive hits.
4. If no hits were found for a given segment, the algorithm advances 10 bp to continue the search.
5. The selected hits were stored in a list for downstream analysis and plotted using ggplot2 package (Wickham *et al*., 2016).

This approach ensured that the most relevant hits were retained while reducing redundancy and ensuring broad coverage of each chromosome. A region within the mtDNA of *Mitrastemon* was classified as foreign if it fulfilled at least one of the following criteria: (1) BLAST hits were exclusively found in members of the host family (Fagaceae) or (2) BLAST hits to the host lineage (Fagaceae) had higher sequence identity and were longer than those from other angiosperms (Sanchez-Puerta *et al*., 2019).

### Phylogenetic inference

Phylogenetic analyses under Maximum Likelihood (ML) were performed for each protein-coding gene and MTPT. For protein-coding genes nucleotide sequences of diverse angiosperms (Table S5C) were aligned with Muscle in AliView v.1.27 (Larsson, 2014). For the MTPTs alignments, we incorporated chloroplast data from angiosperm species, as well as other MTPTs identified in additional angiosperms. ML trees were obtained with RAXML v.8.2.11 (Stamatakis, 2014) using the GTRGAMMA model, including 1,000 rapid bootstrap pseudoreplicates. The best-fitting nucleotide substitution models (GTRGAMMA) were estimated by MODELTEST version 3.7 using the Akaike information criterion (Posada & Crandall, 1998). The resulting trees were visualized with Figtree v.1.4.4 http://tree.bio.ed.ac.uk/software/figtree/.

For each foreign gene identified in the *Mitrastemon* mtDNA, we conducted an approximately unbiased (AU) test (Shimodaira, 2002). We generated constrained topologies (Figure S5), enforcing the foreign copy of *Mitrastemon* to cluster with Ericales while excluding its association with Fagaceae, as would be expected under vertical inheritance. Site likelihoods for both constrained and unconstrained trees were estimated using PAUP v.4.0 (Swofford *et al*., 2002), and AU *p-*values were calculated with CONSEL.

Besides, to evaluate the presence of chimeric genes in *Mitrastemon*, gene alignments were analyzed using GeneConv v.1.81a (Sawyer, 1989). The alignments incorporated gene sequences from multiple Ericales and host species (Fagaceae) (Table S5D). Additionally, phylogenetic analyses were carried out on specific gene subregions, and BLASTn searches were performed on flanking sequences.

### Transcription and RNA editing of foreign mitochondrial genes

To evaluate the expression of foreign mitochondrial genes identified in *Mitrastemon* mtDNA, we aligned RNA sequencing data from another individual of *M. yamamotoi* available in NCBI (SRR28027597) (Carruthers *et al*., 2024). The RNA read depth was calculated as described above.There was sufficient RNA coverage for *atp1*, but not for *atp6*, *ccmFN*, or *rps12*. We predicted the non-synonymous RNA editing sites in *atp1* with PREPACT3 v.3.12.0 (Lenz *et al*., 2018) and evaluated the actual editing by counting the number of Us in the RNA reads aligned to the predicted editing cytosines.

### Analysis of host contamination in the mtDNA of *Mitrastemon yamamotoi*

The possibility that the mitochondrial genome of *Mitrastemon yamamotoi* includes sequences derived from host contamination was carefully evaluated and effectively ruled out based on multiple independent lines of evidence. First, DNA was extracted from flower tissues of *Mitrastemon*, which are developmentally derived from the parasite and not in direct contact with host tissue, thereby minimizing the risk of contamination during sampling. Second, phylogenetic analyses revealed that the foreign sequences found in the *Mitrastemon* mtDNA consistently cluster as sister groups to members of the Fagaceae family, rather than grouping specifically with the host species (*Castanopsis* sp.) as expected in case of contamination. Third, DNA read mapping analysis demonstrated even and consistent read depth across all mitochondrial chromosomes, including those identified as foreign or with foreign genes. No evidence of abnormal read depth or gaps was observed in these regions, which supports their genuine presence of HGT in the mtDNA. Fourth, RNA sequencing data from a different individual of *M. yamamotoi* (SRR28027597) presents the same foreign genes as those reported in the individual studied here. Collectively, these observations provide robust support for the authenticity of the foreign sequences and confirm that they are the result of true HGT events rather than sample contamination.

### Identification of DNA-RRR genes with roles in the mitochondrion in the transcriptome of *Mitrastemon*

To identify DNA-RRR genes whose products are targeted to the mitochondrion, we *de novo* assembled the transcriptome of *Mitrastemon* with RNAseq data available at NCBI (SRR28027597, Carruthers et al., 2024) using Trinity v.2.15.0 with default parameters except for --min_contig_length 100. Transdecoder was used to find ORFs for each transcript sequence. To assess the completeness of the transcriptome assembly, the BUSCO v.5.8.2 (Waterhouse *et al*., 2018) was executed in genome mode, based on the eukaryota_odb10 (255 genes), viridiplantae_odb10 (425), embryophyta_odb10 (1,614 genes) and eudicots_odb10 (2,326 genes) datasets, respectively. We focused on 24 DNA-RRR genes whose products are targeted to the mitochondrion (Gualberto & Newton, 2017; Schatz *et al*., 2025). Using the protein sequences of *Arabidopsis* as queries, we conducted BLASTN and TBLASTX to search for homologous sequences in the assembled transcriptome and the identified ORFs of *Mitrastemon* with the parameter -evalue 1e-3. For genes that are duplicated in *Arabidopsis* due to ancestral duplication events and for which multiple homologs were identified in *Mitrastemon*, phylogenetic analyses were performed as described above.

## Acknowledgements

We thank W. Tulle and L. Gatica-Soria for help with the computational analyses. This work used the SARTOI Cluster from IBAM (FCA-UNCuyo) and was supported by grants from Fondo para la Investigación Científica y Tecnológica (grant N°: PICT2020-01018) and Universidad Nacional de Cuyo (grant N°: 06/A092-T1).

## Short legends for Supporting Information

**Table S1.** Repeats with >90% sequence identity detected in the *Mitrastemon yamamotoi* mtDNA.

**Table S2.** Features of ribosomal, transfer RNA genes, and pseudogenes in the *Mitrastemon yamamotoi* mtDNA.

**Table S3.** Features of chloroplast-derived sequences in the *Mitrastemon yamamotoi* mtDNA.

**Table S4.** Proportion of each mitochondrial chromosome from *Mitrastemon yamamotoi* inferred to be foreign, native or of unknown origin.

**Table S5.** Accession numbers of the mitochondrial sequences used in BLASTn, phylogenetic, and GeneConv analyses.

**Table S6.** BUSCO analysis of the transcriptome assembly of *Mitrastemon yamamotoi* and identification of mitochondrial DNA-RRR genes.

**Figure S1.** Read depth of Illumina DNA reads mapped on the mitochondrial chromosomes of *Mitrastemon yamamotoi*.

**Figure S2.** Maximum Likelihood (ML) phylogenetic analyses of coding sequences in the mtDNA of *Mitrastemon yamamotoi*.

**Figure S3.** Visualization of BLASTn searches of the mitochondrial chromosomes in *Mitrastemon yamamotoi*.

**Figure S4.** Phylogenetic origin of the intergenic regions of the mtDNA of *Mitrastemon yamamotoi*.

**Figure S5.** Approximately unbiased (AU) tests to evaluate alternative topologies based on different constrained trees.

**Figure S6.** Maximum Likelihood phylogenetic analyses of chloroplast-derived sequences (MTPTs) in the mtDNA of *Mitrastemon yamamotoi*.

**Figure S7.** Maximum Likelihood phylogenetic analyses of duplicated DNA-RRR genes.

## Notes

**Funding statement** This work was supported by grants from Fondo para la Investigación Científica y Tecnológica (grant # PICT2020-01018) and Universidad Nacional de Cuyo (grant # 06/A092-T1) to MVS-P. This study was supported by the National Natural Science Foundation of China (grant # 31811530297 to R.Z.).

### Competing Interest Statement

The authors have declared no competing interest.

