## Supplementary Figures for "Circle-Mediated HGT shapes the multichromosomal mitochondrial genome of the endoparasite *Mitrastemon yamamotoi*"

Figure S1. Read depth of Illumna DNA reads mapped on mitochondrial chromosomes of *Mitrastemon yamamotoi*.

### Chromosome1

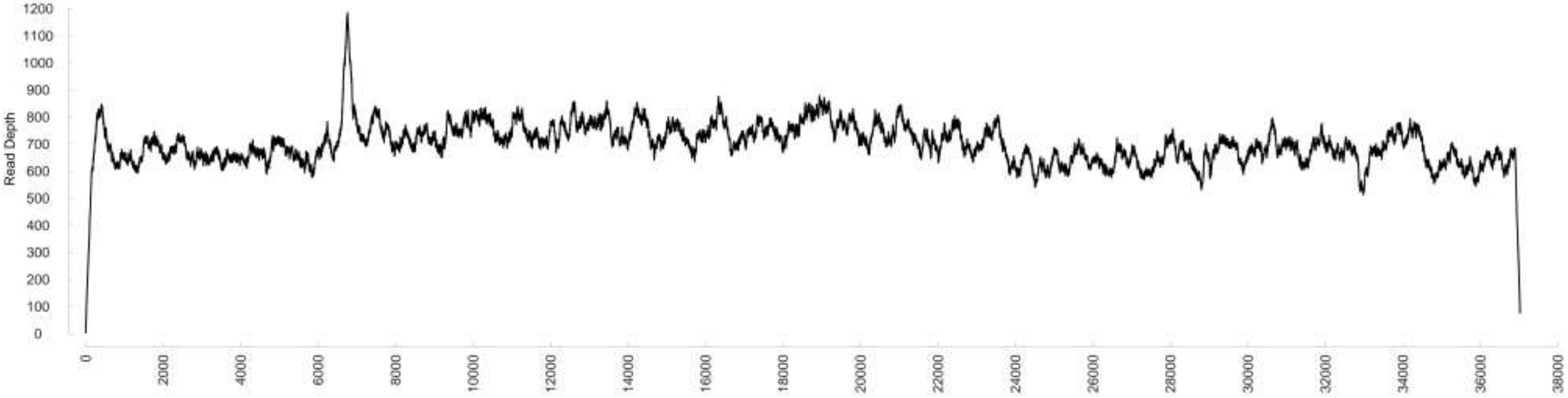

### Chromosome2

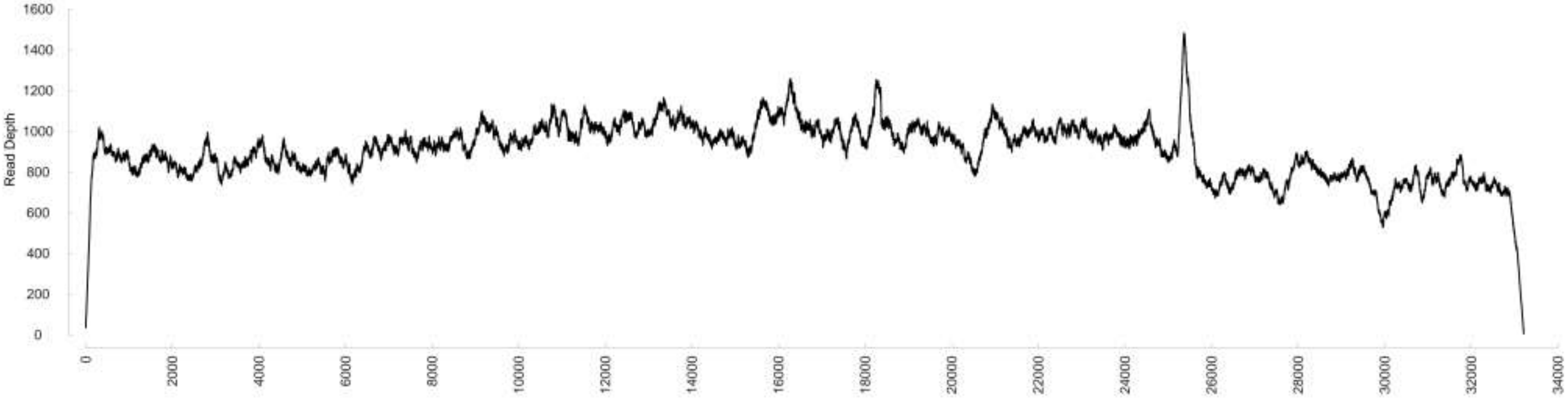

### Chromosome3

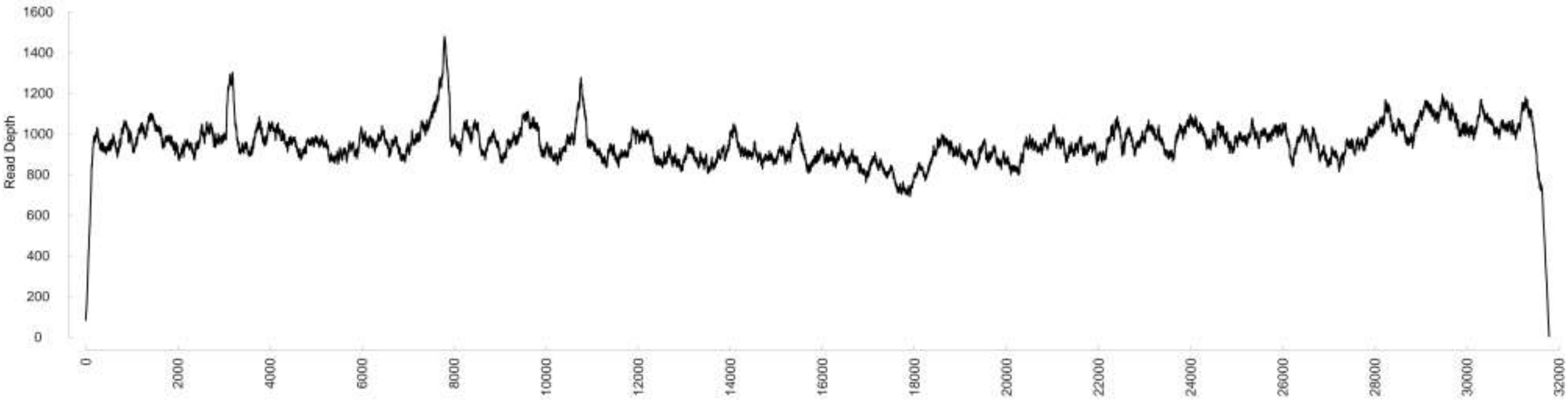

### Chromosome4

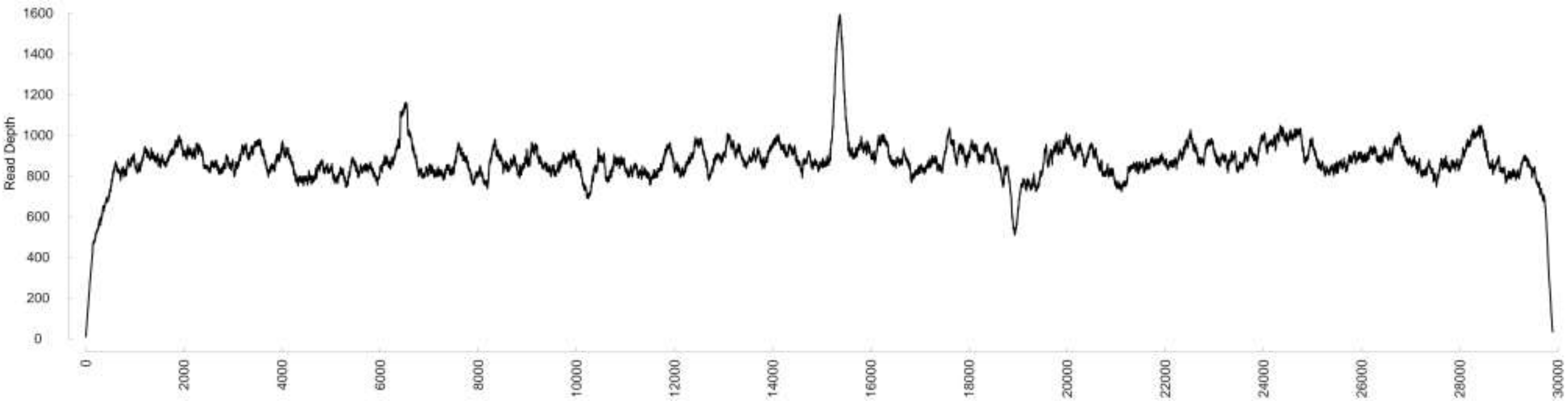

### Chromosome5

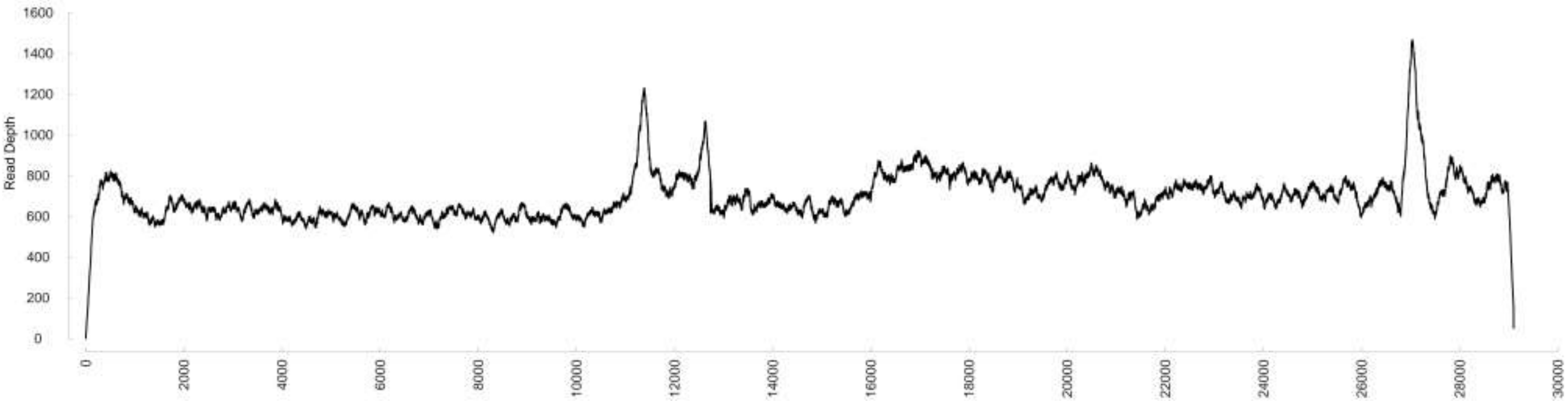

### Chromosome6

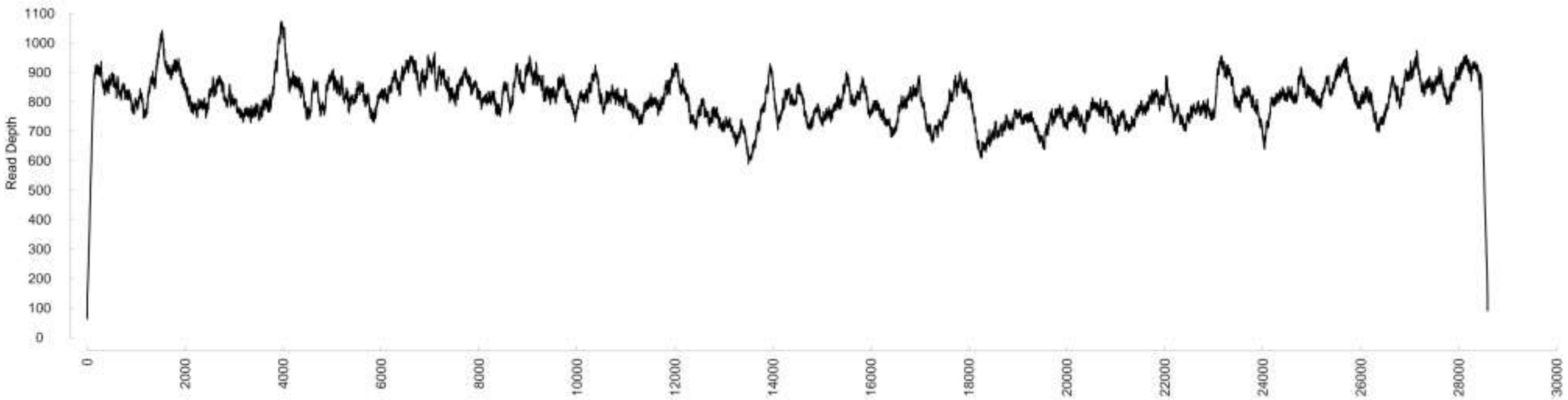

### Chromosome7

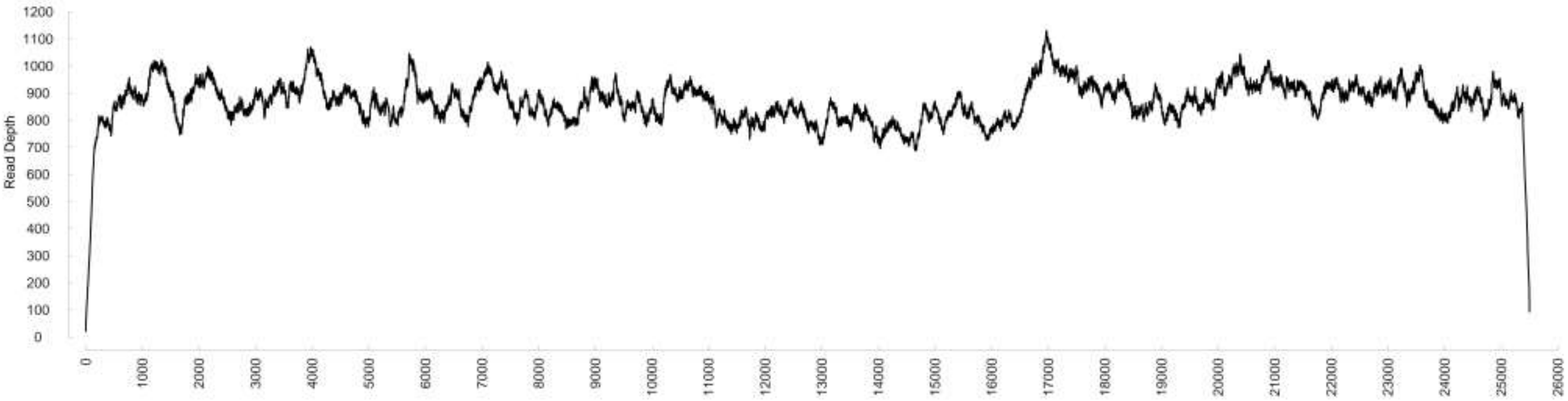

### Chromosome8

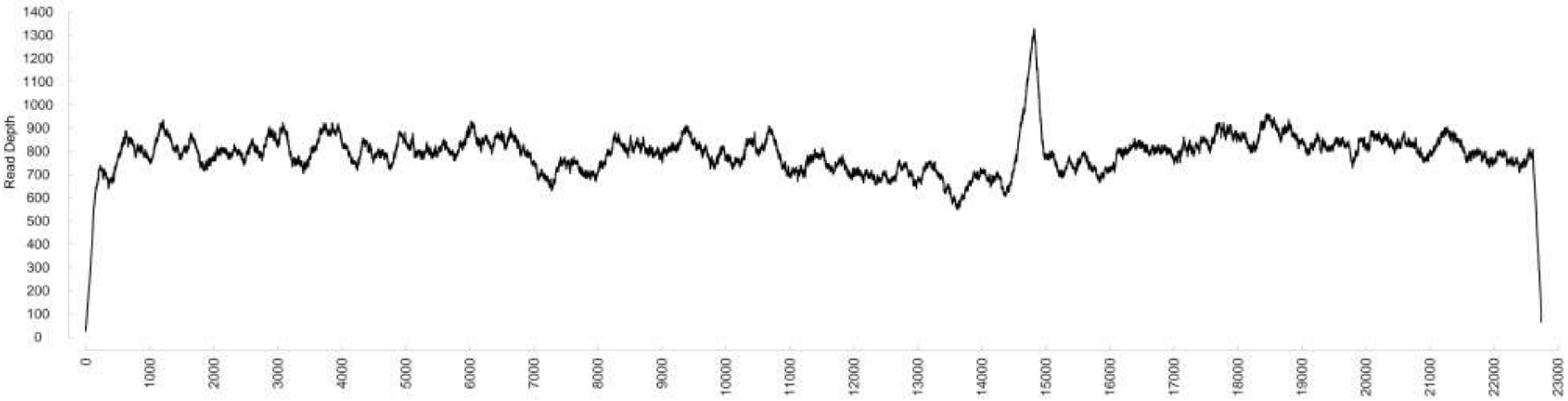

### Chromosome9

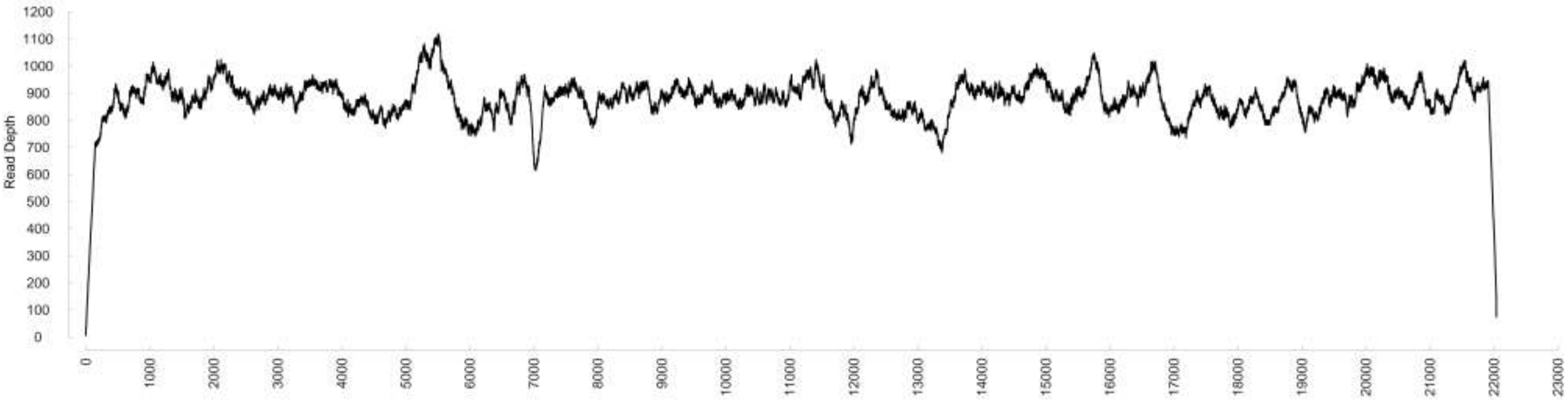

### Chromosome10

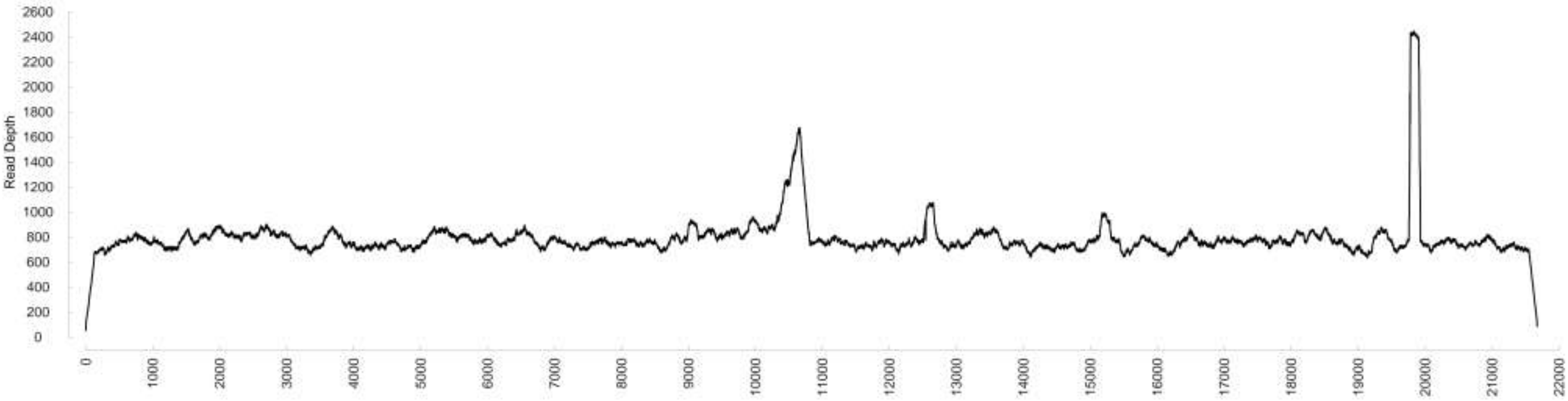

### Chromosome11

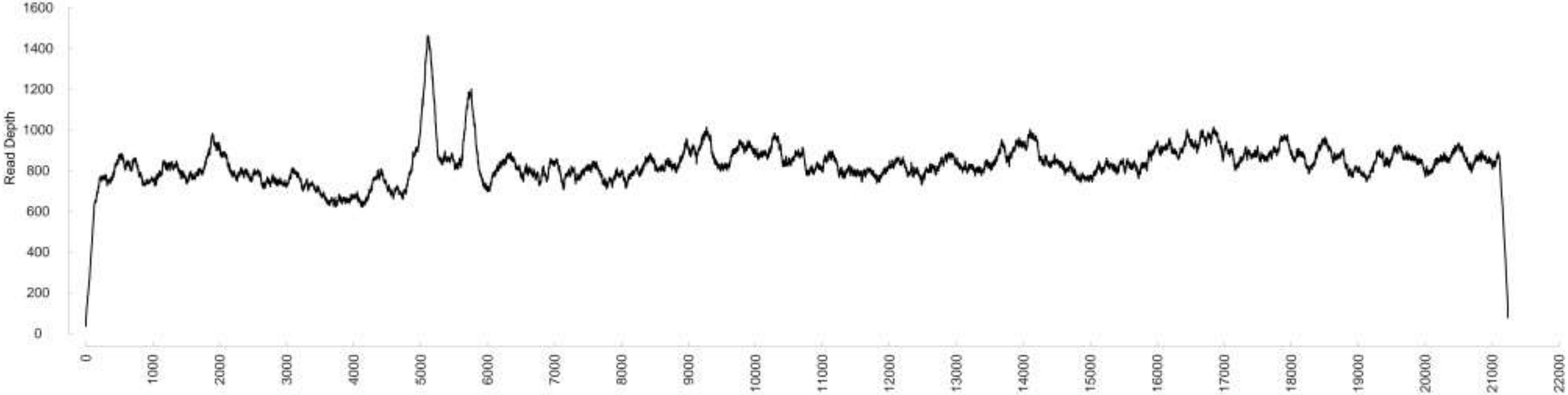

### Chromosome12

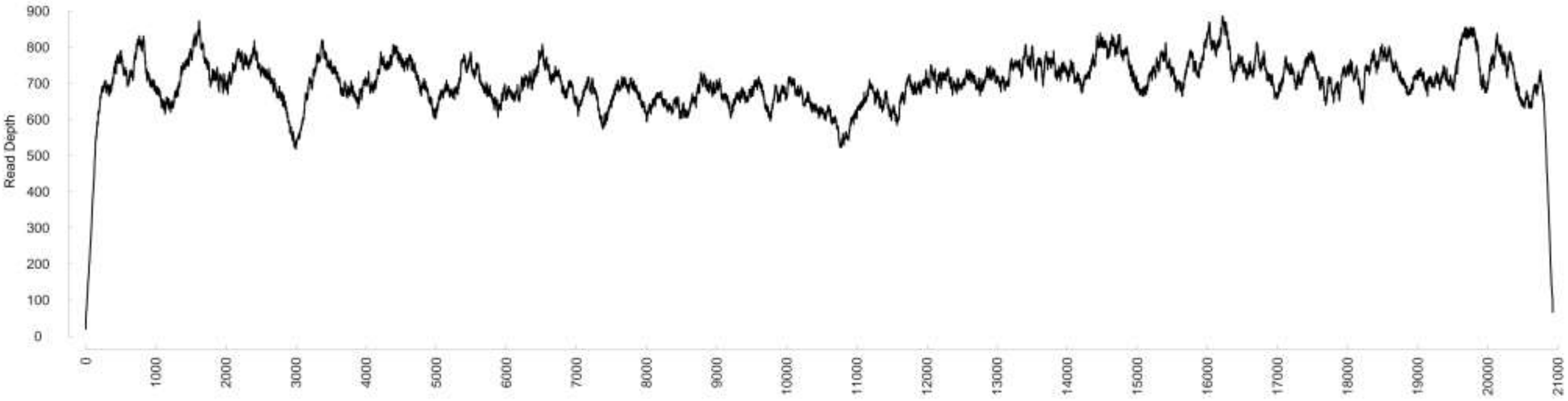

### Chromosome13

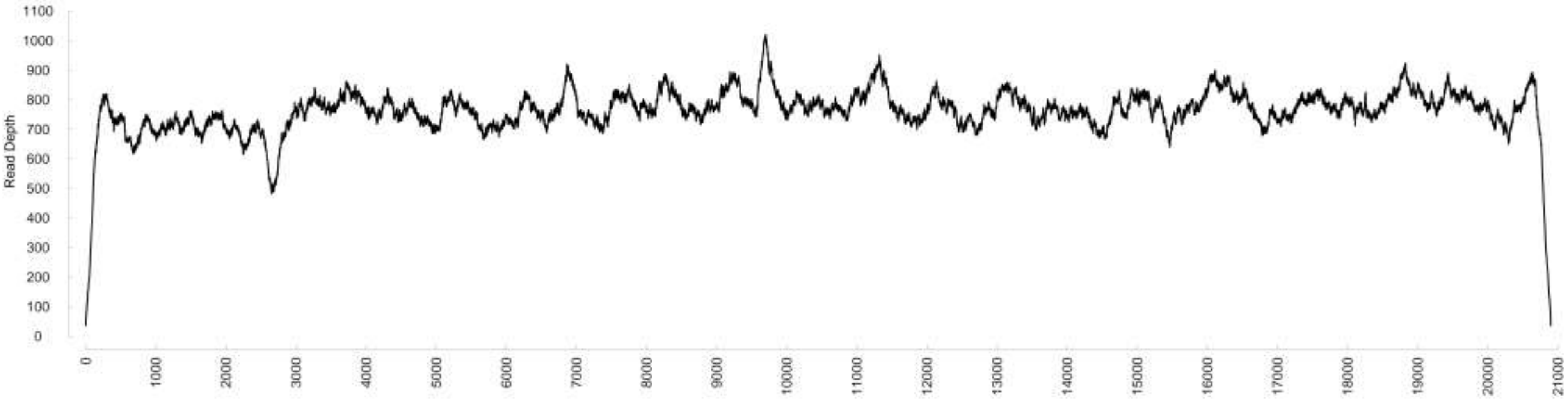

### Chromosome14

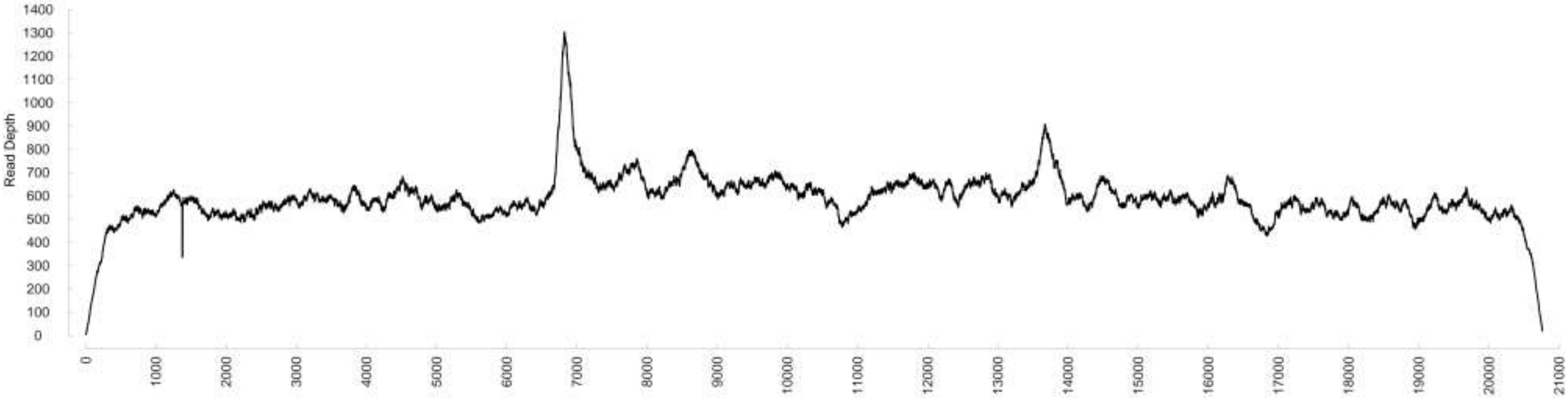

### Chromosome15

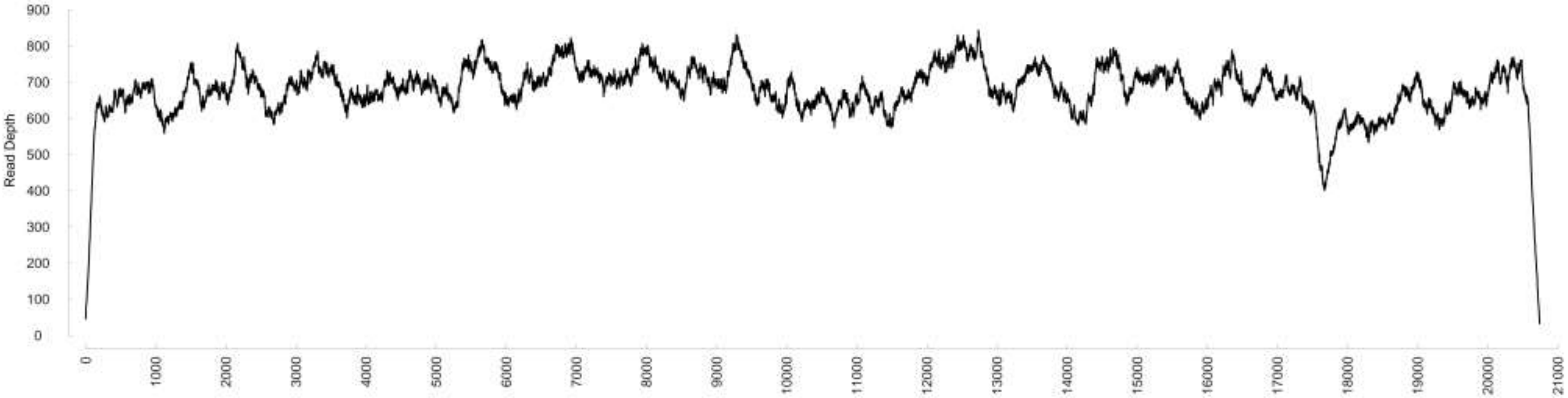

### Chromosome16

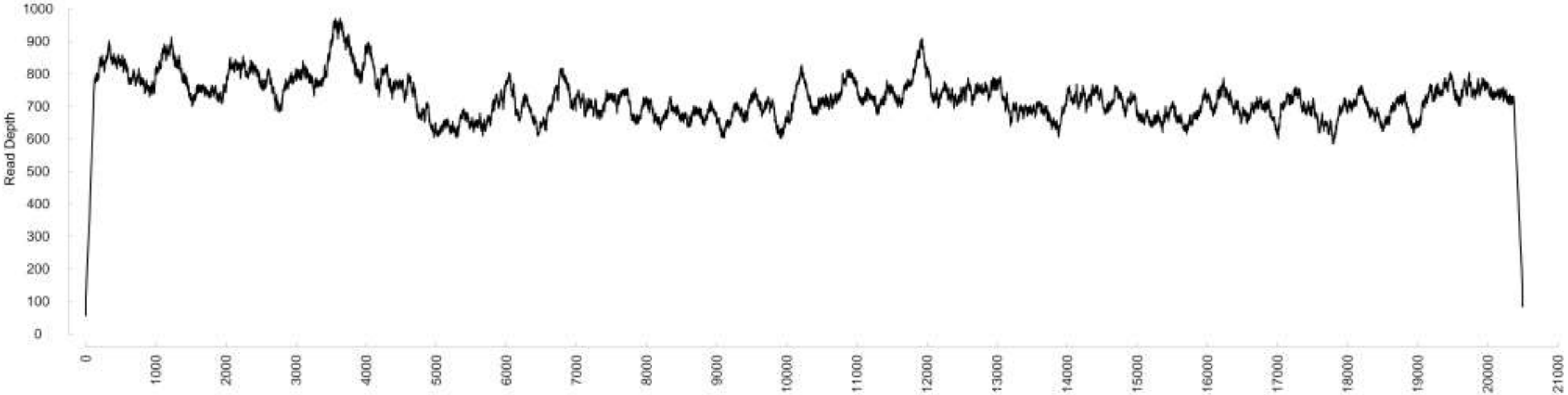

### Chromosome17

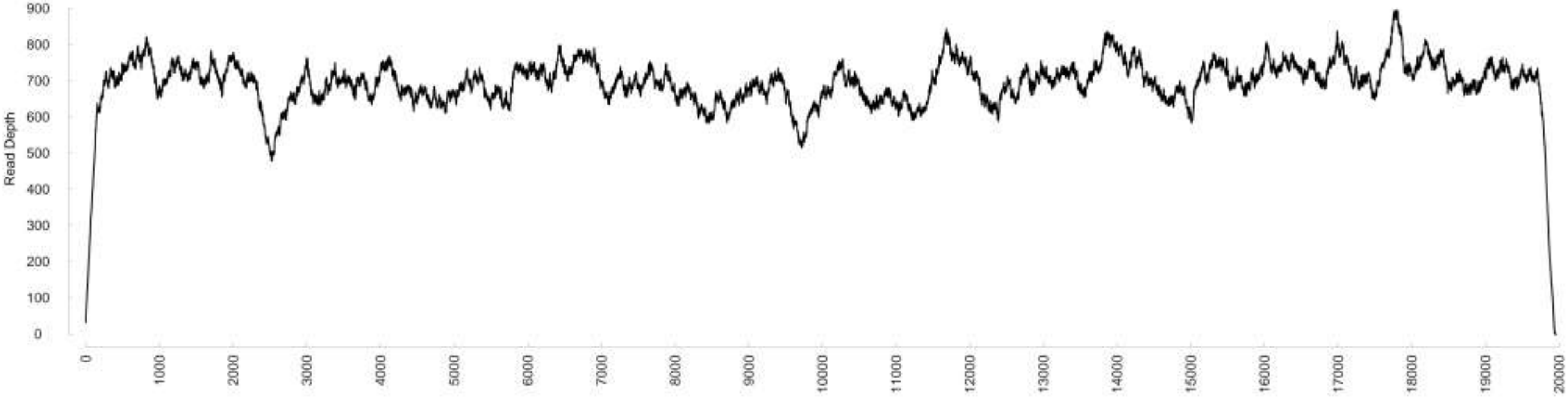

### Chromosome18

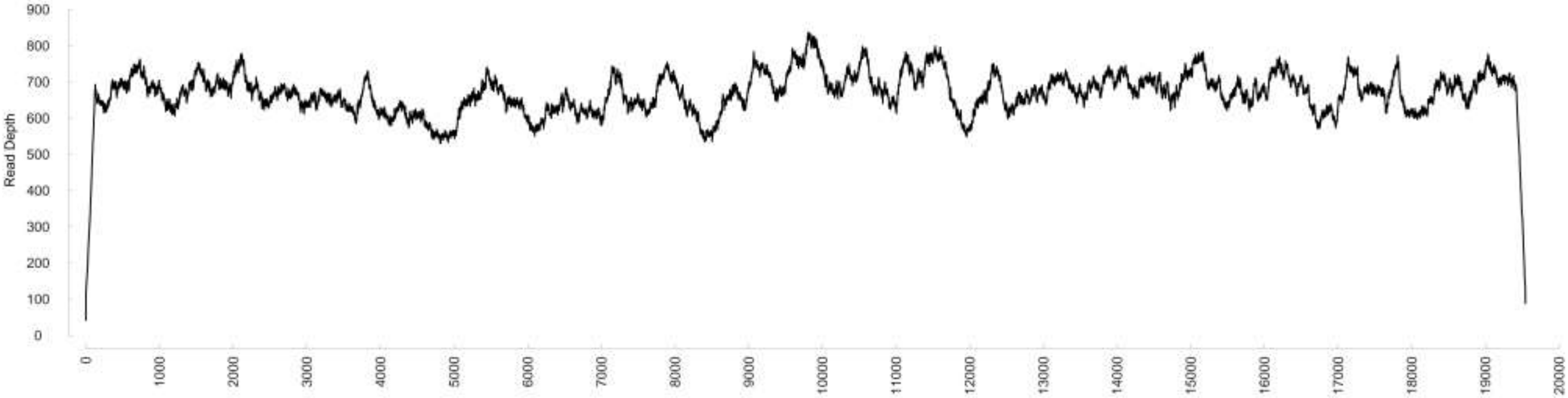

### Chromosome19

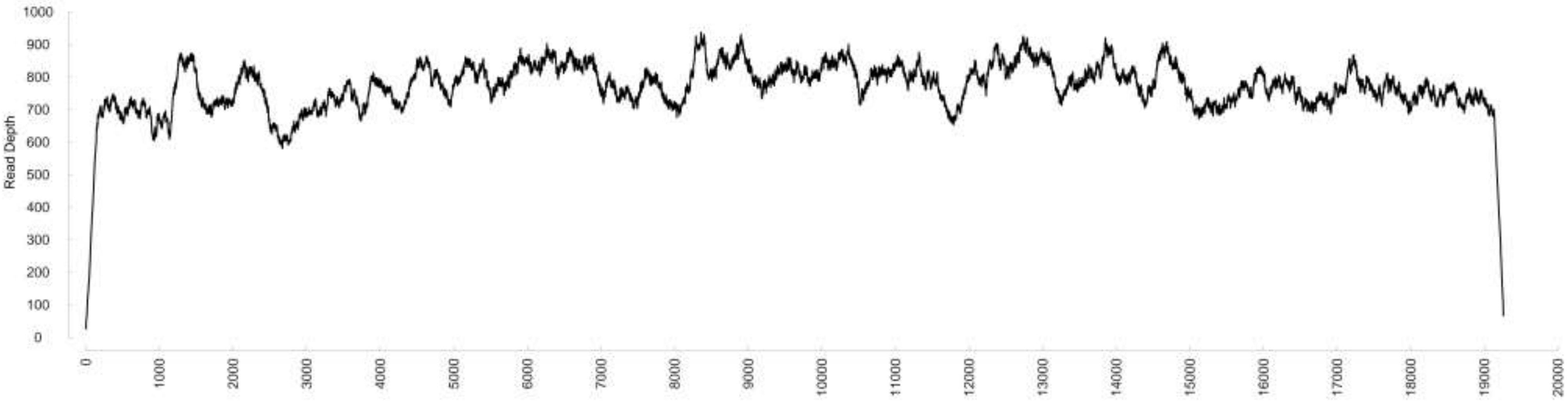

### Chromosome20

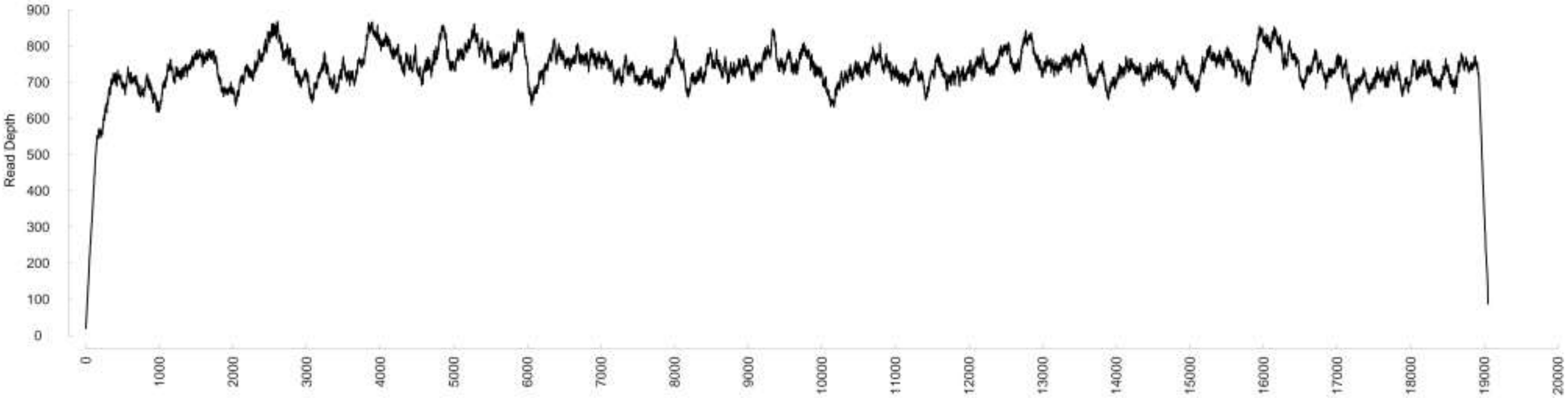

### Chromosome21

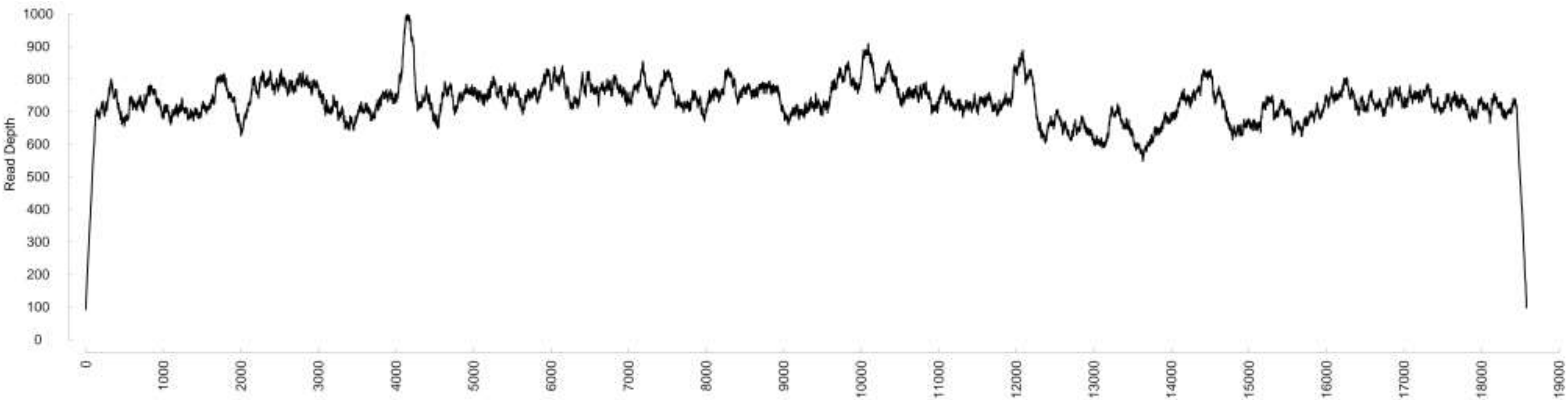

### Chromosome22

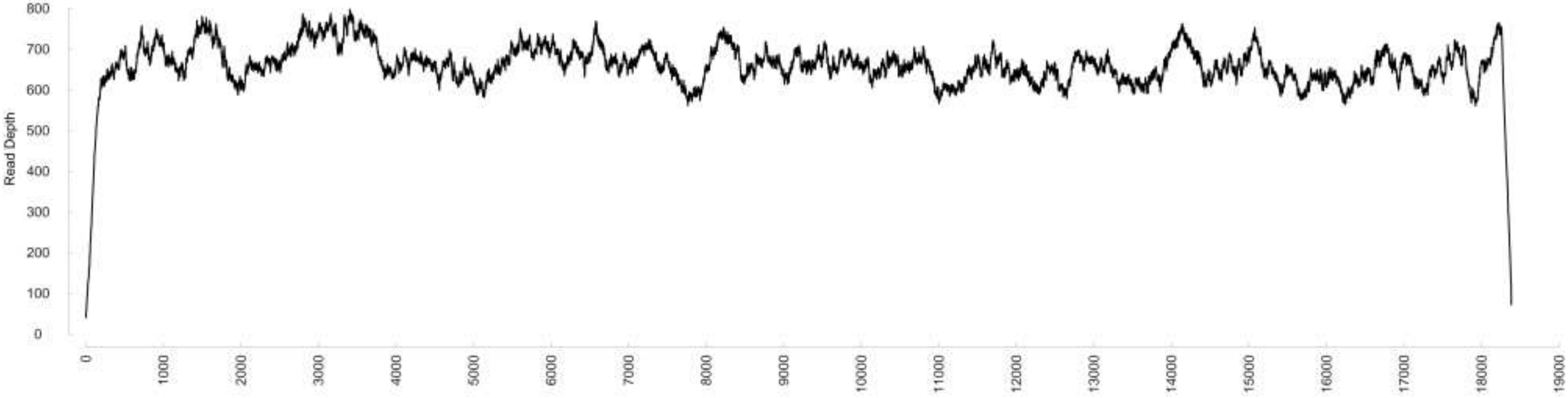

### Chromosome23

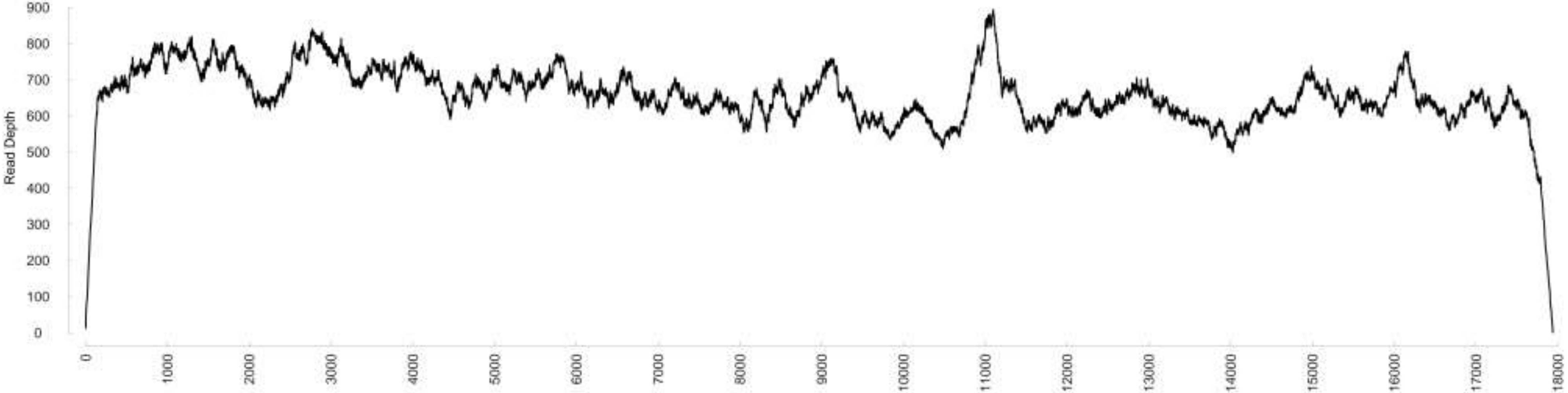

### Chromosome24

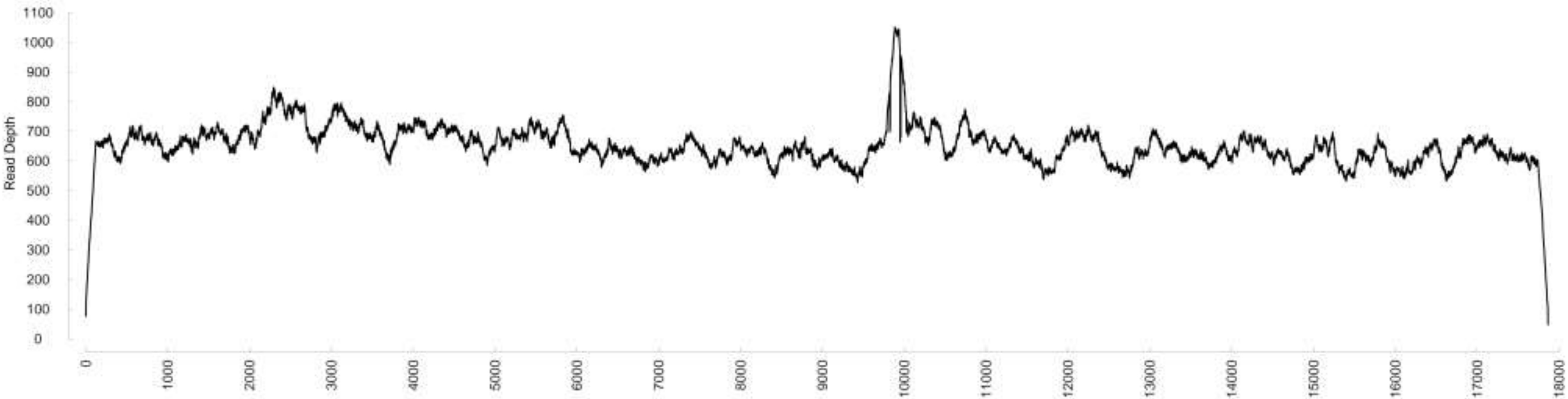

### Chromosome25

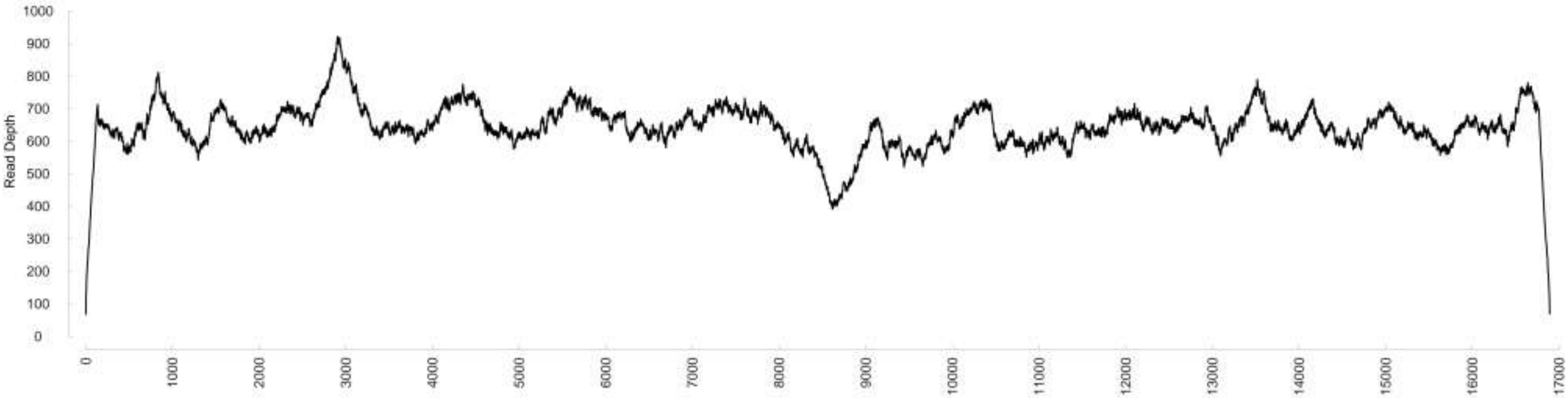

### Chromosome26

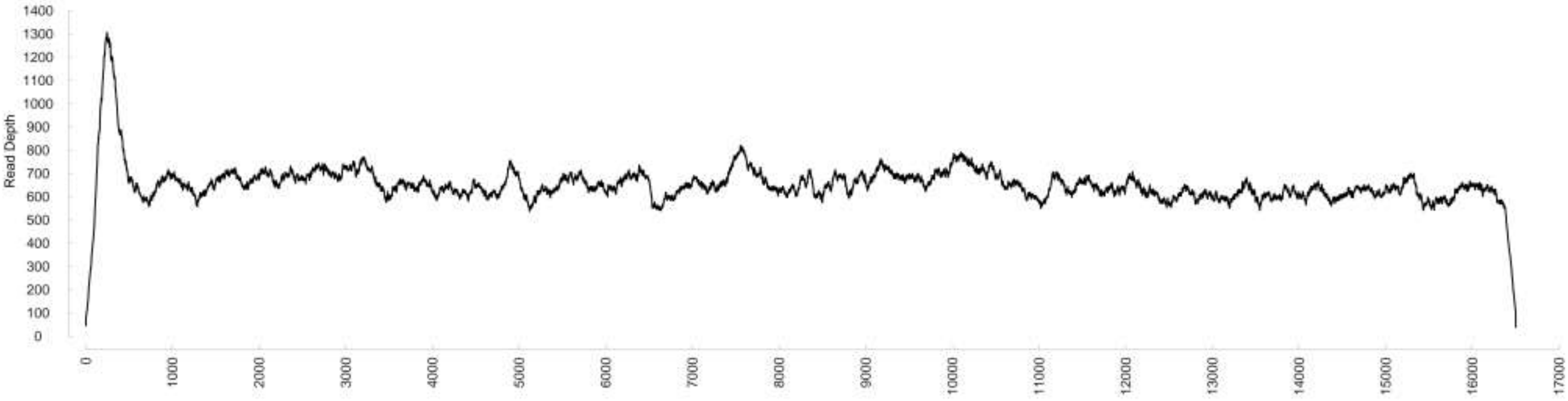

### Chromosome27

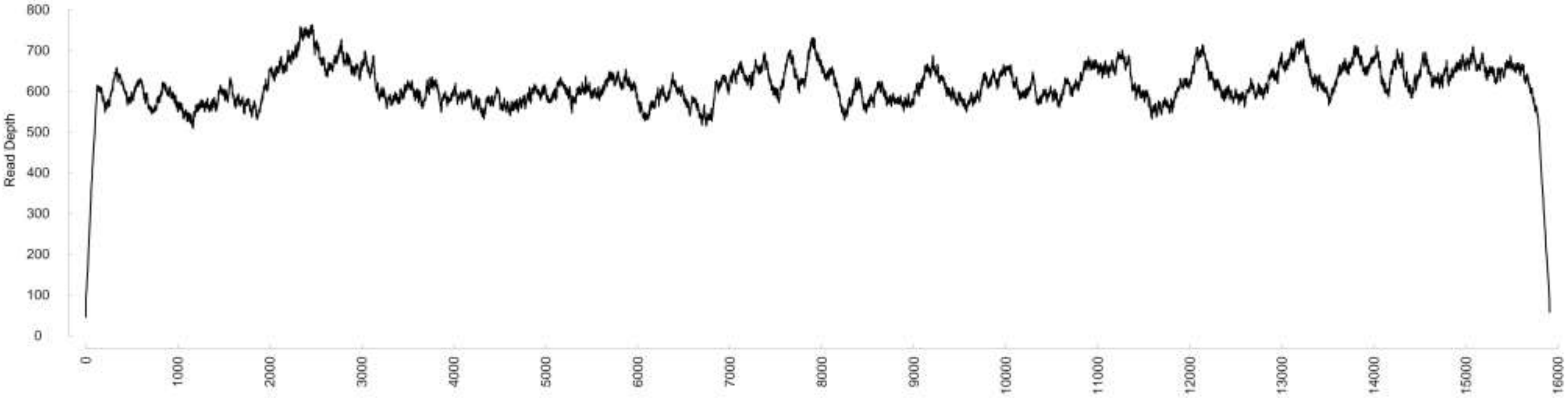

### Chromosome28

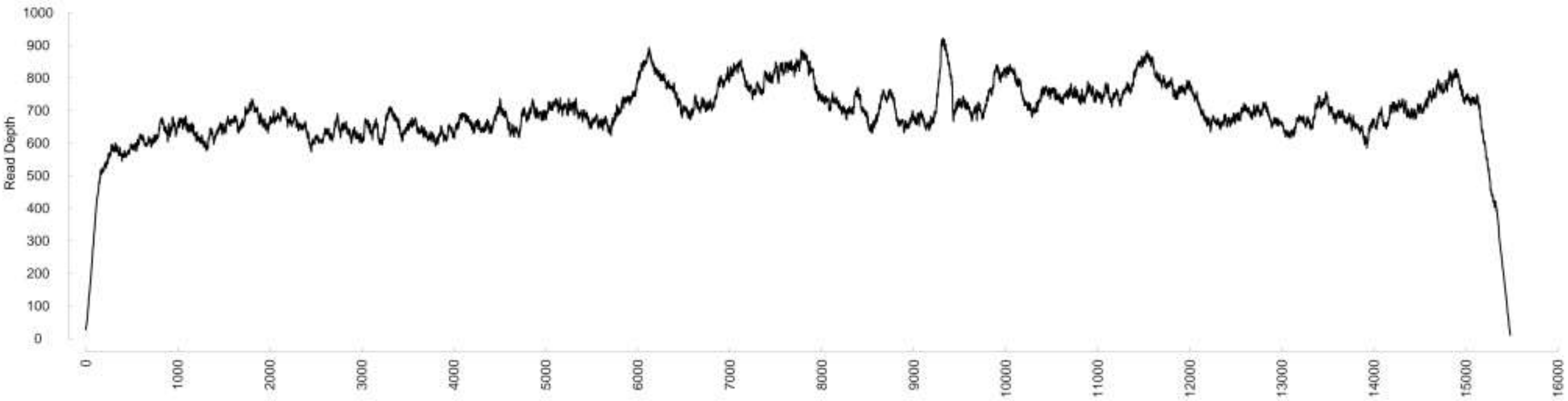

### Chromosome29

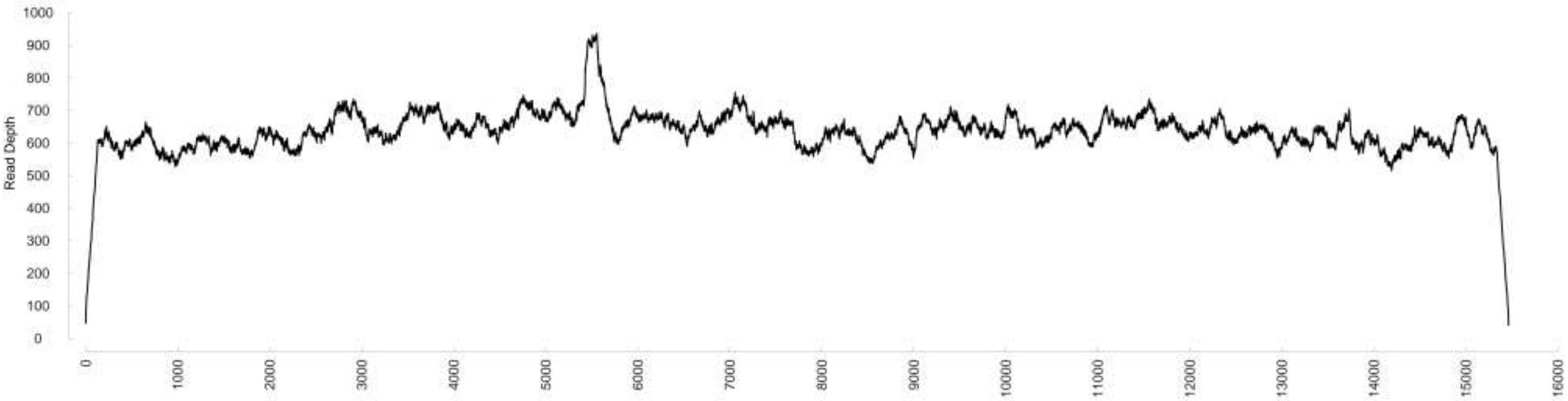

### Chromosome30

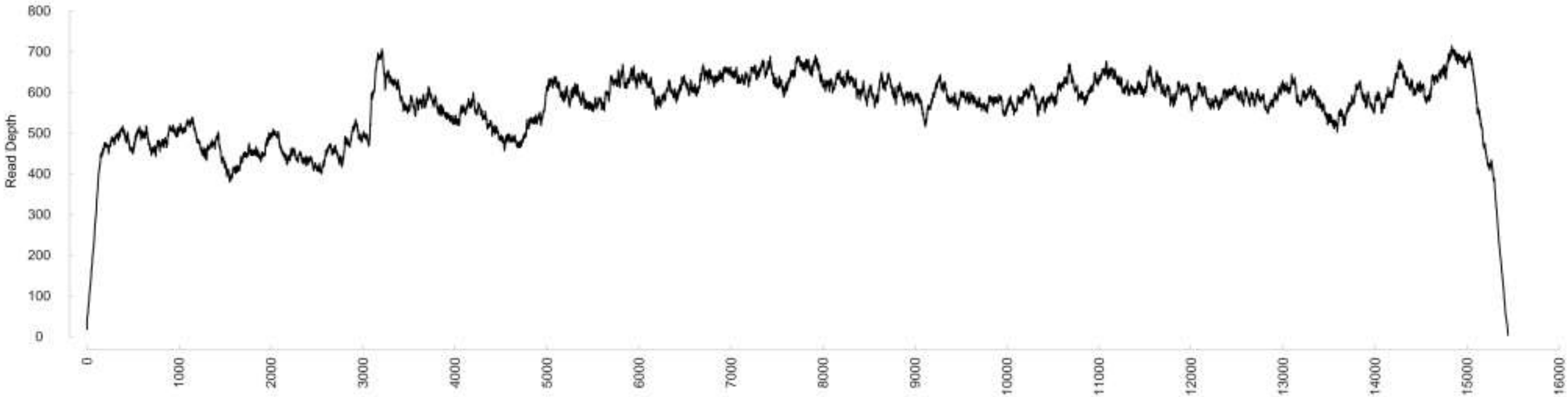

### Chromosome31

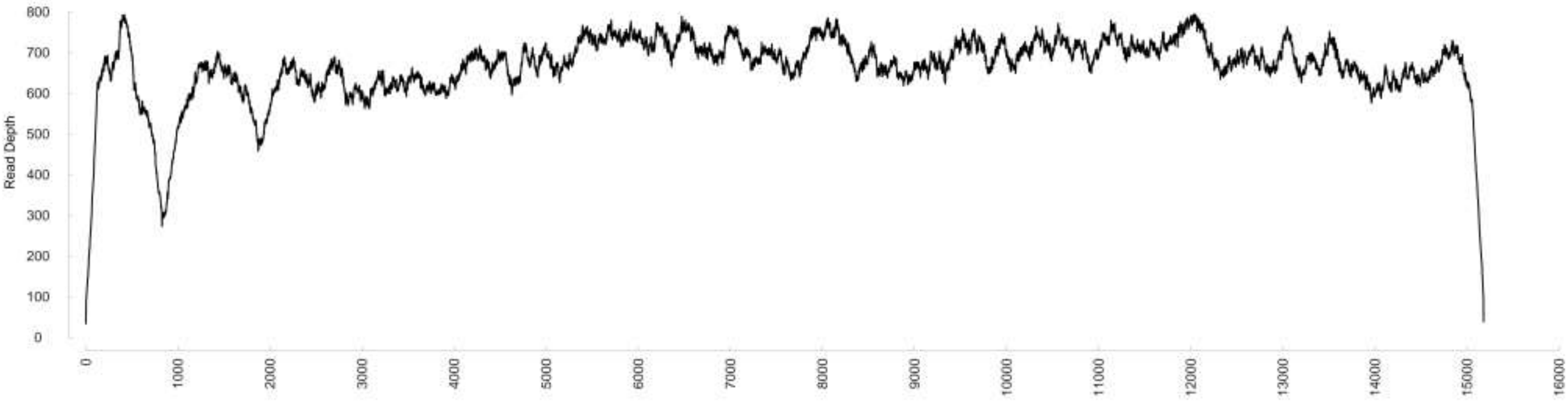

### Chromosome32

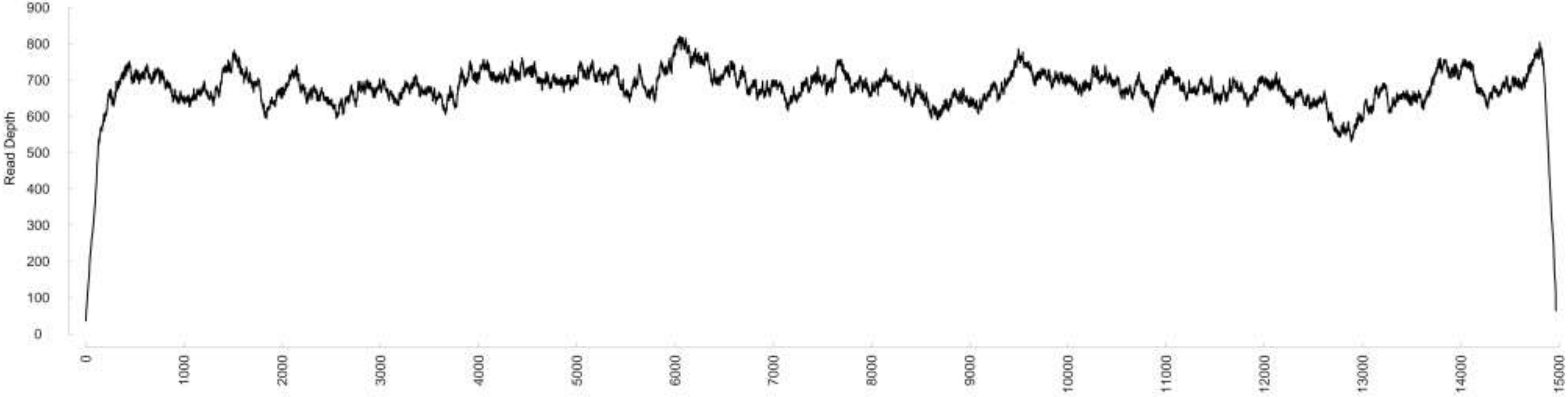

### Chromosome33

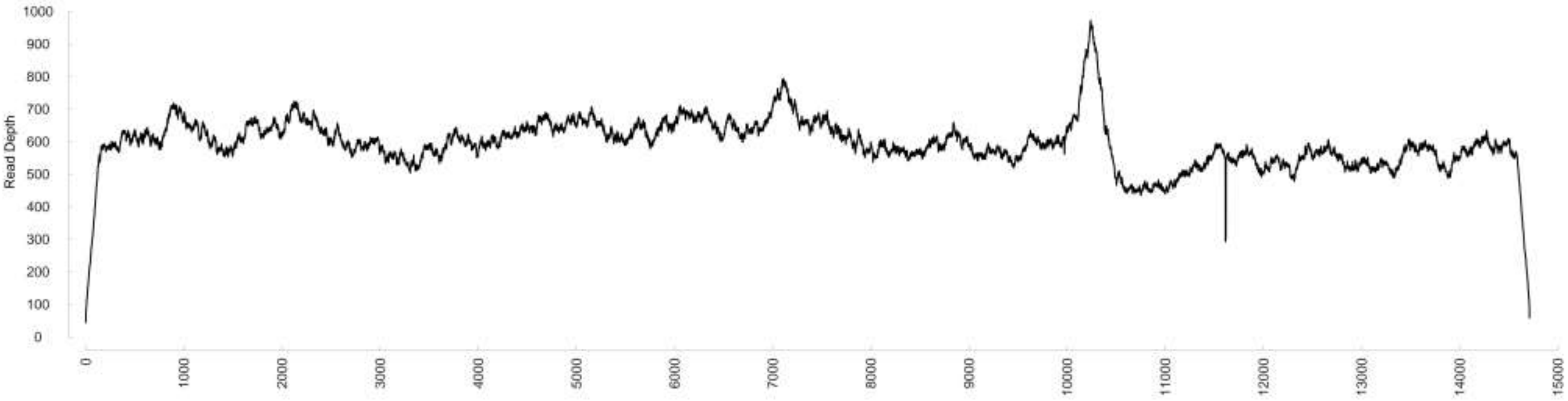

### Chromosome34

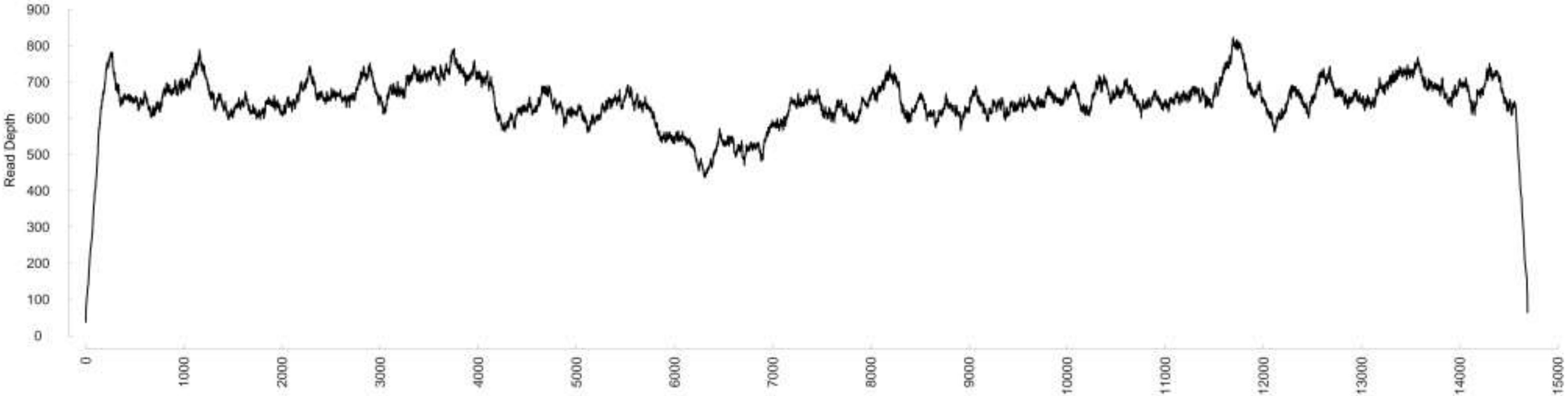

### Chromosome35

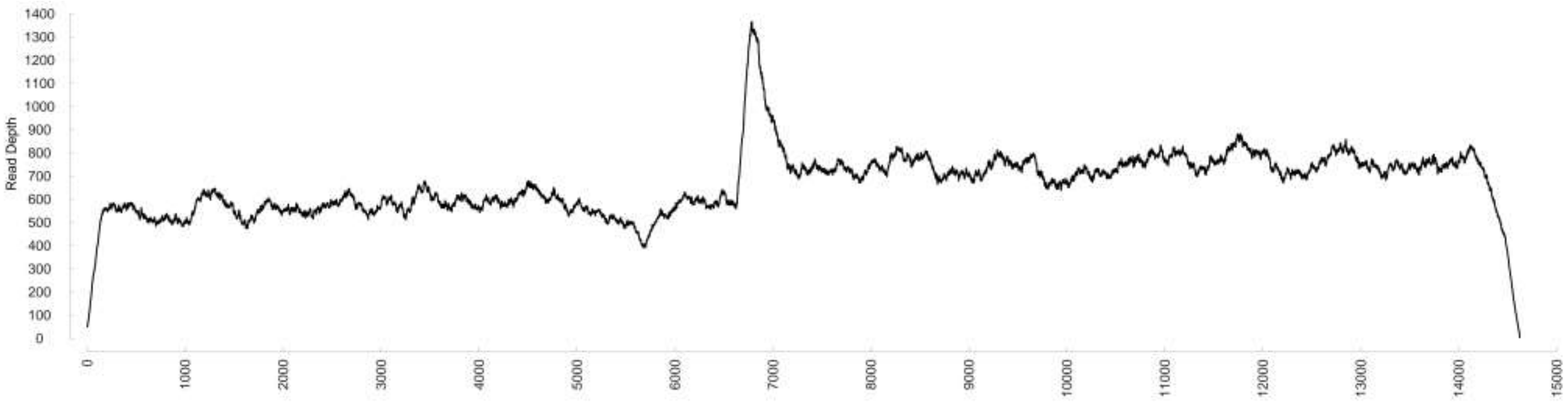

### Chromosome36

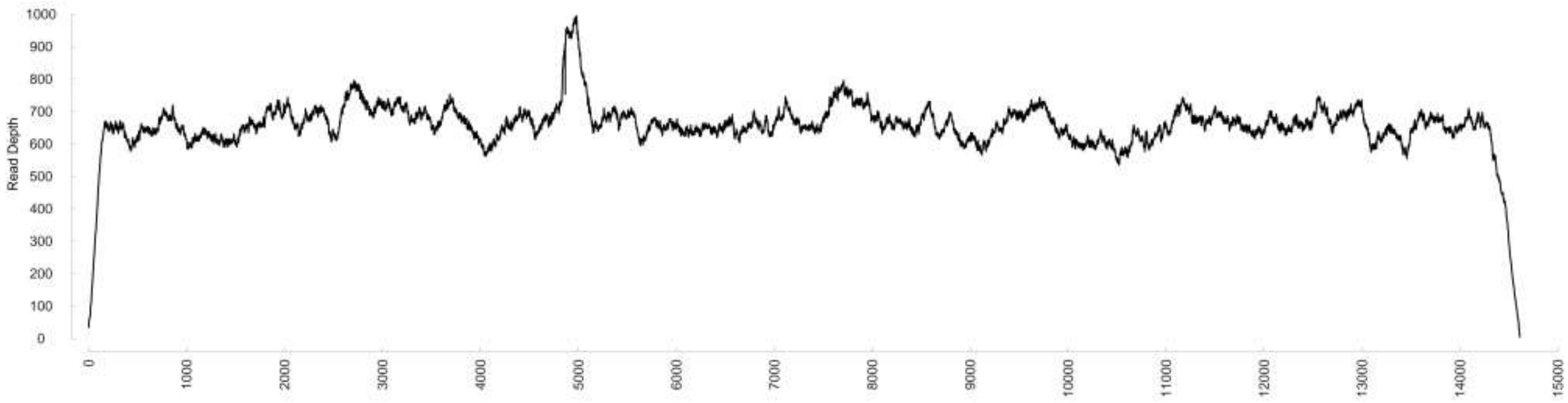

### Chromosome37

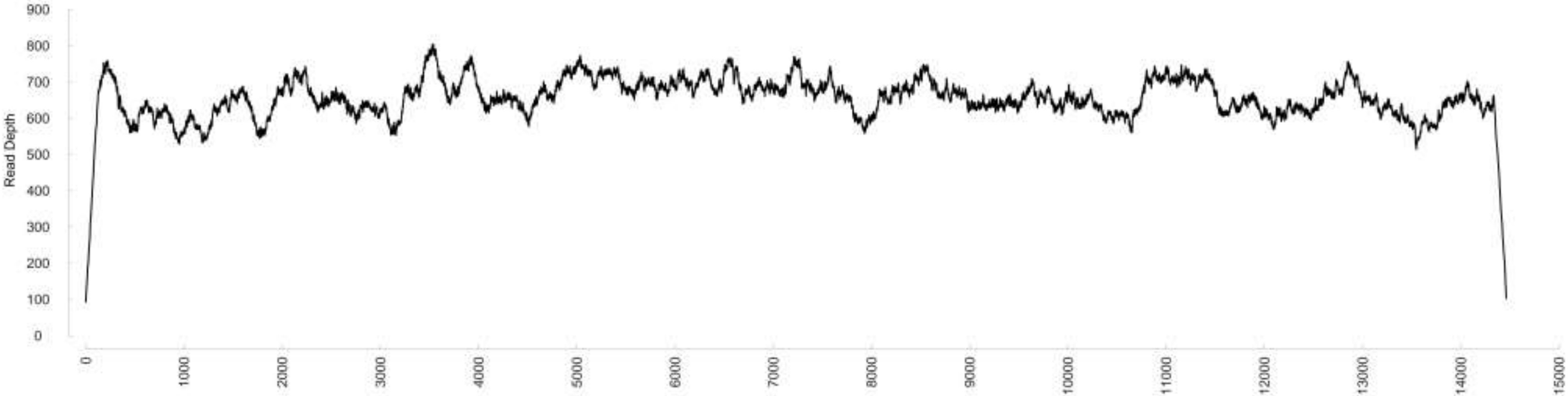

### Chromosome38

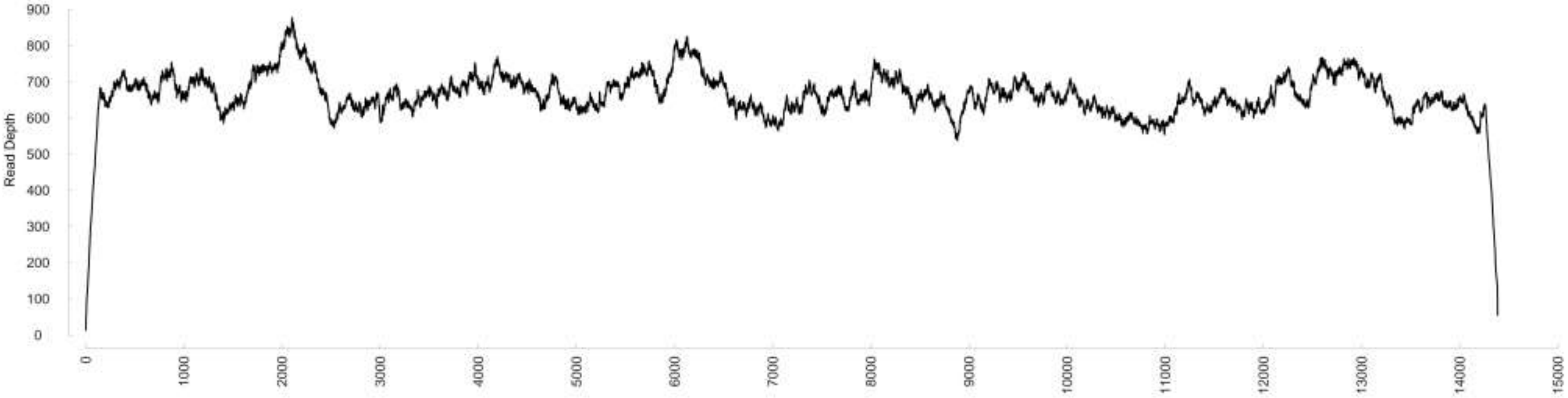

### Chromosome39

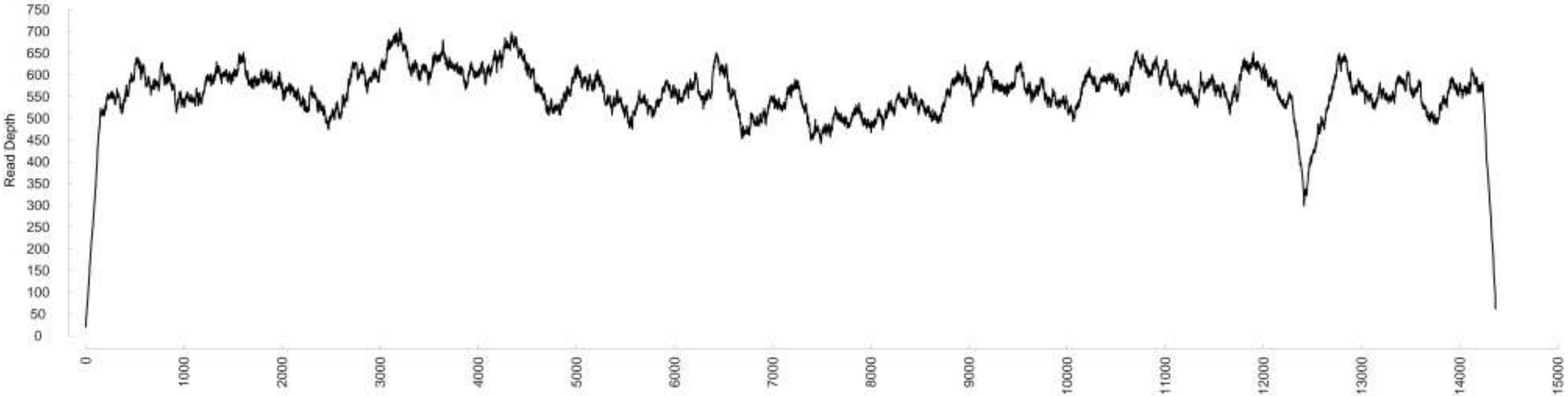

### Chromosome40

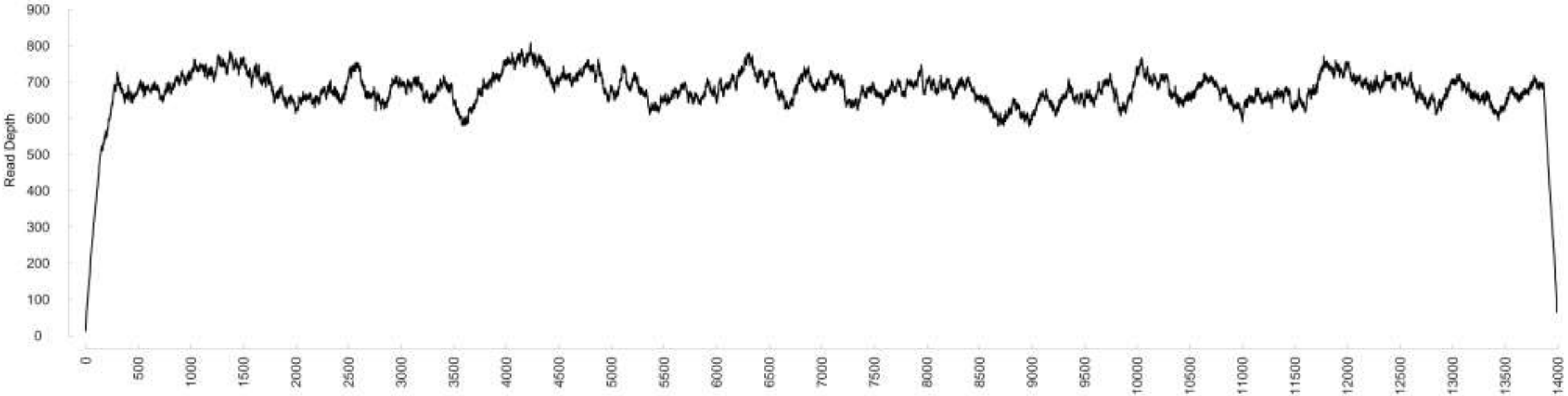

### Chromosome41

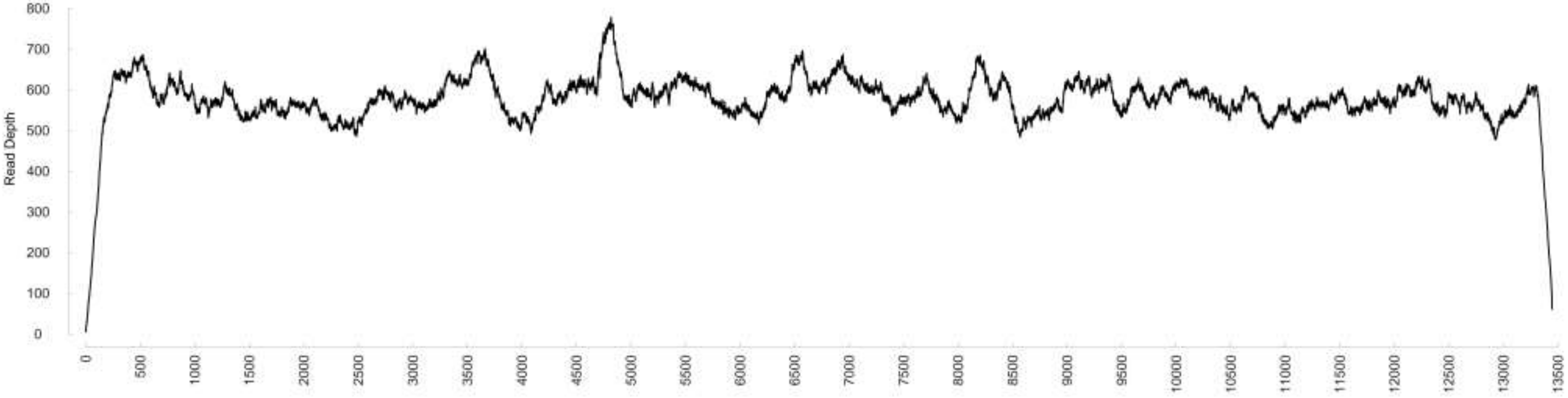

### Chromosome42

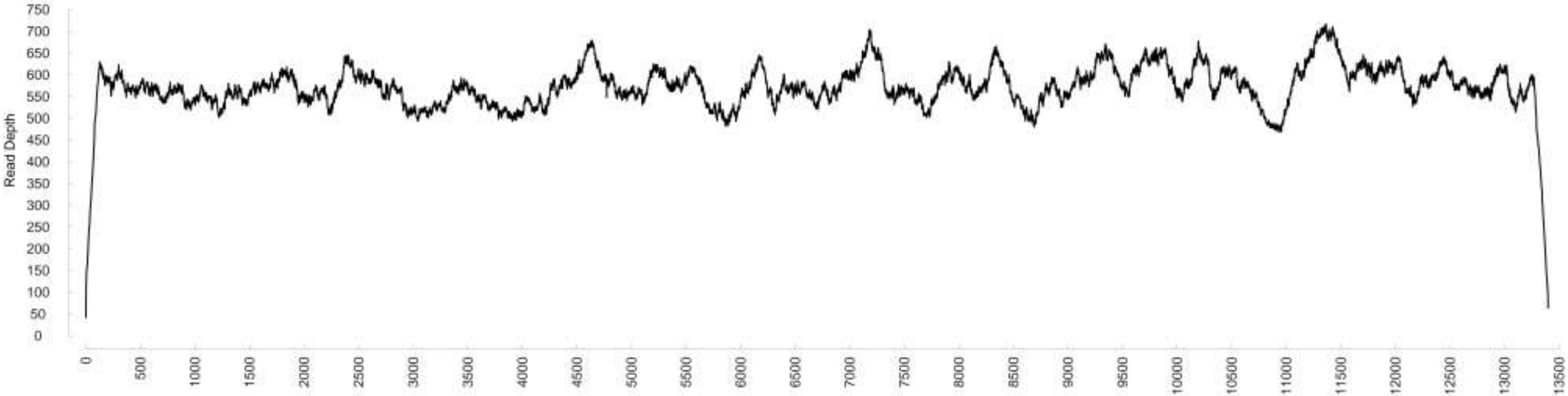

### Chromosome43

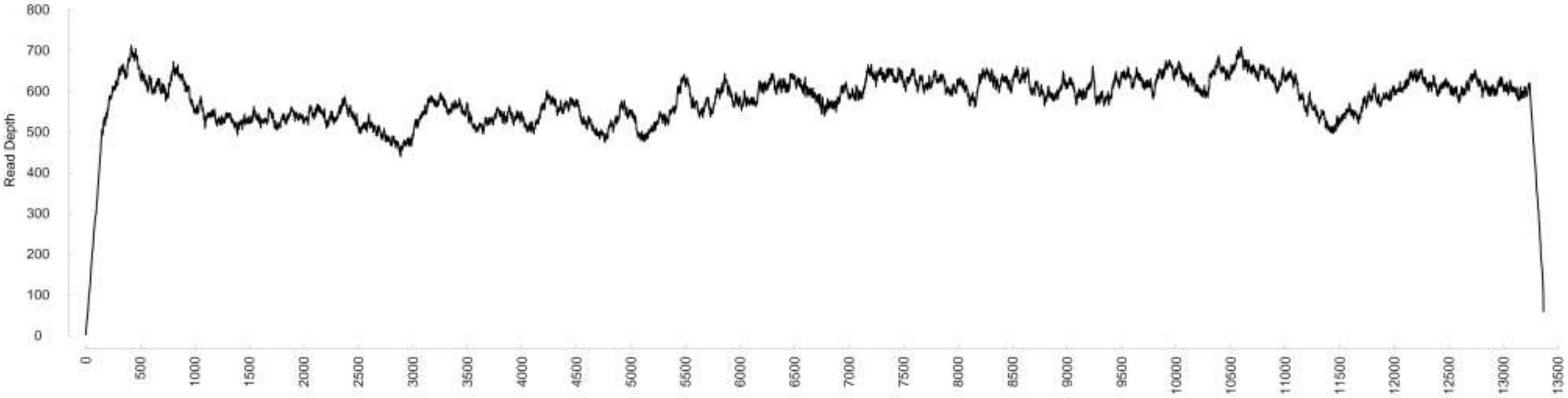

### Chromosome44

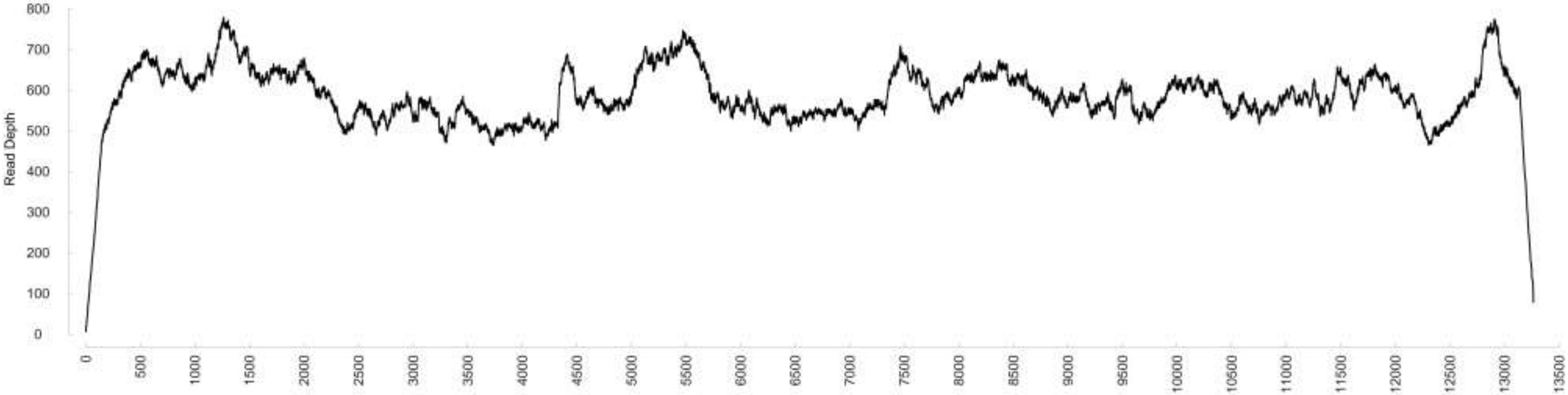

### Chromosome45

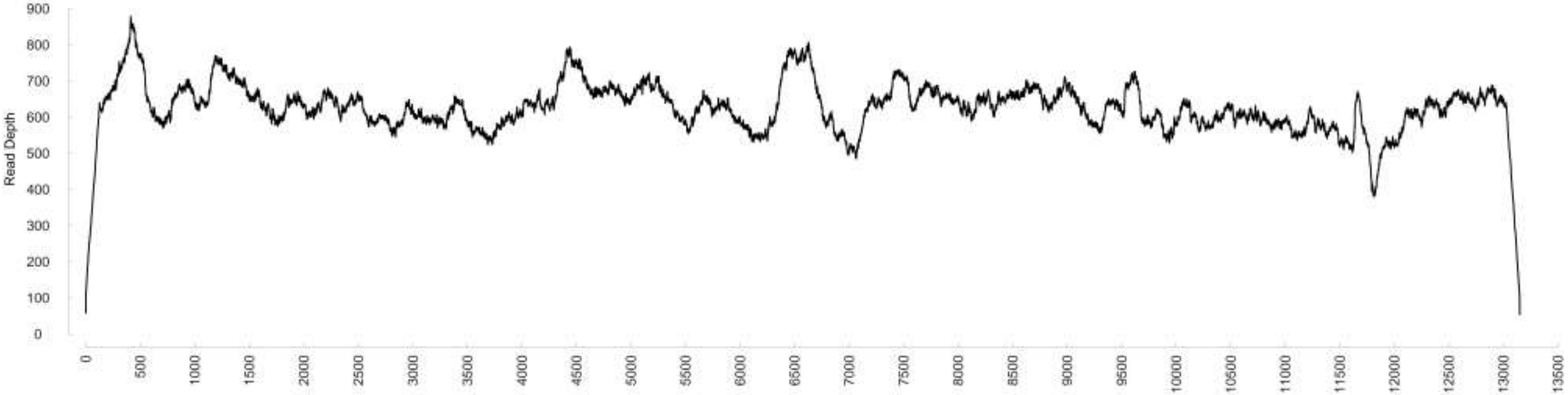

### Chromosome46

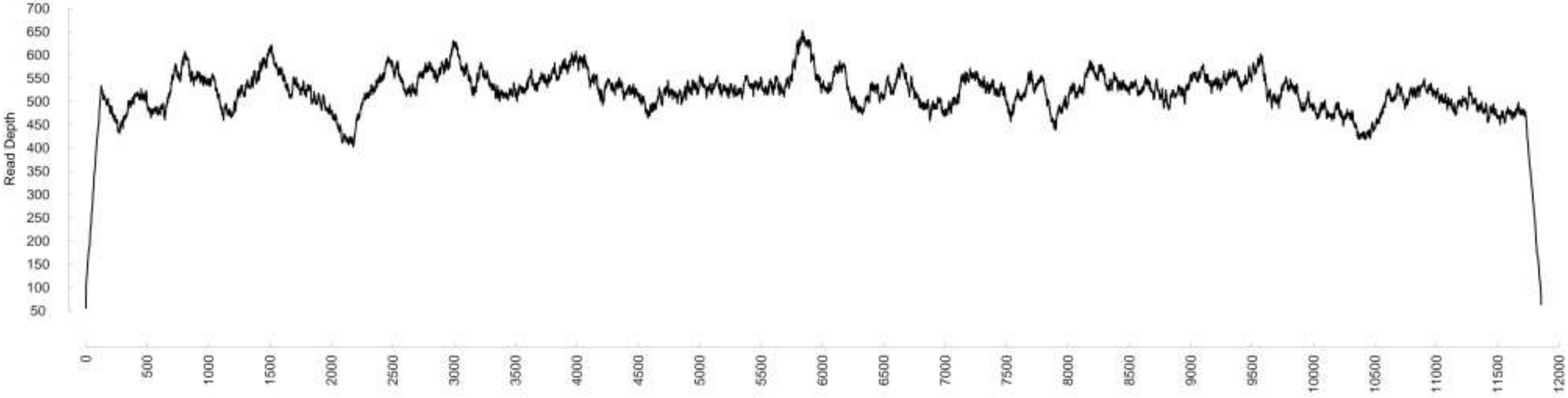

### Chromosome47

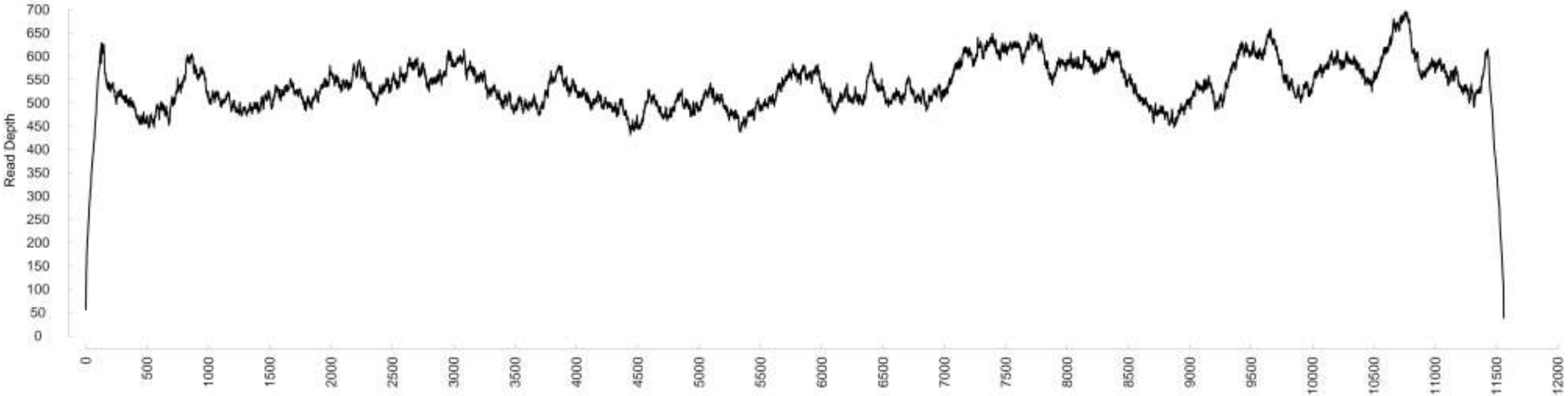

### Chromosome48

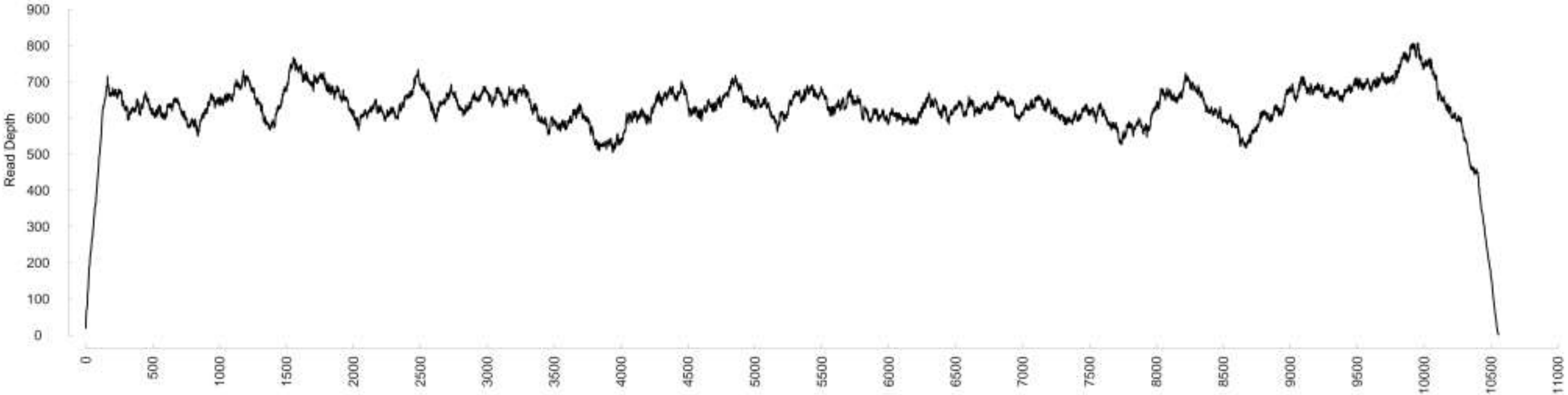

Chromosome49

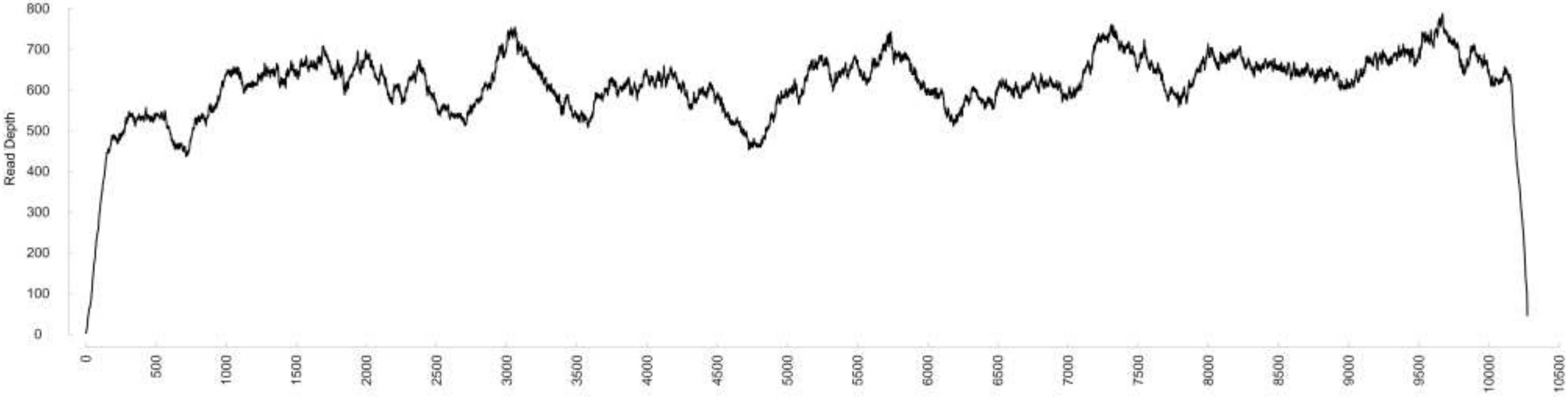

### Chromosome50

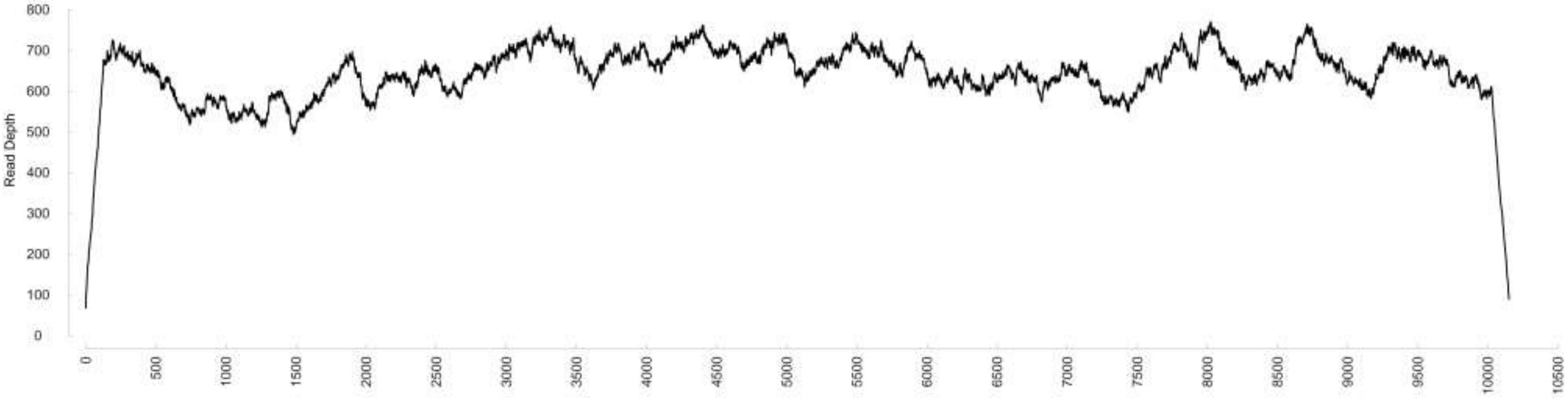

### Chromosome51

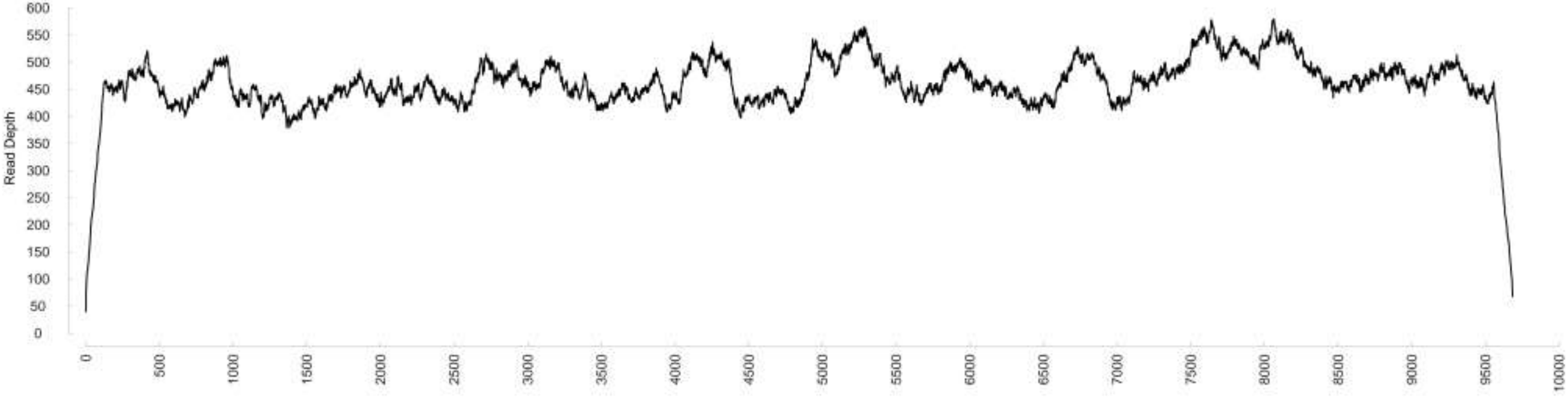

**Figure S2**. Maximum Likelihood (ML) phylogenetic analyses of coding sequences in the mtDNA of *Mitrastemon yamamotoi*. ML bootstrap support values ≥50% are shown. Scale bars correspond to substitutions per site.

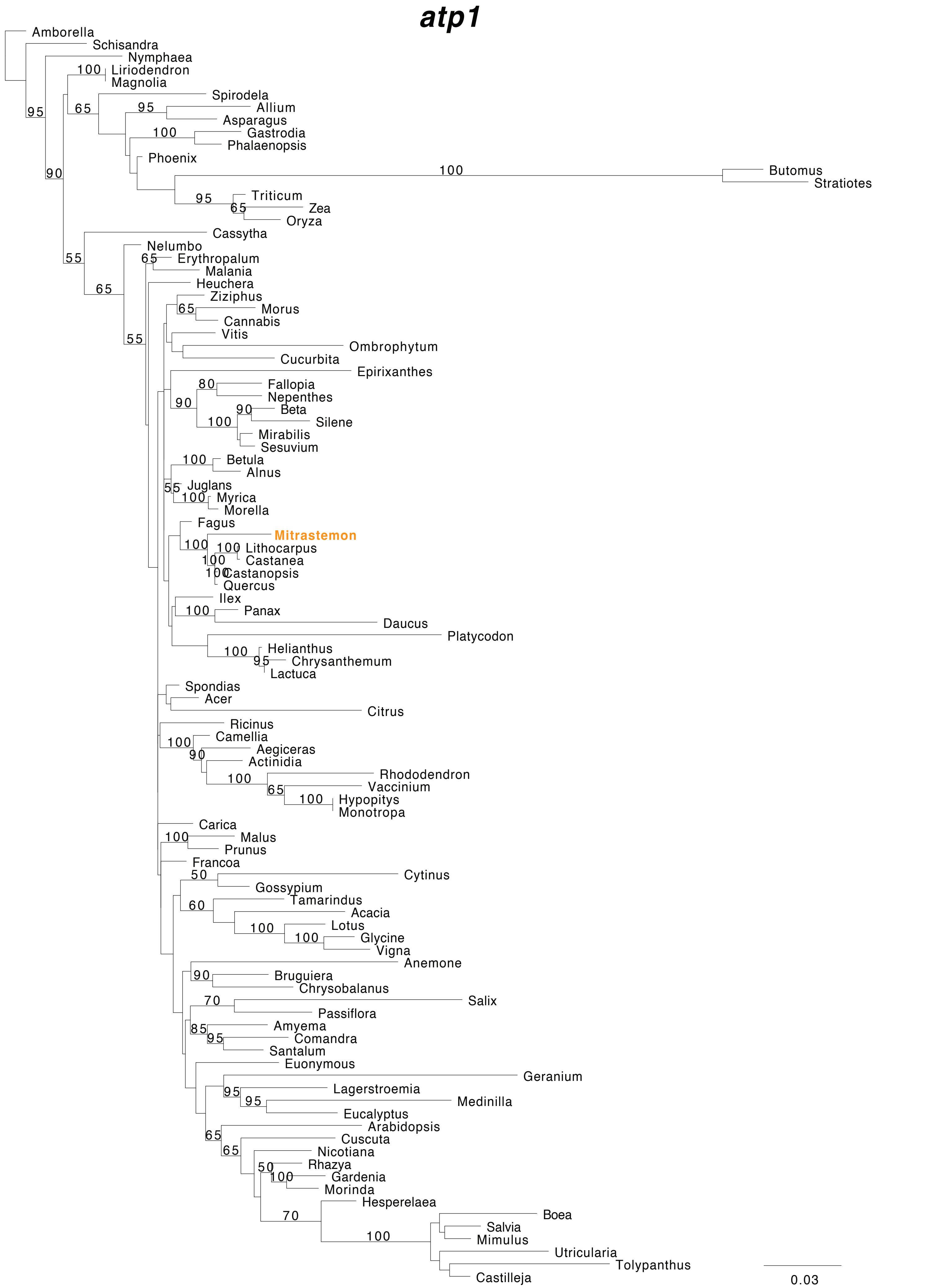

### atp4

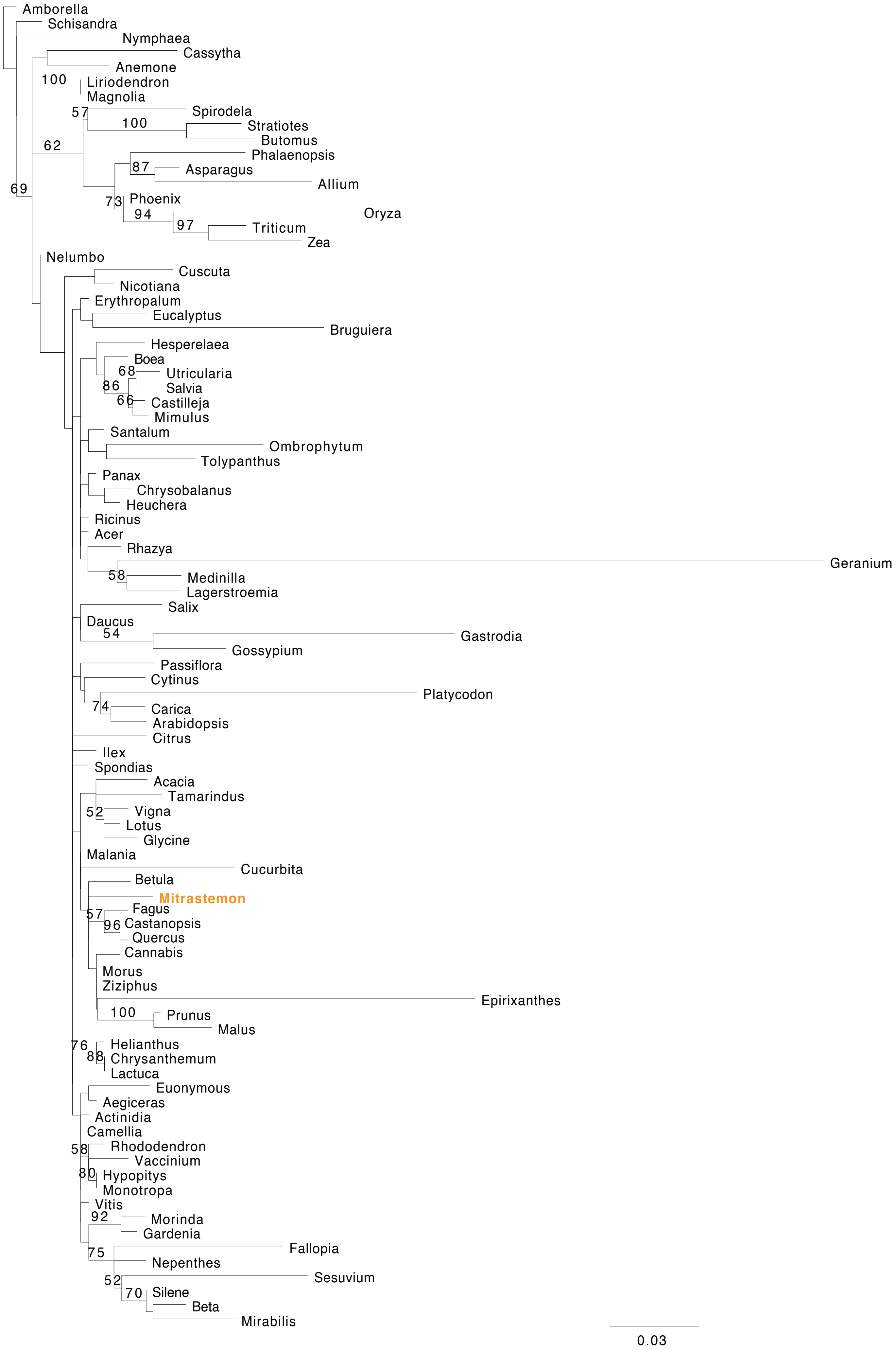
Amborella
Schisandra
Nymphaea
Cassytha
Anemone
100
Liriodendron
Magnolia
57
Spirodela
100
Stratiotes
Butomus
62
Phalaenopsis
87
Asparagus
Allium
69
73
Phoenix
94
Oryza
97
Triticum
Zea
Nelumbo
Cuscuta
Nicotiana
Erythropalum
Eucalyptus
Bruguiera
Hesperelaea
Boea
68
Utricularia
86
Salvia
66
Castilleja
Mimulus
Santalum
Ombrophytum
Tolypanthus
Panax
Chrysobalanus
Heuchera
Ricinus
Acer
Rhazya
Geranium
58
Medinilla
Lagerstroemia
Salix
Daucus
54
Gastrodia
Gossypium
Passiflora
Cytinus
Platycodon
74
Carica
Arabidopsis
Citrus
Ilex
Spondias
Acacia
Tamarindus
52
Vigna
Lotus
Glycine
Malania
Cucurbita
Betula
Mitrastemon
57
Fagus
96
Castanopsis
Quercus
Cannabis
Morus
Ziziphus
Epirixanthes
100
Prunus
Malus
76
Helianthus
88
Chrysanthemum
Lactuca
Euonymous
Aegiceras
Actinidia
Camellia
58
Rhododendron
Vaccinium
80
Hypopitys
Monotropa
Vitis
92
Morinda
Gardenia
75
Fallopia
Nepenthes
52
Sesuvium
70
Silene
Beta
Mirabilis
0.03

### *atp6*

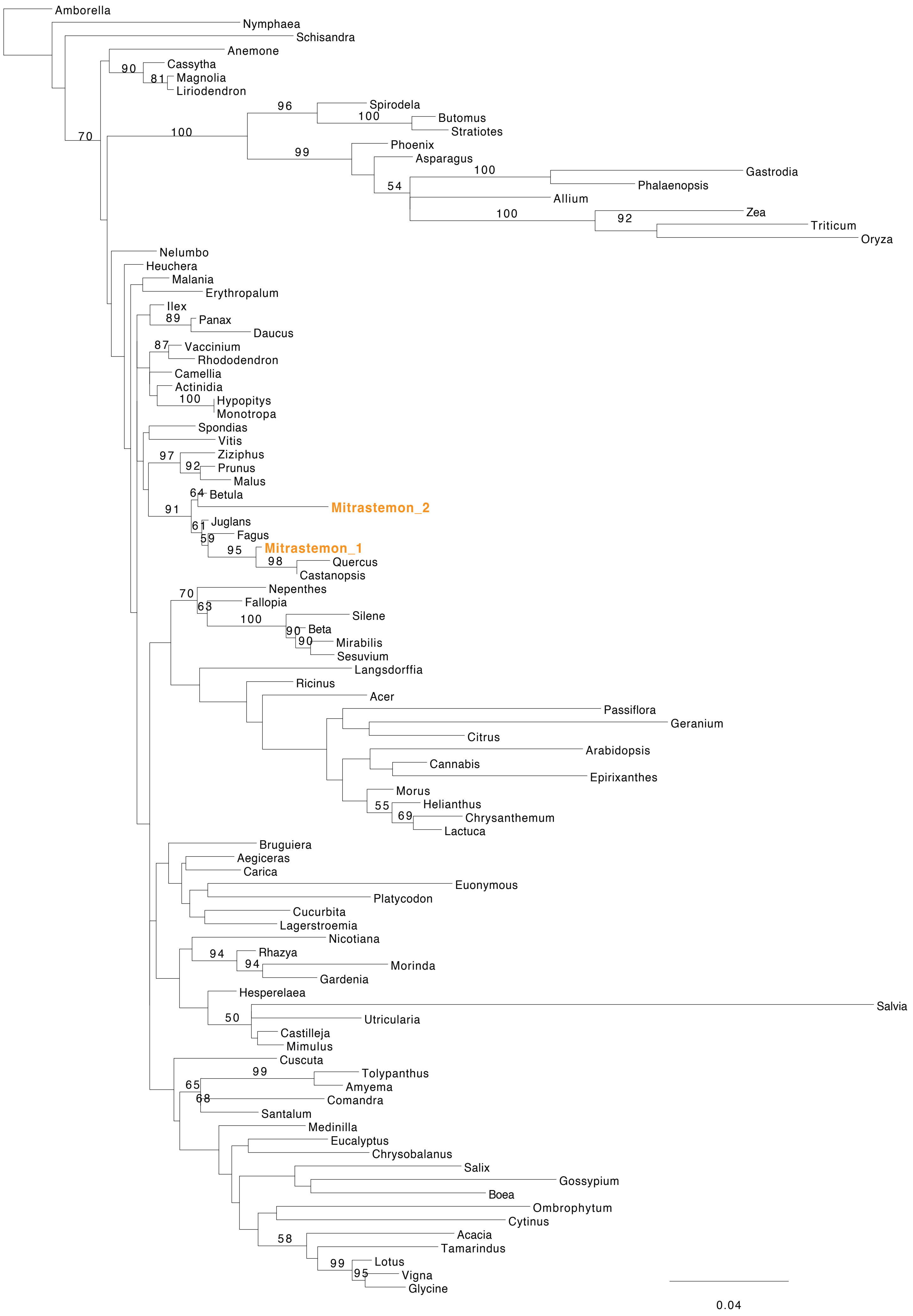
Amborella
Nymphaea
Schisandra
Anemone
90
Cassytha
81
Magnolia
Liriodendron
70
96
Spirodela
100
Butomus
Stratiotes
100
99
Phoenix
Asparagus
100
Gastrodia
Phalaenopsis
54
Allium
100
Zea
92
Triticum
Oryza
Nelumbo
Heuchera
Malania
Erythropalum
Ilex
89
Panax
Daucus
87
Vaccinium
Rhododendron
Camellia
Actinidia
100
Hypopitys
Monotropa
Spondias
Vitis
97
Ziziphus
92
Prunus
Malus
64
Betula
Mitrastemon_2
91
61
Juglans
59
Fagus
95
Mitrastemon_1
98
Quercus
Castanopsis
70
Nepenthes
63
Fallopia
100
Silene
90
Beta
90
Mirabilis
Sesuvium
Langsdorffia
Ricinus
Acer
Passiflora
Geranium
Citrus
Arabidopsis
Cannabis
Epirixanthes
Morus
55
Helianthus
69
Chrysanthemum
Lactuca
Bruguiera
Aegiceras
Carica
Euonymous
Platycodon
Cucurbita
Lagerstroemia
Nicotiana
94
Rhazya
94
Morinda
Gardenia
Hesperelaea
Salvia
50
Utricularia
Castilleja
Mimulus
Cuscuta
99
Tolypanthus
65
Amyema
68
Comandra
Santalum
Medinilla
Eucalyptus
Chrysobalanus
Salix
Gossypium
Boea
Ombrophytum
Cytinus
Acacia
58
Tamarindus
99
Lotus
95
Vigna
Glycine
0.04

### *atp8*

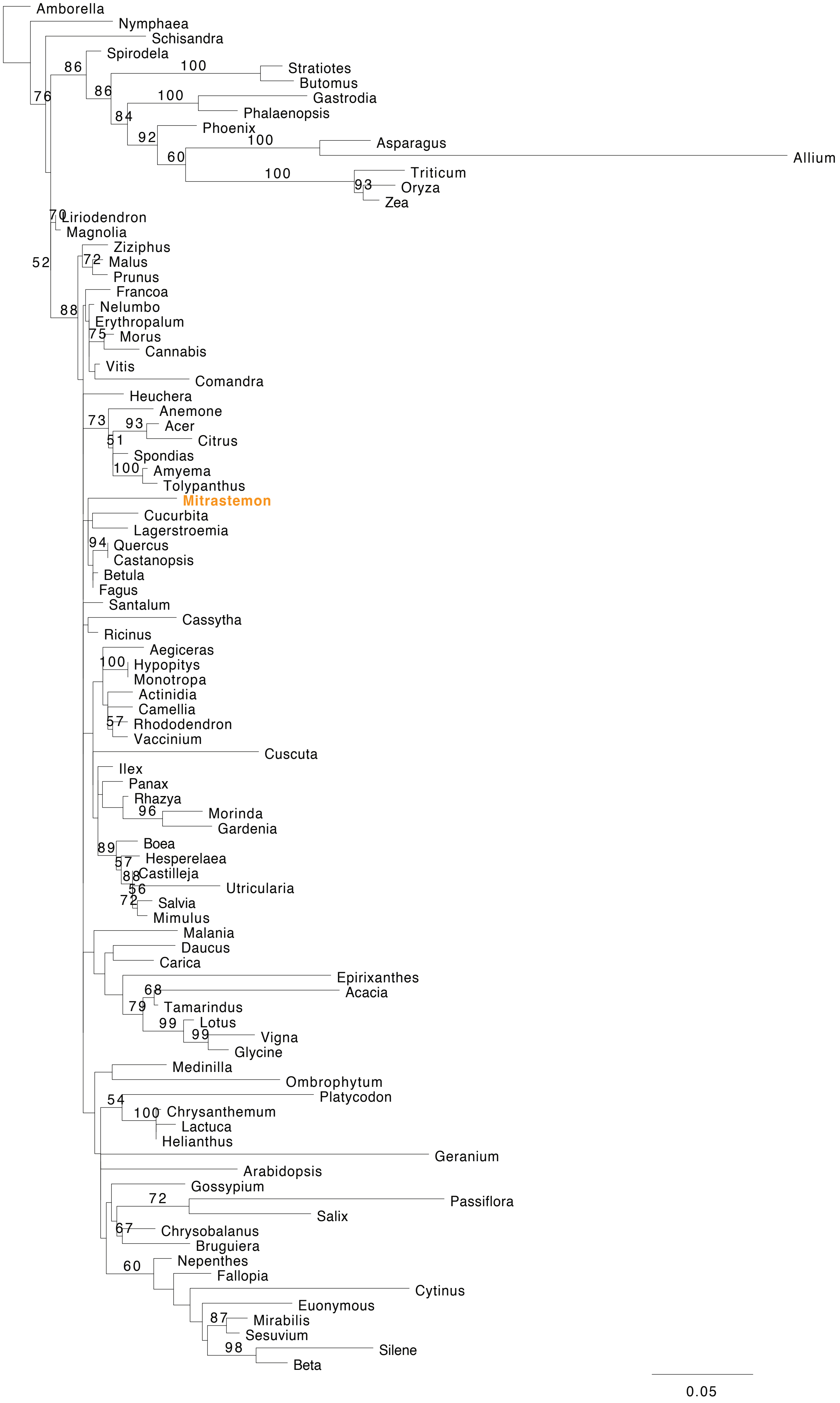

Amborella
Nymphaea
Schisandra
Spirodela
Stratiotes
Butomus
Gastrodia
Phalaenopsis
Phoenix
Asparagus
Allium
Triticum
Oryza
Zea
Liriodendron
Magnolia
Ziziphus
Malus
Prunus
Francoa
Nelumbo
Erythropalum
Morus
Cannabis
Vitis
Comandra
Heuchera
Anemone
Acer
Citrus
Spondias
Amyema
Tolypanthus
Mitrastemon
Cucurbita
Lagerstroemia
Quercus
Castanopsis
Betula
Fagus
Santalum
Cassytha
Ricinus
Aegiceras
Hypopitys
Monotropa
Actinidia
Camellia
Rhododendron
Vaccinium
Cuscuta
Ilex
Panax
Rhazya
Morinda
Gardenia
Boea
Hesperelaea
Castilleja
Utricularia
Salvia
Mimulus
Malania
Daucus
Carica
Epirixanthes
Acacia
Tamarindus
Lotus
Vigna
Glycine
Medinilla
Ombrophytum
Platycodon
Chrysanthemum
Lactuca
Helianthus
Geranium
Arabidopsis
Gossypium
Passiflora
Salix
Chrysobalanus
Bruguiera
Nepenthes
Fallopia
Cytinus
Euonymous
Mirabilis
Sesuvium
Silene
Beta
0.05

### *atp9*

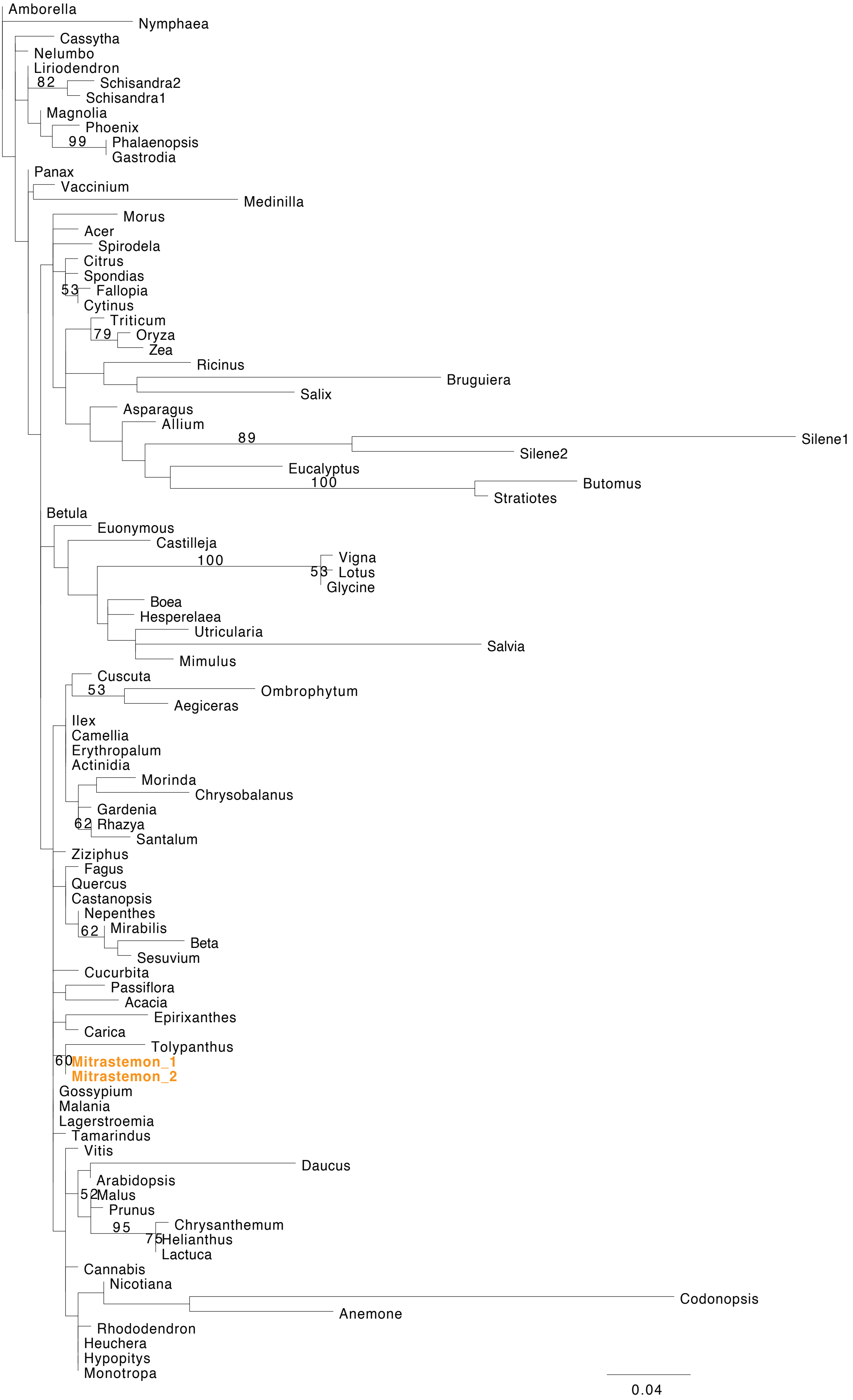

Amborella
Nymphaea
Cassytha
Nelumbo
Liriodendron
82
Schisandra2
Schisandra1
Magnolia
Phoenix
99
Phalaenopsis
Gastrodia
Panax
Vaccinium
Medinilla
Morus
Acer
Spirodela
Citrus
Spondias
53
Fallopia
Cytinus
Triticum
79
Oryza
Zea
Ricinus
Bruguiera
Salix
Asparagus
Allium
89
Silene1
Silene2
Eucalyptus
100
Butomus
Stratiotes
Betula
Euonymous
Castilleja
100
Vigna
53
Lotus
Glycine
Boea
Hesperelaea
Utricularia
Salvia
Mimulus
Cuscuta
53
Ombrophytum
Aegiceras
Ilex
Camellia
Erythropalum
Actinidia
Morinda
Chrysobalanus
Gardenia
62
Rhazya
Santalum
Ziziphus
Fagus
Quercus
Castanopsis
Nepenthes
62
Mirabilis
Beta
Sesuvium
Cucurbita
Passiflora
Acacia
Epirixanthes
Carica
Tolypanthus
60
Mitrastemon_1
Mitrastemon_2
Gossypium
Malania
Lagerstroemia
Tamarindus
Vitis
Daucus
Arabidopsis
52
Malus
Prunus
95
Chrysanthemum
75
Helianthus
Lactuca
Cannabis
Nicotiana
Codonopsis
Anemone
Rhododendron
Heuchera
Hypopitys
Monotropa
0.04

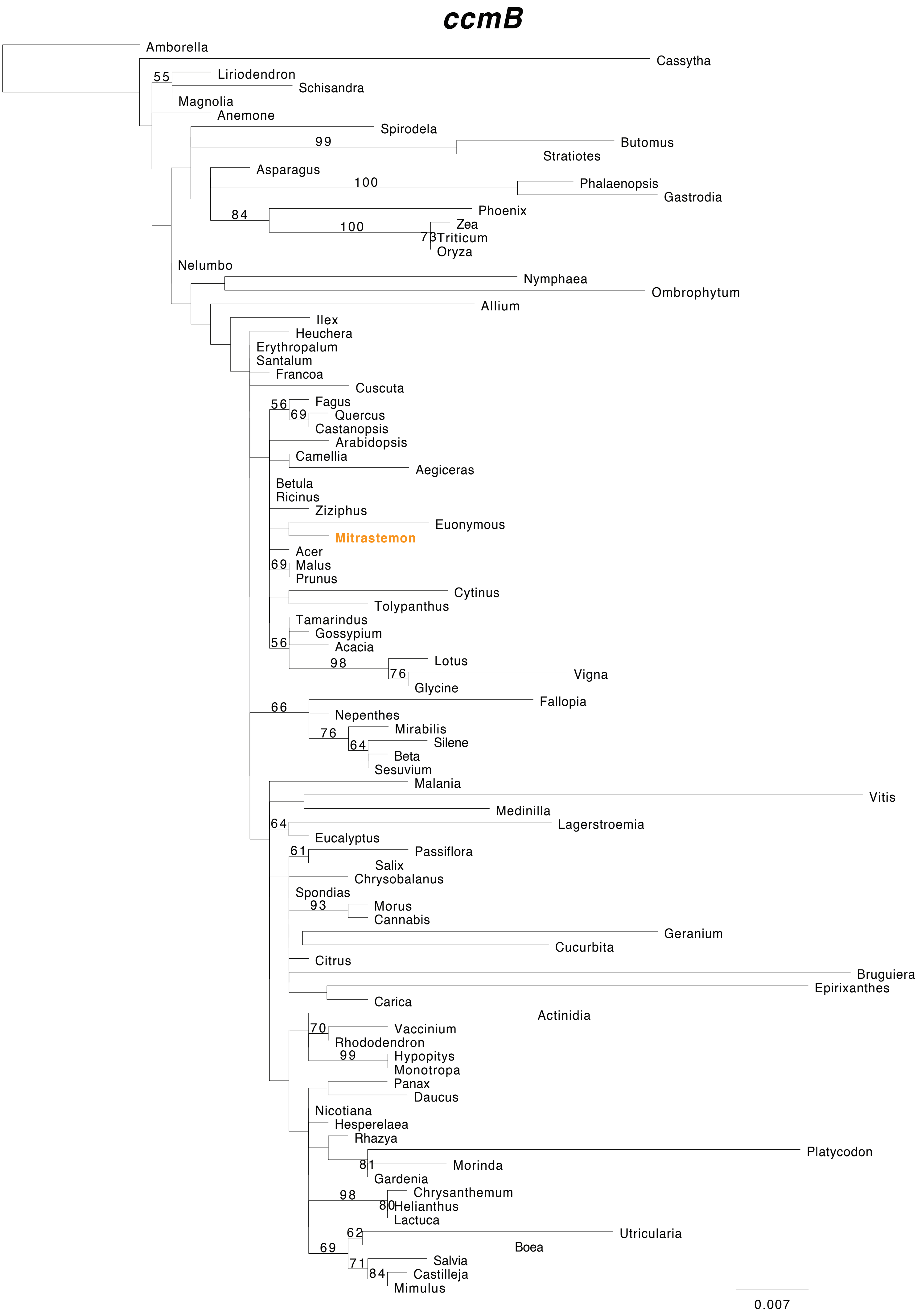
ccmB
Amborella
Cassytha
55
Liriodendron
Schisandra
Magnolia
Anemone
Spirodela
99
Butomus
Stratiotes
Asparagus
100
Phalaenopsis
Gastrodia
84
Phoenix
100
Zea
73
Triticum
Oryza
Nelumbo
Nymphaea
Ombrophytum
Allium
Ilex
Heuchera
Erythropalum
Santalum
Francoa
Cuscuta
56
Fagus
69
Quercus
Castanopsis
Arabidopsis
Camellia
Aegiceras
Betula
Ricinus
Ziziphus
Euonymous
Mitrastemon
Acer
69
Malus
Prunus
Cytinus
Tolypanthus
Tamarindus
Gossypium
56
Acacia
98
Lotus
76
Vigna
Glycine
66
Fallopia
Nepenthes
76
Mirabilis
64
Silene
Beta
Sesuvium
Malania
Vitis
Medinilla
64
Lagerstroemia
Eucalyptus
61
Passiflora
Salix
Chrysobalanus
Spondias
93
Morus
Cannabis
Geranium
Cucurbita
Citrus
Bruguiera
Epirixanthes
Carica
Actinidia
70
Vaccinium
Rhododendron
99
Hypopitys
Monotropa
Panax
Daucus
Nicotiana
Hesperelaea
Rhazya
Platycodon
81
Morinda
Gardenia
98
Chrysanthemum
80
Helianthus
Lactuca
62
Utricularia
69
Boea
71
Salvia
84
Castilleja
Mimulus
0.007

***ccmC***

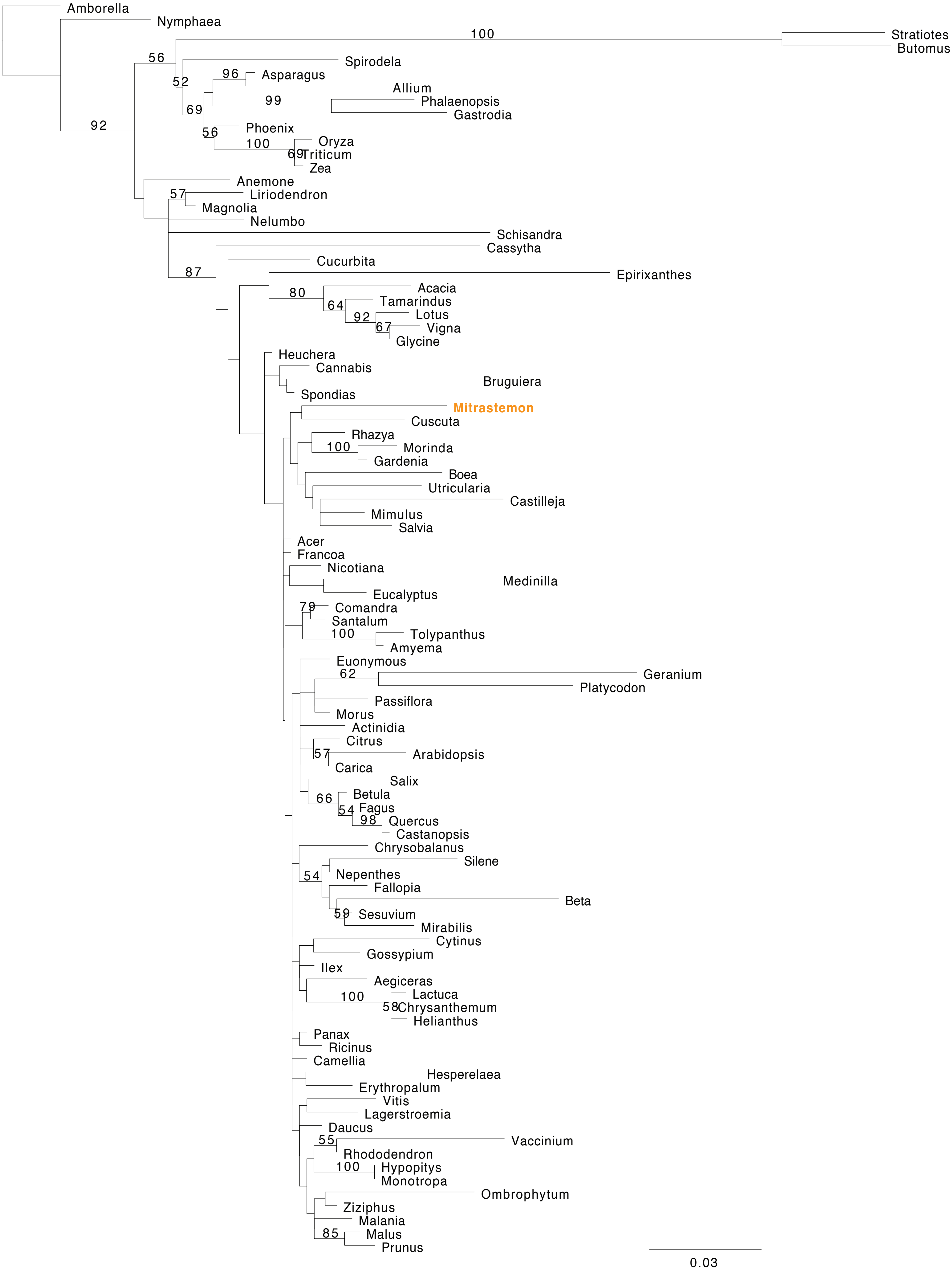
Amborella
Nymphaea
100
Stratiotes
Butomus
56
Spirodela
52
96
Asparagus
Allium
99
Phalaenopsis
Gastrodia
69
92
Phoenix
56
100
Oryza
69
Triticum
Zea
Anemone
57
Liriodendron
Magnolia
Nelumbo
Schisandra
Cassytha
87
Cucurbita
Epirixanthes
80
Acacia
64
Tamarindus
92
Lotus
67
Vigna
Glycine
Heuchera
Cannabis
Bruguiera
Spondias
Mitrastemon
Cuscuta
Rhazya
100
Morinda
Gardenia
Boea
Utricularia
Castilleja
Mimulus
Salvia
Acer
Francoa
Nicotiana
Medinilla
Eucalyptus
79
Comandra
Santalum
100
Tolypanthus
Amyema
Euonymous
62
Geranium
Platycodon
Passiflora
Morus
Actinidia
Citrus
57
Arabidopsis
Carica
Salix
66
Betula
54
Fagus
98
Quercus
Castanopsis
Chrysobalanus
Silene
54
Nepenthes
Fallopia
Beta
59
Sesuvium
Mirabilis
Cytinus
Gossypium
Ilex
Aegiceras
100
Lactuca
58
Chrysanthemum
Helianthus
Panax
Ricinus
Camellia
Hesperelaea
Erythropalum
Vitis
Lagerstroemia
Daucus
55
Vaccinium
Rhododendron
100
Hypopitys
Monotropa
Ombrophytum
Ziziphus
Malania
85
Malus
Prunus
0.03

### ccmFC

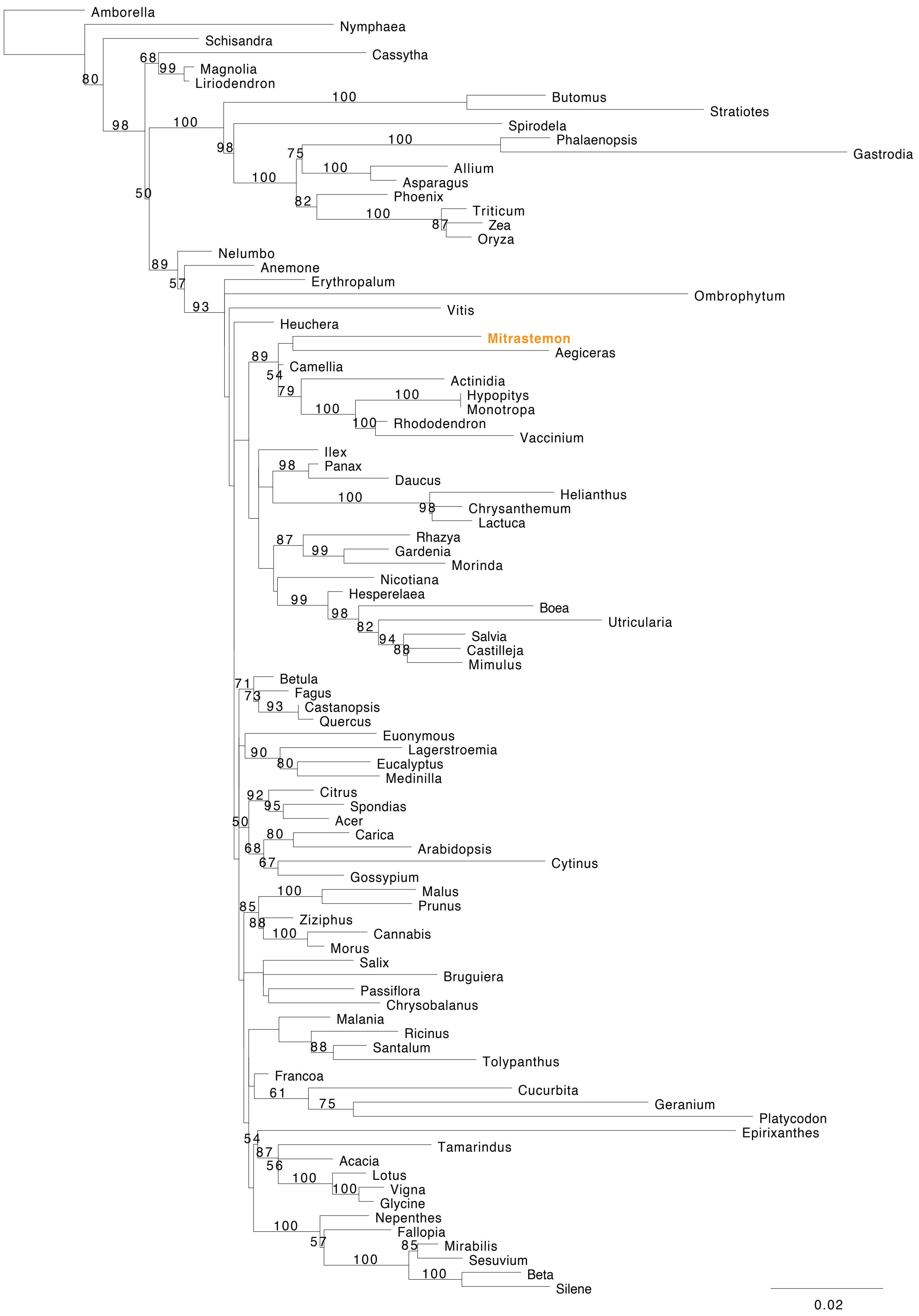
Amborella
Nymphaea
Schisandra
Cassytha
Magnolia
Liriodendron
Butomus
Stratiotes
Spirodela
Phalaenopsis
Gastrodia
Allium
Asparagus
Phoenix
Triticum
Zea
Oryza
Nelumbo
Anemone
Erythropalum
Ombrophytum
Vitis
Heuchera
Mitrastemon
Aegiceras
Camellia
Actinidia
Hypopitys
Monotropa
Rhododendron
Vaccinium
Ilex
Panax
Daucus
Helianthus
Chrysanthemum
Lactuca
Rhazya
Gardenia
Morinda
Nicotiana
Hesperelaea
Boea
Utricularia
Salvia
Castilleja
Mimulus
Betula
Fagus
Castanopsis
Quercus
Euonymous
Lagerstroemia
Eucalyptus
Medinilla
Citrus
Spondias
Acer
Carica
Arabidopsis
Cytinus
Gossypium
Malus
Prunus
Ziziphus
Cannabis
Morus
Salix
Bruguiera
Passiflora
Chrysobalanus
Malania
Ricinus
Santalum
Tolypanthus
Francoa
Cucurbita
Geranium
Platycodon
Epirixanthes
Tamarindus
Acacia
Lotus
Vigna
Glycine
Nepenthes
Fallopia
Mirabilis
Sesuvium
Beta
Silene
0.02

### *ccmFN*

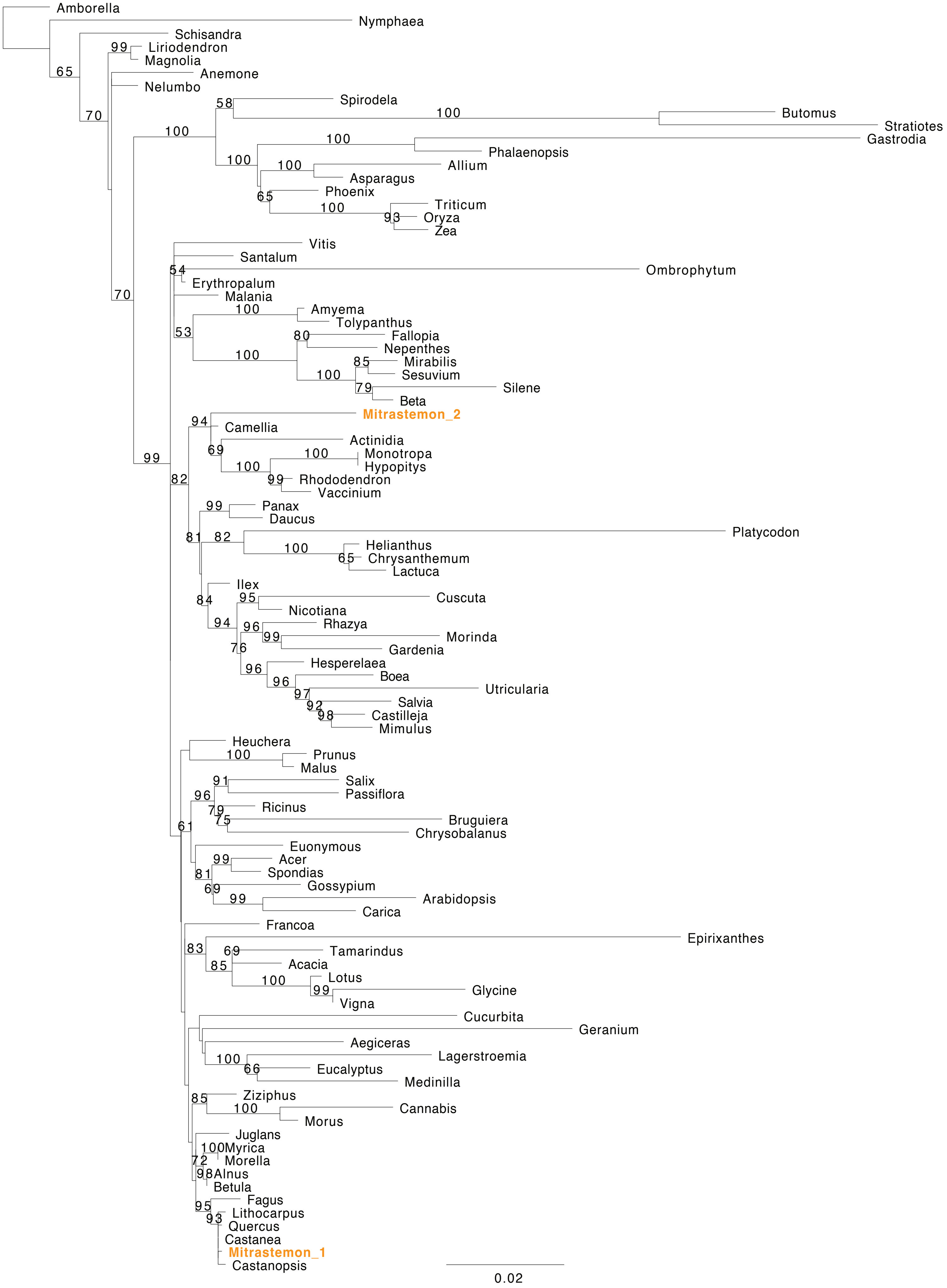
Amborella
Nymphaea
Schisandra
Liriodendron
Magnolia
Anemone
Nelumbo
Spirodela
Butomus
Stratiotes
Gastrodia
Phalaenopsis
Allium
Asparagus
Phoenix
Triticum
Oryza
Zea
Vitis
Santalum
Ombrophytum
Erythropalum
Malania
Amyema
Tolypanthus
Fallopia
Nepenthes
Mirabilis
Sesuvium
Silene
Beta
Mitrastemon_2
Camellia
Actinidia
Monotropa
Hypopitys
Rhododendron
Vaccinium
Panax
Daucus
Platycodon
Helianthus
Chrysanthemum
Lactuca
Ilex
Cuscuta
Nicotiana
Rhazya
Morinda
Gardenia
Hesperelaea
Boea
Utricularia
Salvia
Castilleja
Mimulus
Heuchera
Prunus
Malus
Salix
Passiflora
Ricinus
Bruguiera
Chrysobalanus
Euonymous
Acer
Spondias
Gossypium
Arabidopsis
Carica
Francoa
Epirixanthes
Tamarindus
Acacia
Lotus
Glycine
Vigna
Cucurbita
Geranium
Aegiceras
Lagerstroemia
Eucalyptus
Medinilla
Ziziphus
Cannabis
Morus
Juglans
Myrica
Morella
Alnus
Betula
Fagus
Lithocarpus
Quercus
Castanea
Mitrastemon_1
Castanopsis
0.02

### cob

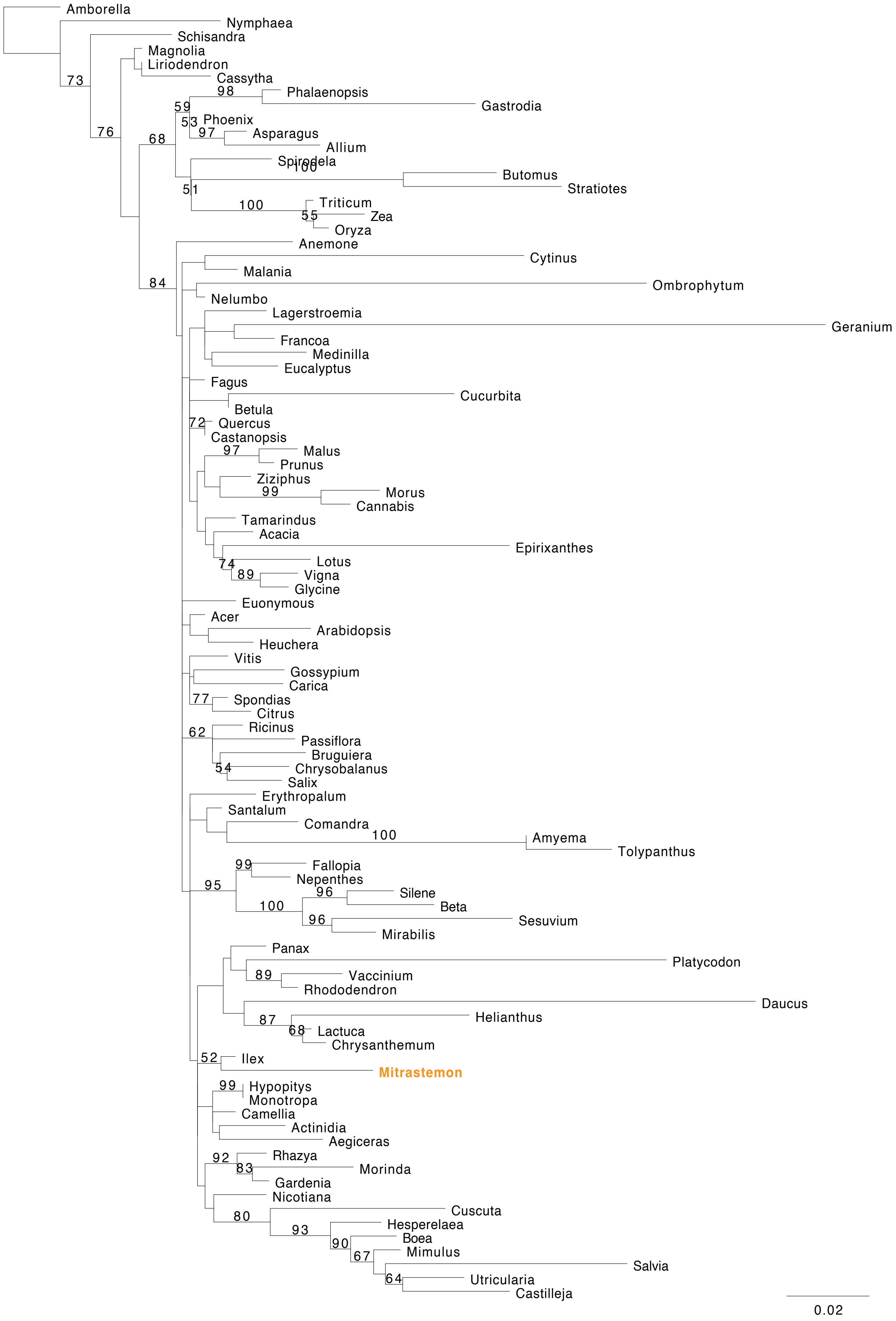
Amborella
Nymphaea
Schisandra
Magnolia
Liriodendron
Cassytha
Phalaenopsis
Gastrodia
Phoenix
Asparagus
Allium
Spirodela
Butomus
Stratiotes
Triticum
Zea
Oryza
Anemone
Cytinus
Malania
Ombrophytum
Nelumbo
Lagerstroemia
Geranium
Francoa
Medinilla
Eucalyptus
Fagus
Cucurbita
Betula
Quercus
Castanopsis
Malus
Prunus
Ziziphus
Morus
Cannabis
Tamarindus
Acacia
Epirixanthes
Lotus
Vigna
Glycine
Euonymous
Acer
Arabidopsis
Heuchera
Vitis
Gossypium
Carica
Spondias
Citrus
Ricinus
Passiflora
Bruguiera
Chrysobalanus
Salix
Erythropalum
Santalum
Comandra
Amyema
Tolypanthus
Fallopia
Nepenthes
Silene
Beta
Sesuvium
Mirabilis
Panax
Platycodon
Vaccinium
Rhododendron
Daucus
Helianthus
Lactuca
Chrysanthemum
Ilex
Mitrastemon
Hypopitys
Monotropa
Camellia
Actinidia
Aegiceras
Rhazya
Morinda
Gardenia
Nicotiana
Cuscuta
Hesperelaea
Boea
Mimulus
Salvia
Utricularia
Castilleja
0.02

### *cox1*

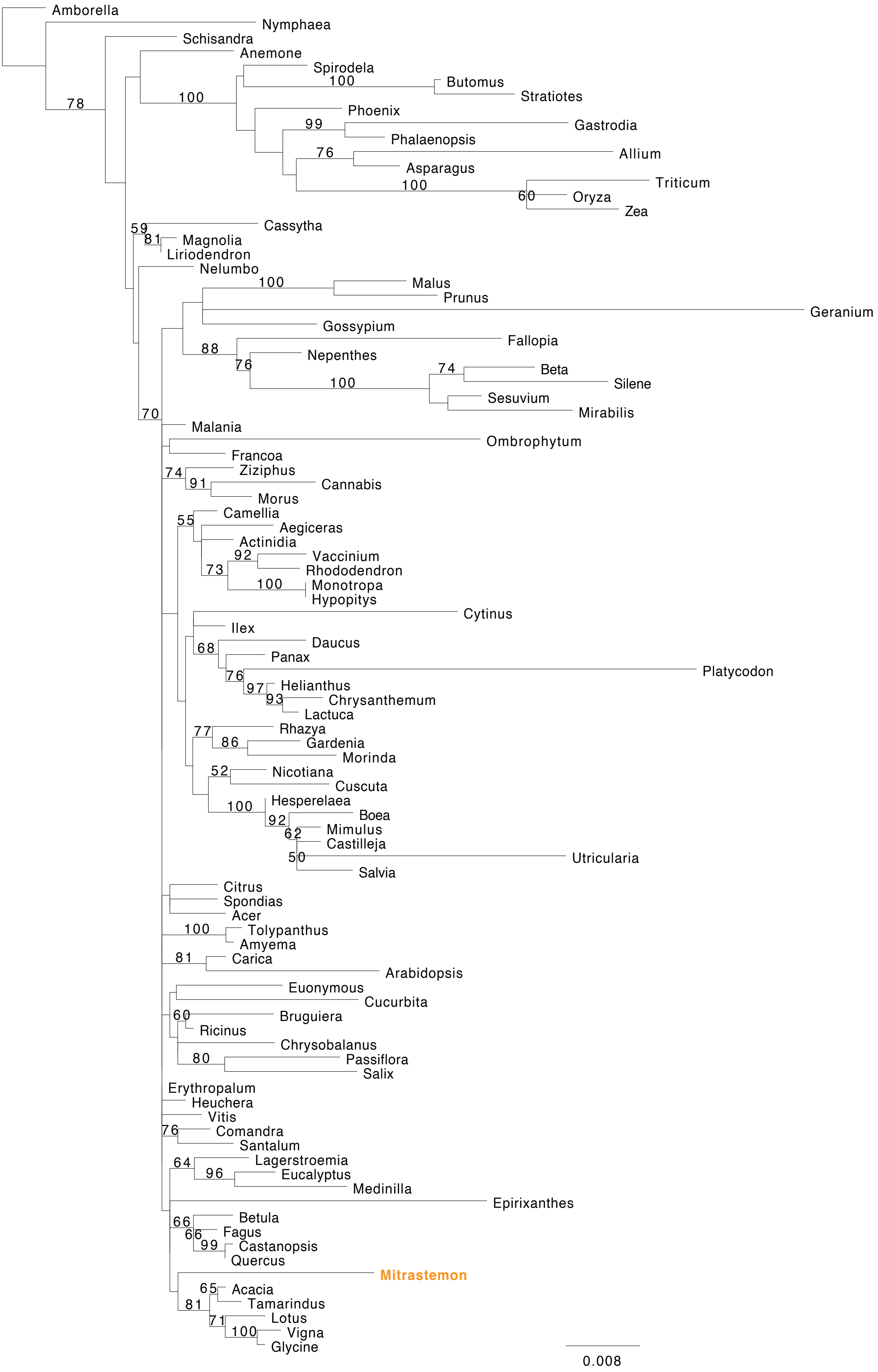

Amborella
Nymphaea
Schisandra
Anemone
Spirodela
100
Butomus
Stratiotes
78
100
Phoenix
99
Gastrodia
Phalaenopsis
76
Allium
Asparagus
100
Triticum
60
Oryza
Zea
59
Cassytha
81
Magnolia
Liriodendron
Nelumbo
100
Malus
Prunus
Geranium
Gossypium
Fallopia
88
Nepenthes
76
74
Beta
Silene
100
Sesuvium
Mirabilis
70
Malania
Ombrophytum
Francoa
74
Ziziphus
91
Cannabis
Morus
55
Camellia
Aegiceras
Actinidia
92
Vaccinium
73
Rhododendron
100
Monotropa
Hypopitys
Cytinus
Ilex
68
Daucus
Panax
76
Platycodon
97
Helianthus
93
Chrysanthemum
Lactuca
77
Rhazya
86
Gardenia
Morinda
52
Nicotiana
Cuscuta
100
Hesperelaea
92
Boea
62
Mimulus
Castilleja
50
Utricularia
Salvia
Citrus
Spondias
Acer
100
Tolypanthus
Amyema
81
Carica
Arabidopsis
Euonymous
Cucurbita
60
Bruguiera
Ricinus
Chrysobalanus
80
Passiflora
Salix
Erythropalum
Heuchera
Vitis
76
Comandra
Santalum
64
Lagerstroemia
96
Eucalyptus
Medinilla
Epirixanthes
66
Betula
66
Fagus
99
Castanopsis
Quercus
Mitrastemon
65
Acacia
81
Tamarindus
71
Lotus
100
Vigna
Glycine
0.008

### *cox2*

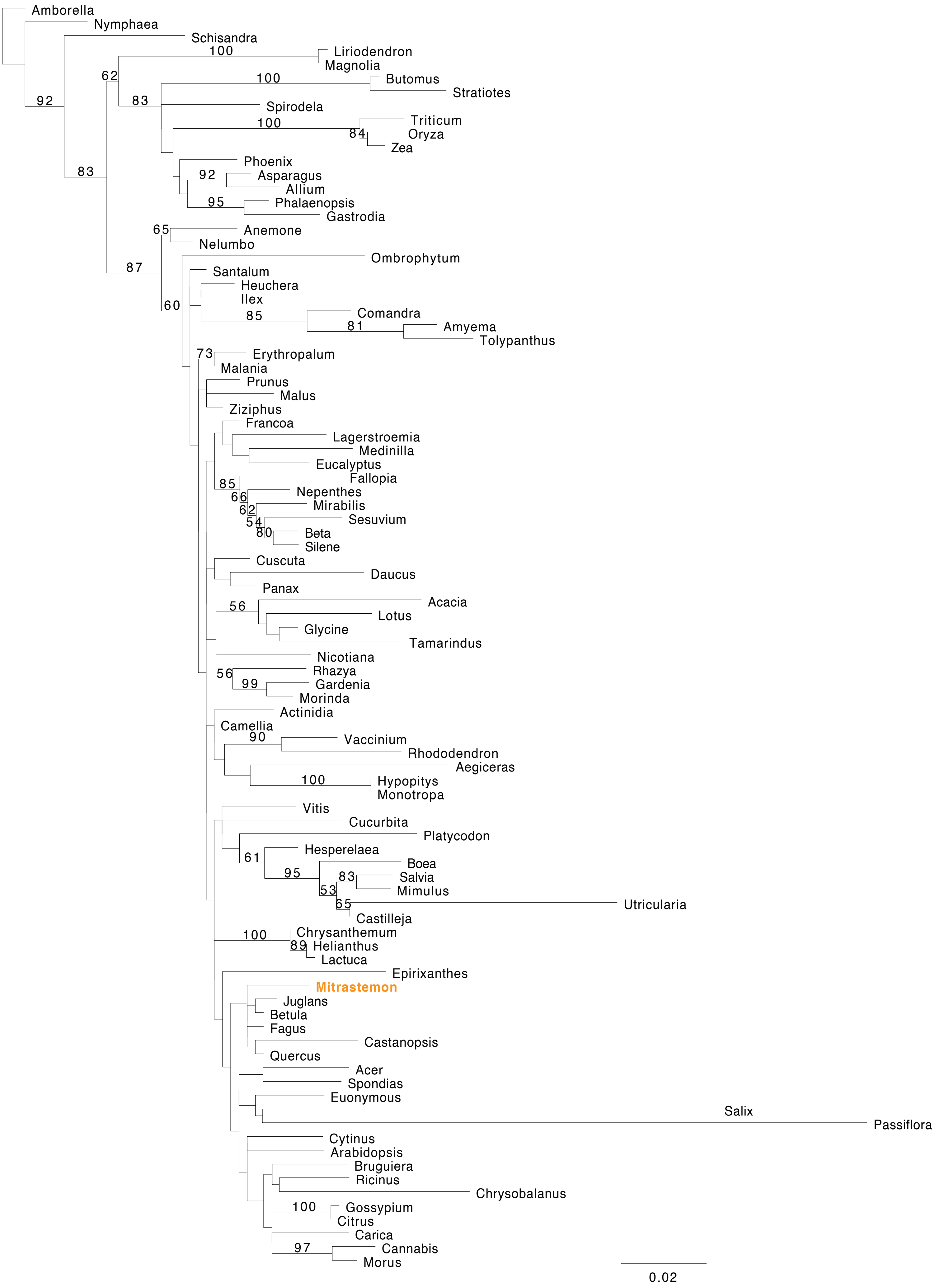

Amborella
Nymphaea
Schisandra
Liriodendron
Magnolia
Butomus
Stratiotes
Spirodela
Triticum
Oryza
Zea
Phoenix
Asparagus
Allium
Phalaenopsis
Gastrodia
Anemone
Nelumbo
Ombrophytum
Santalum
Heuchera
Ilex
Comandra
Amyema
Tolypanthus
Erythropalum
Malania
Prunus
Malus
Ziziphus
Francoa
Lagerstroemia
Medinilla
Eucalyptus
Fallopia
Nepenthes
Mirabilis
Sesuvium
Beta
Silene
Cuscuta
Daucus
Panax
Acacia
Lotus
Glycine
Tamarindus
Nicotiana
Rhazya
Gardenia
Morinda
Actinidia
Camellia
Vaccinium
Rhododendron
Aegiceras
Hypopitys
Monotropa
Vitis
Cucurbita
Platycodon
Hesperelaea
Boea
Salvia
Mimulus
Utricularia
Castilleja
Chrysanthemum
Helianthus
Lactuca
Epirixanthes
Mitrastemon
Juglans
Betula
Fagus
Castanopsis
Quercus
Acer
Spondias
Euonymous
Salix
Passiflora
Cytinus
Arabidopsis
Bruguiera
Ricinus
Chrysobalanus
Gossypium
Citrus
Carica
Cannabis
Morus
0.02

### *cox3*

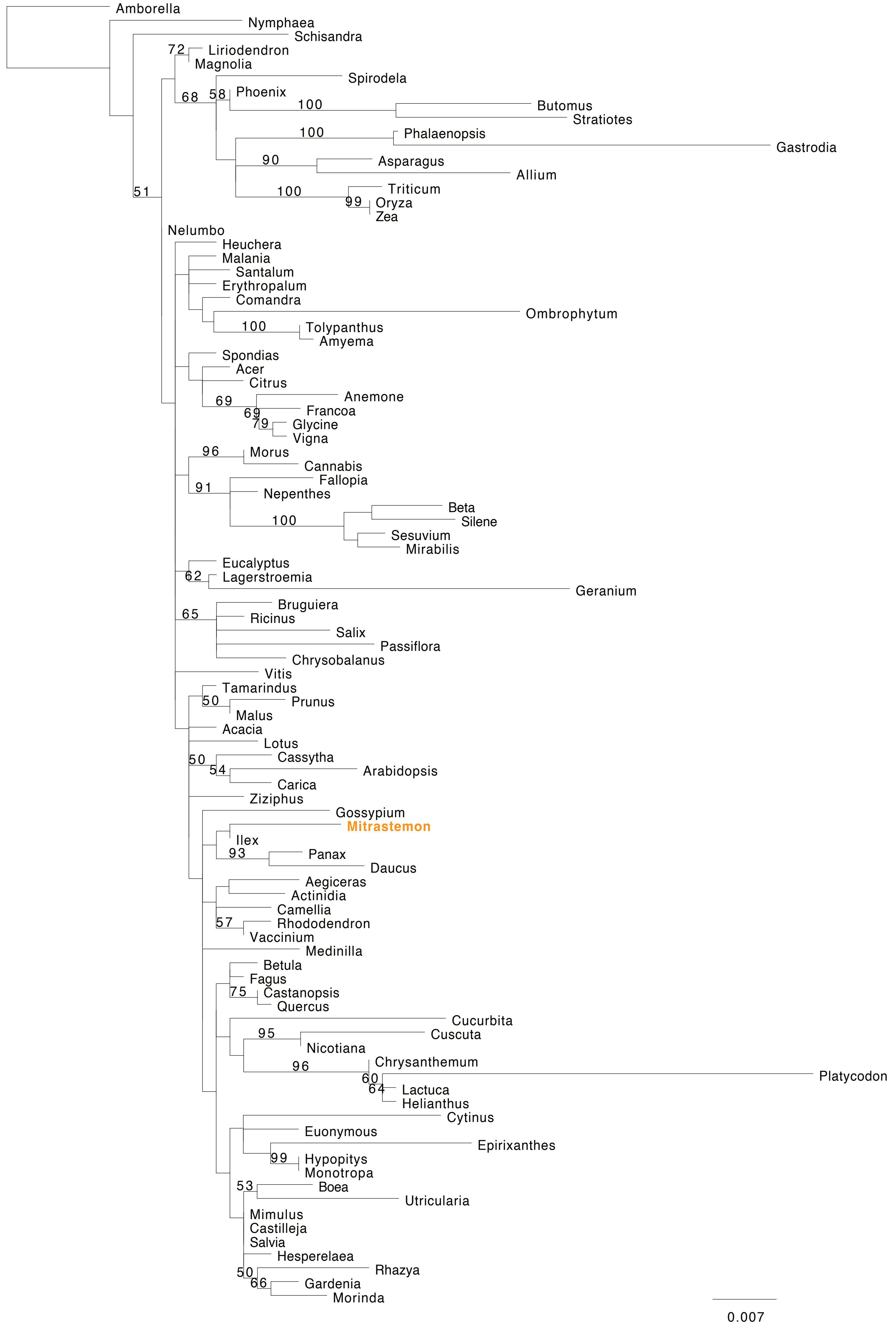
Amborella
Nymphaea
Schisandra
72
Liriodendron
Magnolia
Spirodela
68
58
Phoenix
100
Butomus
Stratiotes
100
Phalaenopsis
Gastrodia
90
Asparagus
Allium
51
100
Triticum
99
Oryza
Zea
Nelumbo
Heuchera
Malania
Santalum
Erythropalum
Comandra
Ombrophytum
100
Tolypanthus
Amyema
Spondias
Acer
Citrus
69
Anemone
69
Francoa
79
Glycine
Vigna
96
Morus
Cannabis
91
Fallopia
Nepenthes
Beta
Silene
100
Sesuvium
Mirabilis
Eucalyptus
62
Lagerstroemia
Geranium
Bruguiera
65
Ricinus
Salix
Passiflora
Chrysobalanus
Vitis
Tamarindus
50
Prunus
Malus
Acacia
Lotus
Cassytha
50
54
Arabidopsis
Carica
Ziziphus
Gossypium
Mitrastemon
Ilex
93
Panax
Daucus
Aegiceras
Actinidia
Camellia
57
Rhododendron
Vaccinium
Medinilla
Betula
Fagus
75
Castanopsis
Quercus
Cucurbita
95
Cuscuta
Nicotiana
96
Chrysanthemum
60
Platycodon
64
Lactuca
Helianthus
Cytinus
Euonymous
Epirixanthes
99
Hypopitys
Monotropa
53
Boea
Utricularia
Mimulus
Castilleja
Salvia
Hesperelaea
50
Rhazya
66
Gardenia
Morinda
0.007

### *matR*

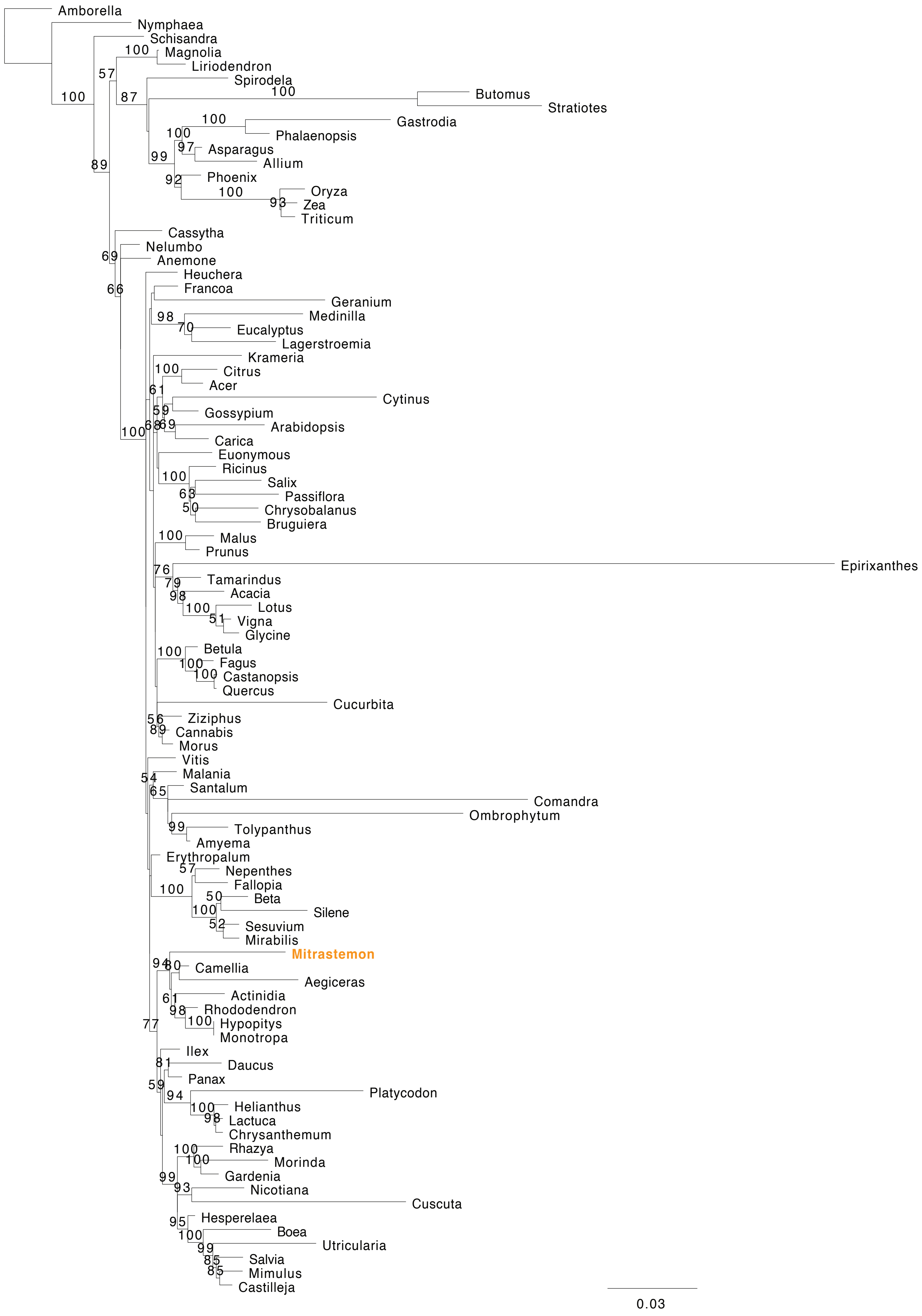

Amborella
Nymphaea
Schisandra
Magnolia
Liriodendron
Spirodela
Butomus
Stratiotes
Gastrodia
Phalaenopsis
Asparagus
Allium
Phoenix
Oryza
Zea
Triticum
Cassytha
Nelumbo
Anemone
Heuchera
Francoa
Geranium
Medinilla
Eucalyptus
Lagerstroemia
Krameria
Citrus
Acer
Cytinus
Gossypium
Arabidopsis
Carica
Euonymous
Ricinus
Salix
Passiflora
Chrysobalanus
Bruguiera
Malus
Prunus
Epirixanthes
Tamarindus
Acacia
Lotus
Vigna
Glycine
Betula
Fagus
Castanopsis
Quercus
Cucurbita
Ziziphus
Cannabis
Morus
Vitis
Malania
Santalum
Comandra
Ombrophytum
Tolypanthus
Amyema
Erythropalum
Nepenthes
Fallopia
Beta
Silene
Sesuvium
Mirabilis
Mitrastemon
Camellia
Aegiceras
Actinidia
Rhododendron
Hypopitys
Monotropa
Ilex
Daucus
Panax
Platycodon
Helianthus
Lactuca
Chrysanthemum
Rhazya
Morinda
Gardenia
Nicotiana
Cuscuta
Hesperelaea
Boea
Utricularia
Salvia
Mimulus
Castilleja
0.03

### *mttB*

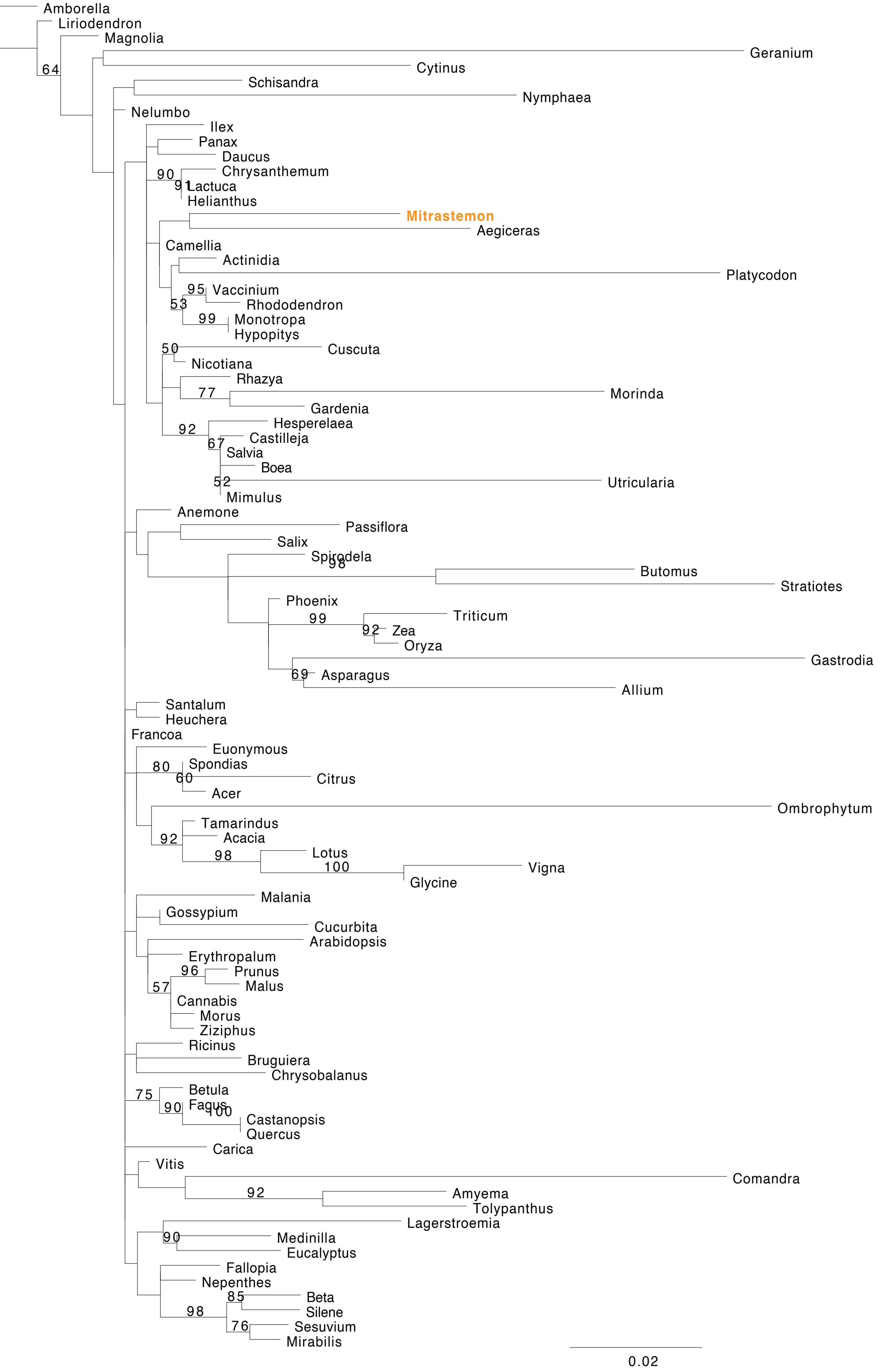

Amborella
Liriodendron
Magnolia
64
Geranium
Cytinus
Schisandra
Nymphaea
Nelumbo
Ilex
Panax
Daucus
90
Chrysanthemum
91
Lactuca
Helianthus
Mitrastemon
Aegiceras
Camellia
Actinidia
Platycodon
95
Vaccinium
53
Rhododendron
99
Monotropa
Hypopitys
50
Cuscuta
Nicotiana
Rhazya
77
Morinda
Gardenia
92
Hesperelaea
67
Castilleja
Salvia
Boea
52
Utricularia
Mimulus
Anemone
Passiflora
Salix
Spirodela
98
Butomus
Stratiotes
Phoenix
99
Triticum
92
Zea
Oryza
Gastrodia
69
Asparagus
Allium
Santalum
Heuchera
Francoa
Euonymous
80
Spondias
60
Citrus
Acer
Ombrophytum
Tamarindus
92
Acacia
98
Lotus
100
Vigna
Glycine
Malania
Gossypium
Cucurbita
Arabidopsis
Erythropalum
96
Prunus
57
Malus
Cannabis
Morus
Ziziphus
Ricinus
Bruguiera
Chrysobalanus
75
Betula
90
Fagus
100
Castanopsis
Quercus
Carica
Vitis
Comandra
92
Amyema
Tolypanthus
Lagerstroemia
90
Medinilla
Eucalyptus
Fallopia
Nepenthes
85
Beta
98
Silene
76
Sesuvium
Mirabilis
0.02

### *nad1*

#### *nad1x1*

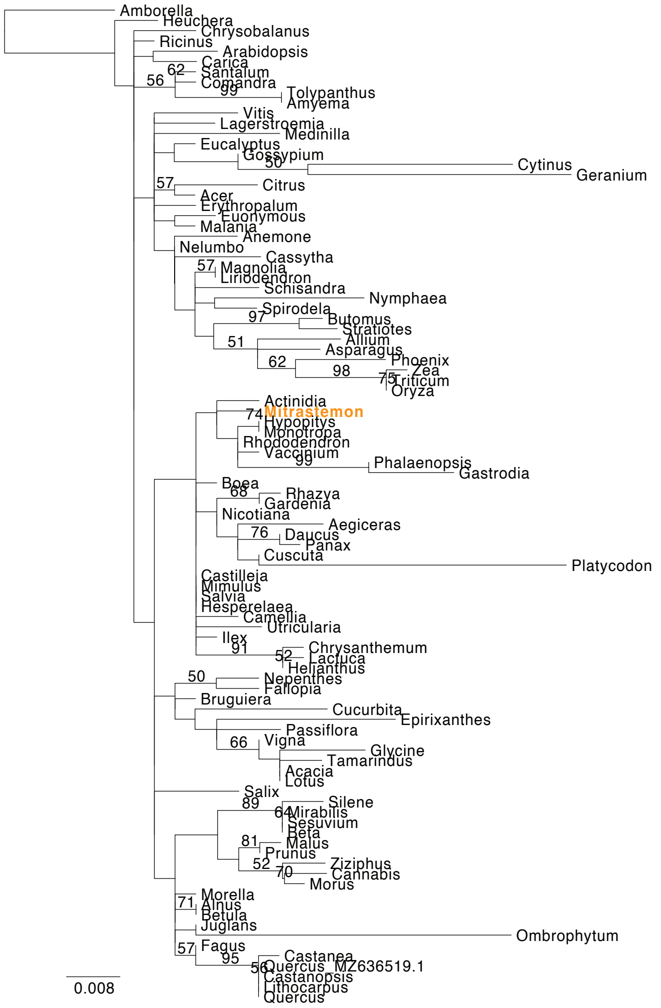


#### *nad1x2x3*

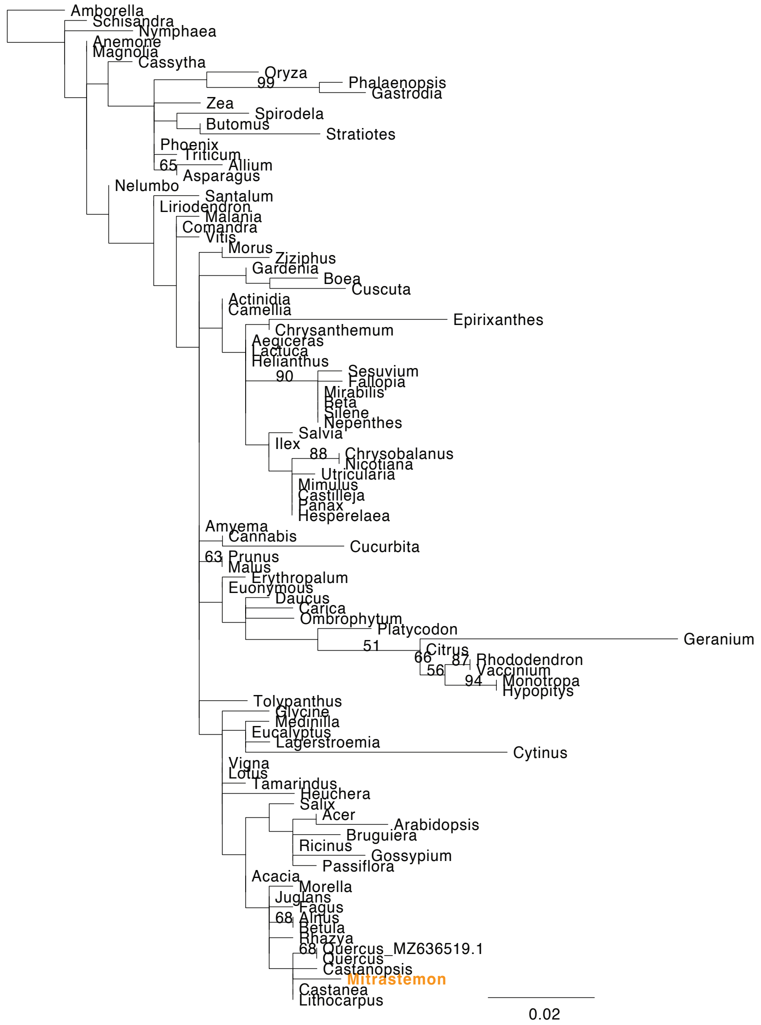


#### *nad1x4x5*

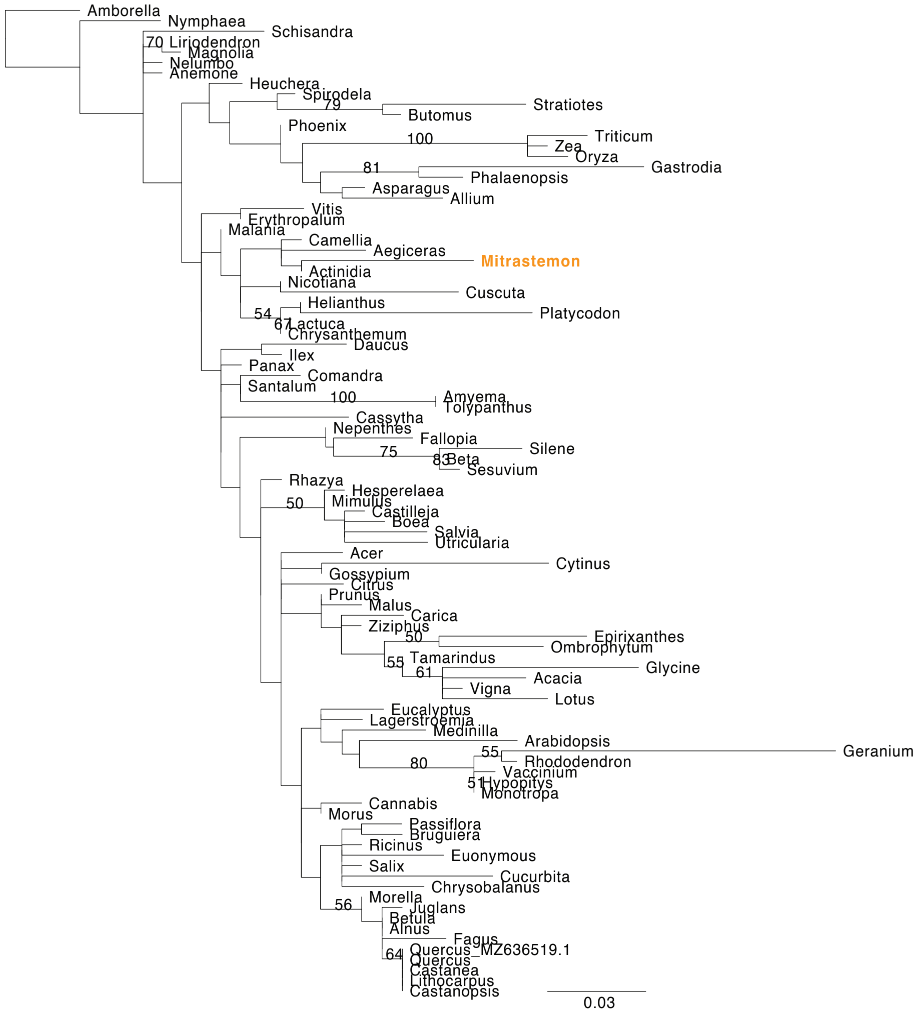

### nad2

##### nad2x1x2

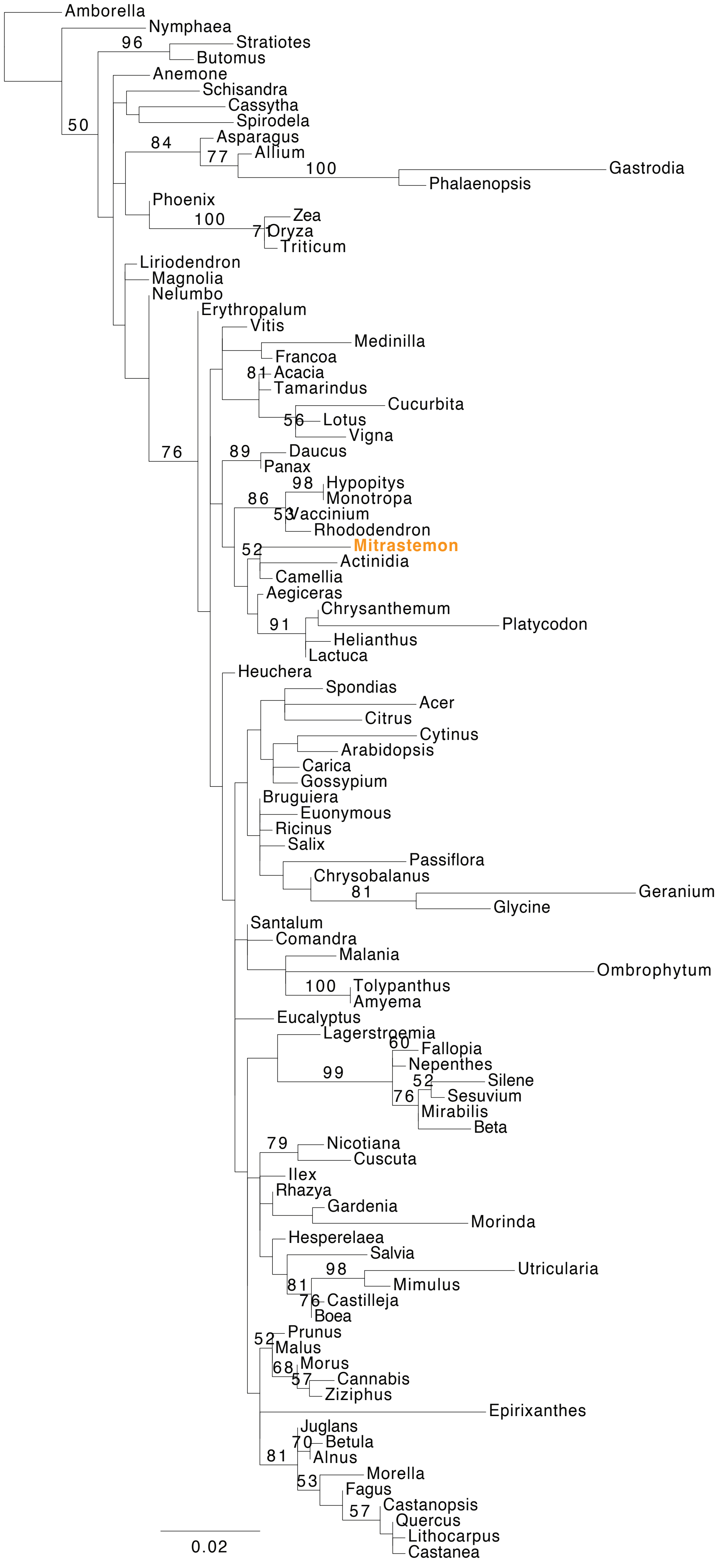


##### nad2x3x4x5

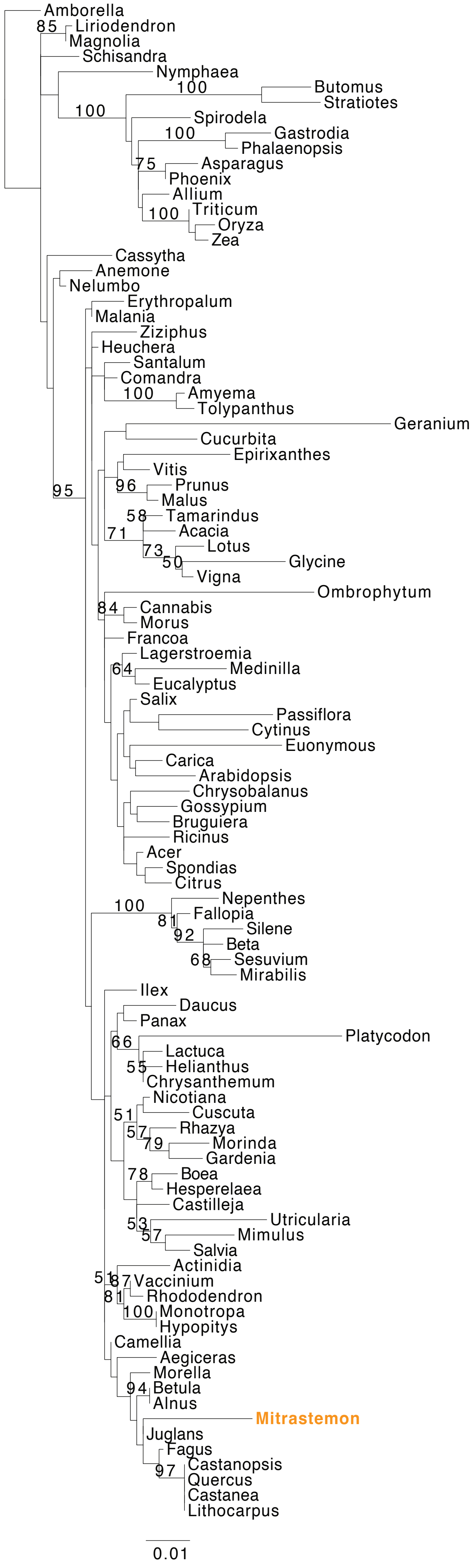

### *nad3*

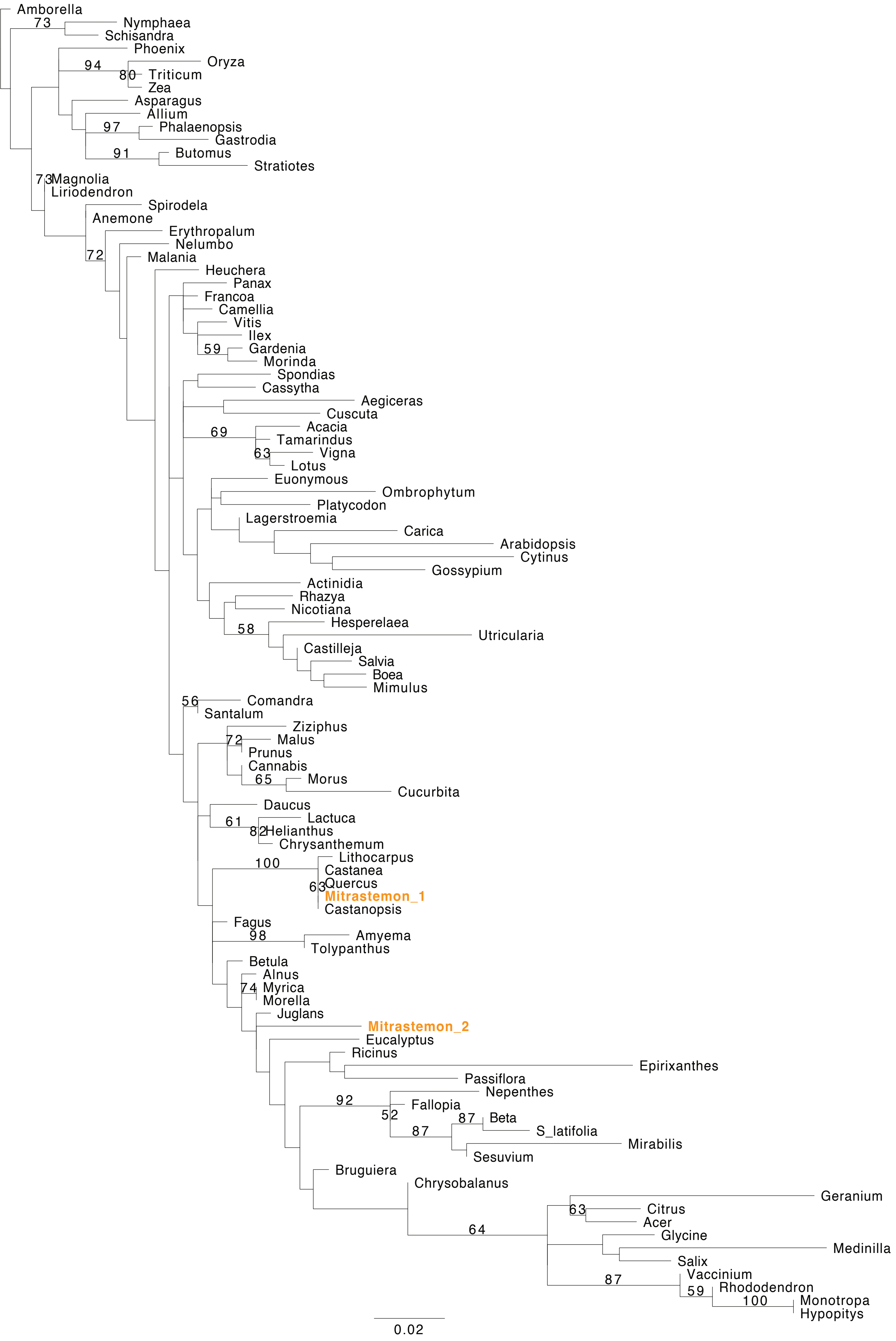
Amborella
73
Nymphaea
Schisandra
Phoenix
94
Oryza
80
Triticum
Zea
Asparagus
Allium
97
Phalaenopsis
Gastrodia
91
Butomus
Stratiotes
73
Magnolia
Liriodendron
Spirodela
Anemone
Erythropalum
Nelumbo
72
Malania
Heuchera
Panax
Francoa
Camellia
Vitis
Ilex
59
Gardenia
Morinda
Spondias
Cassytha
Aegiceras
Cuscuta
69
Acacia
Tamarindus
63
Vigna
Lotus
Euonymous
Ombrophytum
Platycodon
Lagerstroemia
Carica
Arabidopsis
Cytinus
Gossypium
Actinidia
Rhazya
Nicotiana
58
Hesperelaea
Utricularia
Castilleja
Salvia
Boea
Mimulus
56
Comandra
Santalum
Ziziphus
72
Malus
Prunus
Cannabis
65
Morus
Cucurbita
Daucus
61
Lactuca
82
Helianthus
Chrysanthemum
100
Lithocarpus
Castanea
63
Quercus
Mitrastemon_1
Castanopsis
Fagus
98
Amyema
Tolypanthus
Betula
Alnus
74
Myrica
Morella
Juglans
Mitrastemon_2
Eucalyptus
Ricinus
Epirixanthes
Passiflora
92
Nepenthes
Fallopia
52
87
Beta
87
S_latifolia
Mirabilis
Sesuvium
Bruguiera
Chrysobalanus
Geranium
63
Citrus
Acer
64
Glycine
Medinilla
Salix
87
Vaccinium
59
Rhododendron
100
Monotropa
Hypopitys
0.02

### *nad4*

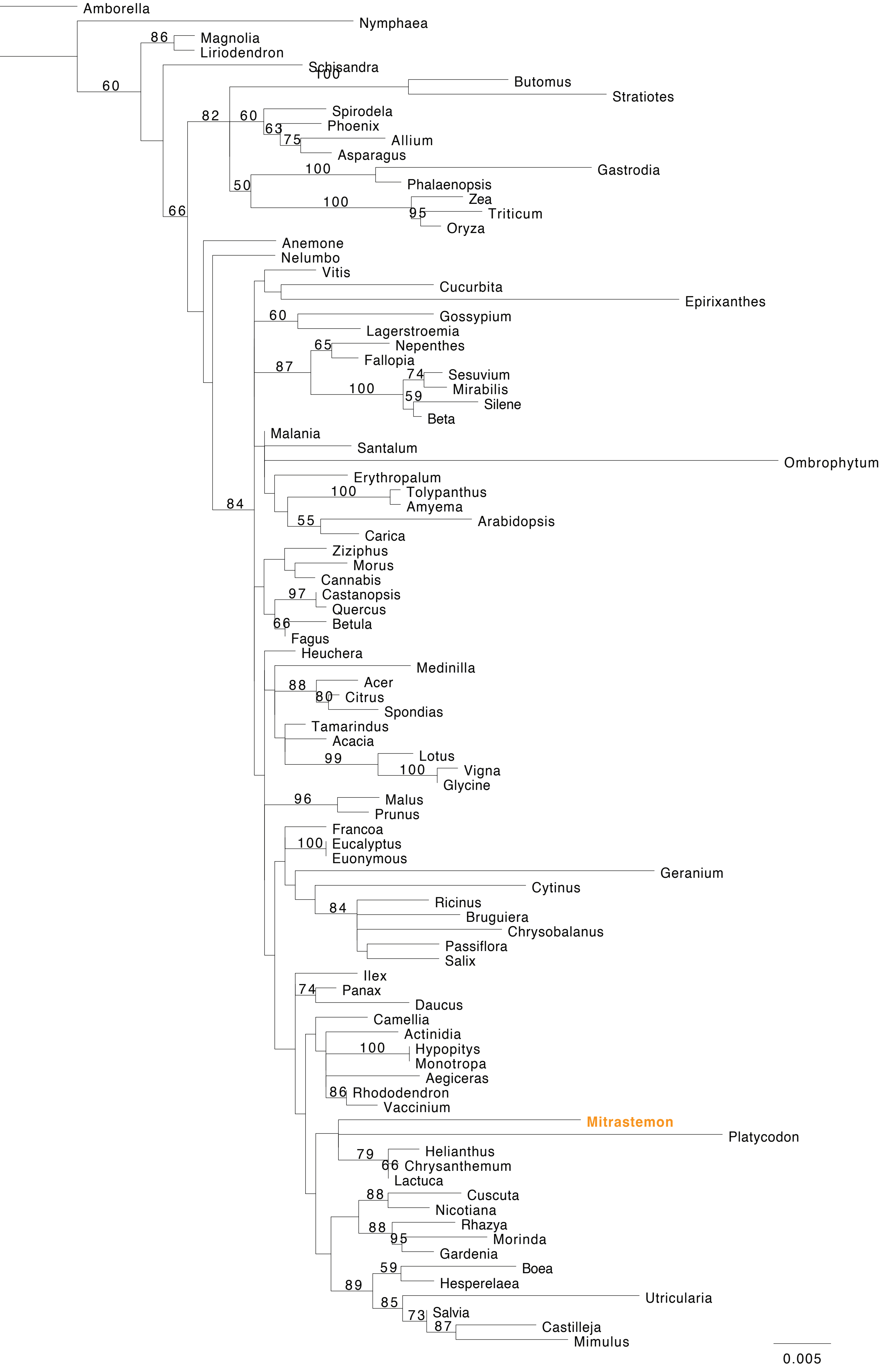
Amborella
Nymphaea
Magnolia
Liriodendron
Schisandra
Butomus
Stratiotes
Spirodela
Phoenix
Allium
Asparagus
Gastrodia
Phalaenopsis
Zea
Triticum
Oryza
Anemone
Nelumbo
Vitis
Cucurbita
Epirixanthes
Gossypium
Lagerstroemia
Nepenthes
Fallopia
Sesuvium
Mirabilis
Silene
Beta
Malania
Santalum
Ombrophytum
Erythropalum
Tolypanthus
Amyema
Arabidopsis
Carica
Ziziphus
Morus
Cannabis
Castanopsis
Quercus
Betula
Fagus
Heuchera
Medinilla
Acer
Citrus
Spondias
Tamarindus
Acacia
Lotus
Vigna
Glycine
Malus
Prunus
Francoa
Eucalyptus
Euonymous
Geranium
Cytinus
Ricinus
Bruguiera
Chrysobalanus
Passiflora
Salix
Ilex
Panax
Daucus
Camellia
Actinidia
Hypopitys
Monotropa
Aegiceras
Rhododendron
Vaccinium
Mitrastemon
Platycodon
Helianthus
Chrysanthemum
Lactuca
Cuscuta
Nicotiana
Rhazya
Morinda
Gardenia
Boea
Hesperelaea
Utricularia
Salvia
Castilleja
Mimulus
0.005

### *nad4 intron - Chimeric - Native region*

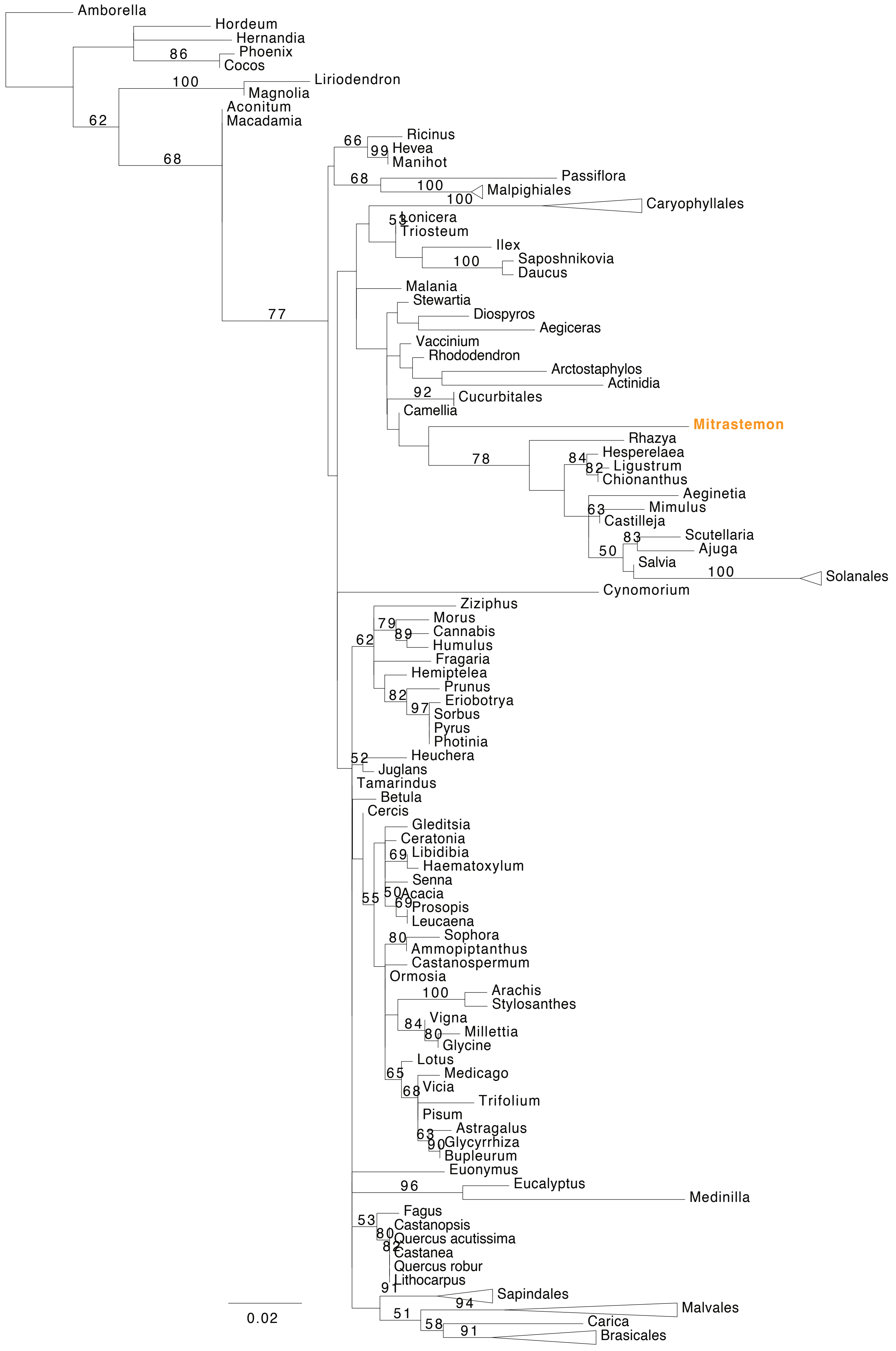

### *nad4 intron- Chimeric - Foreign region*

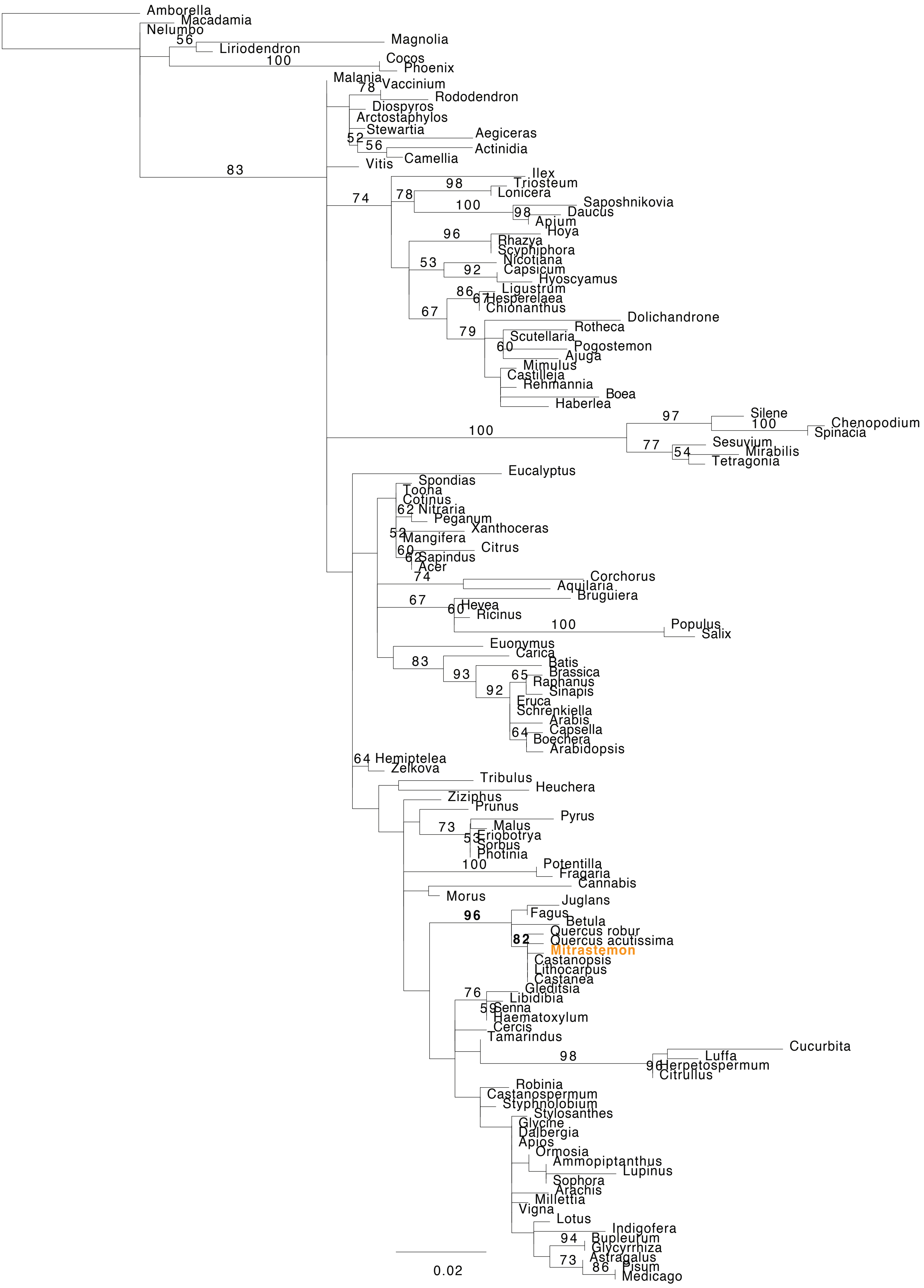

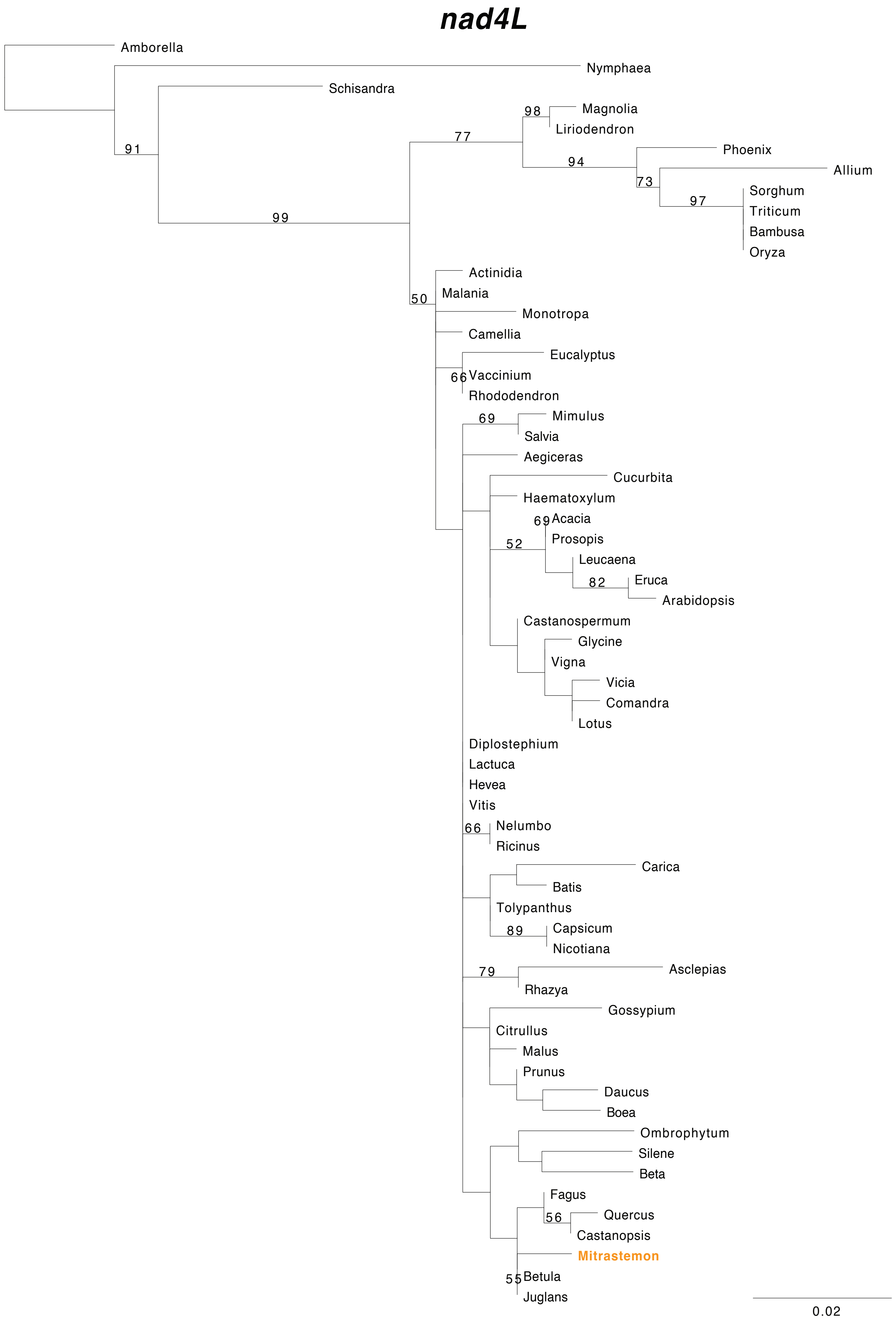
nad4L
Amborella
Nymphaea
Schisandra
91
99
98
Magnolia
Liriodendron
77
94
Phoenix
Allium
73
97
Sorghum
Triticum
Bambusa
Oryza
50
Actinidia
Malania
Monotropa
Camellia
Eucalyptus
66
Vaccinium
Rhododendron
69
Mimulus
Salvia
Aegiceras
Cucurbita
Haematoxylum
69
Acacia
Prosopis
52
Leucaena
82
Eruca
Arabidopsis
Castanospermum
Glycine
Vigna
Vicia
Comandra
Lotus
Diplostephium
Lactuca
Hevea
Vitis
66
Nelumbo
Ricinus
Carica
Batis
Tolypanthus
89
Capsicum
Nicotiana
79
Asclepias
Rhazya
Gossypium
Citrullus
Malus
Prunus
Daucus
Boea
Ombrophytum
Silene
Beta
Fagus
56
Quercus
Castanopsis
Mitrastemon
55
Betula
Juglans
0.02

### nad5

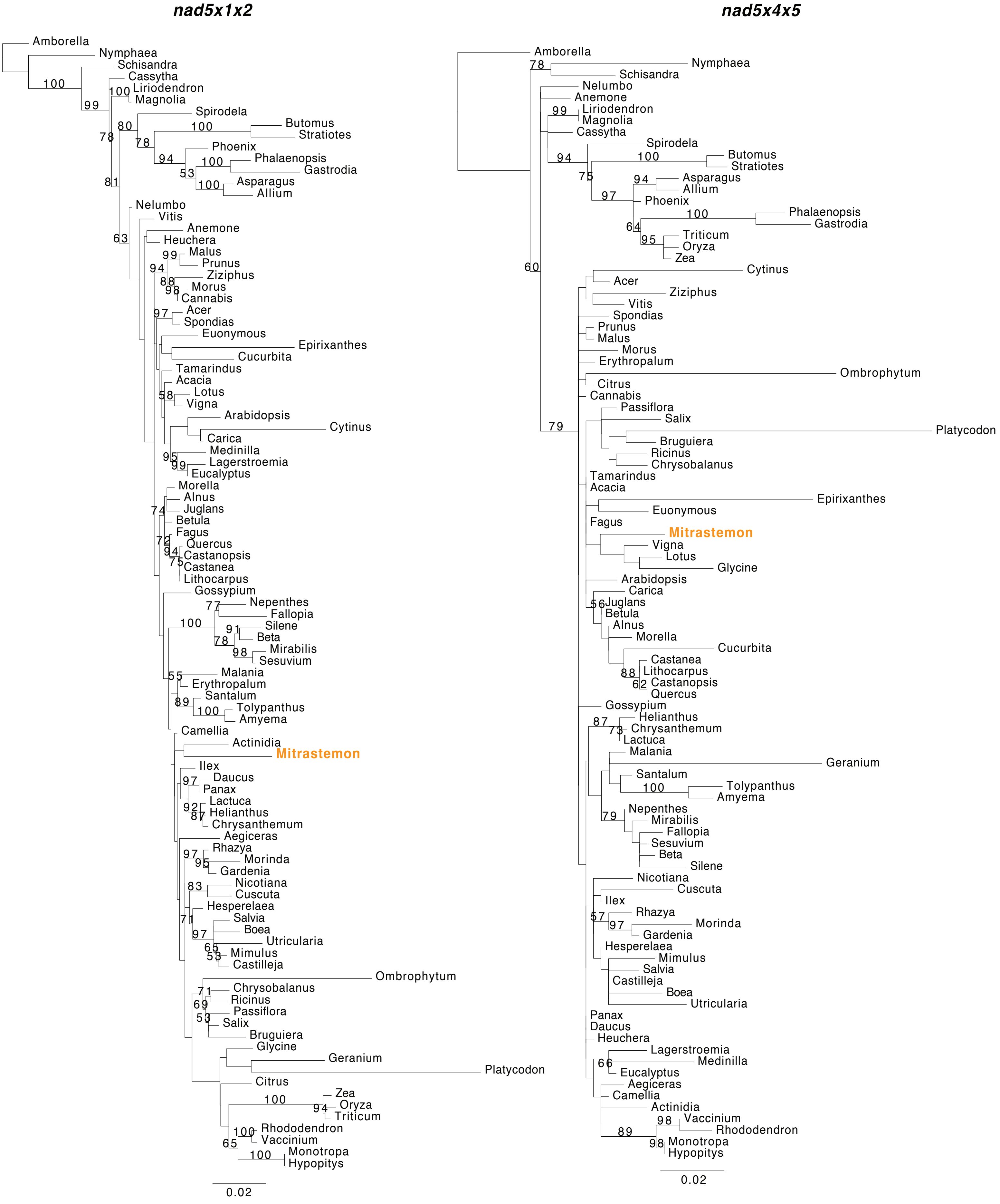
nad5x1x2
Amborella
Nymphaea
Schisandra
Cassytha
Liriodendron
Magnolia
Spirodela
Butomus
Stratiotes
Phoenix
Phalaenopsis
Gastrodia
Asparagus
Allium
Nelumbo
Vitis
Anemone
Heuchera
Malus
Prunus
Ziziphus
Morus
Cannabis
Acer
Spondias
Euonymous
Epirixanthes
Cucurbita
Tamarindus
Acacia
Lotus
Vigna
Arabidopsis
Cytinus
Carica
Medinilla
Lagerstroemia
Eucalyptus
Morella
Alnus
Juglans
Betula
Fagus
Quercus
Castanopsis
Castanea
Lithocarpus
Gossypium
Nepenthes
Fallopia
Silene
Beta
Mirabilis
Sesuvium
Malania
Erythropalum
Santalum
Tolypanthus
Amyema
Camellia
Actinidia
Mitrastemon
Ilex
Daucus
Panax
Lactuca
Helianthus
Chrysanthemum
Aegiceras
Rhazya
Morinda
Gardenia
Nicotiana
Cuscuta
Hesperelaea
Salvia
Boea
Utricularia
Mimulus
Castilleja
Ombrophytum
Chrysobalanus
Ricinus
Passiflora
Salix
Bruguiera
Glycine
Geranium
Platycodon
Citrus
Zea
Oryza
Triticum
Rhododendron
Vaccinium
Monotropa
Hypopitys
0.02
nad5x4x5
Amborella
Nymphaea
Schisandra
Nelumbo
Anemone
Liriodendron
Magnolia
Cassytha
Spirodela
Butomus
Stratiotes
Asparagus
Allium
Phoenix
Phalaenopsis
Gastrodia
Triticum
Oryza
Zea
Cytinus
Acer
Ziziphus
Vitis
Spondias
Prunus
Malus
Morus
Erythropalum
Ombrophytum
Citrus
Cannabis
Passiflora
Salix
Platycodon
Bruguiera
Ricinus
Chrysobalanus
Tamarindus
Acacia
Epirixanthes
Euonymous
Fagus
Mitrastemon
Vigna
Lotus
Glycine
Arabidopsis
Carica
Juglans
Betula
Alnus
Morella
Cucurbita
Castanea
Lithocarpus
Castanopsis
Quercus
Gossypium
Helianthus
Chrysanthemum
Lactuca
Malania
Geranium
Santalum
Tolypanthus
Amyema
Nepenthes
Mirabilis
Fallopia
Sesuvium
Beta
Silene
Nicotiana
Cuscuta
Ilex
Rhazya
Morinda
Gardenia
Hesperelaea
Mimulus
Salvia
Castilleja
Boea
Utricularia
Panax
Daucus
Heuchera
Lagerstroemia
Medinilla
Eucalyptus
Aegiceras
Camellia
Actinidia
Vaccinium
Rhododendron
Monotropa
Hypopitys
0.02

### *nad6*

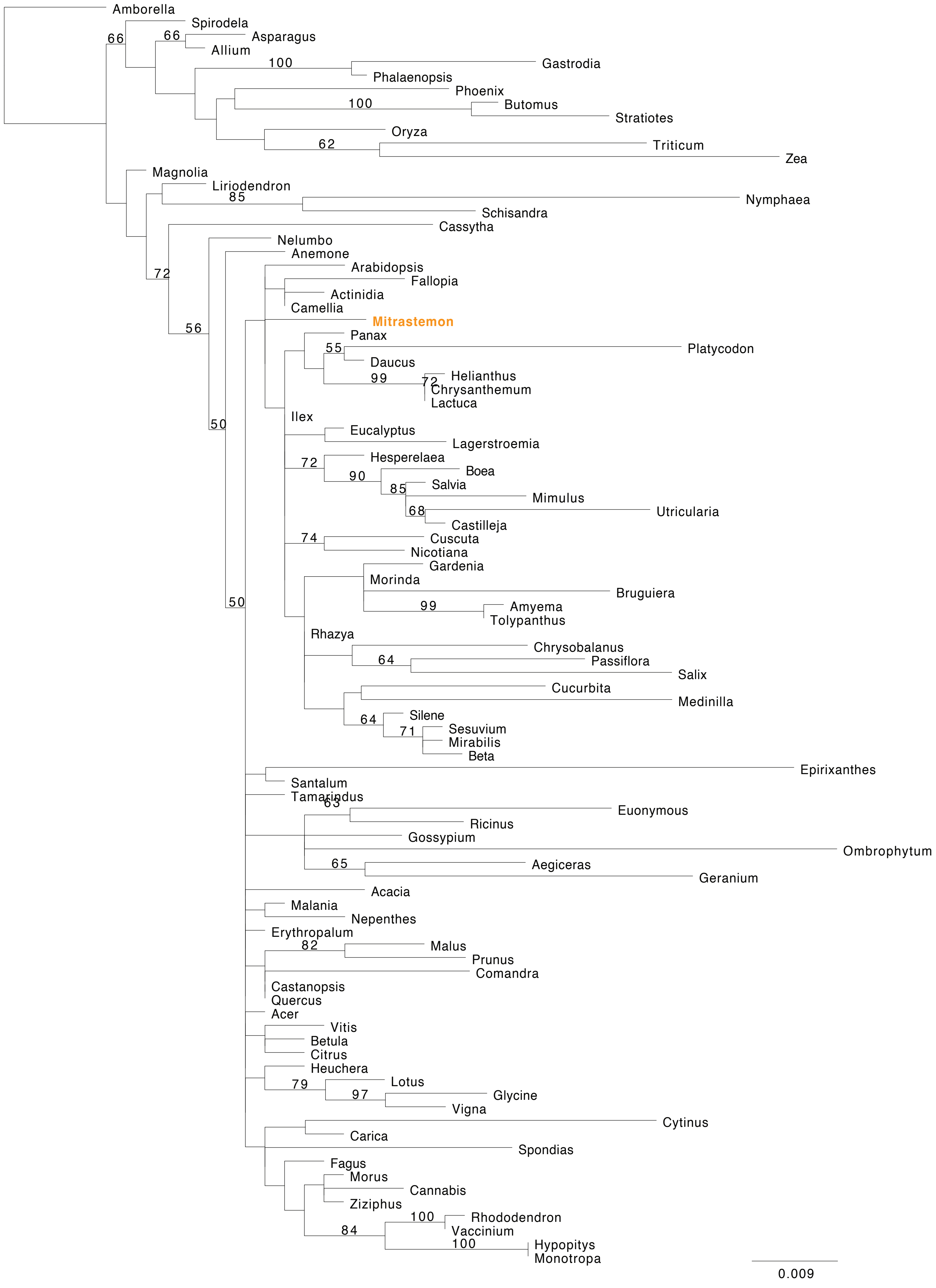

Amborella
Spirodela
66
66
Asparagus
Allium
100
Gastrodia
Phalaenopsis
Phoenix
100
Butomus
Stratiotes
Oryza
62
Triticum
Zea
Magnolia
Liriodendron
85
Nymphaea
Schisandra
Cassytha
72
Nelumbo
Anemone
Arabidopsis
Fallopia
Actinidia
Camellia
56
Mitrastemon
Panax
55
Platycodon
Daucus
99
72
Helianthus
Chrysanthemum
Lactuca
50
Ilex
Eucalyptus
Lagerstroemia
72
Hesperelaea
90
Boea
85
Salvia
Mimulus
68
Utricularia
Castilleja
74
Cuscuta
Nicotiana
Gardenia
Morinda
Bruguiera
99
Amyema
Tolypanthus
50
Rhazya
Chrysobalanus
64
Passiflora
Salix
Cucurbita
Medinilla
64
Silene
71
Sesuvium
Mirabilis
Beta
Epirixanthes
Santalum
Tamarindus
63
Euonymous
Ricinus
Gossypium
Ombrophytum
65
Aegiceras
Geranium
Acacia
Malania
Nepenthes
Erythropalum
82
Malus
Prunus
Comandra
Castanopsis
Quercus
Acer
Vitis
Betula
Citrus
Heuchera
79
Lotus
97
Glycine
Vigna
Cytinus
Carica
Spondias
Fagus
Morus
Cannabis
Ziziphus
100
Rhododendron
Vaccinium
84
100
Hypopitys
Monotropa
0.009

### nad7

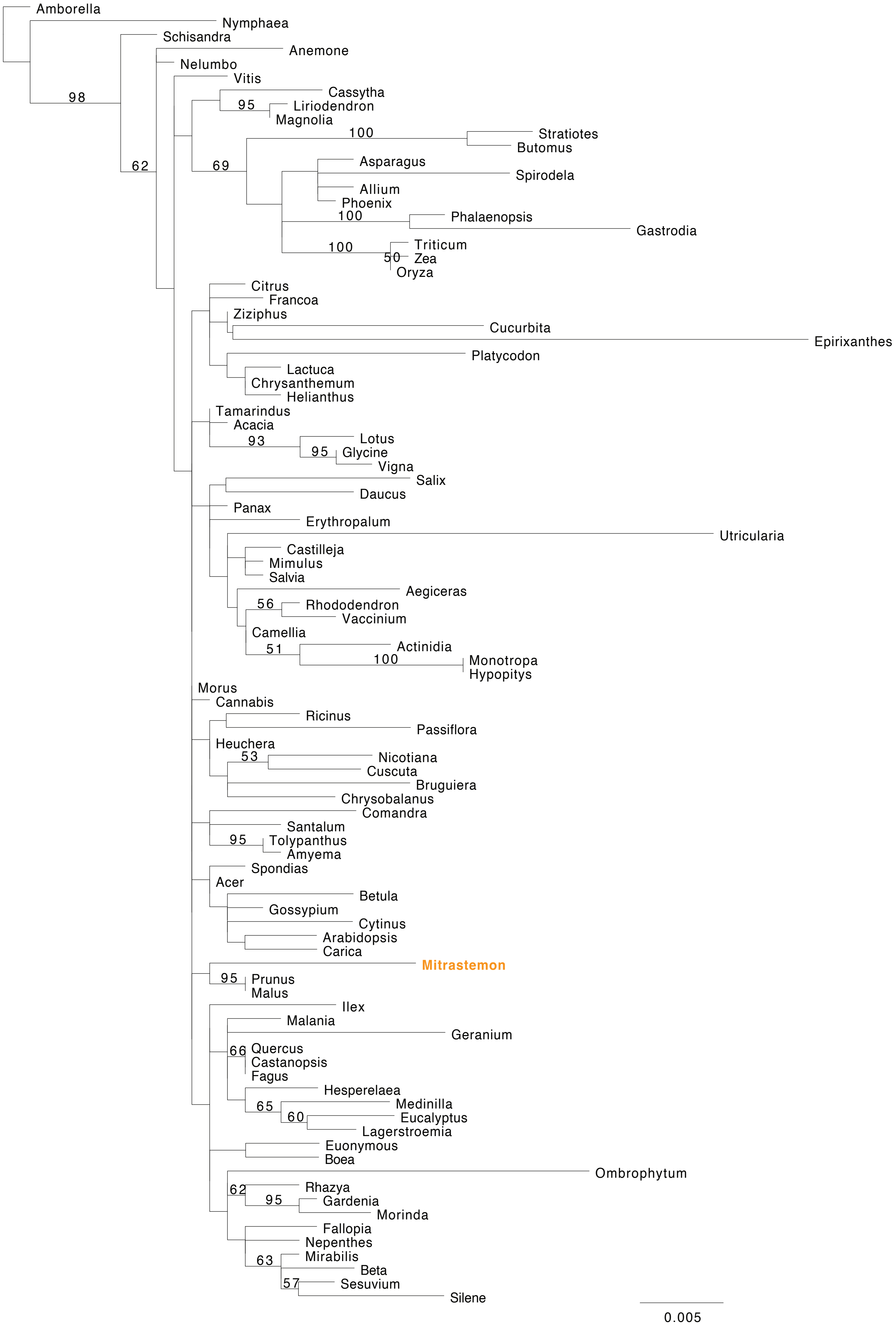
Amborella
Nymphaea
Schisandra
Anemone
Nelumbo
Vitis
Cassytha
Liriodendron
Magnolia
Stratiotes
Butomus
Asparagus
Spirodela
Allium
Phoenix
Phalaenopsis
Gastrodia
Triticum
Zea
Oryza
Citrus
Francoa
Ziziphus
Cucurbita
Epirixanthes
Platycodon
Lactuca
Chrysanthemum
Helianthus
Tamarindus
Acacia
Lotus
Glycine
Vigna
Salix
Daucus
Panax
Erythropalum
Utricularia
Castilleja
Mimulus
Salvia
Aegiceras
Rhododendron
Vaccinium
Camellia
Actinidia
Monotropa
Hypopitys
Morus
Cannabis
Ricinus
Passiflora
Heuchera
Nicotiana
Cuscuta
Bruguiera
Chrysobalanus
Comandra
Santalum
Tolypanthus
Amyema
Spondias
Acer
Betula
Gossypium
Cytinus
Arabidopsis
Carica
Mitrastemon
Prunus
Malus
Ilex
Malania
Geranium
Quercus
Castanopsis
Fagus
Hesperelaea
Medinilla
Eucalyptus
Lagerstroemia
Euonymous
Boea
Ombrophytum
Rhazya
Gardenia
Morinda
Fallopia
Nepenthes
Mirabilis
Beta
Sesuvium
Silene
98
62
95
100
69
100
100
50
93
95
56
51
100
53
95
95
66
65
60
62
95
63
57
0.005

### nad9

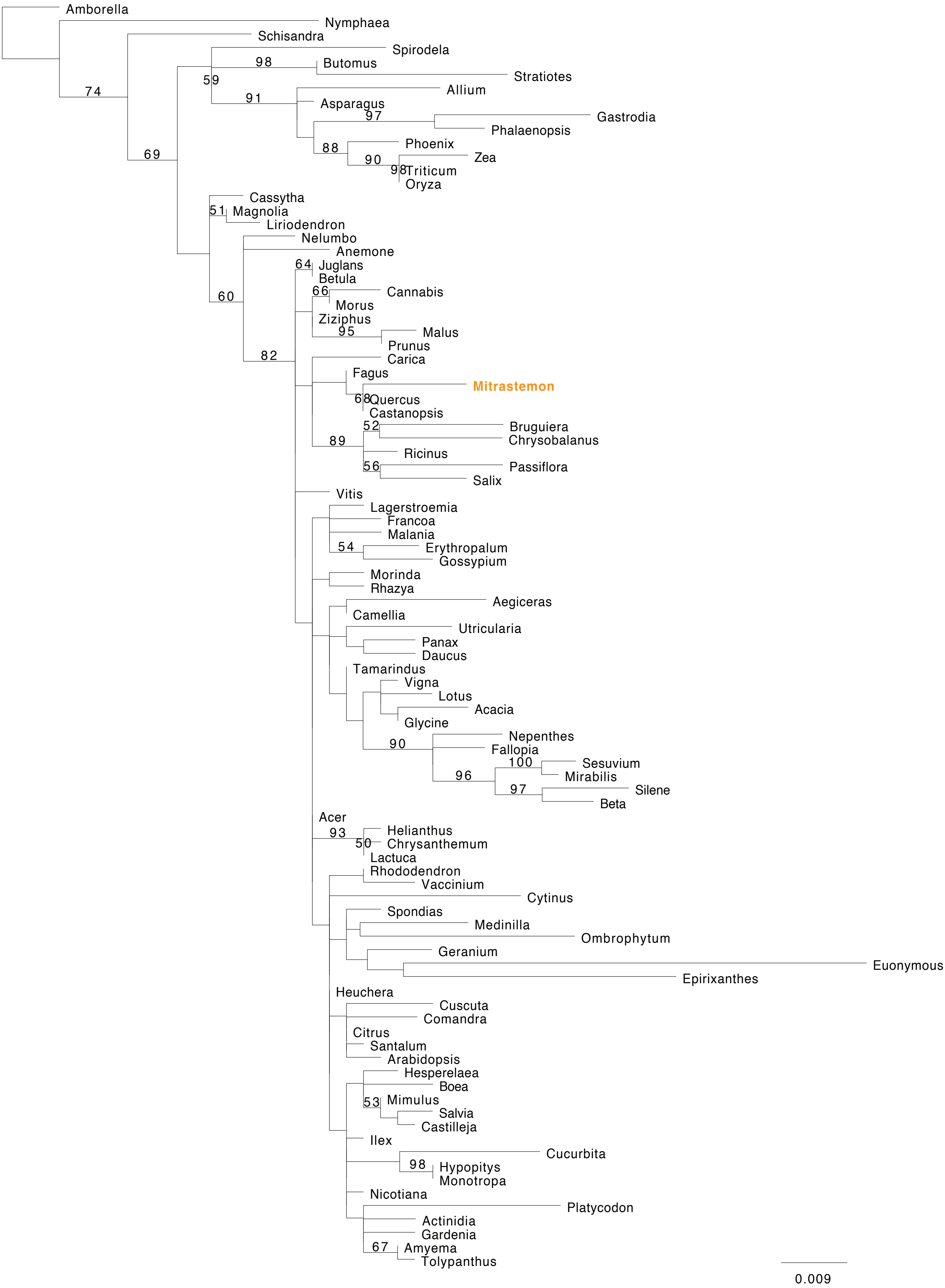

Amborella
Nymphaea
Schisandra
Spirodela
Butomus
Stratiotes
Allium
Asparagus
Gastrodia
Phalaenopsis
Phoenix
Zea
Triticum
Oryza
Cassytha
Magnolia
Liriodendron
Nelumbo
Anemone
Juglans
Betula
Cannabis
Morus
Ziziphus
Malus
Prunus
Carica
Fagus
Mitrastemon
Quercus
Castanopsis
Bruguiera
Chrysobalanus
Ricinus
Passiflora
Salix
Vitis
Lagerstroemia
Francoa
Malania
Erythropalum
Gossypium
Morinda
Rhazya
Aegiceras
Camellia
Utricularia
Panax
Daucus
Tamarindus
Vigna
Lotus
Acacia
Glycine
Nepenthes
Fallopia
Sesuvium
Mirabilis
Silene
Beta
Acer
Helianthus
Chrysanthemum
Lactuca
Rhododendron
Vaccinium
Cytinus
Spondias
Medinilla
Ombrophytum
Geranium
Euonymous
Epirixanthes
Heuchera
Cuscuta
Comandra
Citrus
Santalum
Arabidopsis
Hesperelaea
Boea
Mimulus
Salvia
Castilleja
Ilex
Cucurbita
Hypopitys
Monotropa
Nicotiana
Platycodon
Actinidia
Gardenia
Amyema
Tolypanthus
74
69
59
98
91
97
88
90
98
51
60
82
64
66
95
68
52
89
56
54
90
100
96
97
93
50
53
98
67
0.009

***rpl2***

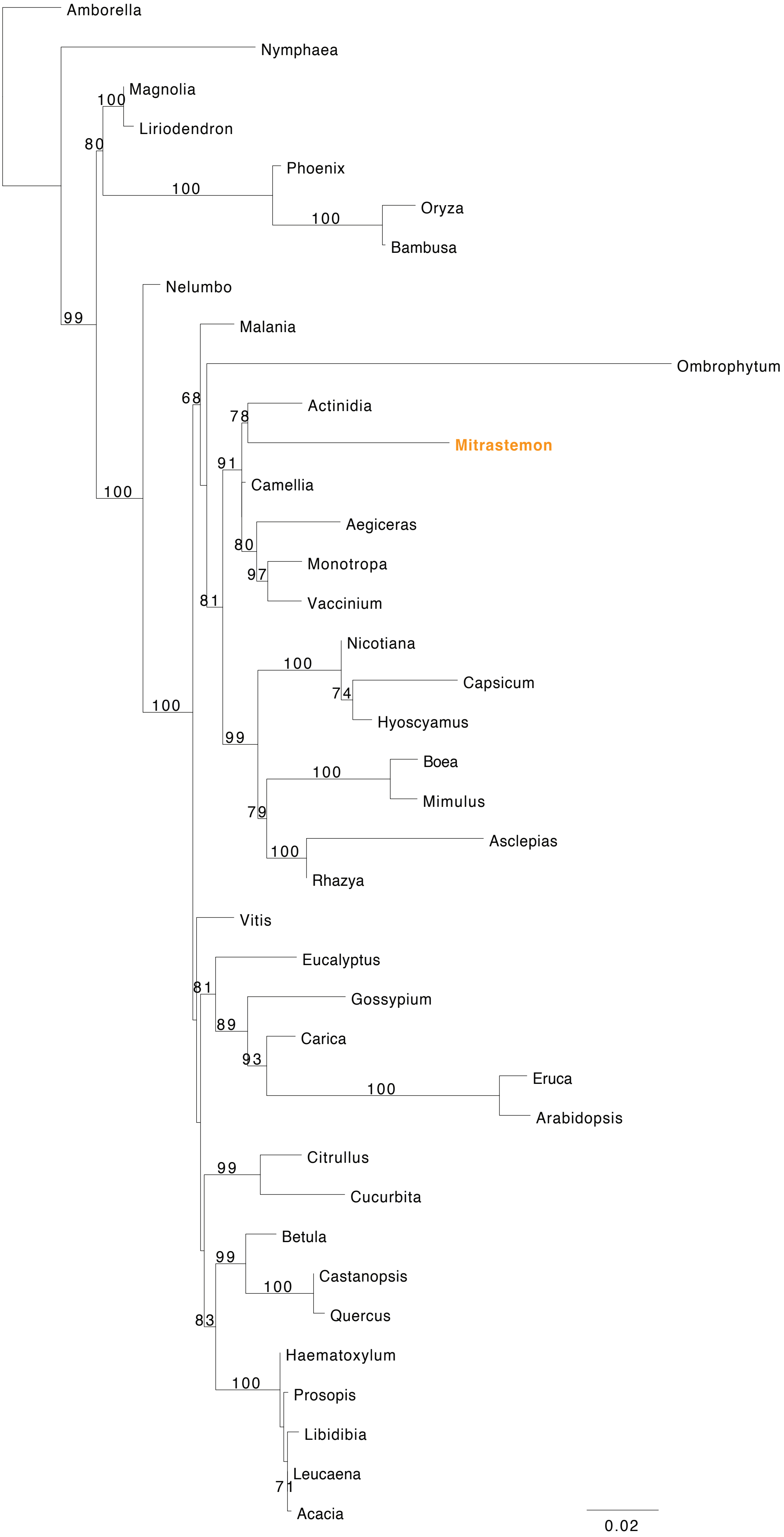
Amborella
Nymphaea
Magnolia
100
Liriodendron
80
Phoenix
100
Oryza
100
Bambusa
Nelumbo
99
Malania
Ombrophytum
68
Actinidia
78
Mitrastemon
91
Camellia
100
Aegiceras
80
Monotropa
97
Vaccinium
81
Nicotiana
100
Capsicum
74
Hyoscyamus
100
99
Boea
100
Mimulus
79
Asclepias
100
Rhazya
Vitis
Eucalyptus
81
Gossypium
89
Carica
93
Eruca
100
Arabidopsis
Citrullus
99
Cucurbita
Betula
99
Castanopsis
100
Quercus
83
Haematoxylum
100
Prosopis
Libidibia
Leucaena
71
Acacia
0.02

### *rpl5*

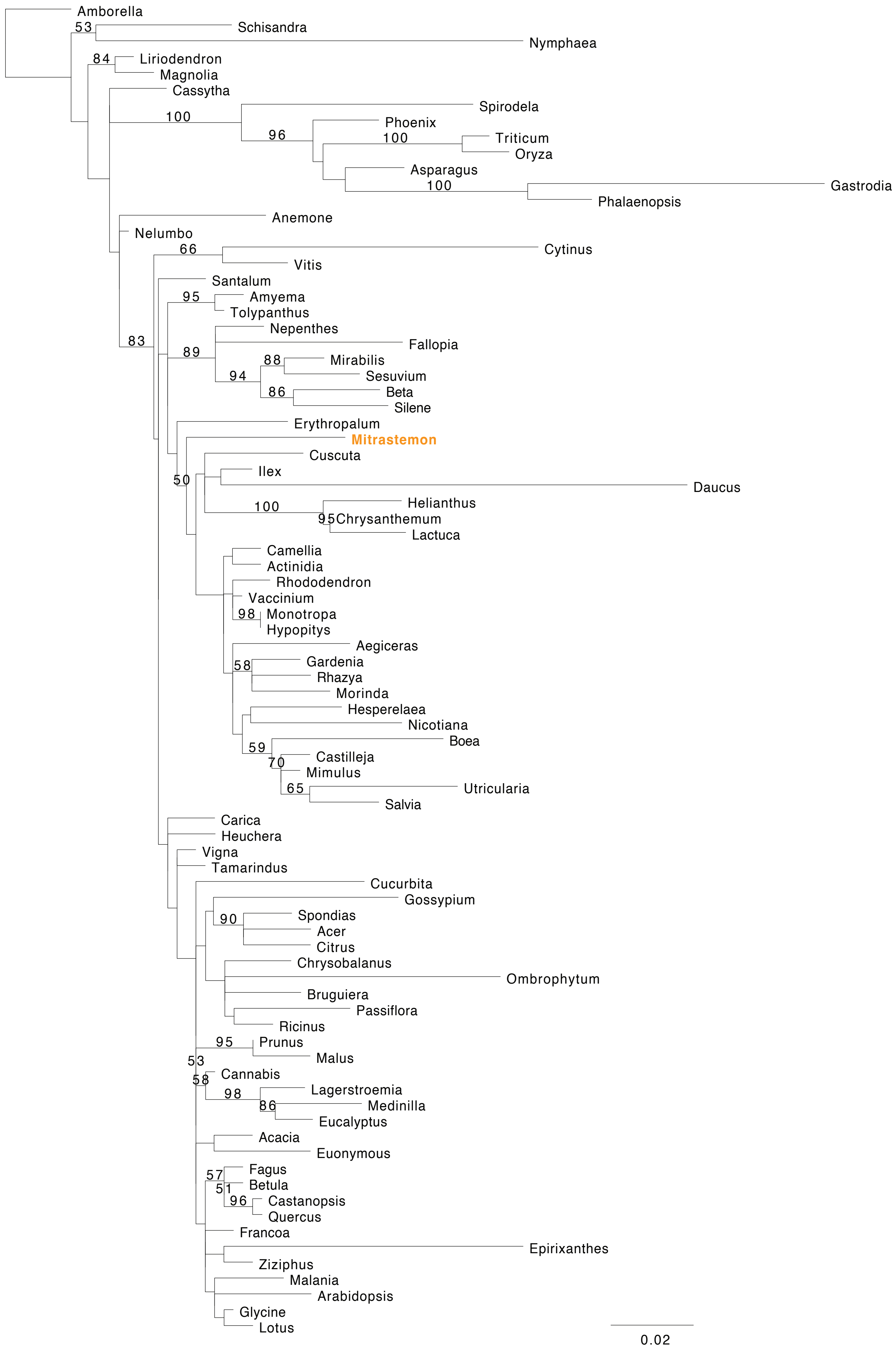

Amborella
53
Schisandra
Nymphaea
84
Liriodendron
Magnolia
Cassytha
Spirodela
100
Phoenix
96
Triticum
100
Oryza
Asparagus
100
Gastrodia
Phalaenopsis
Anemone
Nelumbo
66
Cytinus
Vitis
Santalum
95
Amyema
Tolypanthus
Nepenthes
83
Fallopia
89
88
Mirabilis
94
Sesuvium
86
Beta
Silene
Erythropalum
Mitrastemon
Cuscuta
50
Ilex
Daucus
100
Helianthus
95
Chrysanthemum
Lactuca
Camellia
Actinidia
Rhododendron
Vaccinium
98
Monotropa
Hypopitys
Aegiceras
58
Gardenia
Rhazya
Morinda
Hesperelaea
Nicotiana
59
Boea
70
Castilleja
Mimulus
65
Utricularia
Salvia
Carica
Heuchera
Vigna
Tamarindus
Cucurbita
Gossypium
90
Spondias
Acer
Citrus
Chrysobalanus
Ombrophytum
Bruguiera
Passiflora
Ricinus
95
Prunus
53
Malus
58
Cannabis
98
Lagerstroemia
86
Medinilla
Eucalyptus
Acacia
Euonymous
57
Fagus
51
Betula
96
Castanopsis
Quercus
Francoa
Epirixanthes
Ziziphus
Malania
Arabidopsis
Glycine
Lotus
0.02

### *rpl10*

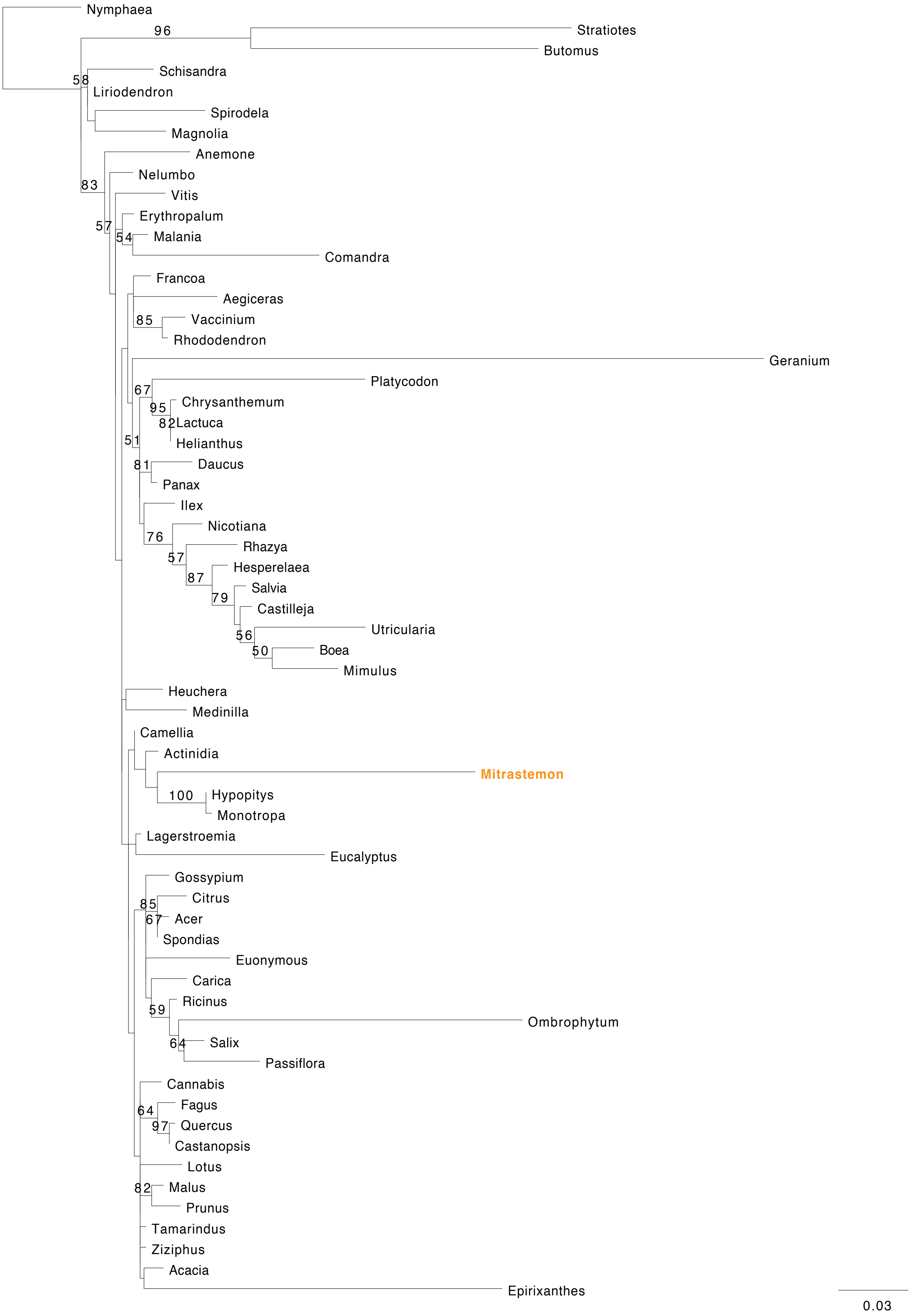
Nymphaea
96
Stratiotes
Butomus
58
Schisandra
Liriodendron
Spirodela
Magnolia
Anemone
83
Nelumbo
Vitis
57
Erythropalum
54
Malania
Comandra
Francoa
Aegiceras
85
Vaccinium
Rhododendron
Geranium
67
Platycodon
95
Chrysanthemum
82
Lactuca
51
Helianthus
81
Daucus
Panax
Ilex
76
Nicotiana
57
Rhazya
87
Hesperelaea
79
Salvia
Castilleja
56
Utricularia
50
Boea
Mimulus
Heuchera
Medinilla
Camellia
Actinidia
Mitrastemon
100
Hypopitys
Monotropa
Lagerstroemia
Eucalyptus
Gossypium
85
Citrus
67
Acer
Spondias
Euonymous
Carica
59
Ricinus
Ombrophytum
64
Salix
Passiflora
Cannabis
64
Fagus
97
Quercus
Castanopsis
Lotus
82
Malus
Prunus
Tamarindus
Ziziphus
Acacia
Epirixanthes
0.03

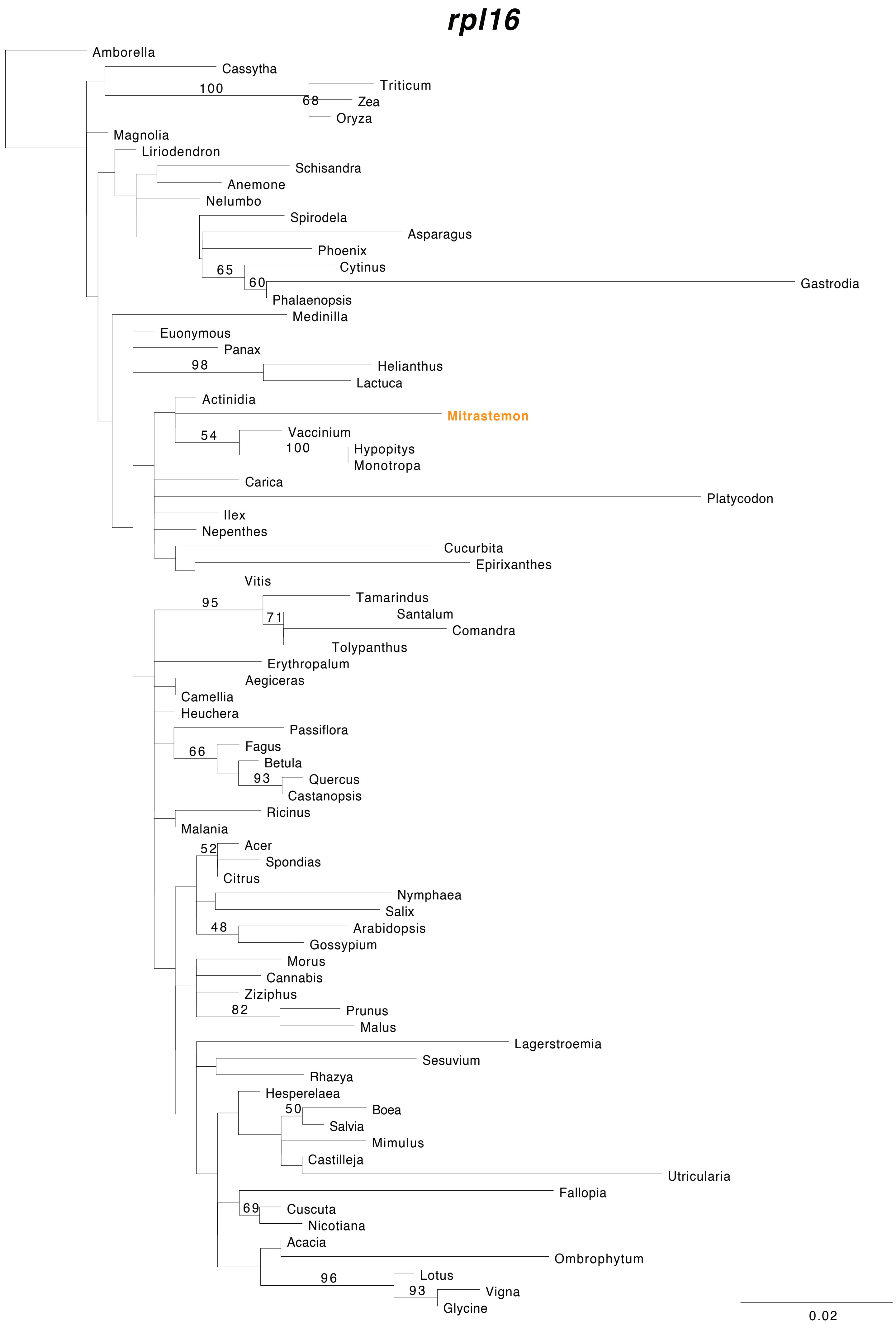
rpl16
Amborella
Cassytha
100
Triticum
68
Zea
Oryza
Magnolia
Liriodendron
Schisandra
Anemone
Nelumbo
Spirodela
Asparagus
Phoenix
65
Cytinus
60
Gastrodia
Phalaenopsis
Medinilla
Euonymous
Panax
98
Helianthus
Lactuca
Actinidia
Mitrastemon
54
Vaccinium
100
Hypopitys
Monotropa
Carica
Platycodon
Ilex
Nepenthes
Cucurbita
Epirixanthes
Vitis
95
Tamarindus
71
Santalum
Comandra
Tolypanthus
Erythropalum
Aegiceras
Camellia
Heuchera
Passiflora
66
Fagus
Betula
93
Quercus
Castanopsis
Ricinus
Malania
52
Acer
Spondias
Citrus
Nymphaea
Salix
48
Arabidopsis
Gossypium
Morus
Cannabis
Ziziphus
82
Prunus
Malus
Lagerstroemia
Sesuvium
Rhazya
Hesperelaea
50
Boea
Salvia
Mimulus
Castilleja
Utricularia
Fallopia
69
Cuscuta
Nicotiana
Acacia
Ombrophytum
96
Lotus
93
Vigna
Glycine
0.02

### *rps1*

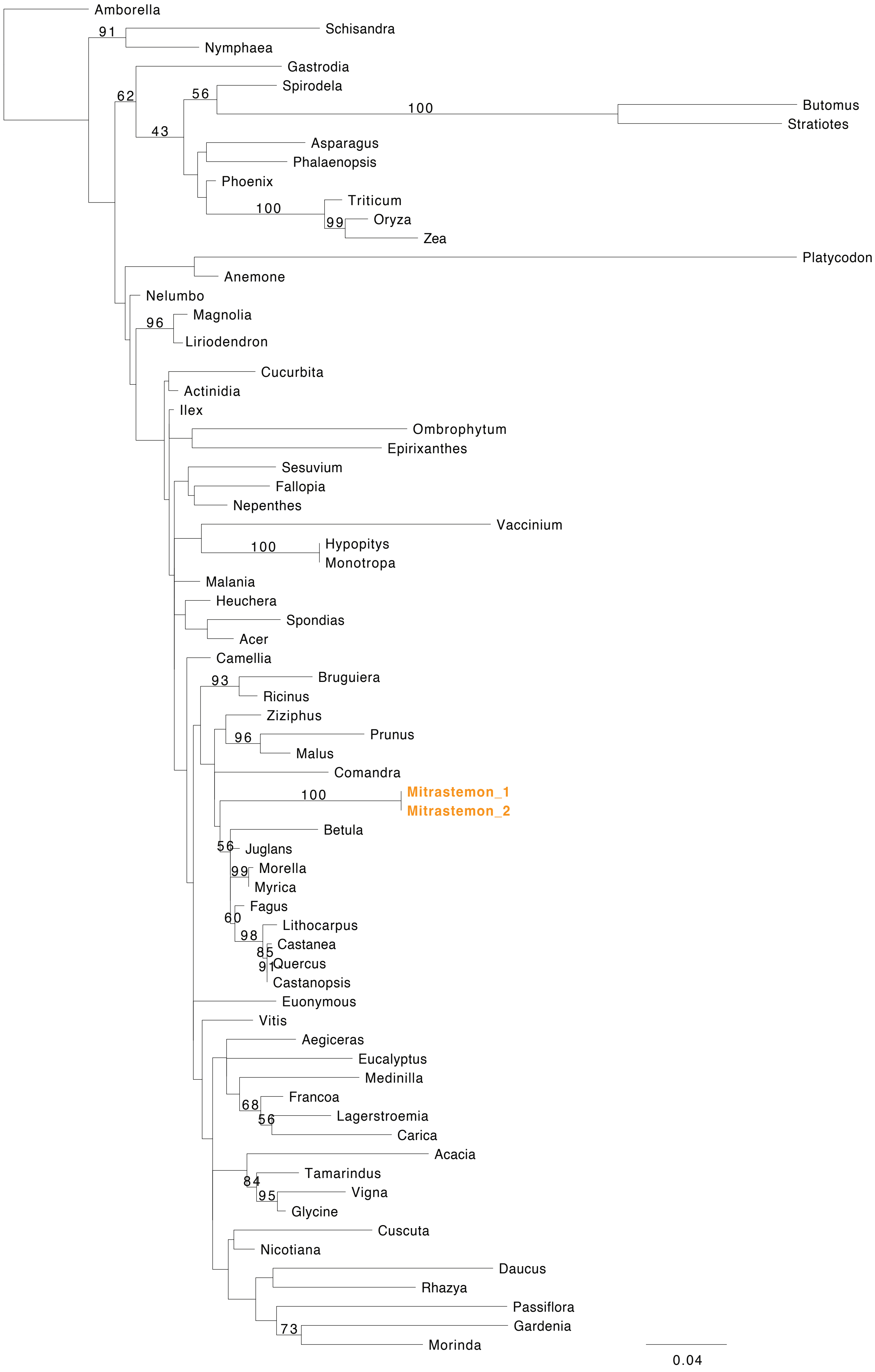
Amborella
91
Schisandra
Nymphaea
Gastrodia
Spirodela
62
56
100
Butomus
Stratiotes
43
Asparagus
Phalaenopsis
Phoenix
Triticum
100
99
Oryza
Zea
Platycodon
Anemone
Nelumbo
Magnolia
96
Liriodendron
Cucurbita
Actinidia
Ilex
Ombrophytum
Epirixanthes
Sesuvium
Fallopia
Nepenthes
Vaccinium
100
Hypopitys
Monotropa
Malania
Heuchera
Spondias
Acer
Camellia
93
Bruguiera
Ricinus
Ziziphus
96
Prunus
Malus
Comandra
100
Mitrastemon_1
Mitrastemon_2
Betula
56
Juglans
99
Morella
Myrica
Fagus
60
Lithocarpus
98
Castanea
85
91
Quercus
Castanopsis
Euonymous
Vitis
Aegiceras
Eucalyptus
Medinilla
Francoa
68
56
Lagerstroemia
Carica
Acacia
Tamarindus
84
95
Vigna
Glycine
Cuscuta
Nicotiana
Daucus
Rhazya
Passiflora
Gardenia
73
Morinda
0.04

### *rps3*

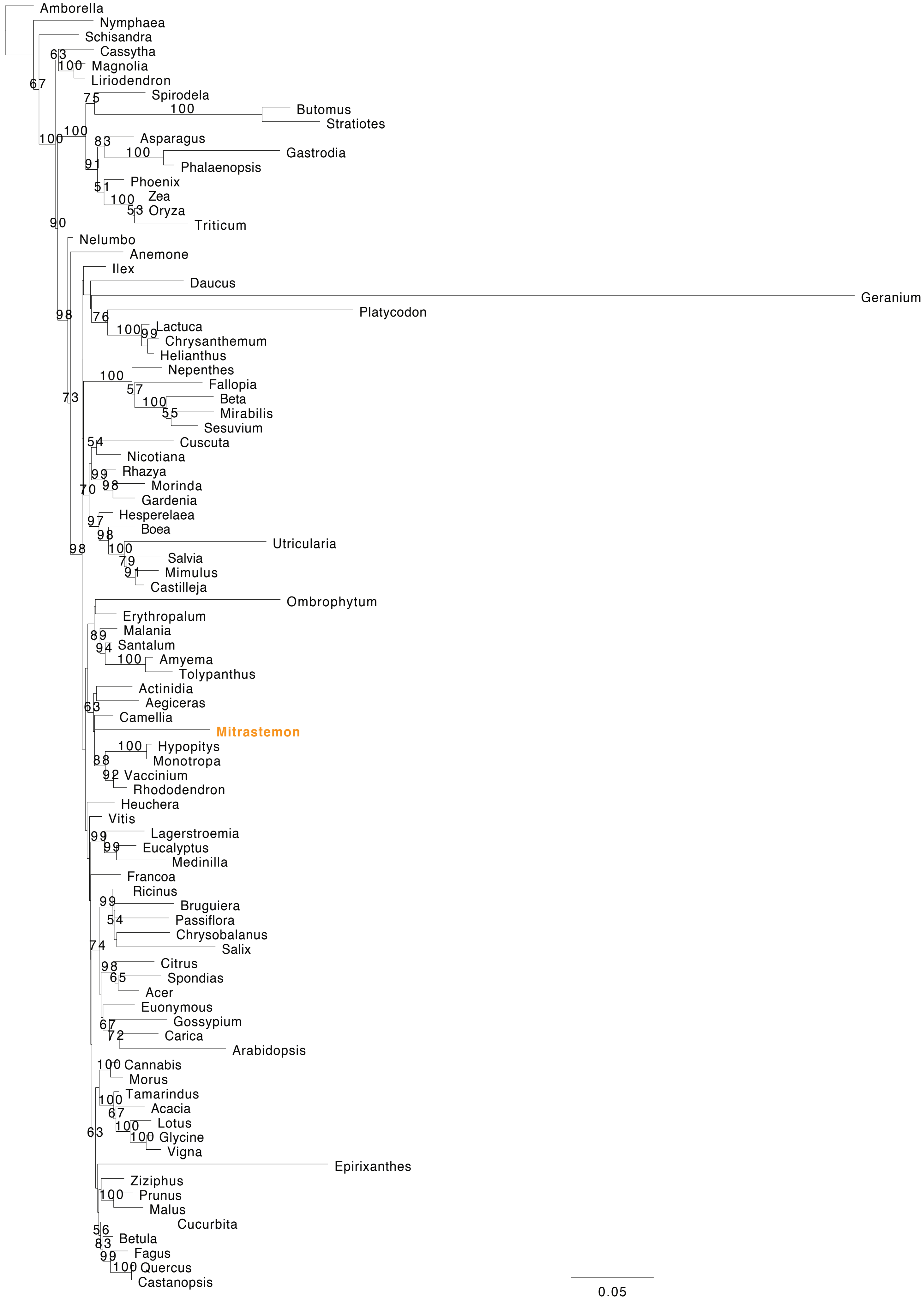

Amborella
Nymphaea
Schisandra
Cassytha
Magnolia
Liriodendron
Spirodela
Butomus
Stratiotes
Asparagus
Gastrodia
Phalaenopsis
Phoenix
Zea
Oryza
Triticum
Nelumbo
Anemone
Ilex
Daucus
Geranium
Platycodon
Lactuca
Chrysanthemum
Helianthus
Nepenthes
Fallopia
Beta
Mirabilis
Sesuvium
Cuscuta
Nicotiana
Rhazya
Morinda
Gardenia
Hesperelaea
Boea
Utricularia
Salvia
Mimulus
Castilleja
Ombrophytum
Erythropalum
Malania
Santalum
Amyema
Tolypanthus
Actinidia
Aegiceras
Camellia
Mitrastemon
Hypopitys
Monotropa
Vaccinium
Rhododendron
Heuchera
Vitis
Lagerstroemia
Eucalyptus
Medinilla
Francoa
Ricinus
Bruguiera
Passiflora
Chrysobalanus
Salix
Citrus
Spondias
Acer
Euonymous
Gossypium
Carica
Arabidopsis
Cannabis
Morus
Tamarindus
Acacia
Lotus
Glycine
Vigna
Epirixanthes
Ziziphus
Prunus
Malus
Cucurbita
Betula
Fagus
Quercus
Castanopsis
0.05

### *rps4*

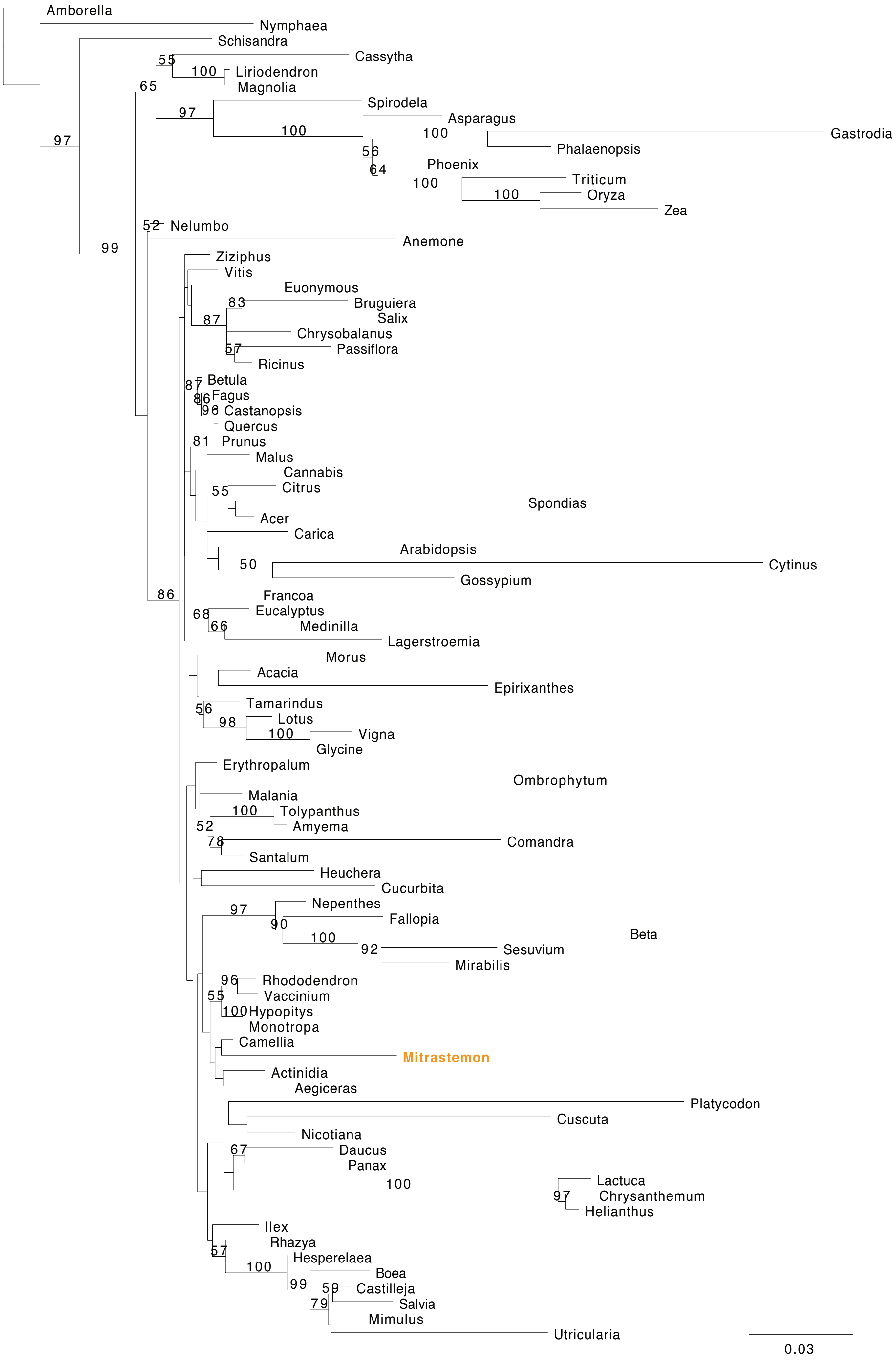

Amborella
Nymphaea
Schisandra
Cassytha
Liriodendron
Magnolia
Spirodela
Asparagus
Gastrodia
Phalaenopsis
Phoenix
Triticum
Oryza
Zea
Nelumbo
Anemone
Ziziphus
Vitis
Euonymous
Bruguiera
Salix
Chrysobalanus
Passiflora
Ricinus
Betula
Fagus
Castanopsis
Quercus
Prunus
Malus
Cannabis
Citrus
Spondias
Acer
Carica
Arabidopsis
Cytinus
Gossypium
Francoa
Eucalyptus
Medinilla
Lagerstroemia
Morus
Acacia
Epirixanthes
Tamarindus
Lotus
Vigna
Glycine
Erythropalum
Ombrophytum
Malania
Tolypanthus
Amyema
Comandra
Santalum
Heuchera
Cucurbita
Nepenthes
Fallopia
Beta
Sesuvium
Mirabilis
Rhododendron
Vaccinium
Hypopitys
Monotropa
Camellia
Mitrastemon
Actinidia
Aegiceras
Platycodon
Cuscuta
Nicotiana
Daucus
Panax
Lactuca
Chrysanthemum
Helianthus
Ilex
Rhazya
Hesperelaea
Boea
Castilleja
Salvia
Mimulus
Utricularia
55
100
65
97
100
100
56
64
100
100
97
52
99
83
87
57
87
86
96
81
55
50
86
68
66
56
98
100
100
52
78
97
90
100
92
96
55
100
67
100
97
57
100
99
59
79
0.03

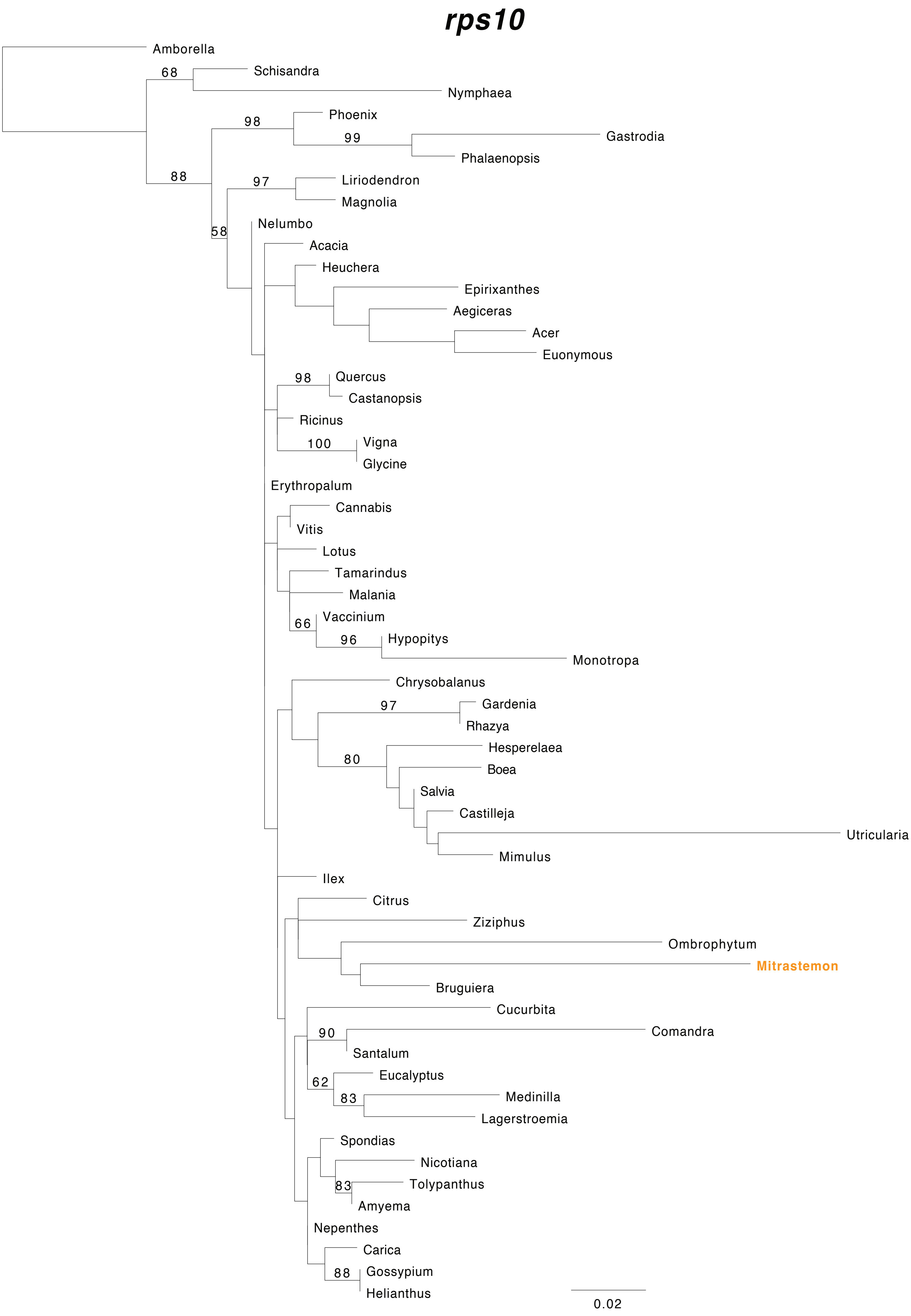
rps10
Amborella
Schisandra
Nymphaea
Phoenix
Gastrodia
Phalaenopsis
Liriodendron
Magnolia
Nelumbo
Acacia
Heuchera
Epirixanthes
Aegiceras
Acer
Euonymous
Quercus
Castanopsis
Ricinus
Vigna
Glycine
Erythropalum
Cannabis
Vitis
Lotus
Tamarindus
Malania
Vaccinium
Hypopitys
Monotropa
Chrysobalanus
Gardenia
Rhazya
Hesperelaea
Boea
Salvia
Castilleja
Utricularia
Mimulus
Ilex
Citrus
Ziziphus
Ombrophytum
Mitrastemon
Bruguiera
Cucurbita
Comandra
Santalum
Eucalyptus
Medinilla
Lagerstroemia
Spondias
Nicotiana
Tolypanthus
Amyema
Nepenthes
Carica
Gossypium
Helianthus
68
98
99
88
97
58
98
100
66
96
97
80
90
62
83
83
88
0.02

#### *rps12*

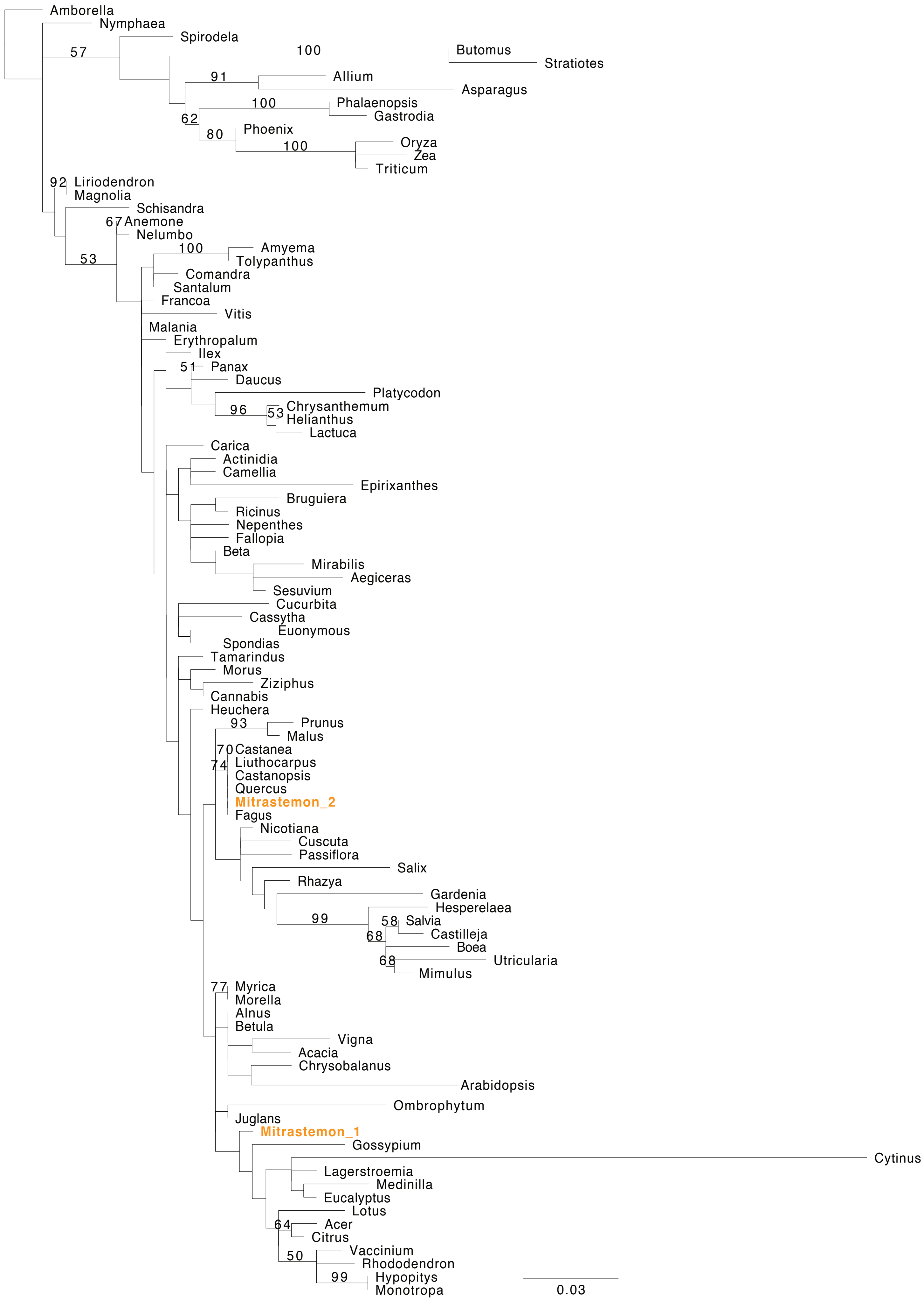
Amborella
Nymphaea
Spirodela
Butomus
Stratiotes
Allium
Asparagus
Phalaenopsis
Gastrodia
Phoenix
Oryza
Zea
Triticum
Liriodendron
Magnolia
Schisandra
Anemone
Nelumbo
Amyema
Tolypanthus
Comandra
Santalum
Francoa
Vitis
Malania
Erythropalum
Ilex
Panax
Daucus
Platycodon
Chrysanthemum
Helianthus
Lactuca
Carica
Actinidia
Camellia
Epirixanthes
Bruguiera
Ricinus
Nepenthes
Fallopia
Beta
Mirabilis
Aegiceras
Sesuvium
Cucurbita
Cassytha
Euonymous
Spondias
Tamarindus
Morus
Ziziphus
Cannabis
Heuchera
Prunus
Malus
Castanea
Liuthocarpus
Castanopsis
Quercus
Mitrastemon_2
Fagus
Nicotiana
Cuscuta
Passiflora
Salix
Rhazya
Gardenia
Hesperelaea
Salvia
Castilleja
Boea
Utricularia
Mimulus
Myrica
Morella
Alnus
Betula
Vigna
Acacia
Chrysobalanus
Arabidopsis
Ombrophytum
Juglans
Mitrastemon_1
Gossypium
Cytinus
Lagerstroemia
Medinilla
Eucalyptus
Lotus
Acer
Citrus
Vaccinium
Rhododendron
Hypopitys
Monotropa
57
100
91
100
62
80
100
92
67
53
100
51
96
53
93
70
74
99
58
68
68
77
64
50
99
0.03

### rps13

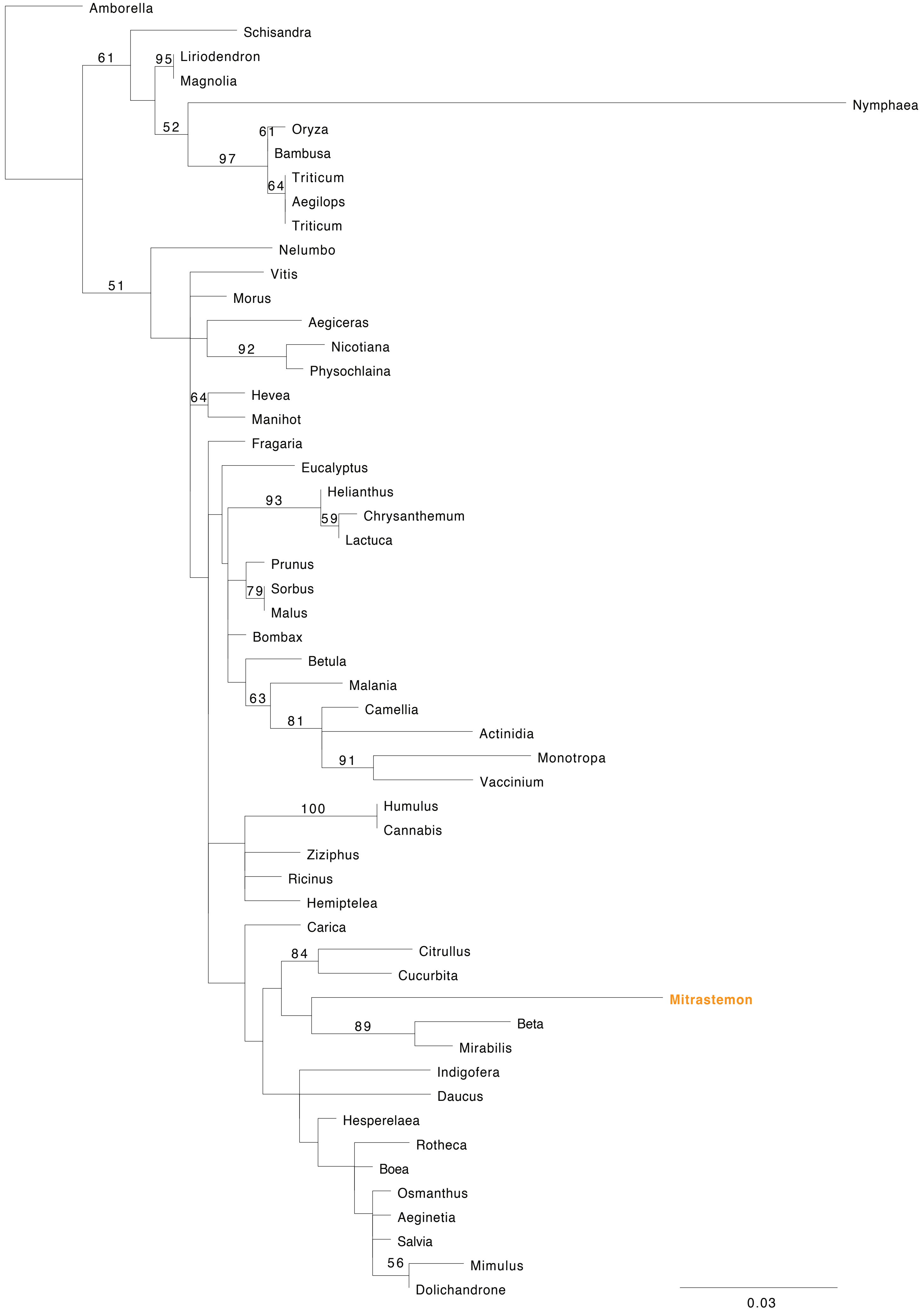
Amborella
Schisandra
61
95
Liriodendron
Magnolia
Nymphaea
52
61
Oryza
Bambusa
97
64
Triticum
Aegilops
Triticum
Nelumbo
Vitis
51
Morus
Aegiceras
92
Nicotiana
Physochlaina
64
Hevea
Manihot
Fragaria
Eucalyptus
93
Helianthus
59
Chrysanthemum
Lactuca
Prunus
79
Sorbus
Malus
Bombax
Betula
Malania
63
Camellia
81
Actinidia
91
Monotropa
Vaccinium
100
Humulus
Cannabis
Ziziphus
Ricinus
Hemiptelea
Carica
84
Citrullus
Cucurbita
Mitrastemon
89
Beta
Mirabilis
Indigofera
Daucus
Hesperelaea
Rotheca
Boea
Osmanthus
Aeginetia
Salvia
56
Mimulus
Dolichandrone
0.03

### *rps19*

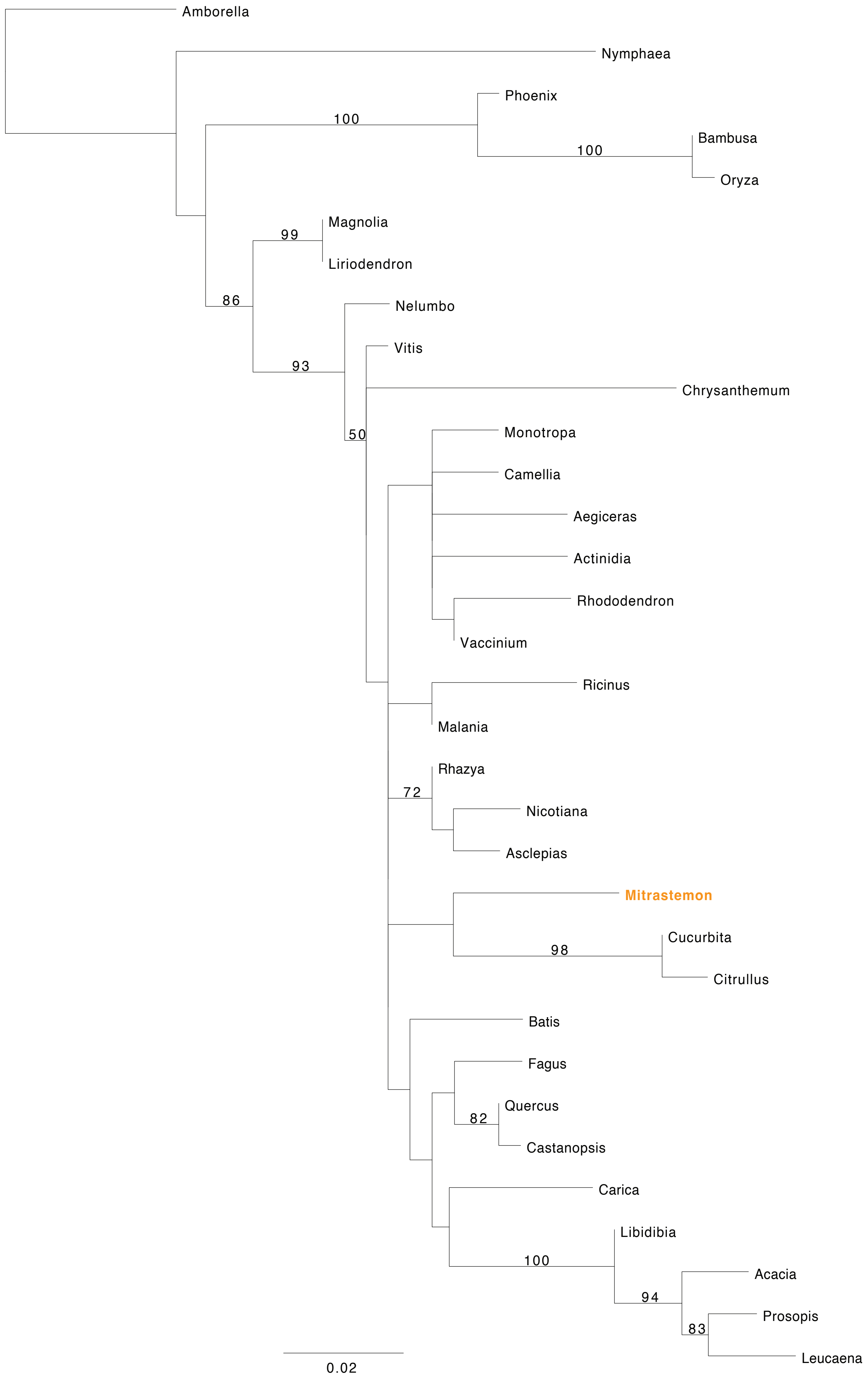
Amborella
Nymphaea
Phoenix
100
Bambusa
100
Oryza
Magnolia
99
Liriodendron
86
Nelumbo
Vitis
93
Chrysanthemum
50
Monotropa
Camellia
Aegiceras
Actinidia
Rhododendron
Vaccinium
Ricinus
Malania
Rhazya
72
Nicotiana
Asclepias
Mitrastemon
Cucurbita
98
Citrullus
Batis
Fagus
Quercus
82
Castanopsis
Carica
Libidibia
100
Acacia
94
Prosopis
83
Leucaena
0.02

### *sdh3*

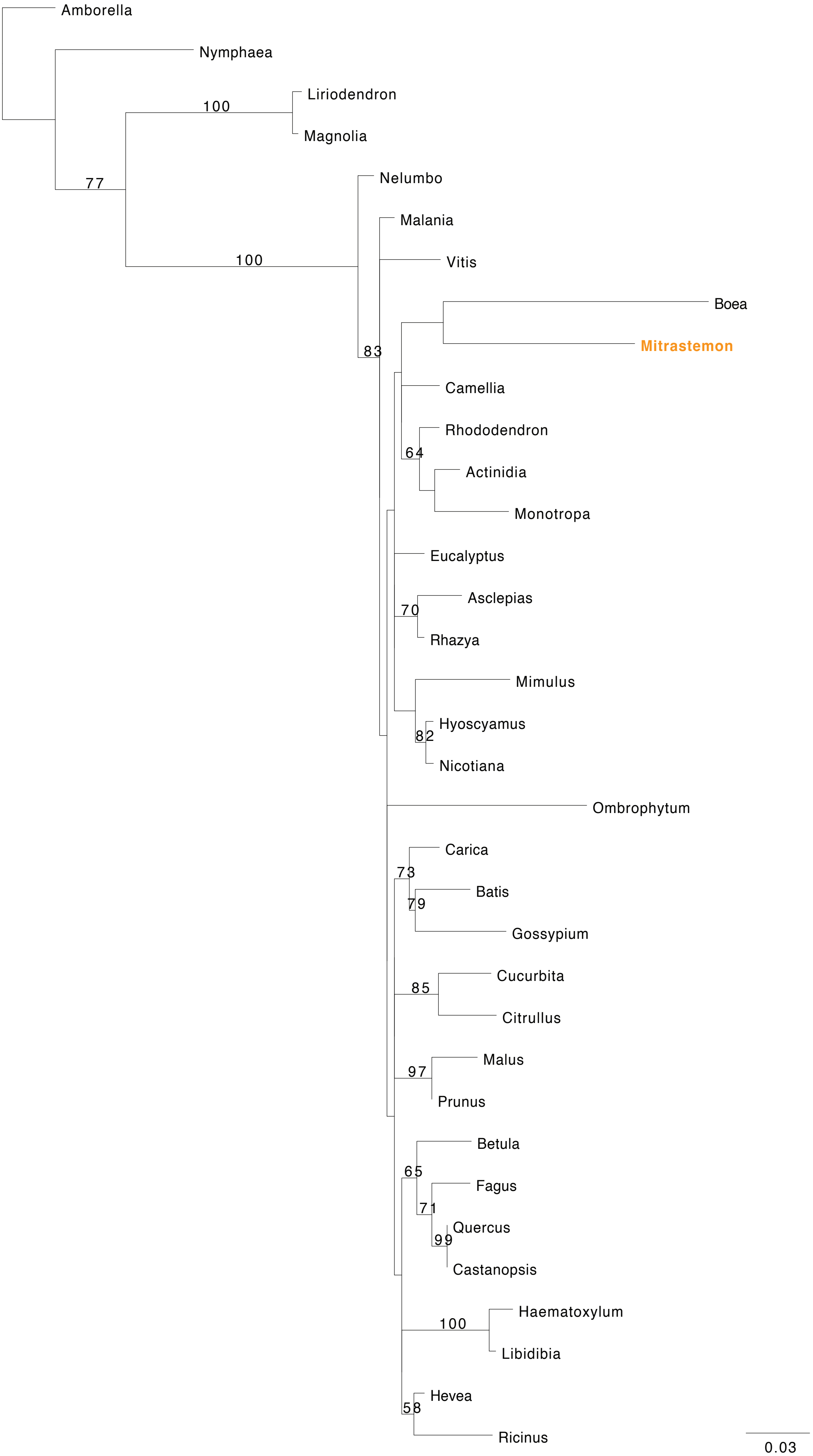
Amborella
Nymphaea
Liriodendron
Magnolia
Nelumbo
Malania
Vitis
Boea
Mitrastemon
Camellia
Rhododendron
Actinidia
Monotropa
Eucalyptus
Asclepias
Rhazya
Mimulus
Hyoscyamus
Nicotiana
Ombrophytum
Carica
Batis
Gossypium
Cucurbita
Citrullus
Malus
Prunus
Betula
Fagus
Quercus
Castanopsis
Haematoxylum
Libidibia
Hevea
Ricinus
100
77
100
83
64
70
82
73
79
85
97
65
71
99
100
58
0.03

### ***sdh4***

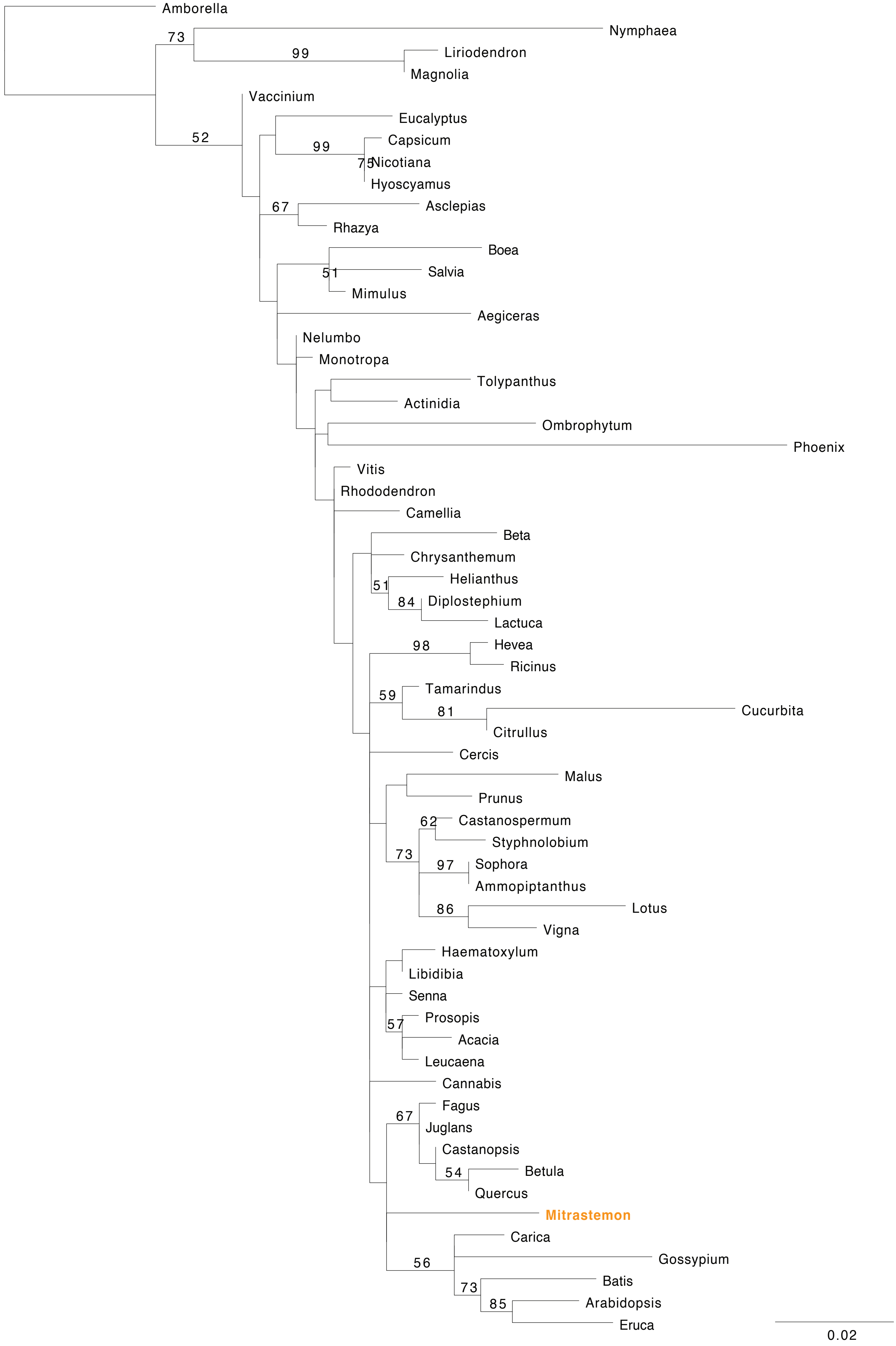

Amborella
73
Nymphaea
99
Liriodendron
Magnolia
Vaccinium
52
Eucalyptus
99
Capsicum
75
Nicotiana
Hyoscyamus
67
Asclepias
Rhazya
Boea
51
Salvia
Mimulus
Aegiceras
Nelumbo
Monotropa
Tolypanthus
Actinidia
Ombrophytum
Phoenix
Vitis
Rhododendron
Camellia
Beta
Chrysanthemum
51
Helianthus
84
Diplostephium
Lactuca
98
Hevea
Ricinus
59
Tamarindus
81
Cucurbita
Citrullus
Cercis
Malus
Prunus
62
Castanospermum
Styphnolobium
73
97
Sophora
Ammopiptanthus
86
Lotus
Vigna
Haematoxylum
Libidibia
Senna
57
Prosopis
Acacia
Leucaena
Cannabis
67
Fagus
Juglans
Castanopsis
54
Betula
Quercus
Mitrastemon
Carica
56
Gossypium
73
Batis
85
Arabidopsis
Eruca
0.02

**Figure S3.** Visualization of BLASTn searches of the mitochondrial chromosomes in *Mitrastemon yamamotoi*. Circular chromosomes are shown linearized for clarity. BLASTn hits between *Mitrastemon* and a custom angiosperm mitochondrial database are organized in groups as described in Table S5: *Castanopsis*, Fagaceae, Fagales, Ericales, and those of other angiosperms. The three rows represent all the BLASTn hits (e-value < $2x10^{-10}$). Colors depict sequence identity of BLASTn hits according to the scale shown on the right. Arrows below each coding region represent the direction of transcription.

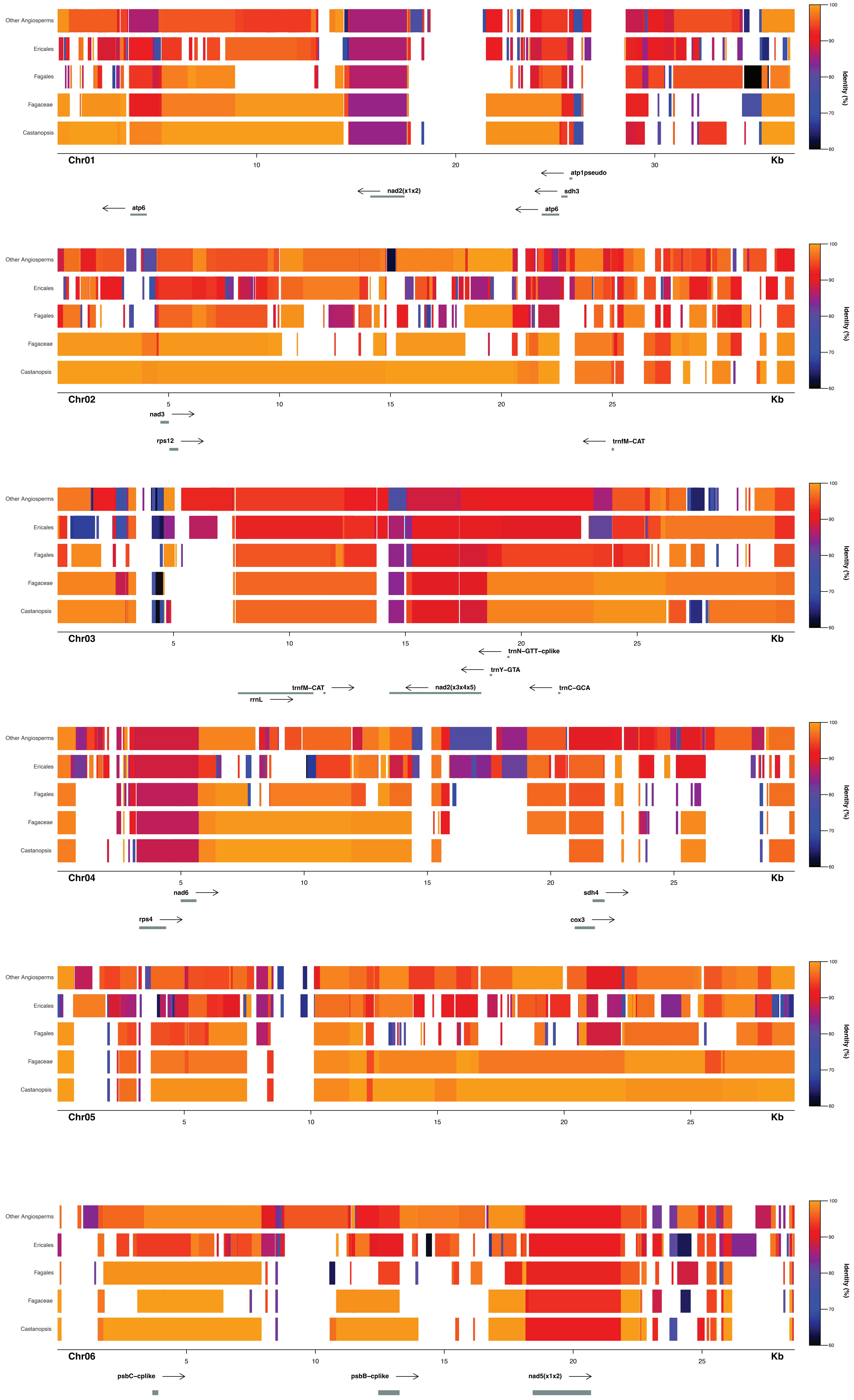

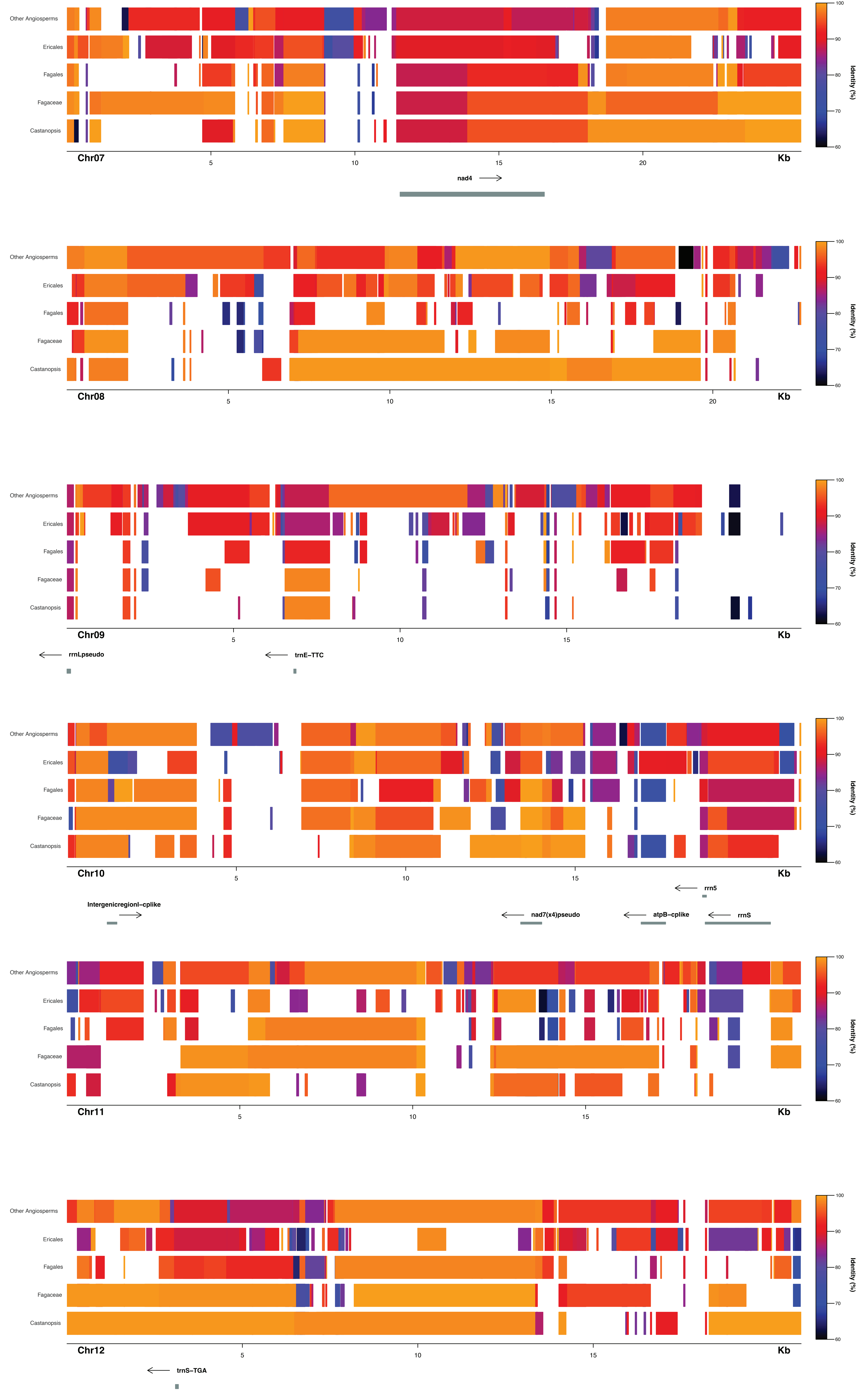
Other Angiosperms
Ericales
Fagales
Fagaceae
Castanopsis
Chr07
5
10
15
20
Kb
nad4
Identity (%)
100
90
80
70
60
Chr08
Chr09
rrnLpseudo
trnE–TTC
Chr10
rrn5
Intergenicregionl–cplike
nad7(x4)pseudo
atpB–cplike
rrnS
Chr11
Chr12
trnS–TGA

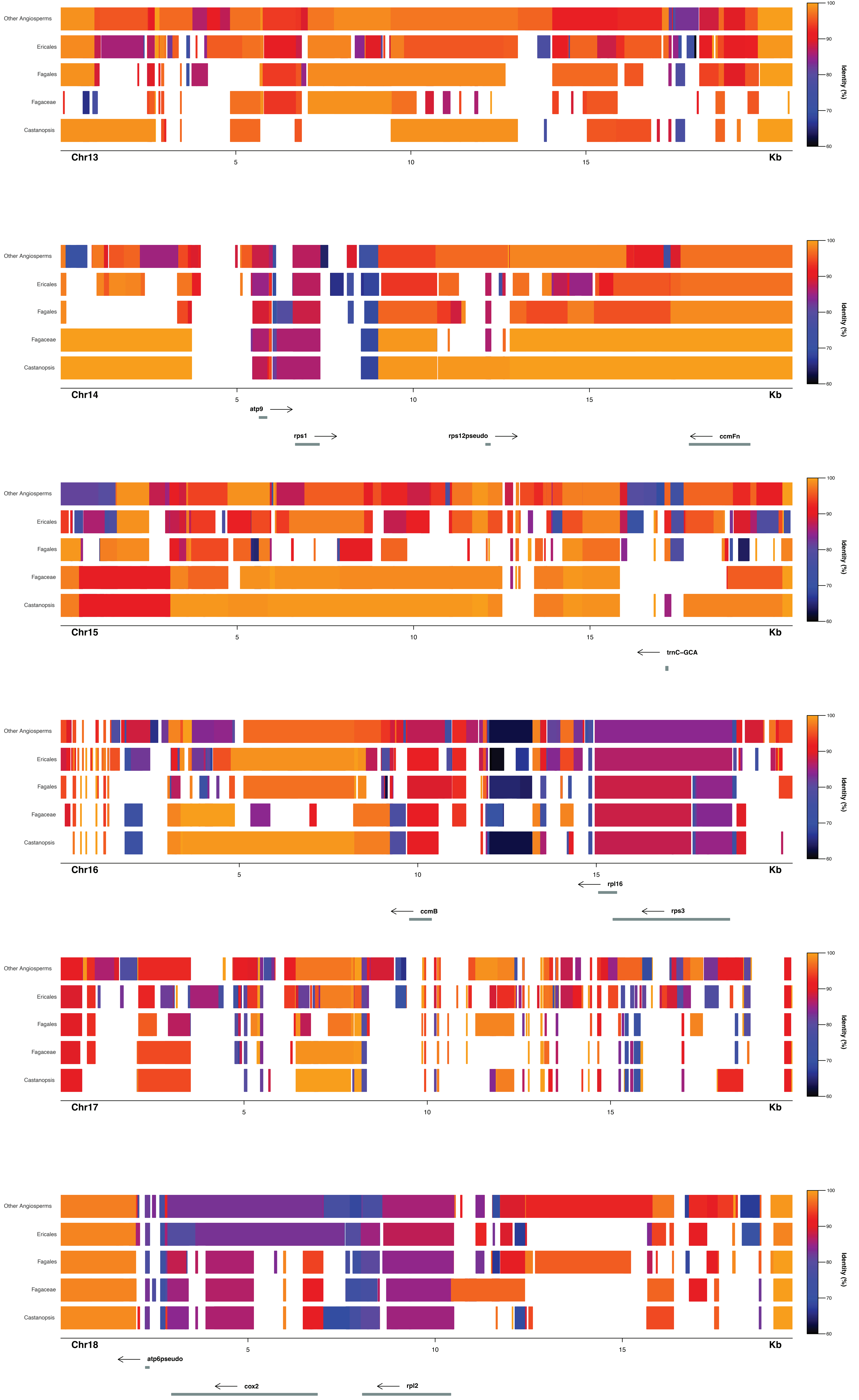
Other Angiosperms
Ericales
Fagales
Fagaceae
Castanopsis
Chr13
5
10
15
Kb
Identity (%)
100
90
80
70
60
Chr14
atp9
rps1
rps12pseudo
ccmFn
Chr15
trnC–GCA
Chr16
rpl16
ccmB
rps3
Chr17
Chr18
atp6pseudo
cox2
rpl2

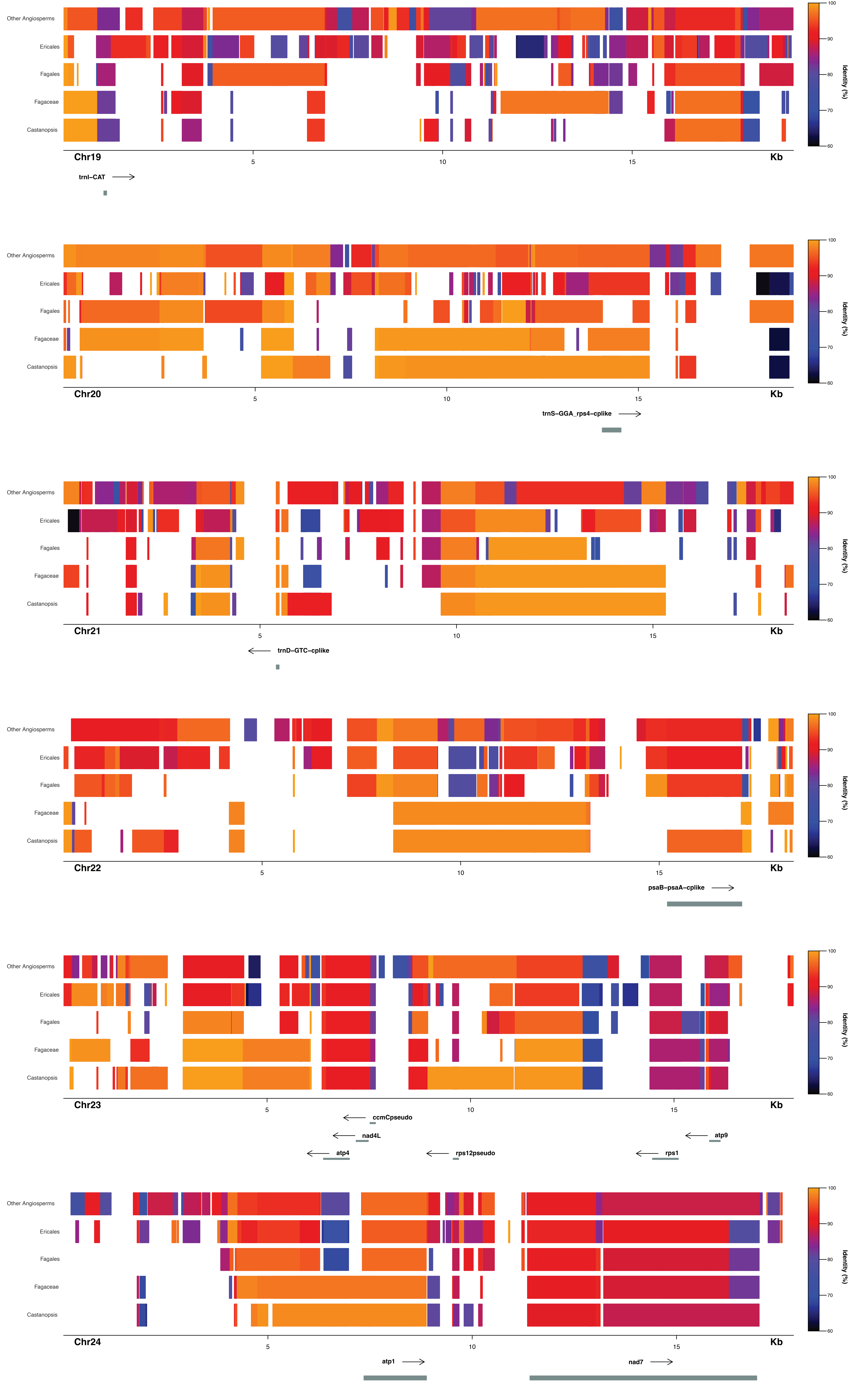
Other Angiosperms
Ericales
Fagales
Fagaceae
Castanopsis
Chr19
5
10
15
Kb
trnI–CAT
Identity (%)
100
90
80
70
60
Other Angiosperms
Ericales
Fagales
Fagaceae
Castanopsis
Chr20
5
10
15
Kb
trnS–GGA_rps4–cplike
Identity (%)
100
90
80
70
60
Other Angiosperms
Ericales
Fagales
Fagaceae
Castanopsis
Chr21
5
10
15
Kb
trnD–GTC–cplike
Identity (%)
100
90
80
70
60
Other Angiosperms
Ericales
Fagales
Fagaceae
Castanopsis
Chr22
5
10
15
Kb
psaB–psaA–cplike
Identity (%)
100
90
80
70
60
Other Angiosperms
Ericales
Fagales
Fagaceae
Castanopsis
Chr23
5
10
15
Kb
ccmCpseudo
nad4L
atp9
atp4
rps12pseudo
rps1
Identity (%)
100
90
80
70
60
Other Angiosperms
Ericales
Fagales
Fagaceae
Castanopsis
Chr24
5
10
15
Kb
atp1
nad7
Identity (%)
100
90
80
70
60

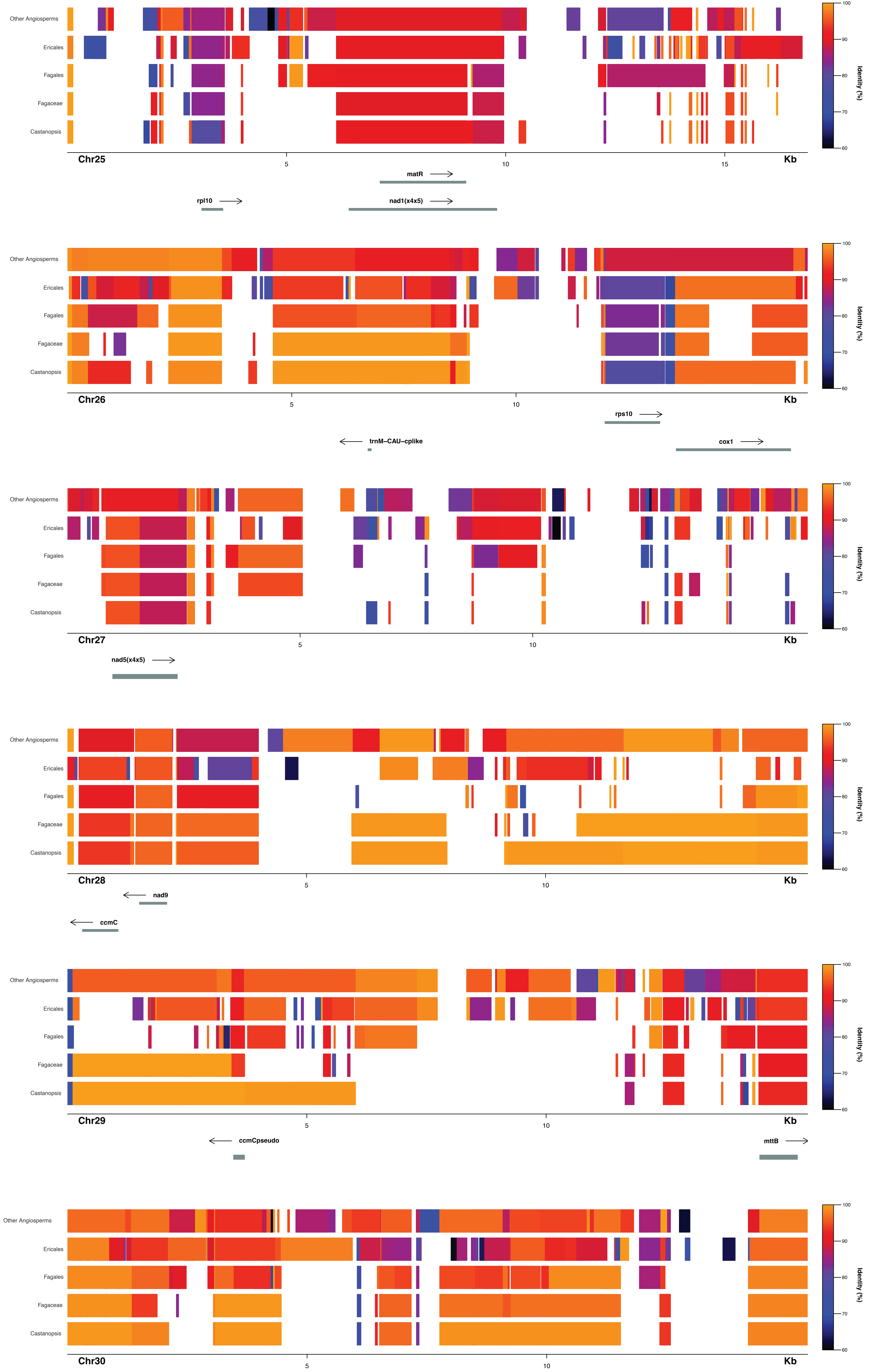

Other Angiosperms
Ericales
Fagales
Fagaceae
Castanopsis
Chr25
Kb
matR
rpl10
nad1(x4x5)
Chr26
rps10
trnM-CAU-cplike
cox1
Chr27
nad5(x4x5)
Chr28
nad9
ccmC
Chr29
ccmCpseudo
mttB
Chr30
Identity (%)

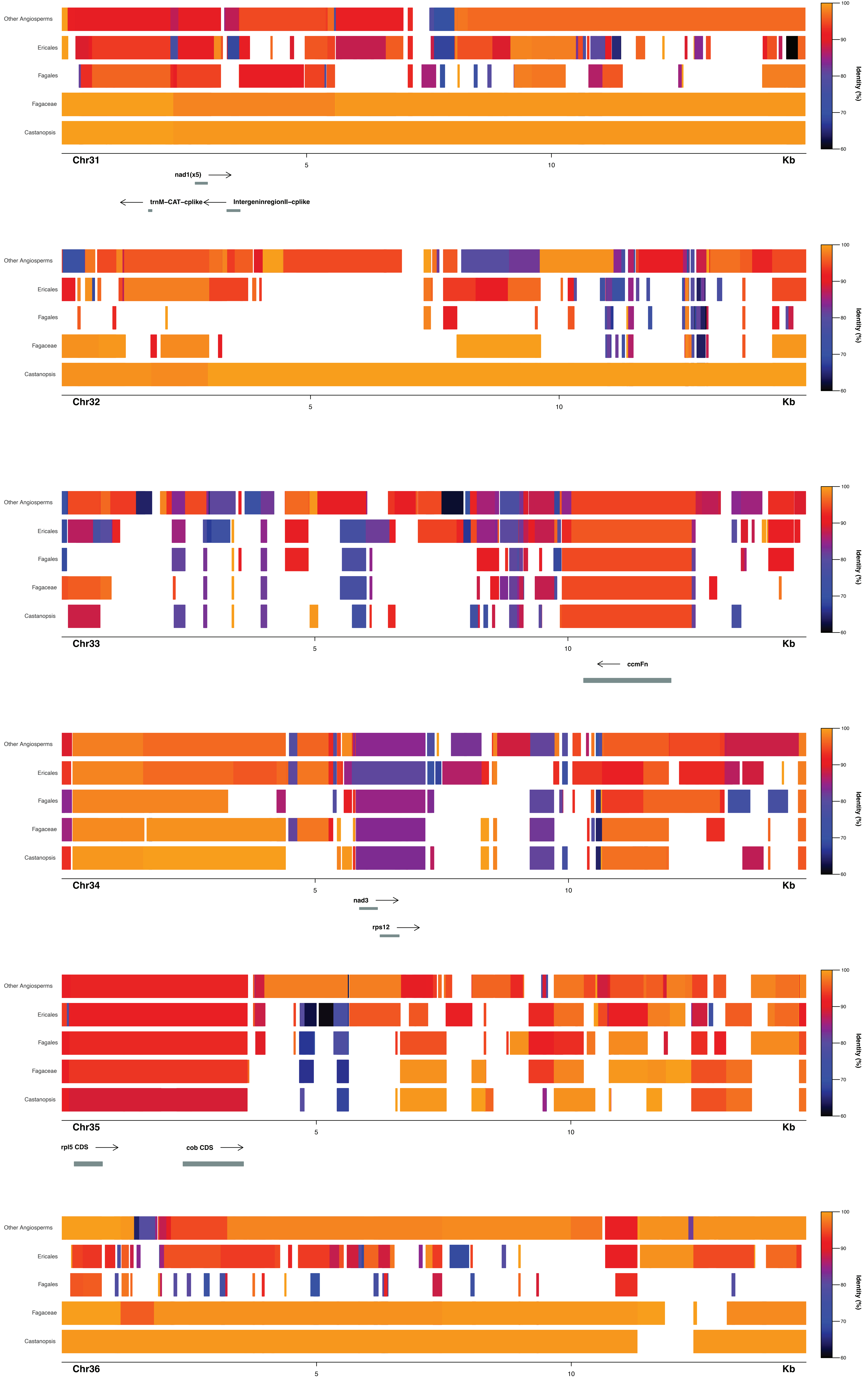
Other Angiosperms
Ericales
Fagales
Fagaceae
Castanopsis
Chr31
5
10
Kb
nad1(x5)
trnM–CAT–cplike
IntergeninregionII–cplike
Chr32
Chr33
ccmFn
Chr34
nad3
rps12
Chr35
rpl5 CDS
cob CDS
Chr36
Identity (%)
100
90
80
70
60

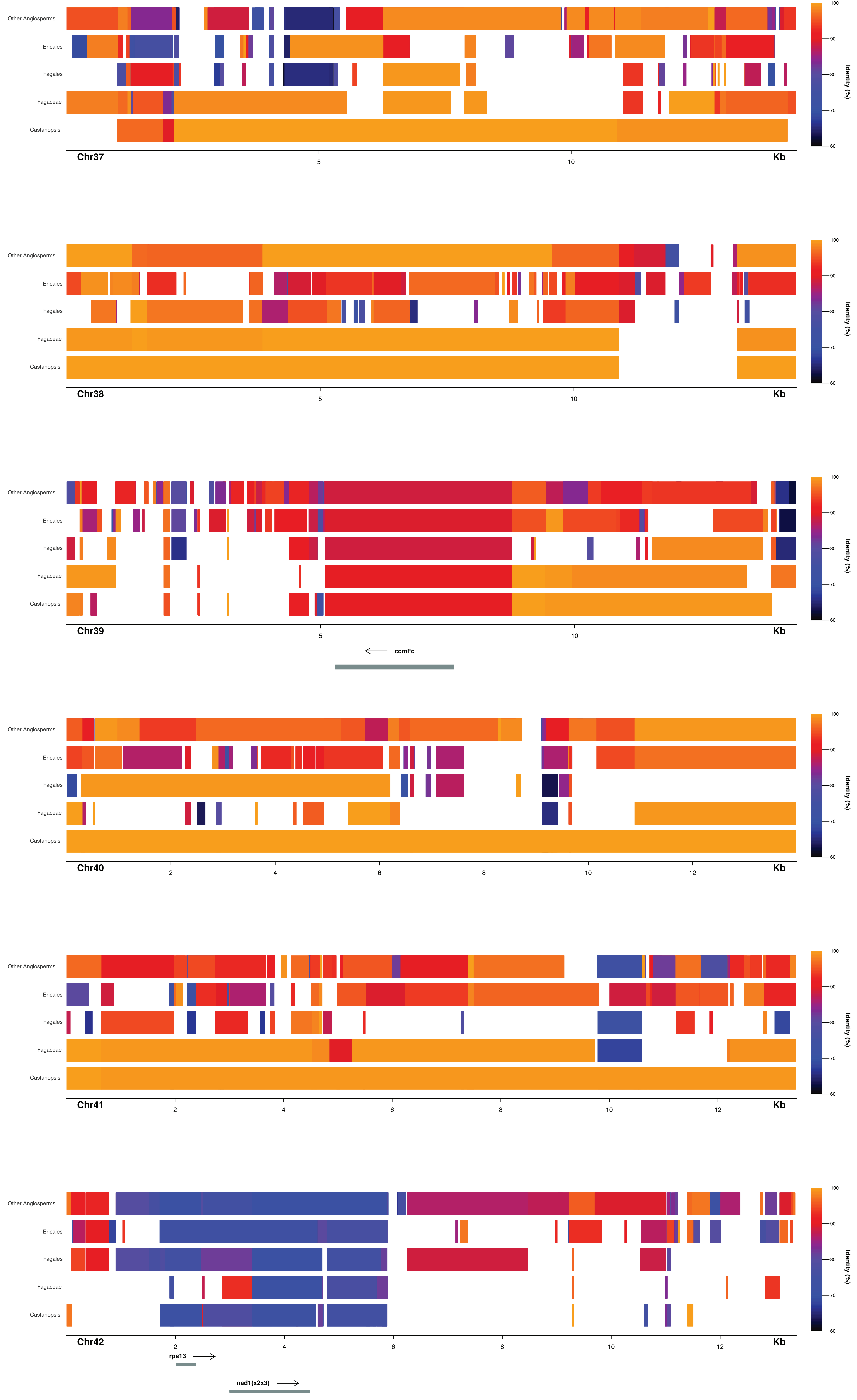

Other Angiosperms
Ericales
Fagales
Fagaceae
Castanopsis
Chr37
5
10
Kb
Identity (%)
100
90
80
70
60
Chr38
Chr39
ccmFc
Chr40
2
4
6
8
12
Chr41
Chr42
rps13
nad1(x2x3)

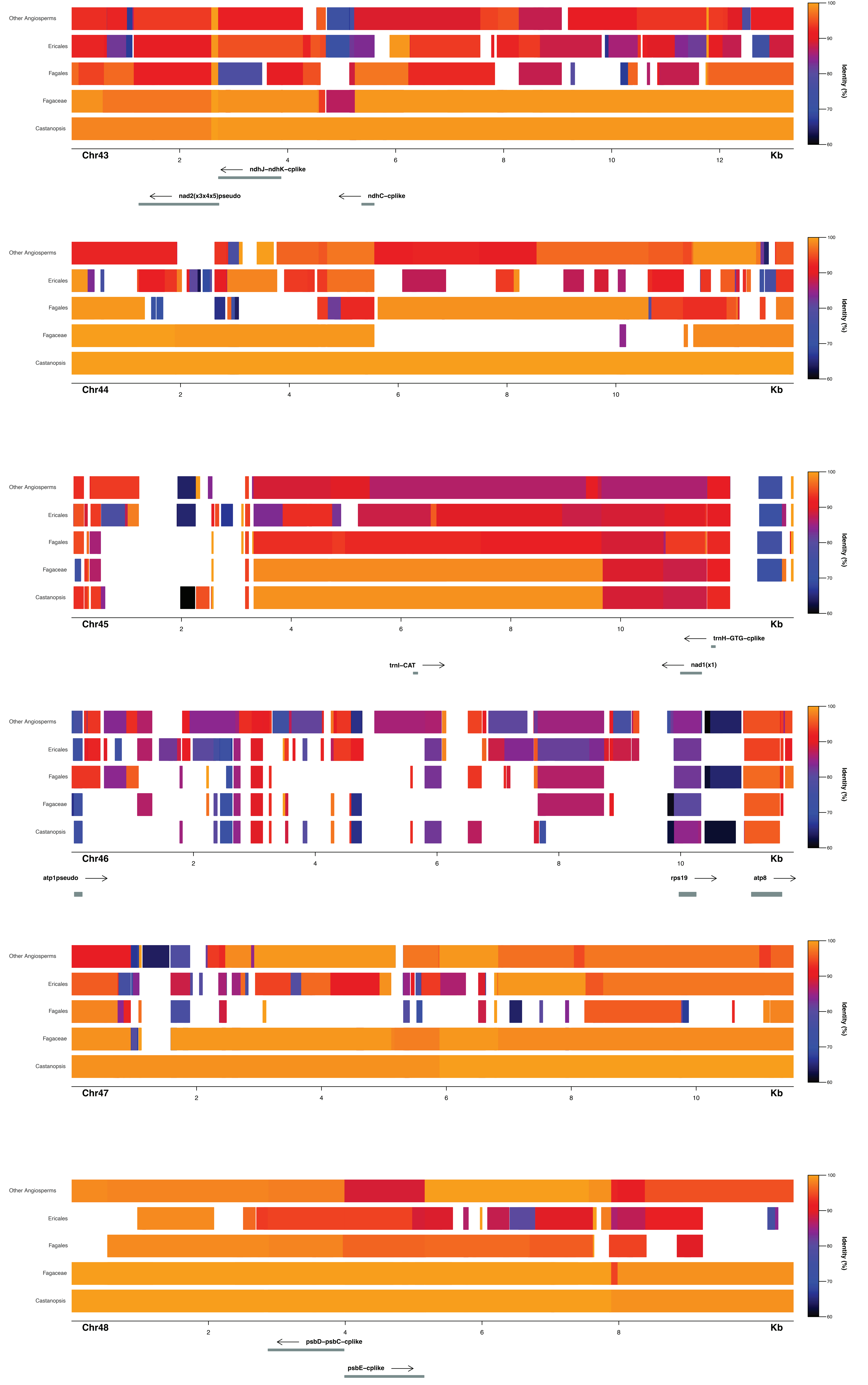
Other Angiosperms
Ericales
Fagales
Fagaceae
Castanopsis
Identity (%)
Chr43
Kb
ndhJ–ndhK–cplike
nad2(x3x4x5)pseudo
ndhC–cplike
Chr44
Chr45
trnH–GTG–cplike
trnI–CAT
nad1(x1)
Chr46
atp1pseudo
rps19
atp8
Chr47
Chr48
psbD–psbC–cplike
psbE–cplike

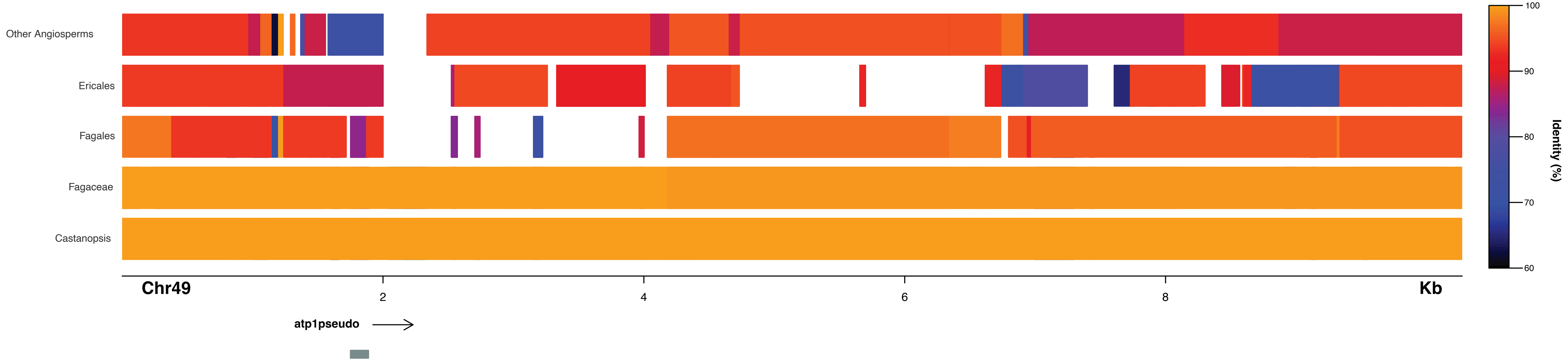
Other Angiosperms
Ericales
Fagales
Fagaceae
Castanopsis
Chr49
2
4
6
8
Kb
atp1pseudo
100
90
80
70
60
Identity (%)

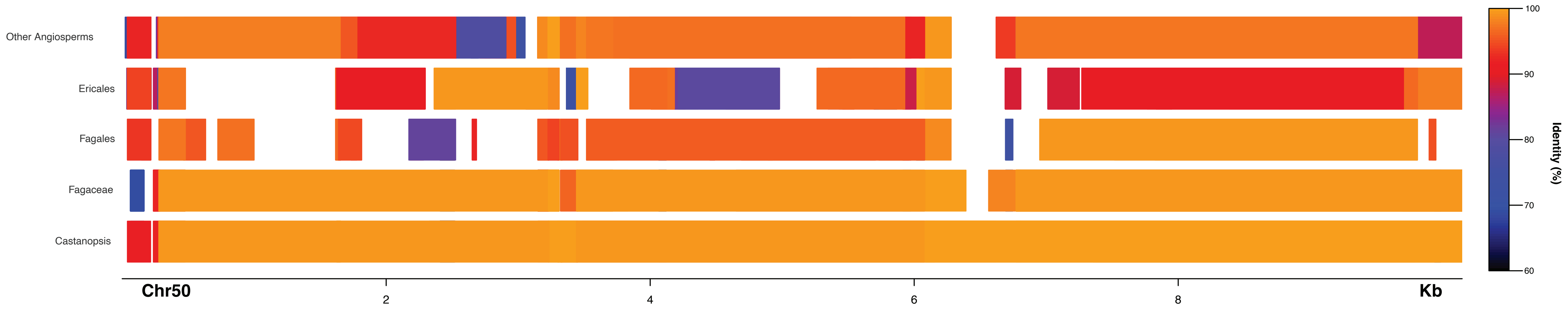
Other Angiosperms
Ericales
Fagales
Fagaceae
Castanopsis
Chr50
2
4
6
8
Kb
100
90
80
70
60
Identity (%)

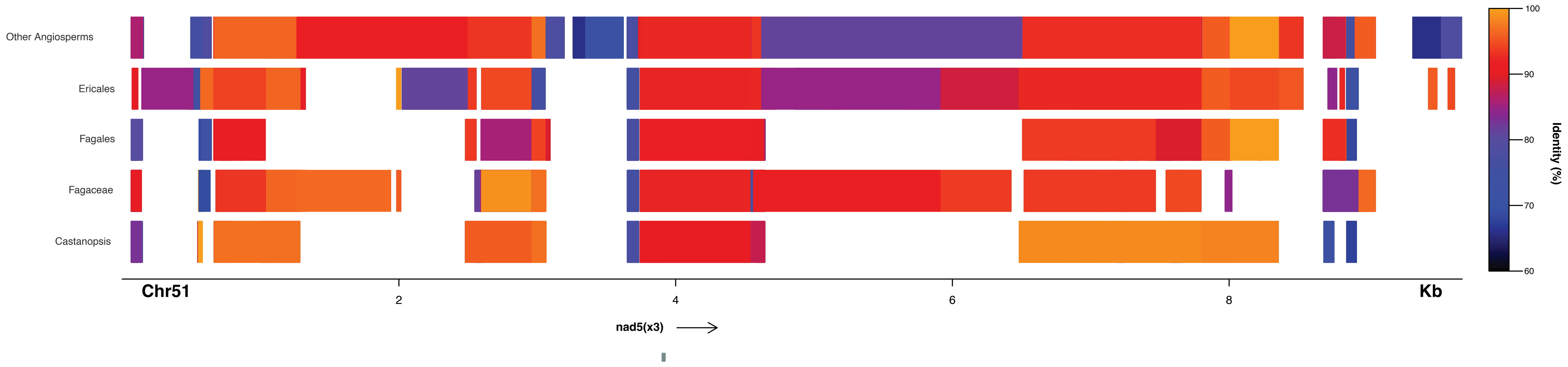
Other Angiosperms
Ericales
Fagales
Fagaceae
Castanopsis
Chr51
2
4
6
8
Kb
nad5(x3)
100
90
80
70
60
Identity (%)

**Figure S4**. Phylogenetic origin of the intergenic regions of the mtDNA of *Mitrastemon yamamotoi*. Circular chromosomes are shown linearized, with annotations in green. The best BLAST hits are shown, selected based on sequence identity and alignment length. Red lines represent BLAST hits using a mitochondrial database from the order Fagales and those regions are inferred to be foreign.Blue and yellow lines represent BLAST hits using a database of Ericales, and of all angiosperms excluding Fagales and Ericales, respectively. The origin of genes has been analyzed by phylogenetic analyses (FigureS2).

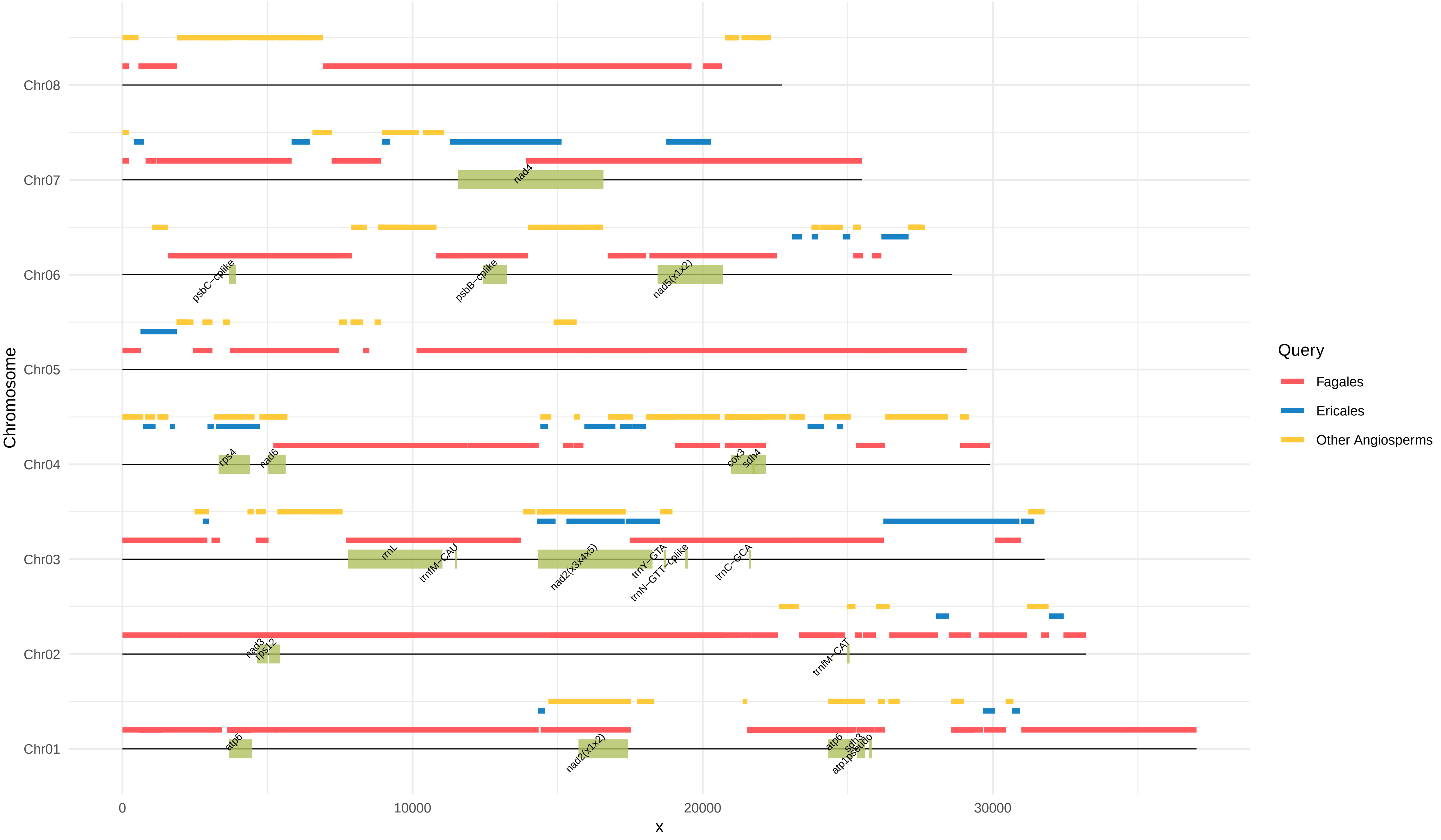

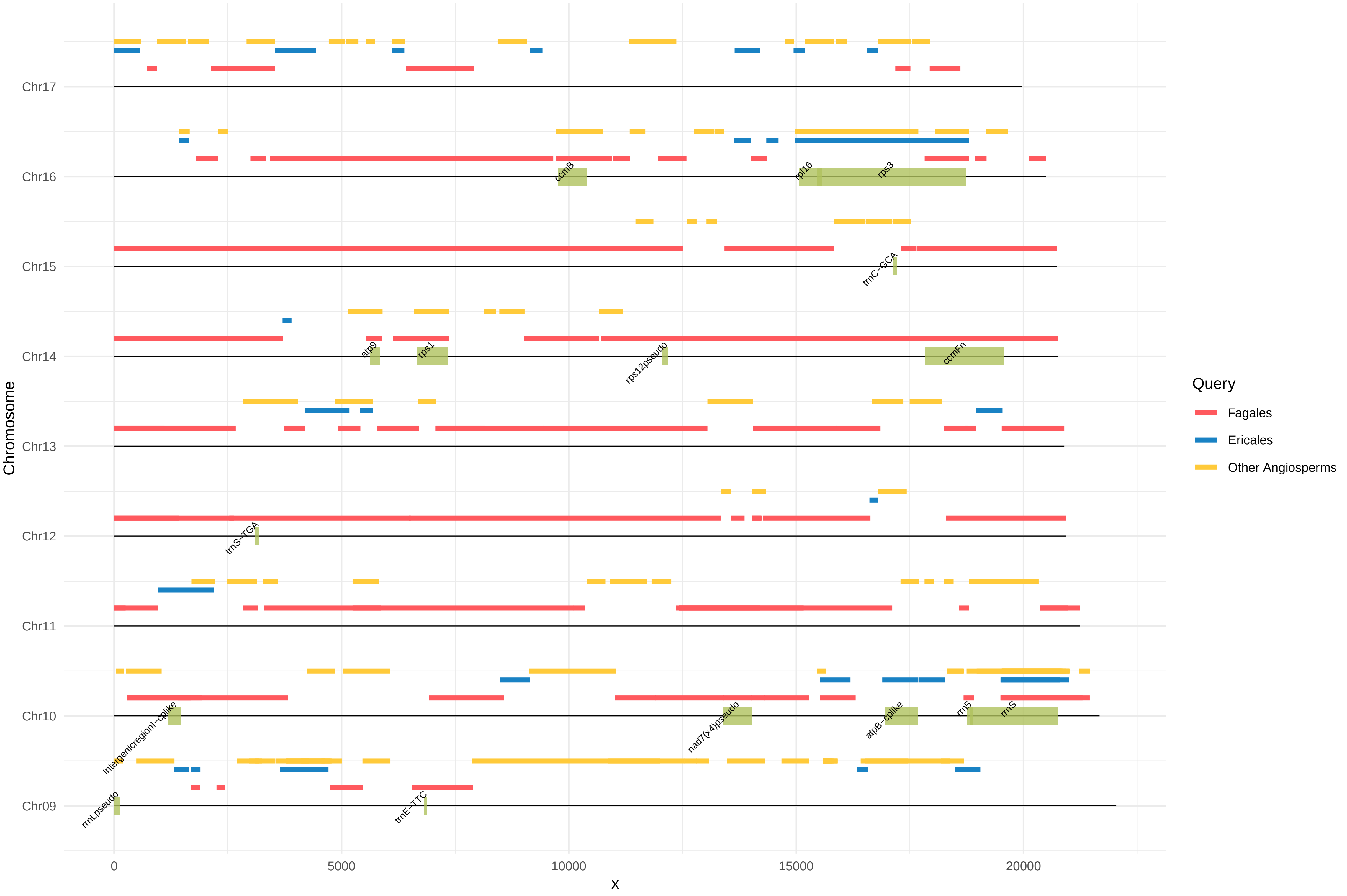
Chr17
Chr16
Chr15
Chr14
Chr13
Chr12
Chr11
Chr10
Chr09
Chromosome
x
0
5000
10000
15000
20000
ccmB
rpl16
rps3
trnC−GCA
atp9
rps1
rps12pseudo
ccmFn
trnS−TGA
Intergenicregionl−cplike
nad7(x4)pseudo
atpB−cplike
rrn5
rrnS
rrnLpseudo
trnE−TTC
Query
Fagales
Ericales
Other Angiosperms

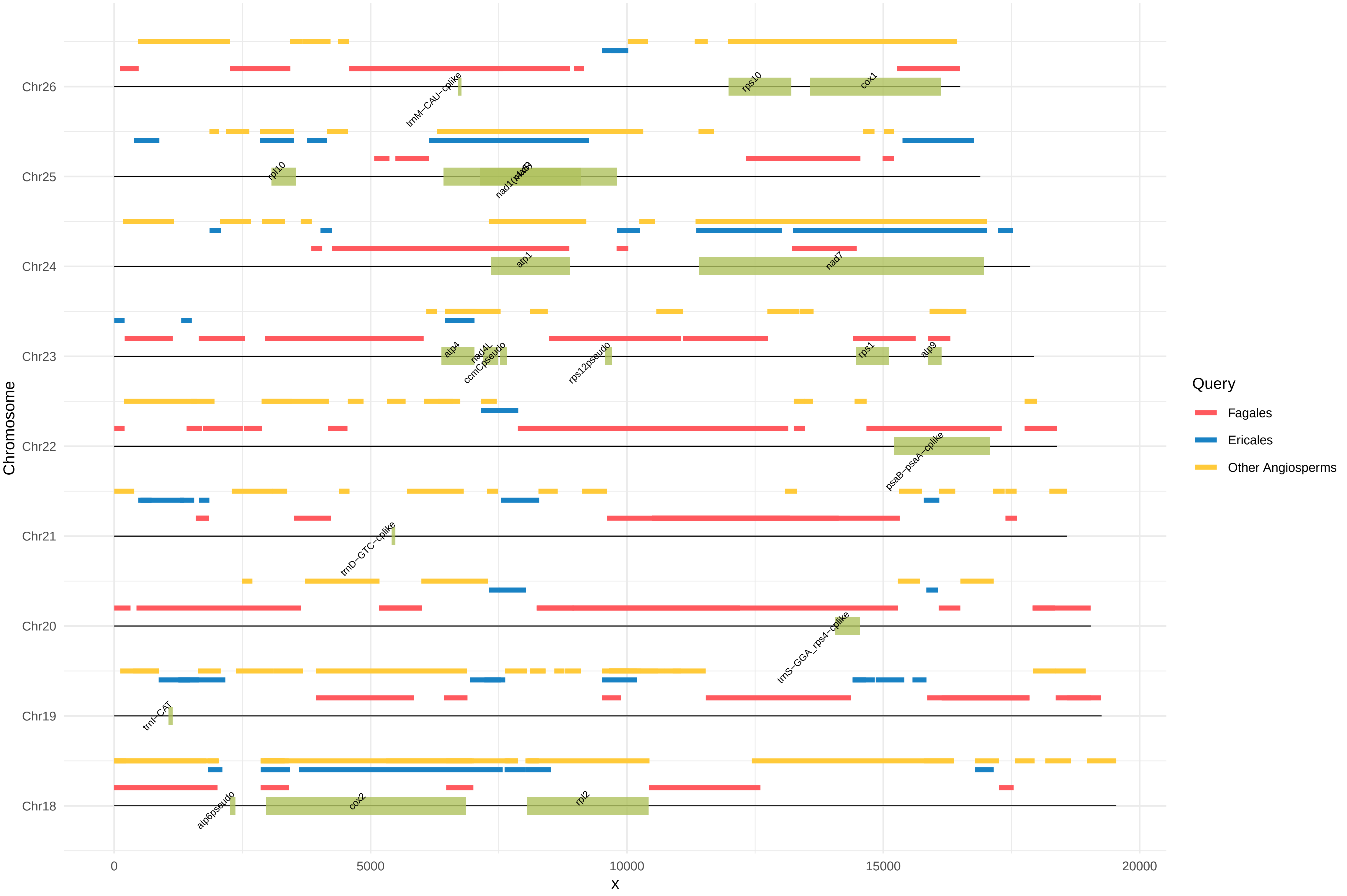

Chr26
Chr25
Chr24
Chr23
Chr22
Chr21
Chr20
Chr19
Chr18
Chromosome
0
5000
10000
15000
20000
x
trnM−CAU−cplike
rps10
cox1
rpl10
nad1(nad5)
atp1
nad7
atp4
nad4L
ccmCpseudo
rps12pseudo
rps1
atp9
psaB−psaA−cplike
trnD−GTC−cplike
trnS−GGA_rps4−cplike
trnI−CAT
atp6pseudo
cox2
rpl2
Query
Fagales
Ericales
Other Angiosperms

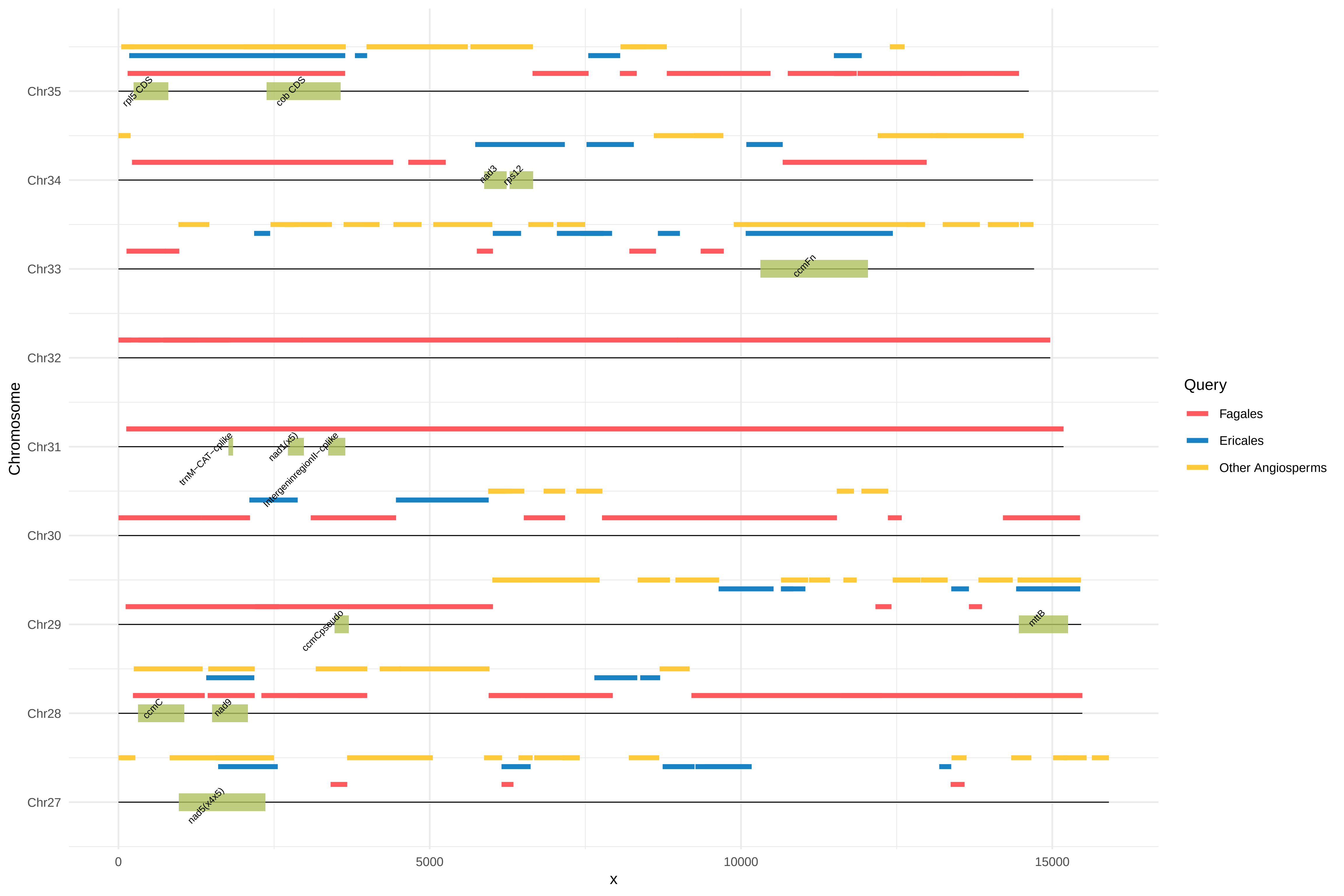

Chr35
Chr34
Chr33
Chr32
Chr31
Chr30
Chr29
Chr28
Chr27
Chromosome
0
5000
10000
15000
x
rpl5 CDS
cob CDS
nad3
rps12
ccmFn
trnM−CAT−cplike
nad1(x5)
intergeninregionII−cplike
ccmCpseudo
mttB
ccmC
nad9
nad5(x4x5)
Query
Fagales
Ericales
Other Angiosperms

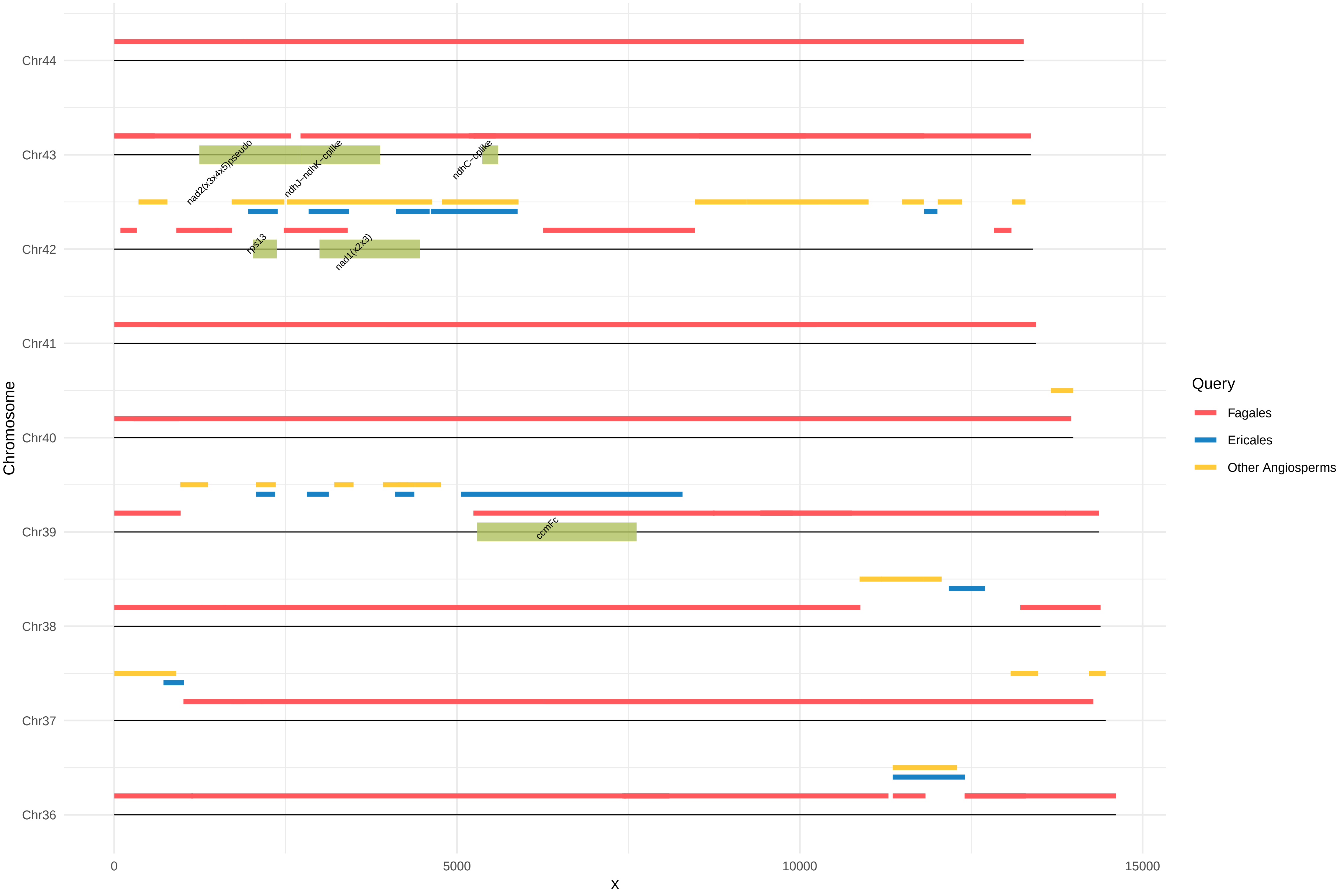

Chr44
Chr43
Chr42
Chr41
Chr40
Chr39
Chr38
Chr37
Chr36
Chromosome
0
5000
10000
15000
x
nad2(x3x4x5)pseudo
ndhJ−ndhK−cplike
ndhC−cplike
rps13
nad1(x2x3)
ccmFc
Query
Fagales
Ericales
Other Angiosperms

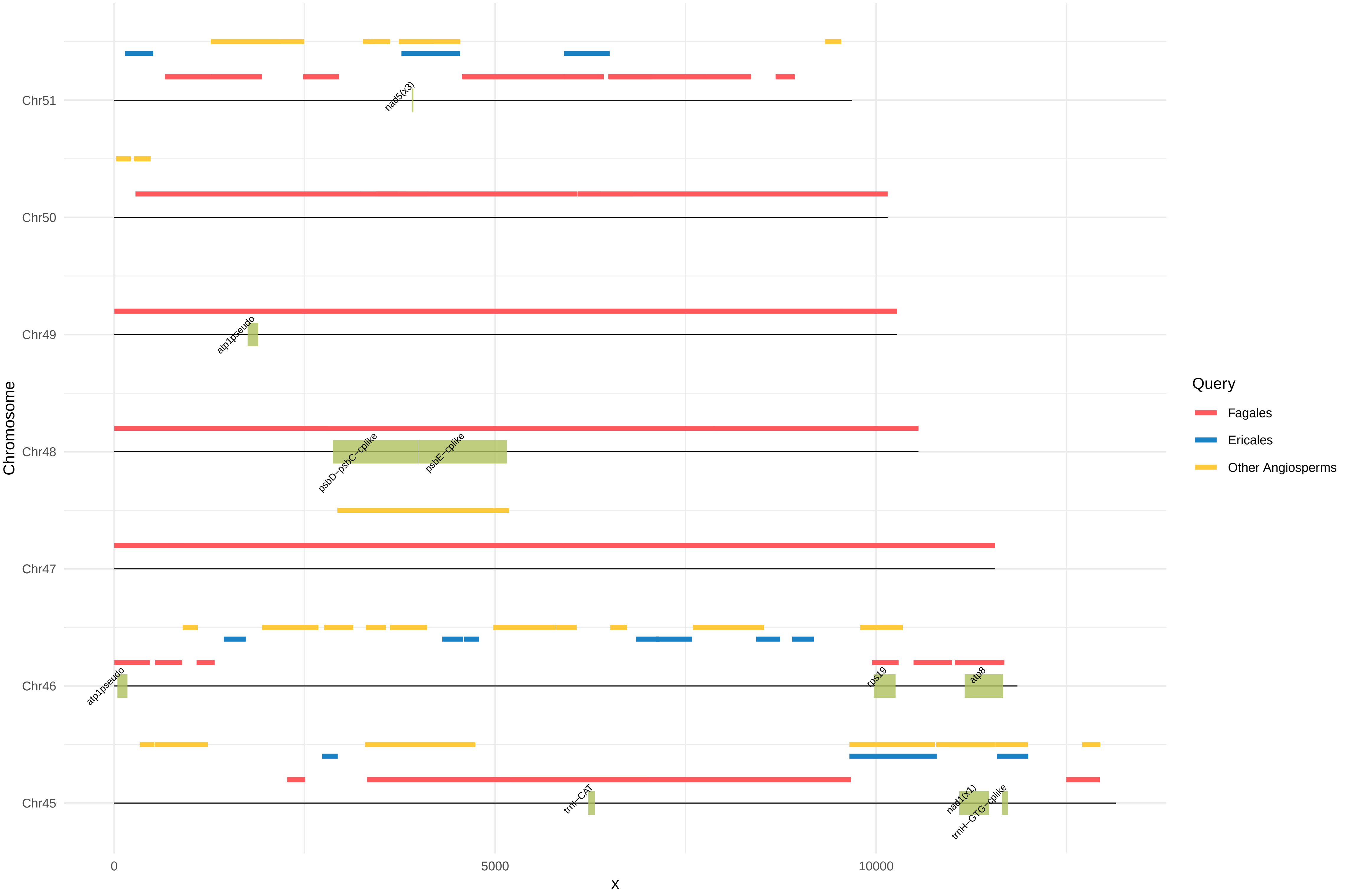

Chr51
Chr50
Chr49
Chr48
Chr47
Chr46
Chr45
Chromosome
0
5000
10000
x
nad5(x3)
atp1pseudo
psbD−psbC−cplike
psbE−cplike
atp1pseudo
rps19
atp8
trnI−CAT
nad1(x1)
trnH−GTG−cplike
Query
Fagales
Ericales
Other Angiosperms

**Figure S5.** Approximately unbiased (AU) tests to evaluate alternative topologies based on different constrained trees. Significant *p*-values are shown in boldface. The constrained topologies were prepared individually for each foreign gene of *Mitrastemon*.

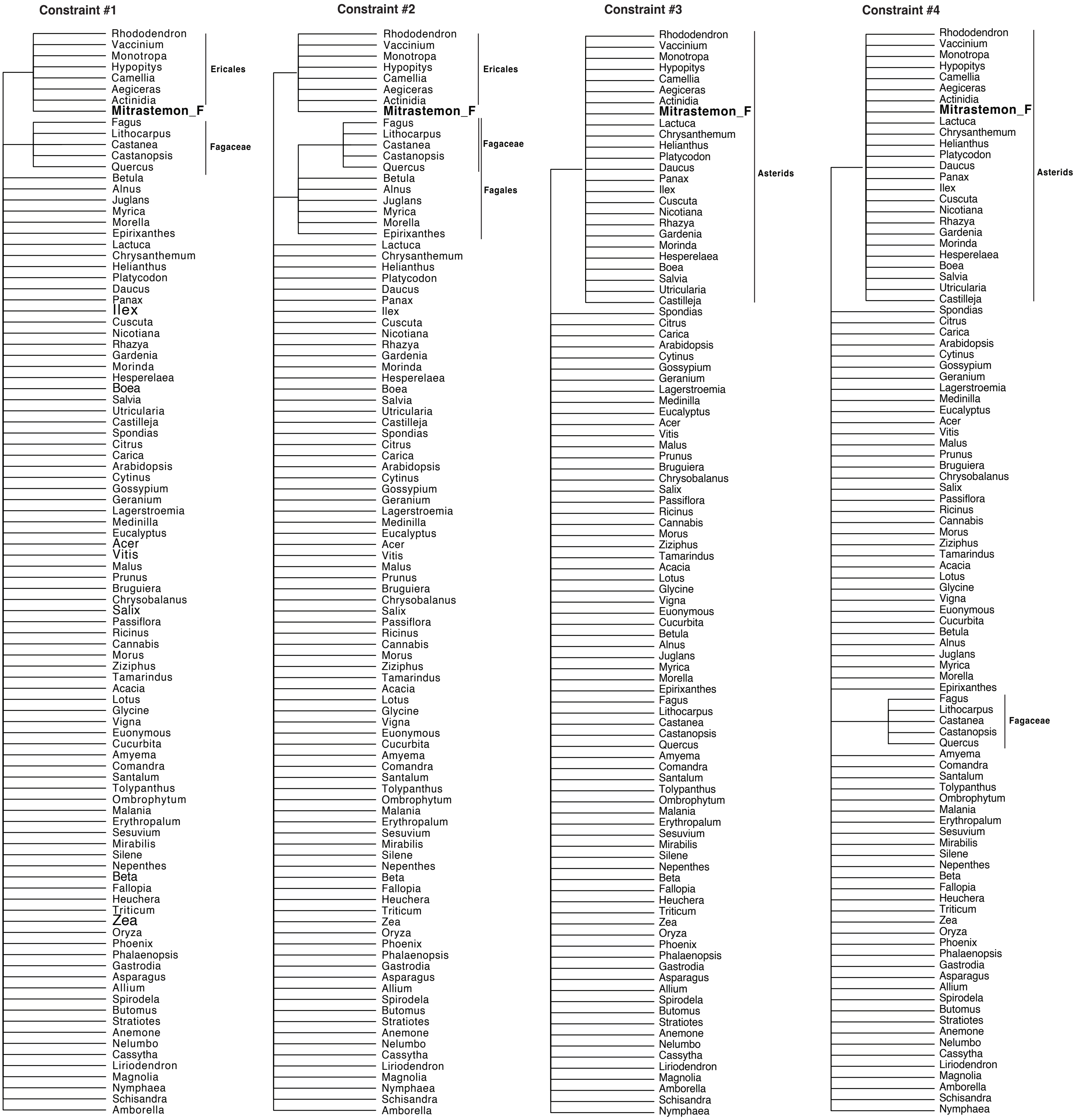


***Ericales** include *Rhododendron, Vaccinium, Monotropa, Hypopitys, Camellia, Aegiceras, Actinidia.*

| Foreign genes | Unconstrained | Constraint #1 | Constraint #2 | Constraint #3 | Constraint #4 |
|---|---|---|---|---|---|
| *atp1* Chr24 | 0.849 | 0.225 | 0.179 | **3e-78** | **1e-46** |
| *ccmFN_1* Chr14 | 1.000 | **2e-05** | **1e-07** | **5e-06** | **1e-05** |
| *rps12_1* Chr02 | 0.857 | 0.121 | **0.014** | **0.005** | **0.002** |

**Figure S6.** Maximum Likelihood phylogenetic analyses of chloroplast-derived sequences (MTPTs) in the mtDNA of *Mitrastemon yamamotoi*. ML bootstrap support values >50% are shown above each branch. Scale bars correspond to substitutions per site.

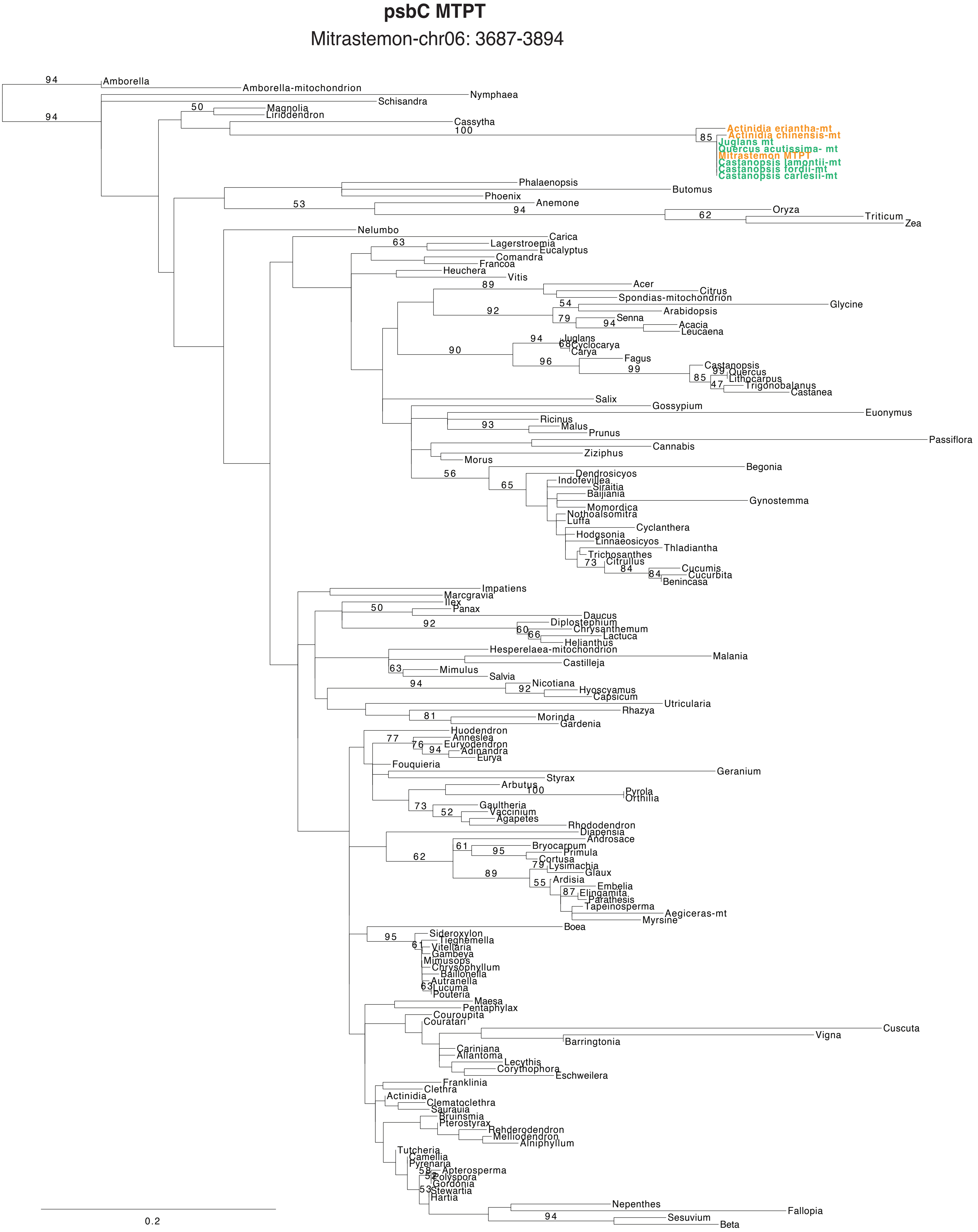

### psbB MTPT

Mitrastemon-chr06: 12448-13255

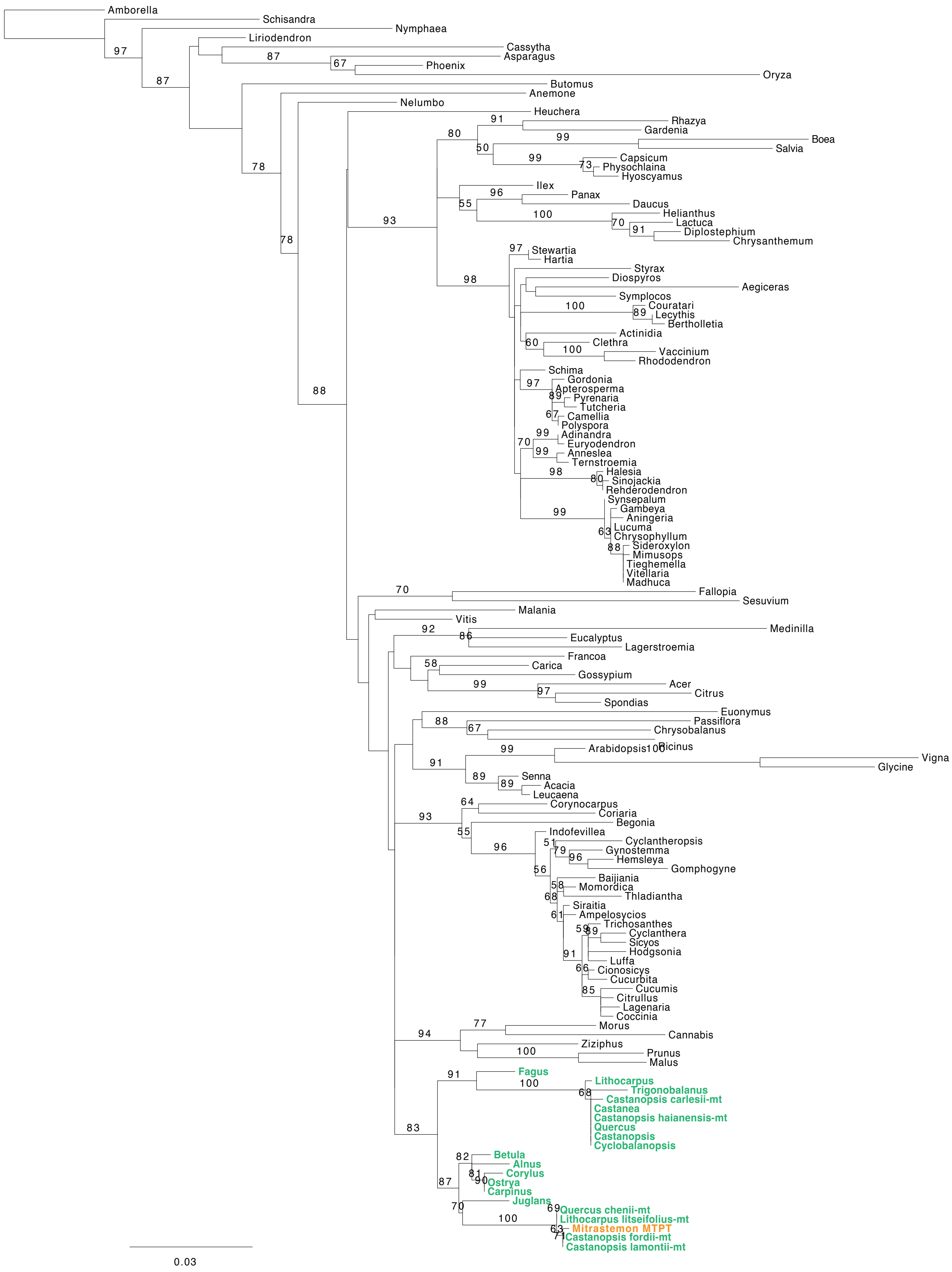

### Intergenic Region I MTPT

Mitrastemon-chr10: 1188-1476

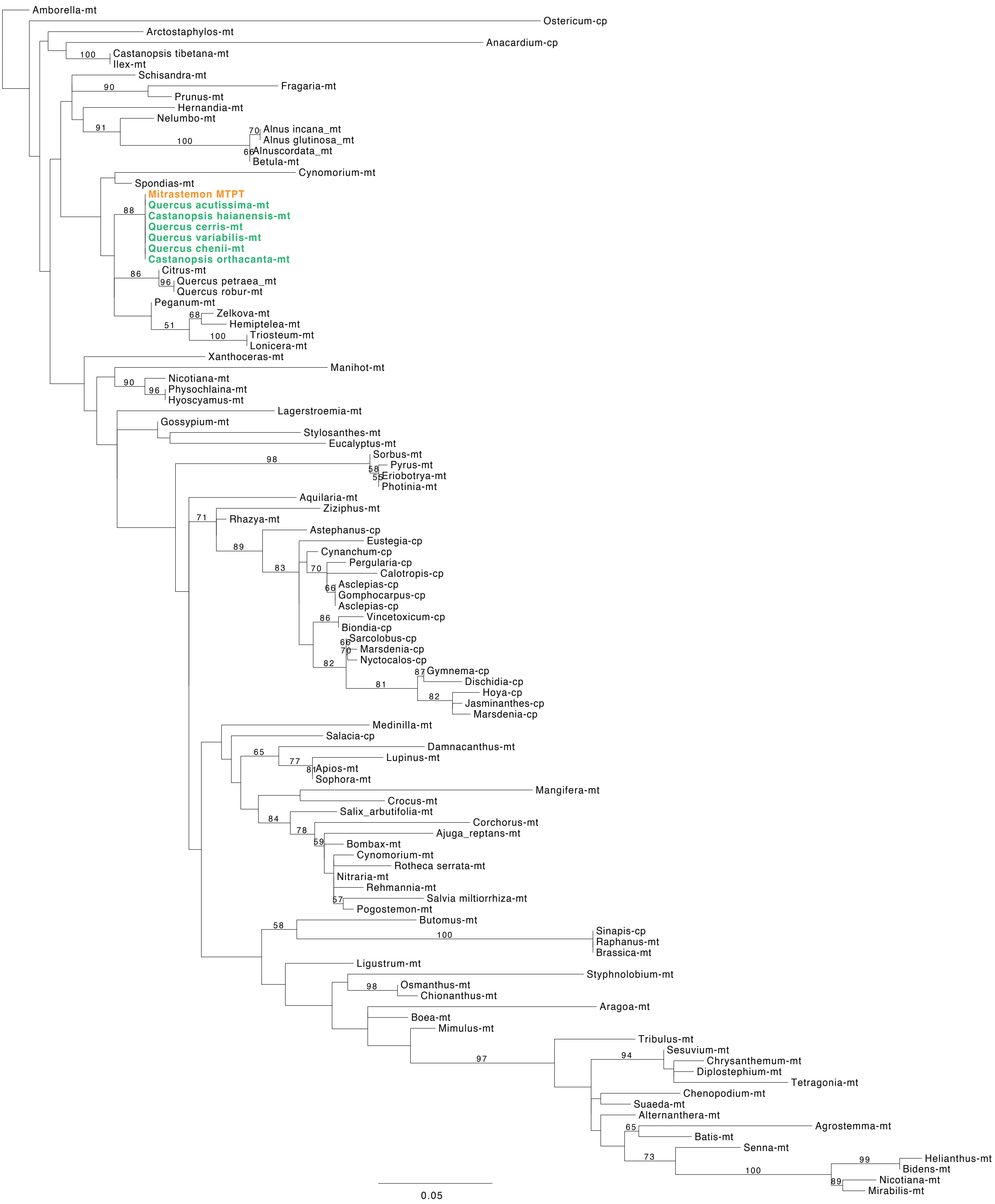

### atpB MTPT

Mitrastemon-chr10: 17672-16947

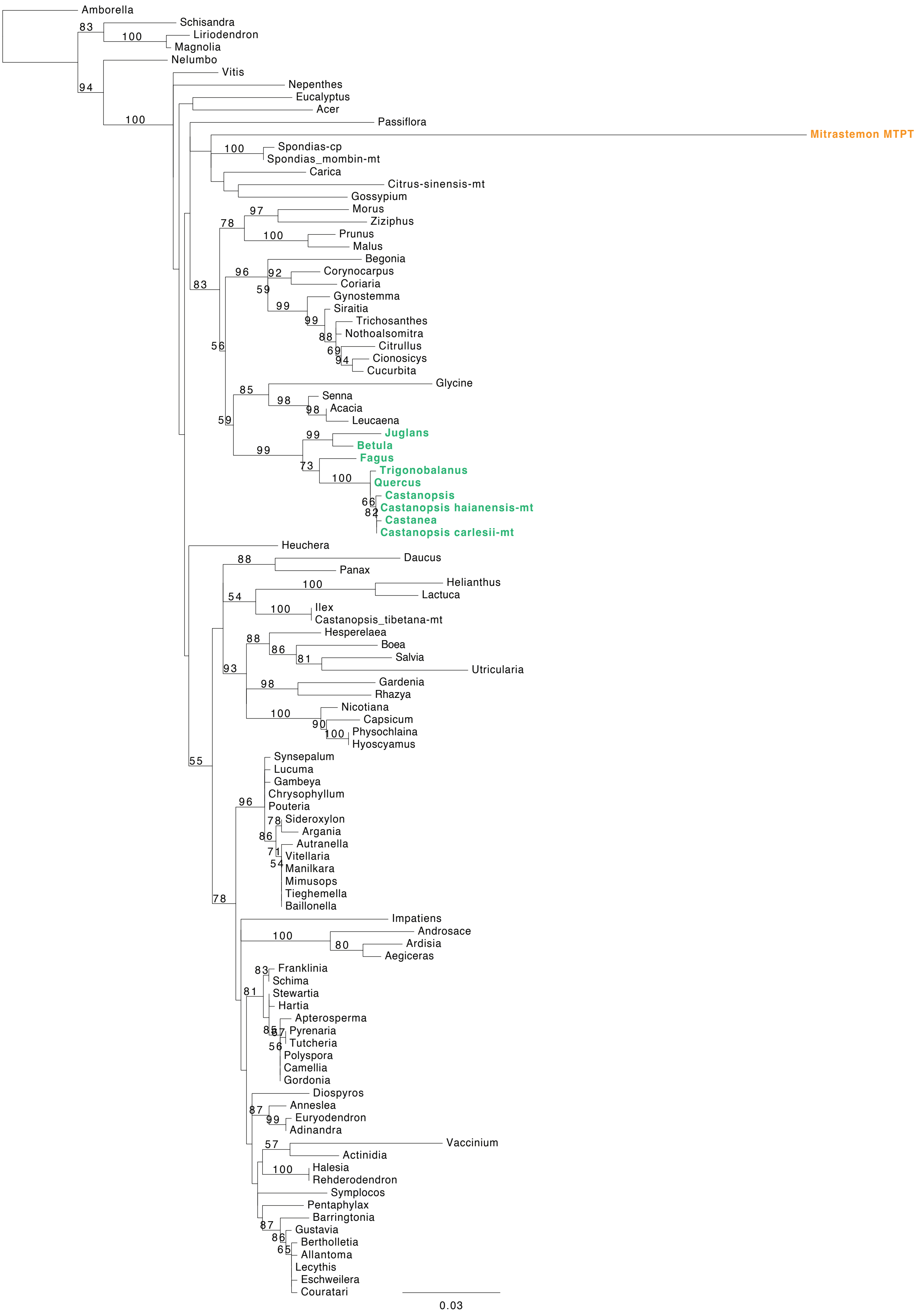

#### trnS-GGA - rps4 MTPT

Mitrastemon-chr20: 14057-14549

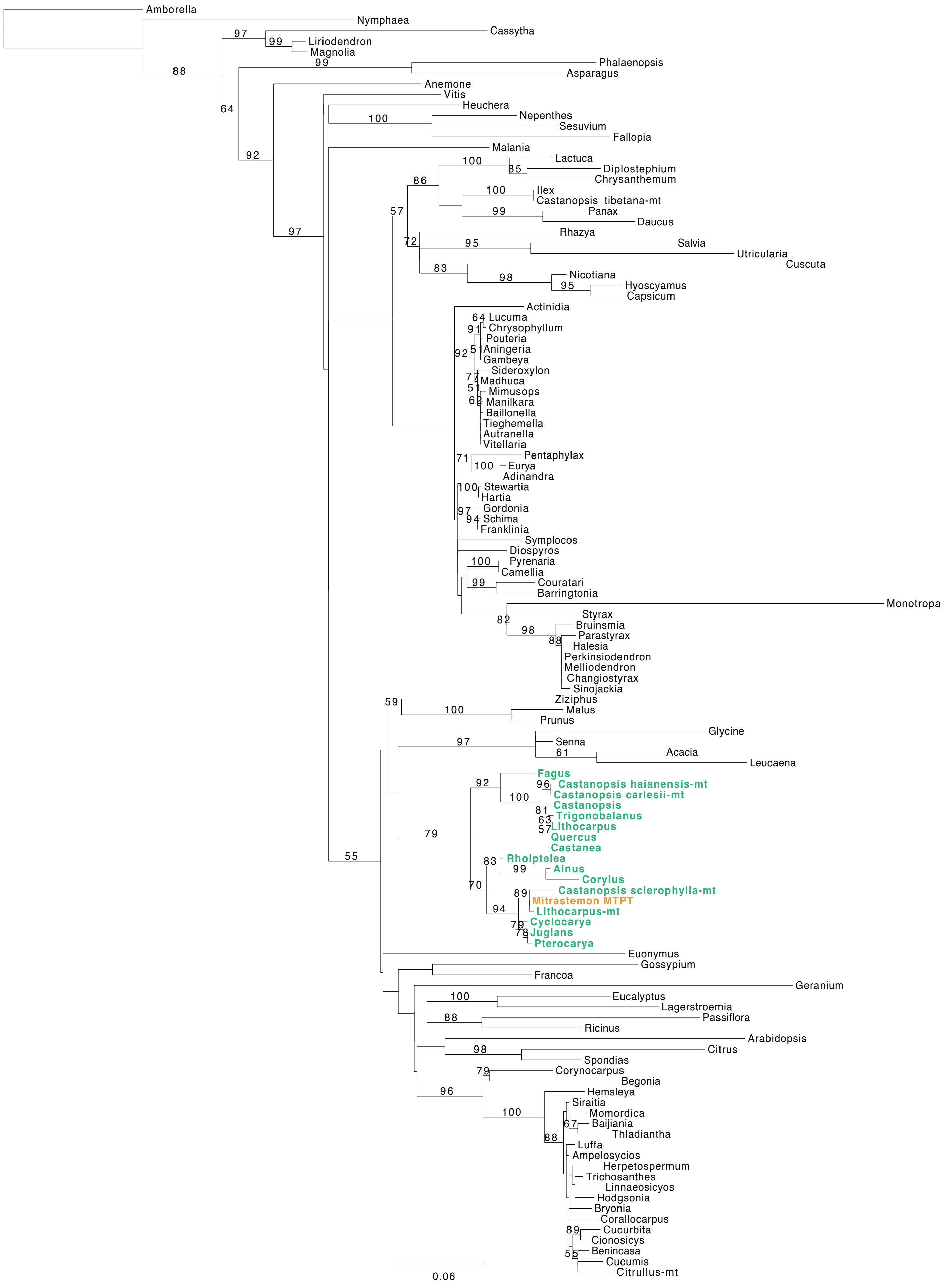

### psaB-psaA MTPT

Mitrastemon-chr22: 15206-17082

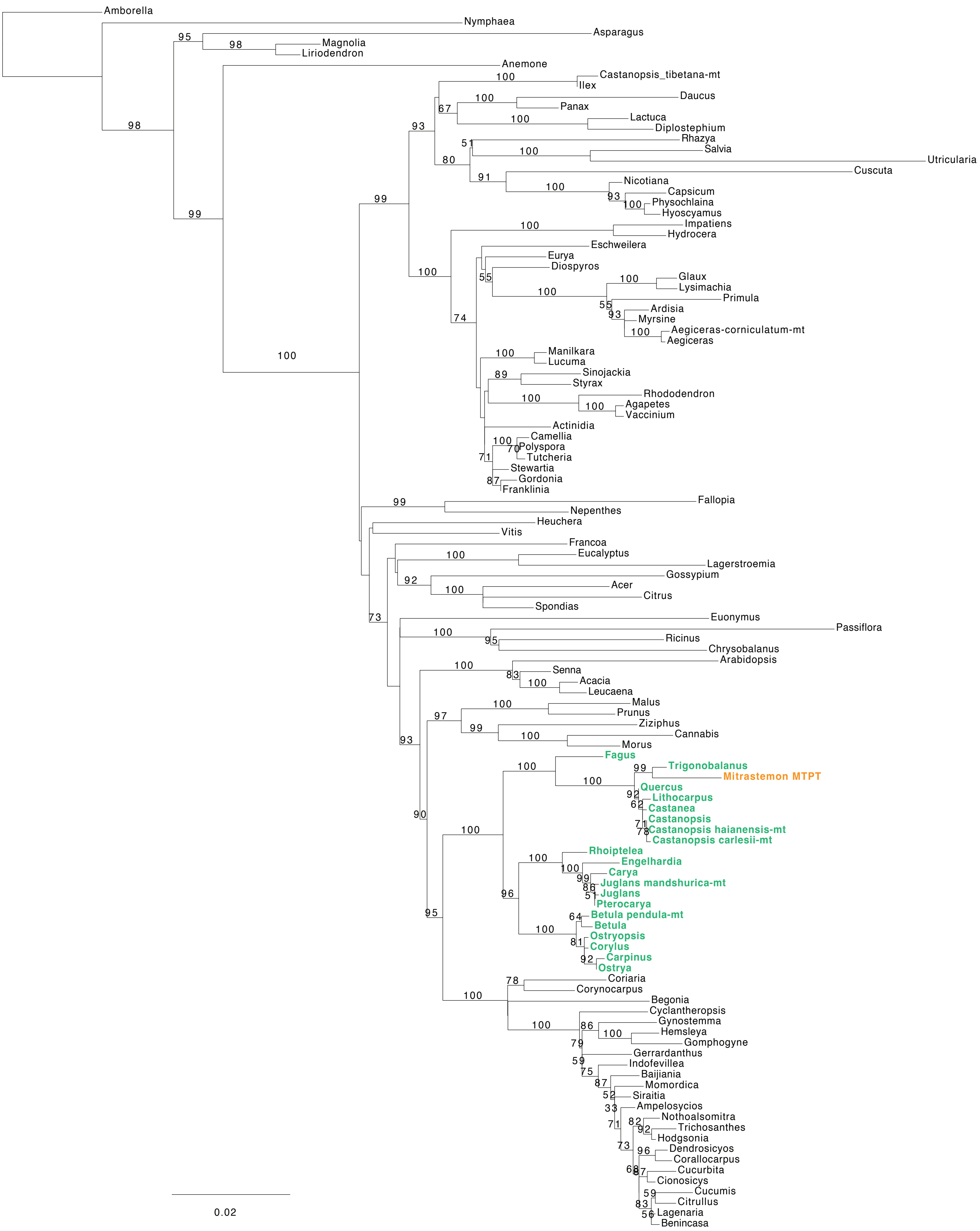

**Intergenic region II MTPT**

Mitrastemon-chr31: 3621-3349

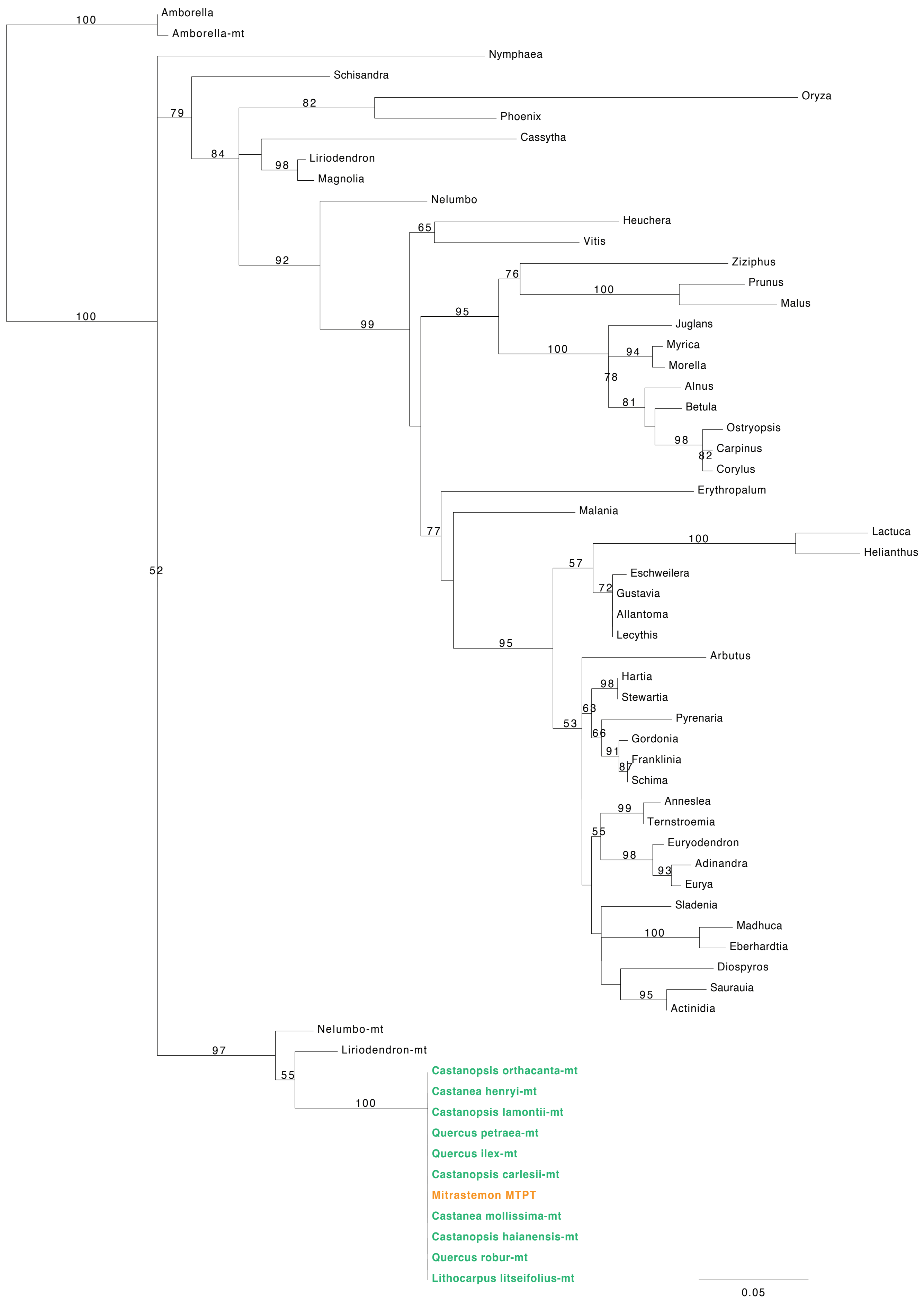

#### ndhJ-ndhK MTPT

Mitrastemon-chr43: 3877-2719

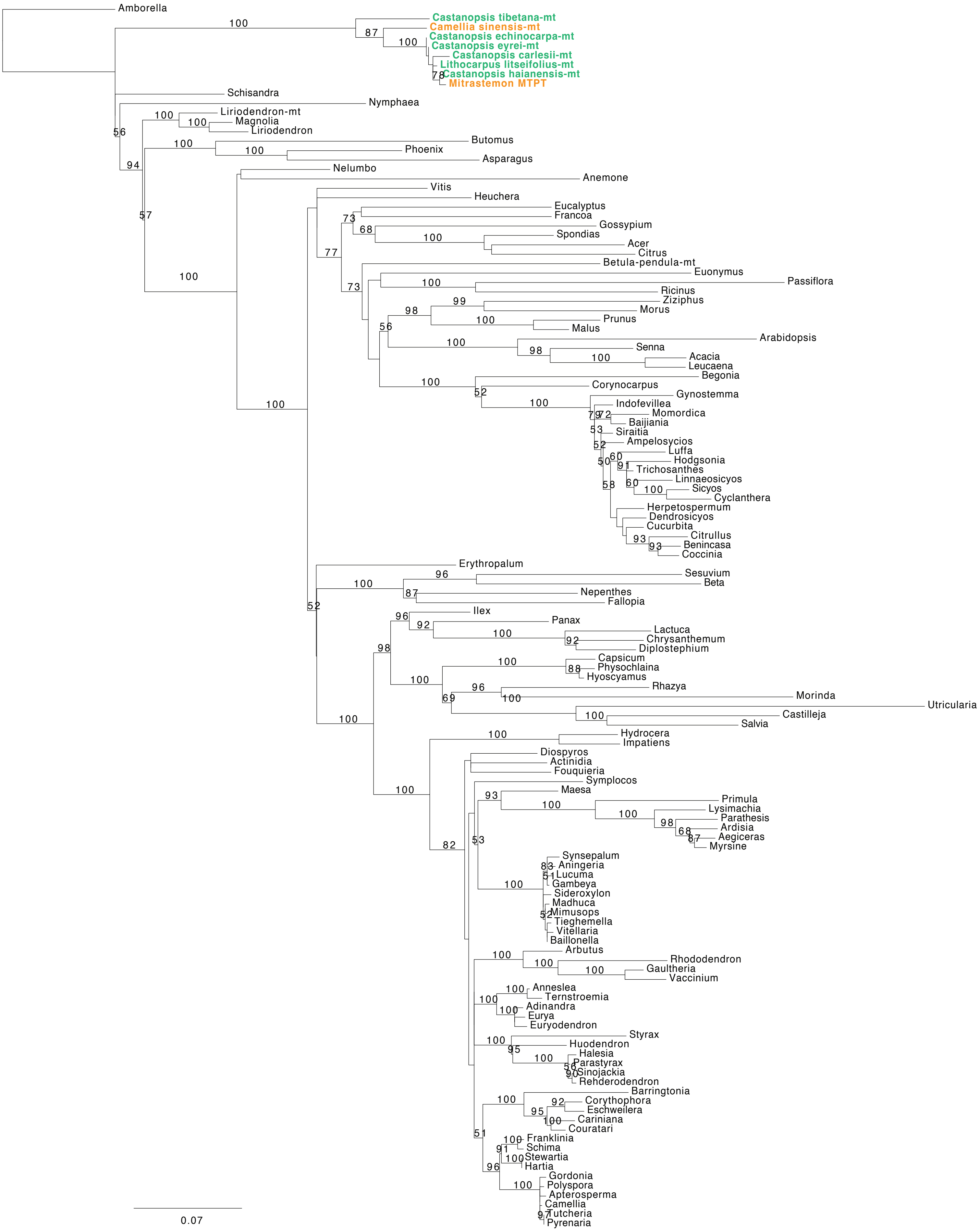

#### ndhC MTPT

Mitrastemon-chr43: 5603-5372

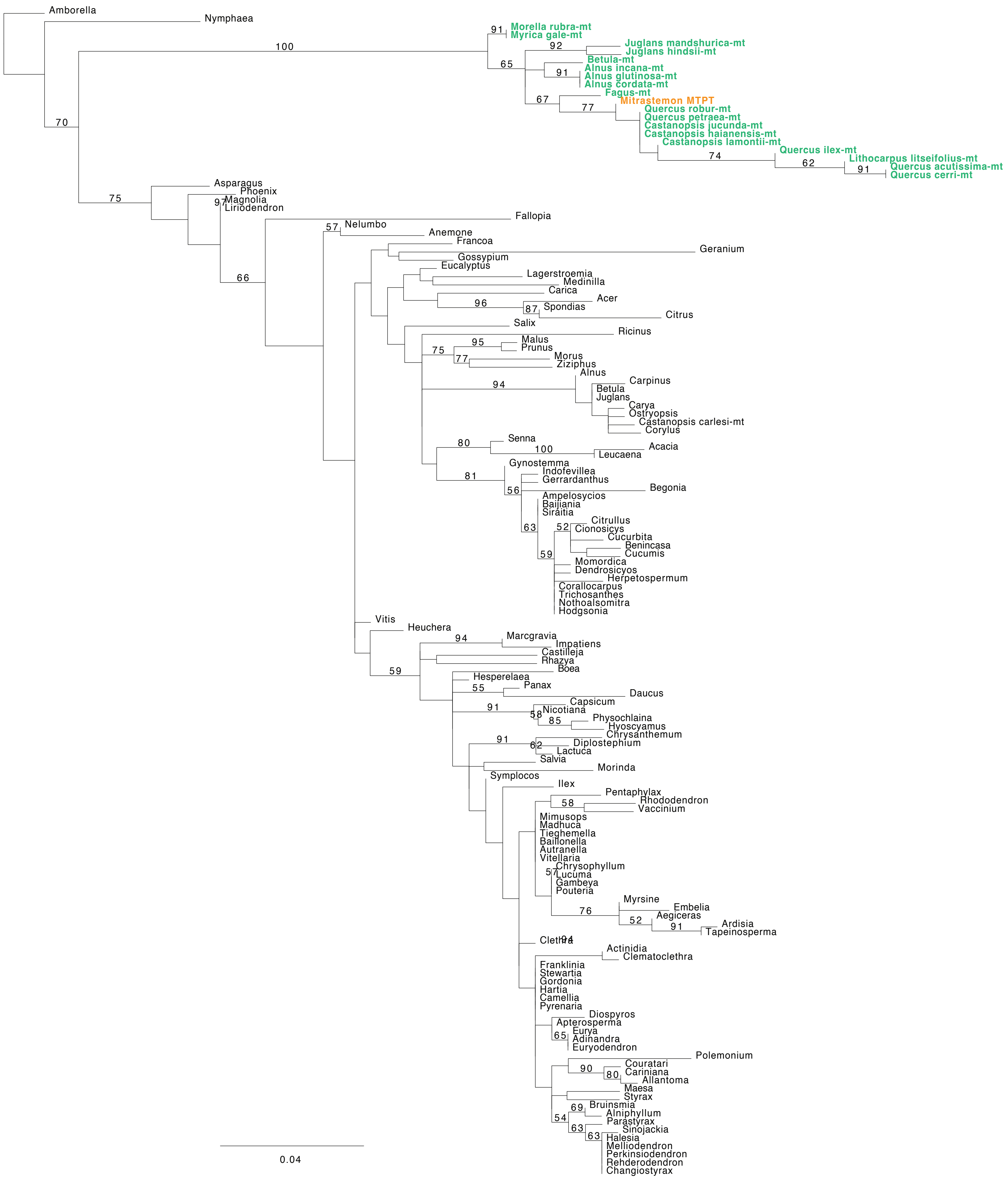

### psbD-psbC MTPT

Mitrastemon-chr48: 3984-2872

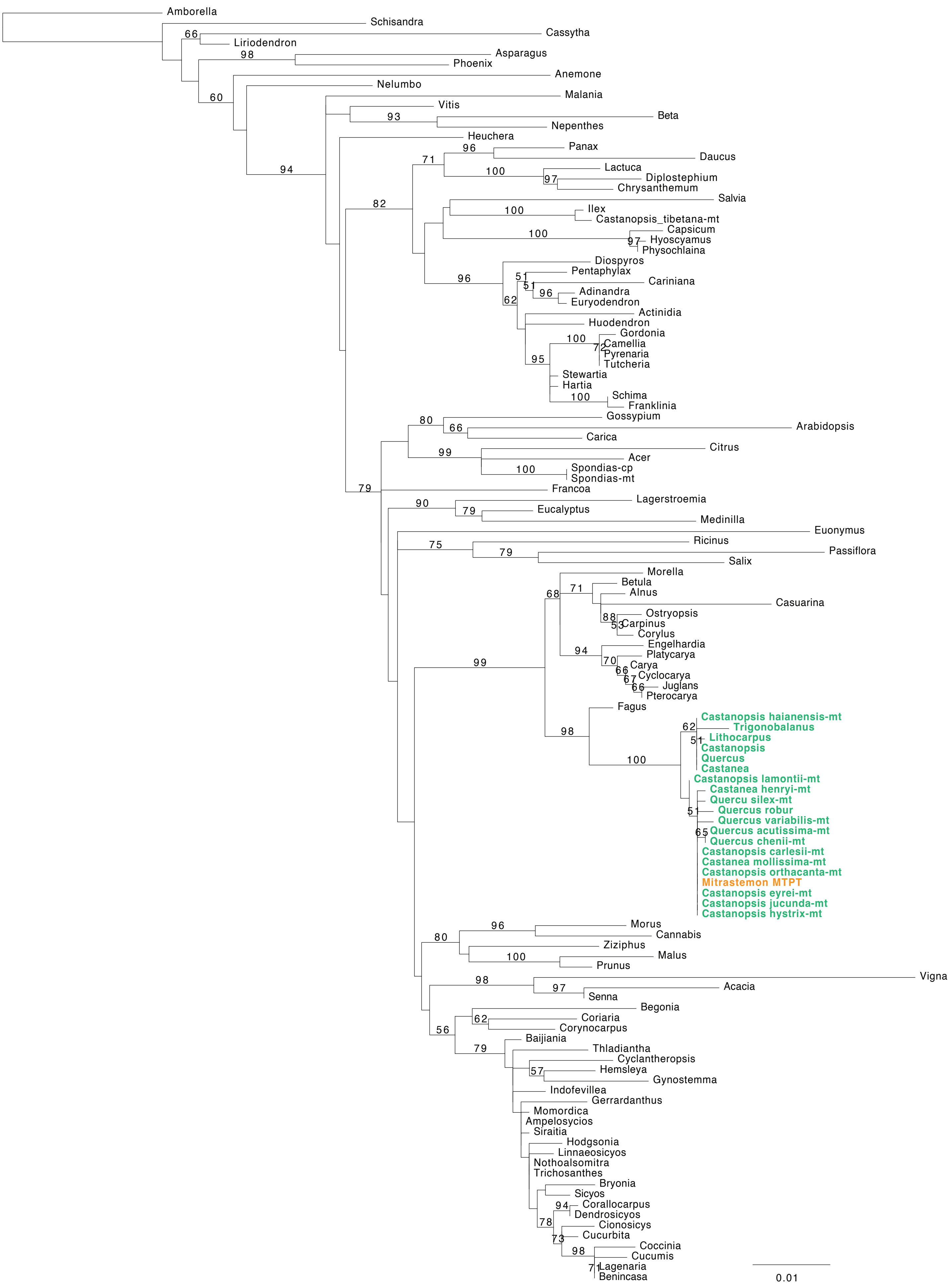

#### psbE MTPT

Mitrastemon-chr48: 3992-5153

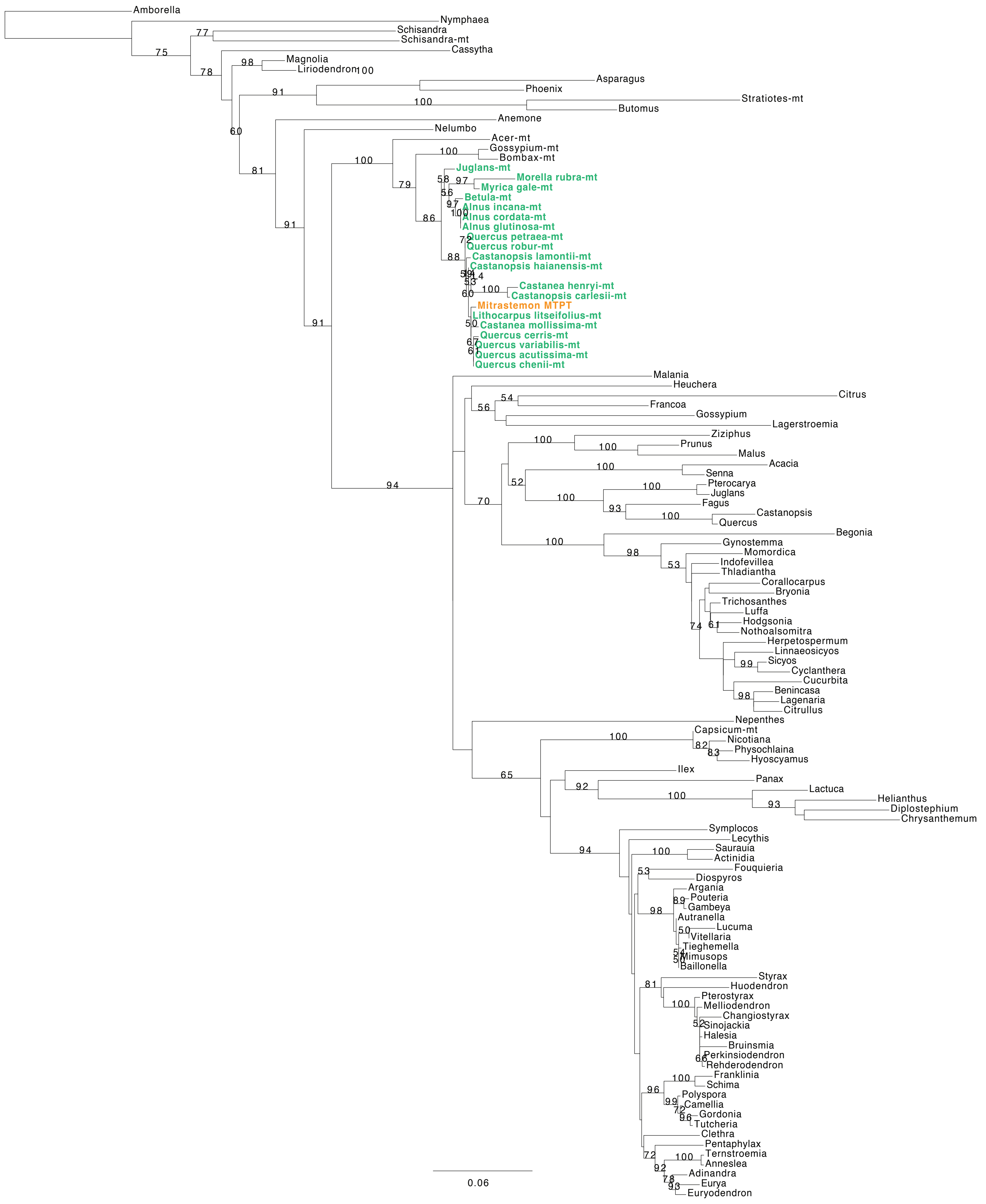

**Figure S7.** Maximum Likelihood phylogenetic analyses of duplicated DNA-RRR genes. ML bootstrap support values are shown above each branch. Scale bars correspond to substitutions per site.

#### A. ODB

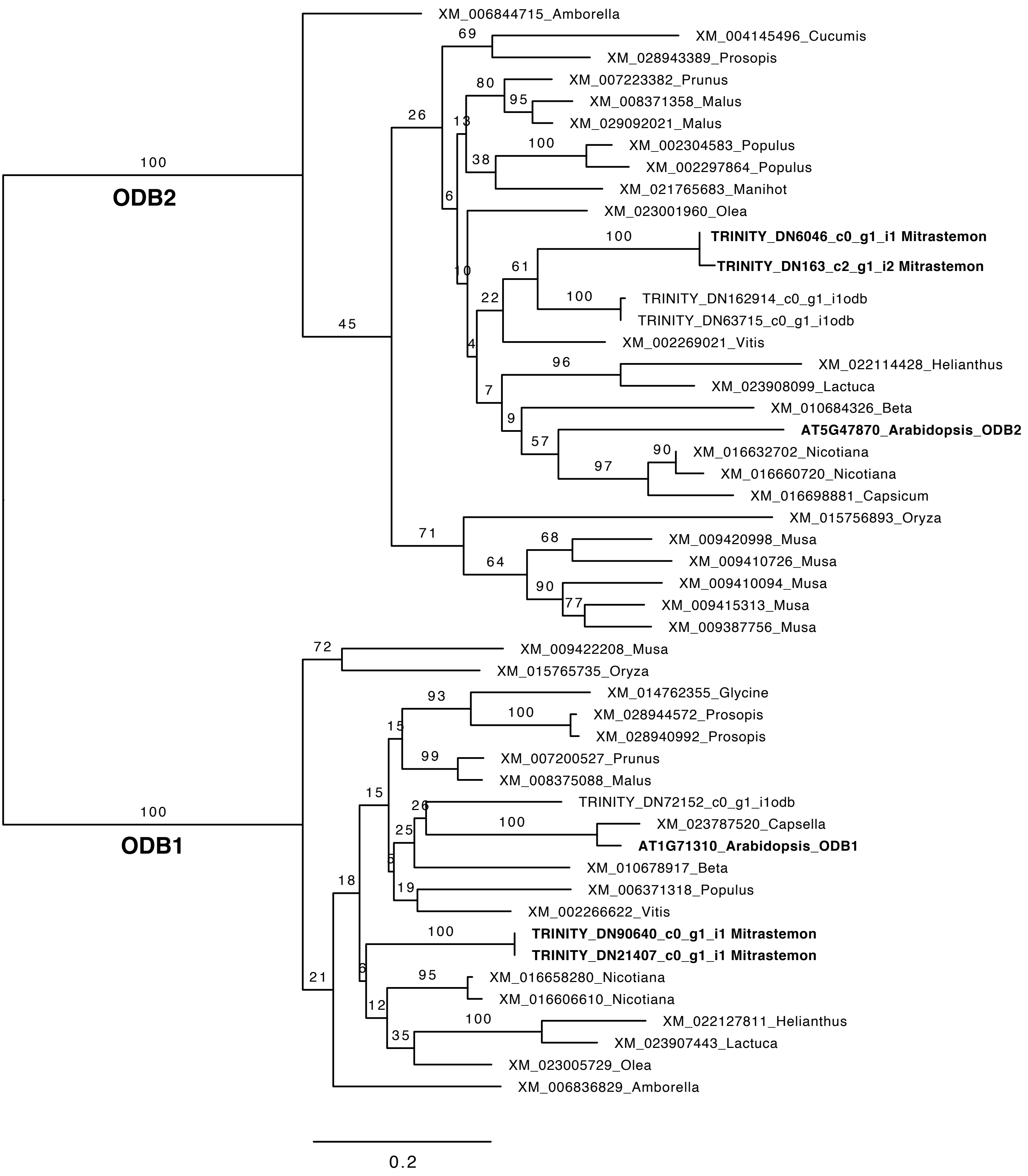

#### B. OSB

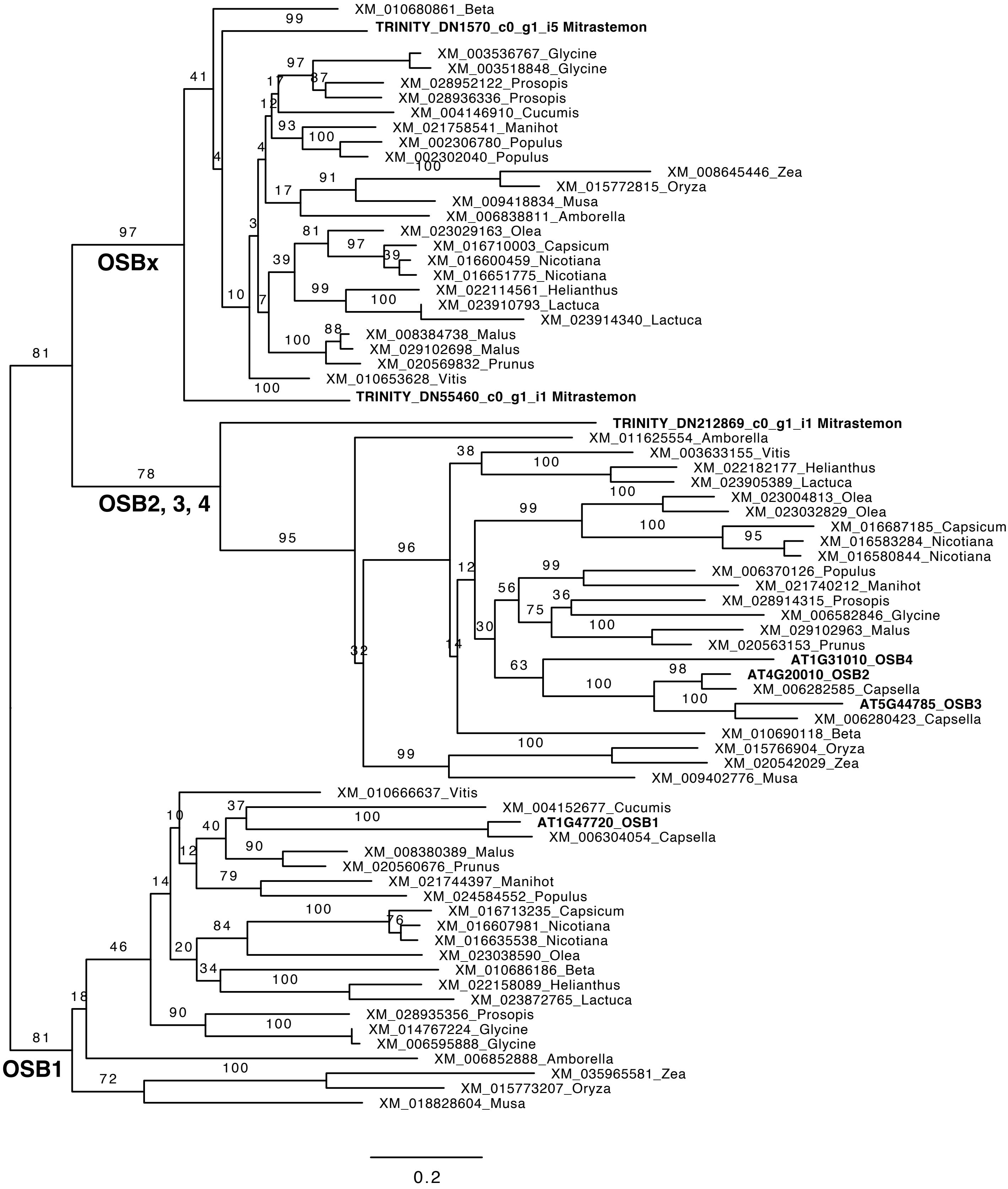

XM_010680861_Beta
TRINITY_DN1570_c0_g1_i5 Mitrastemon
XM_003536767_Glycine
XM_003518848_Glycine
XM_028952122_Prosopis
XM_028936336_Prosopis
XM_004146910_Cucumis
XM_021758541_Manihot
XM_002306780_Populus
XM_002302040_Populus
XM_008645446_Zea
XM_015772815_Oryza
XM_009418834_Musa
XM_006838811_Amborella
XM_023029163_Olea
XM_016710003_Capsicum
XM_016600459_Nicotiana
XM_016651775_Nicotiana
XM_022114561_Helianthus
XM_023910793_Lactuca
XM_023914340_Lactuca
XM_008384738_Malus
XM_029102698_Malus
XM_020569832_Prunus
XM_010653628_Vitis
TRINITY_DN55460_c0_g1_i1 Mitrastemon
OSBx
TRINITY_DN212869_c0_g1_i1 Mitrastemon
XM_011625554_Amborella
XM_003633155_Vitis
XM_022182177_Helianthus
XM_023905389_Lactuca
XM_023004813_Olea
XM_023032829_Olea
XM_016687185_Capsicum
XM_016583284_Nicotiana
XM_016580844_Nicotiana
XM_006370126_Populus
XM_021740212_Manihot
XM_028914315_Prosopis
XM_006582846_Glycine
XM_029102963_Malus
XM_020563153_Prunus
AT1G31010_OSB4
AT4G20010_OSB2
XM_006282585_Capsella
AT5G44785_OSB3
XM_006280423_Capsella
XM_010690118_Beta
XM_015766904_Oryza
XM_020542029_Zea
XM_009402776_Musa
OSB2, 3, 4
XM_010666637_Vitis
XM_004152677_Cucumis
AT1G47720_OSB1
XM_006304054_Capsella
XM_008380389_Malus
XM_020560676_Prunus
XM_021744397_Manihot
XM_024584552_Populus
XM_016713235_Capsicum
XM_016607981_Nicotiana
XM_016635538_Nicotiana
XM_023038590_Olea
XM_010686186_Beta
XM_022158089_Helianthus
XM_023872765_Lactuca
XM_028935356_Prosopis
XM_014767224_Glycine
XM_006595888_Glycine
XM_006852888_Amborella
XM_035965581_Zea
XM_015773207_Oryza
XM_018828604_Musa
OSB1
0.2

**C. SSB**

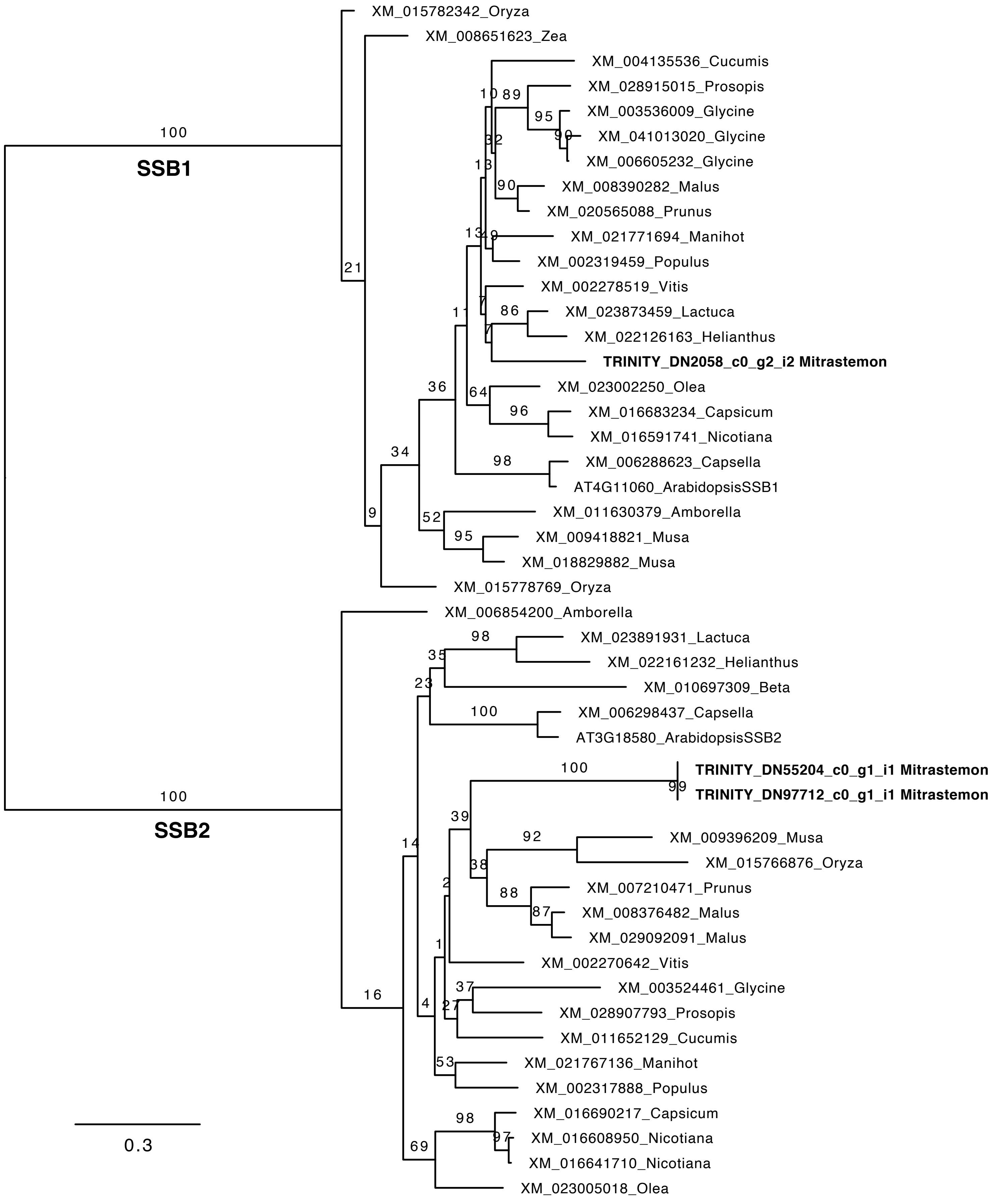
XM_015782342_Oryza
XM_008651623_Zea
XM_004135536_Cucumis
XM_028915015_Prosopis
XM_003536009_Glycine
XM_041013020_Glycine
XM_006605232_Glycine
XM_008390282_Malus
XM_020565088_Prunus
XM_021771694_Manihot
XM_002319459_Populus
XM_002278519_Vitis
XM_023873459_Lactuca
XM_022126163_Helianthus
TRINITY_DN2058_c0_g2_i2 Mitrastemon
XM_023002250_Olea
XM_016683234_Capsicum
XM_016591741_Nicotiana
XM_006288623_Capsella
AT4G11060_ArabidopsisSSB1
XM_011630379_Amborella
XM_009418821_Musa
XM_018829882_Musa
XM_015778769_Oryza
XM_006854200_Amborella
XM_023891931_Lactuca
XM_022161232_Helianthus
XM_010697309_Beta
XM_006298437_Capsella
AT3G18580_ArabidopsisSSB2
TRINITY_DN55204_c0_g1_i1 Mitrastemon
TRINITY_DN97712_c0_g1_i1 Mitrastemon
XM_009396209_Musa
XM_015766876_Oryza
XM_007210471_Prunus
XM_008376482_Malus
XM_029092091_Malus
XM_002270642_Vitis
XM_003524461_Glycine
XM_028907793_Prosopis
XM_011652129_Cucumis
XM_021767136_Manihot
XM_002317888_Populus
XM_016690217_Capsicum
XM_016608950_Nicotiana
XM_016641710_Nicotiana
XM_023005018_Olea
100
SSB1
100
SSB2
0.3

**C. Whirly**

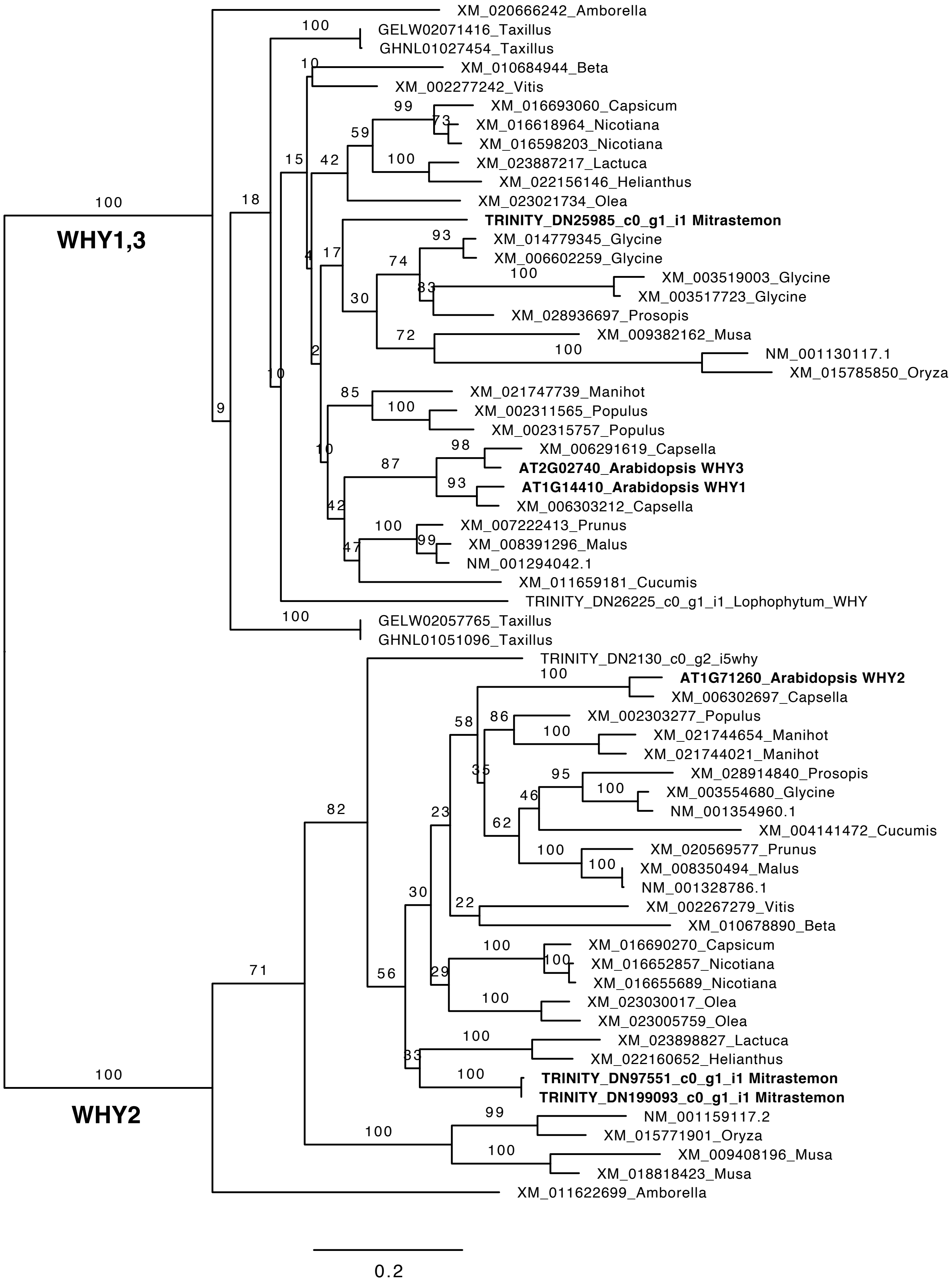
WHY1,3
XM_020666242_Amborella
GELW02071416_Taxillus
GHNL01027454_Taxillus
XM_010684944_Beta
XM_002277242_Vitis
XM_016693060_Capsicum
XM_016618964_Nicotiana
XM_016598203_Nicotiana
XM_023887217_Lactuca
XM_022156146_Helianthus
XM_023021734_Olea
TRINITY_DN25985_c0_g1_i1 Mitrastemon
XM_014779345_Glycine
XM_006602259_Glycine
XM_003519003_Glycine
XM_003517723_Glycine
XM_028936697_Prosopis
XM_009382162_Musa
NM_001130117.1
XM_015785850_Oryza
XM_021747739_Manihot
XM_002311565_Populus
XM_002315757_Populus
XM_006291619_Capsella
AT2G02740_Arabidopsis WHY3
AT1G14410_Arabidopsis WHY1
XM_006303212_Capsella
XM_007222413_Prunus
XM_008391296_Malus
NM_001294042.1
XM_011659181_Cucumis
TRINITY_DN26225_c0_g1_i1_Lophophytum_WHY
GELW02057765_Taxillus
GHNL01051096_Taxillus
WHY2
TRINITY_DN2130_c0_g2_i5why
AT1G71260_Arabidopsis WHY2
XM_006302697_Capsella
XM_002303277_Populus
XM_021744654_Manihot
XM_021744021_Manihot
XM_028914840_Prosopis
XM_003554680_Glycine
NM_001354960.1
XM_004141472_Cucumis
XM_020569577_Prunus
XM_008350494_Malus
NM_001328786.1
XM_002267279_Vitis
XM_010678890_Beta
XM_016690270_Capsicum
XM_016652857_Nicotiana
XM_016655689_Nicotiana
XM_023030017_Olea
XM_023005759_Olea
XM_023898827_Lactuca
XM_022160652_Helianthus
TRINITY_DN97551_c0_g1_i1 Mitrastemon
TRINITY_DN199093_c0_g1_i1 Mitrastemon
NM_001159117.2
XM_015771901_Oryza
XM_009408196_Musa
XM_018818423_Musa
XM_011622699_Amborella
0.2
